# Evolutionary assembly of crown reptile anatomy clarified by late Paleozoic relatives of Neodiapsida

**DOI:** 10.1101/2024.11.25.625221

**Authors:** Xavier A Jenkins, Roger BJ Benson, David P Ford, Claire Browning, Vincent Fernandez, Kathleen Dollman, Timothy Gomes, Elizabeth Griffiths, Jonah N Choiniere, Brandon R Peecook

## Abstract

Living reptiles include more than 20,000 species with disparate ecologies. Direct anatomical evidence from Neodiapsida, which includes the reptile crown-group and its closest extinct relatives, shows that this diversity originates from a single common ancestor that lived over 255 million years ago in the Paleozoic. However, the evolutionary assembly of crown reptile traits is poorly understood due to the lack of anatomically close relatives of Neodiapsida^1–7^. We present a substantially revised phylogenetic hypothesis, informed by new anatomical data from high-resolution synchrotron tomography of Paleozoic stem reptiles^8–9^. We find strong evidence placing the clade Millerettidae as the sister group to Neodiapsida, which uniquely share a suite of derived features. This grouping, for which we name the new clade Parapleurota, replaces previous phylogenetic paradigms by rendering the group Parareptilia as a polyphyletic assemblage of stem-reptiles, of which millerettids are the most crownward. Our findings address long-standing issues in Paleozoic reptile evolution^10–17^, such as firm support for the placement of captorhinids outside of crown Amniota and most varanopids as synapsids. These results greatly improve the fit of early amniote phylogeny to the observed stratigraphic record and reveal stepwise origin of crown reptile anatomy, including a middle Permian origin of tympanic hearing and loss of the lower temporal bar. This evolutionary framework provides a platform for investigating the diversification of the reptile crown group in the Early Triassic that was foundational to the origins of important living and extinct groups including dinosaurs (including birds), marine reptiles, crocodilians, and lepidosaurs.

## Introduction

Living reptiles include lepidosaurs, turtles, crocodilians, and birds^18^ within the reptile crown group, which exploded in diversity ∼250 million years ago in the Triassic Period. This important episode of diversification was preceded by the origin of important functional traits on the reptile stem lineage, whose timing has long been debated, including impedance-matching hearing^19–20^, temporal fenestration^10–11^, and the reorganization of the hindlimb musculature for tail-driven locomotion^12^. Relevant taxa informing the origins of these traits include the late Permian non-saurian neodiapsids such as terrestrial younginids^21^, semiaquatic tangasaurids^22^, and gliding weigeltisaurids^23^. Close neodiapsid outgroups would also greatly increase our leverage for understanding these anatomical transformations but have remained almost completely unknown for much of the Permian, causing major uncertainties about the evolutionary assembly of crown reptile anatomy^2^.

Prevailing hypotheses of Paleozoic reptile evolution^5–7^ generally posit a deep phylogenetic split between ‘near reptiles’ (Parareptilia; an ecomorphologically diverse group that is thought to be only distantly related to crown reptiles), and ‘true reptiles’ (Eureptilia; an assemblage of taxa interpreted as successively closer sister-groups to crown reptiles including Neodiapsida, and including the Late Carboniferous *Hylonomus* (∼318 Ma)) (Fig. 1). The origin of this dichotomy between Parareptilia and Eureptilia lies in the foundational work of Olson^14^, who allied the Seymouriamorpha and Diadectomorpha with Procolophonia and Chelonia (turtles) in the class ‘Parareptilia’, and the Captorhinomorpha, Synapsida (mammal-line amniotes), and fenestrated reptiles in the class ‘Eureptilia’. This hypothesis was largely ignored by Watson^8^, Romer^13^, and Carroll^15–16^ who considered seymouriamorphs and diadectomorphs as earlier-diverging tetrapods more distantly related to amniotes, whereas captorhinomorphs were thought to be either close relatives of Synapsida^8,13^ or outside of crown Amniota. Although Olson’s original hypothesis of a monophyletic Parareptilia containing diadectomorphs and a Eureptilia containing Synapsida has not found support since the advent of cladistics^1–7,24–28^, the concept of a parareptilian group that includes Procolophonia with various other taxa has nevertheless persisted.

**Figure 1.**
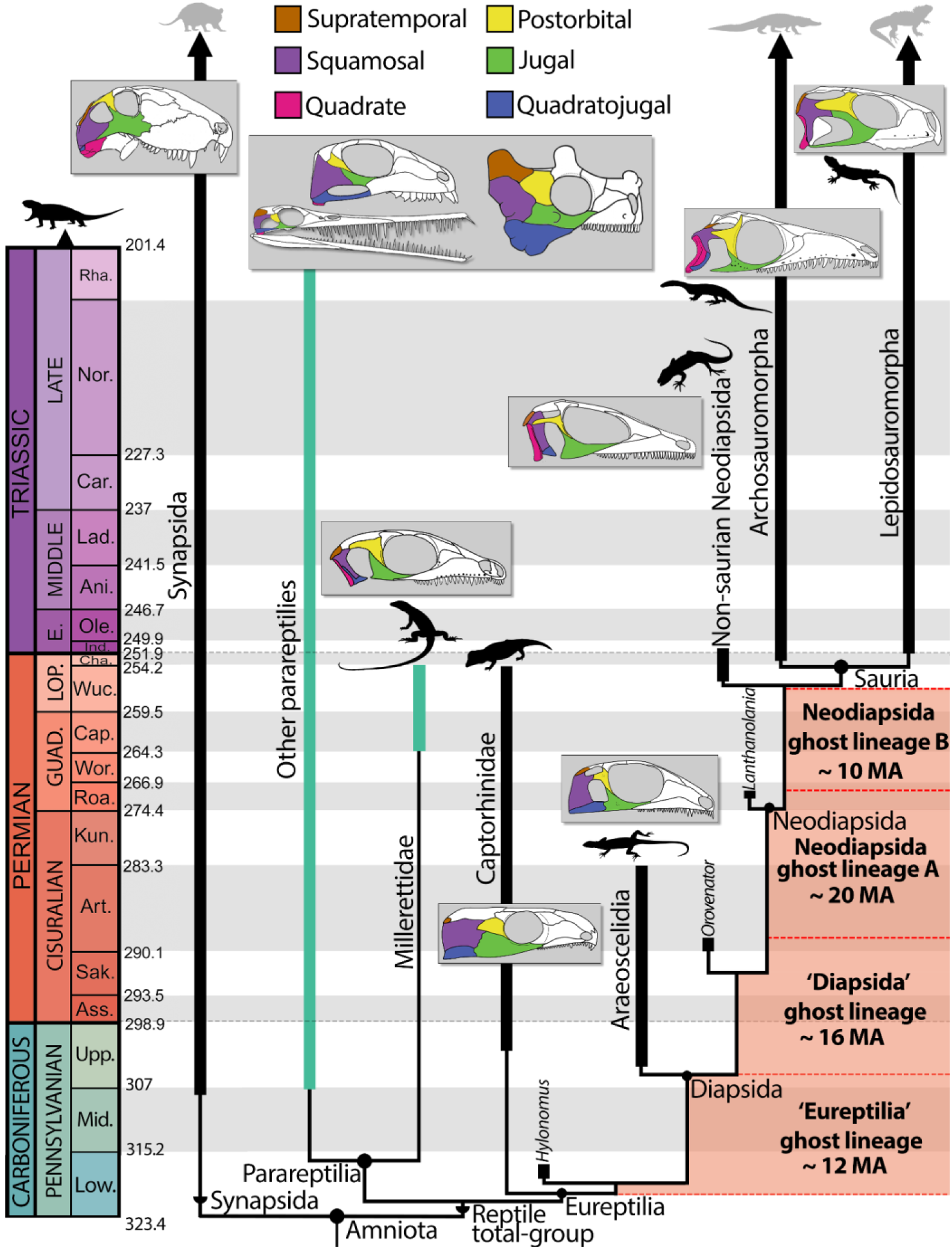
Traditional phylogenetic hypothesis of Late Carboniferous-Triassic amniote relationships, highlighting the long-standing stem-reptile ghost lineages from the Late Carboniferous through late Permian. Green branches represent ‘Parareptilia’. Diagrams of amniote fenestration are not to scale and have been redrawn based on the following reconstructions (left to right): *Dimetrodon*^75^, *Mesosaurus*^74^, *Belebey*^29^, *Milleropsis*^44^, *Bunostegos*^104^, *Captorhinus*^105^, EAZ younginiform (this study), *Petrolacosaurus*^36^, *Prolacerta*^107^, and *Clevosaurus*^108^. Silhouettes are available from Phylopic (www.phylopic.org) under CC BY 3.0 licenses or within public domain.

Early cladistic analyses on amniote origins opted to use ‘Parareptilia’ for a group consisting of Procolophonia and Millerettidae, and ‘Eureptilia’ for a group containing captorhinids, protorothyridids, and diapsid (two temporal fenestrae) reptiles^5^. Soon after, the late

Carboniferous and Permian groups Acleistorhinidae^6^ and Bolosauridae^29^ were also placed within Parareptilia. Tsuji and Müller^30^ proposed the following definition for Eureptilia: ‘the most inclusive clade containing *Captorhinus aguti* and *Petrolacosaurus kansensis* but not *Procolophon trigoniceps*’, and Parareptilia as: ‘the most inclusive clade containing *Milleretta rubidgei* and *Procolophon trigoniceps* but not *Captorhinus aguti*’. This framework has been paradigmatic for studies of early reptile evolution since the early 2000s.

The composition of Parareptilia and Eureptilia under these long-standing definitions has recently been challenged by phylogenetic analyses from multiple working groups, with the most comprehensive studies finding former members of Eureptilia either as stem-amniotes (captorhinids)^24–25^ or the among the earliest-branching stem reptiles (araeoscelidians)^4,26–28^ and members of Parareptilia as a grade of stem-reptiles closer to the reptile crown group^24–25^.

However, substantial uncertainties remain regarding the affinities of some groups to the reptile or even the mammal stem-lineage^4^ and whether some taxa long-regarded as stem-reptiles instead belong to the amniote stem-lineage^24–25^. For example, Varanopidae and Recumbirostra – two groups often placed on the amphibian, amniote, or mammal stem-lineages – have recently been interpreted as stem reptiles^4,31^ (but see^26,32^). Similarly, the ‘Romeriidae’ were originally thought to be ancestral to amniotes^15–16^ and were later reassigned to the reptile stem in the 1980s^7^, but recent studies once again recover them as stem amniotes^24–25^. Challenges to this phylogenetic paradigm of Eureptilia and Parareptilia have yet to achieve consensus^4, 25–26^.

The paradigm of a deep phylogenetic split between Parareptilia and Eureptilia implies the existence of extensive ghost lineages along the reptile stem lineage^33^, particularly for neodiapsid origins^2^ (Fig. 1). These include a ghost lineage for Eureptilia between the appearance of the earliest candidate eureptile *Hylo nomus*^15,34^ (∼318 Ma) and other early reptiles towards the end of the Carboniferous^35–36^ (∼305 Ma) (Fig. 1, ‘Eureptilia’ ghost lineage). More central to the question of crown reptile origins, however, are several ghost lineages in the sporadic fossil record of non-saurian neodiapsids, which are only partially closed by the putative neodiapsids *Orovenator mayorum* (early Permian, ∼289 MA^37^) and *Lanthanolania ivakhnenkoi* (middle Permian, ∼265 MA^38^), both genera represented only by partial skulls (Fig. 1A, B; ‘Diapsida ghost lineage’” and ‘Neodiapsida ghost lineage’ A and B). Within Parareptilia, the phylogenetic position of Millerettidae^9^ as the sister group of a group formed by most or all other parareptiles further exemplifies the poor stratigraphic fit of current phylogenetic hypotheses, millerettids are the geologically youngest group of nominal parareptiles to appear, leading to an expansive basal ghost lineage that spans nearly the entire Permian^33^ (Fig. 1).

The poor stratigraphic fit of these current hypotheses of early reptile evolution makes it difficult to date the series of anatomical transformations involved in the origins of crown reptile anatomy, particularly given the dearth of fossil stem-reptiles bridging the morphological gap between early araeoscelidians in the late Carboniferous^35^ and the first adequately known neodiapsids in the later Permian^21^ (Fig. 1). For instance, key neodiapsid postcranial traits such as an ossified sternum, laterally-directed caudal ribs supporting the caudofemoralis musculature, and the appearance of an outer process on the fifth metatarsal suggesting a restructuring of the m. gastrocnemius femoralis are first documented in late Permian non-saurian neodiapsids such as *Youngina*^21^. Late Permian neodiapsids also contrast with araeoscelidians in the loss of amniote plesiomorphies such as a trough on the dorsal surface of the iliac blade, asymmetrical femoral condyles and the cnemial crest of the tibia^4^–features which are also lost among some groups of middle and late Permian parareptiles, such as millerettids^9^.

Early Permian reptiles relevant to this issue remain stubbornly absent from the fossil record, but their anatomical features could be inferred from geologically younger taxa given robust hypotheses for their phylogenetic relationships. One group of putative stem-reptiles in particular hold latent information that could potentially inform early reptile relationships: crown-reptile-like millerettid parareptiles (Figs. 2–3). Prior to the advent of cladistic methodologies both Watson^8^ and Romer^13^ interpreted millerettids as close relatives of the ‘Eosuchia’, a grade loosely equivalent to early Neodiapsida in current terminology. These historical observations were based on anatomical features suggesting the loss of an upper temporal fenestra and the presence of a tympanic emargination located posteriorly on the skull. Other studies even hypothesized that these derived anatomical features of millerettids indicated a placement within the reptile crown, either on the archosaur ^39^ or on the lepidosaur^9^ stem lineage. However, current phylogenetic hypotheses^4,24^ usually interpret these similarities as homoplasies. New data on Permian taxa such as millerettids therefore have potential to influence phylogenetic hypotheses, with implications for stem reptile relationships and the evolutionary assembly of crown reptile adaptations.

**Figure 2.**
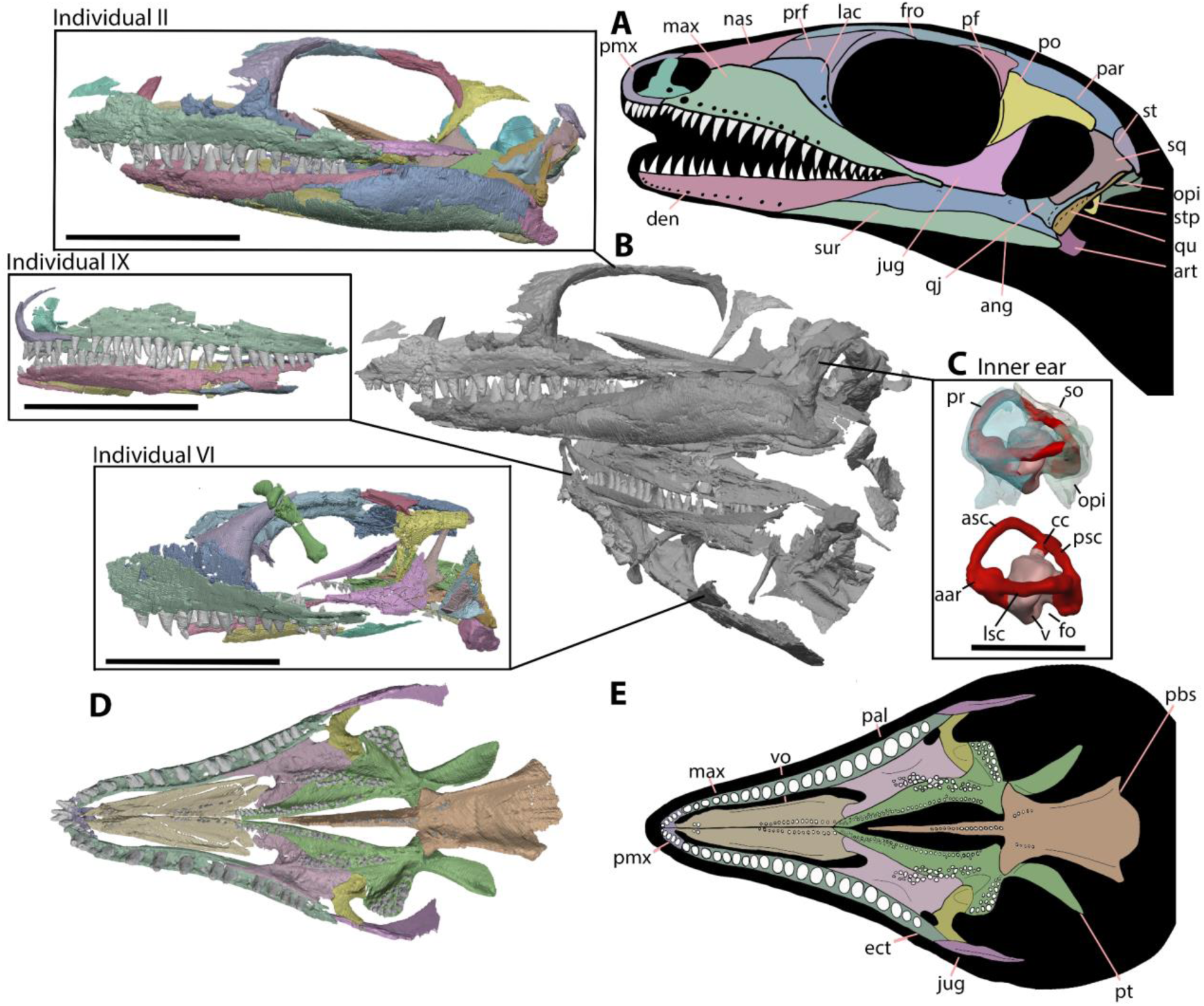
New information and individuals of *Milleropsis* presented here. **A**, line reconstruction of the cranial anatomy of *Milleropsis pricei* BP/1/720. **B**, segmentation of all three individuals (following the numbering of ref. 8) of *Milleropsis* present in BP/1/720, individual II, IX (mirrored), and VI (mirrored), in left lateral views **C,** segmented endosseous labyrinth of *Milleropsis* BP/1/720 individual II in left lateral view with bones of the otic capsules translucent and removed (lagena not reconstructed), **D**, virtual reconstruction of the palate of Individual II (left ectopterygoid mirrored), and **E**, line reconstruction of Individual II in palatal view. Scale bars for skulls are 10 mm whereas inner ear scale bar is 5 mm. Abbreviations: aar, anterior ampullary recess; asc, anterior semicircular canal; ang, angular; art, articular; cc, common crus; den, dentary; ect, ectopterygoid; fro, frontal; fo, fenestra ovalis; jug, jugal; lac, lacrimal; lsc, lateral semicircular canal; max, maxilla; nas, nasal; opi, opisthotic; pal, palatine; par, parietal; pbs, parabasiphenoid; pf, postfrontal; pmx, premaxilla; po, postorbital; pr, prootic; prf, prefrontal; psc, posterior semicircular canal; pt, pterygoid; qj, quadratojugal; qu, quadrate; so, supraoccipital; sq, squamosal; st, supratemporal; stp, stapes; sur, surangular; v, vestibule; and vo, vomer.

**Figure 3.**
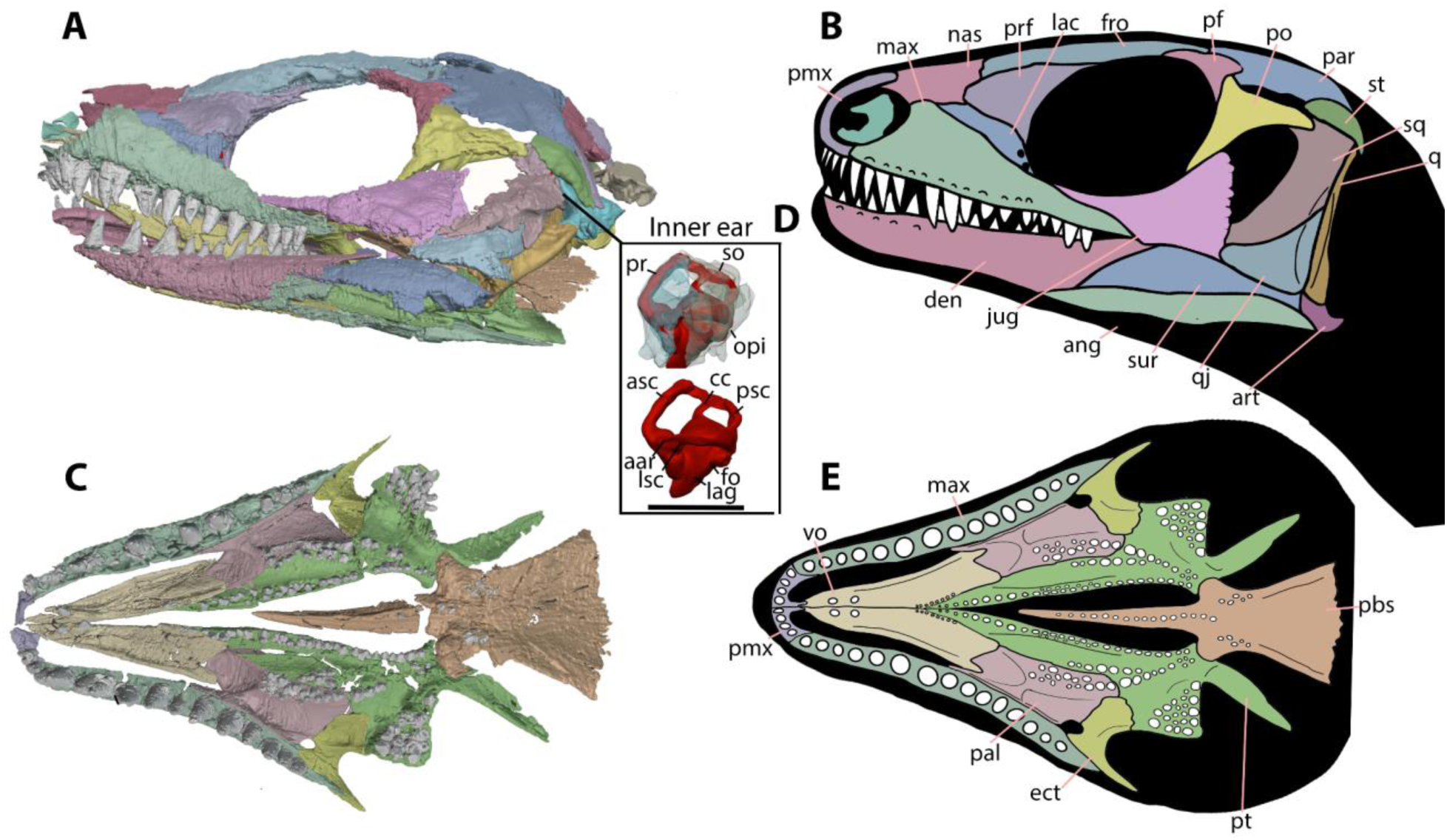
New information on *Milleretta* presented here. See the main text for the discussion of the temporal fenestrae in *Milleretta*, which are secondarily closed during ontogeny. **A**, segmentation of BP/1/3822 in mirrored left lateral view and **B,** corresponding line reconstruction. **C**, segmentation of BP/1/3822 in palatal view; **D**, segmented endosseous labyrinth of *Milleretta* BP/1/3822 in left lateral view with bones of the otic capsules translucent and removed; and **E**, reconstruction of *Milleretta* BP/1/3822 in palatal view. Scale bar for skulls is 10 mm whereas the inner ear scale bar is 5mm. Abbreviations: aar, anterior ampullary recess; asc anterior semicircular canal; ang, angular; art, articular; cc, common crus; den, dentary; ect, ectopterygoid; fro, frontal; fo, fenestra ovalis; jug, jugal; lac, lacrimal; lag, lagena; lsc, lateral semicircular canal; max, maxilla; nas, nasal; opi, opisthotic; pal, palatine; par, parietal; pbs, parabasiphenoid; pf, postfrontal; pmx, premaxilla; po, postorbital; pr, prootic; prf, prefrontal; psc, posterior semicircular canal; pt, pterygoid; qj, quadratojugal; qu, quadrate; so, supraoccipital; st, supratemporal; sq, squamosal; sur, surangular; and vo, vomer.

We reevaluate the evolutionary relationships of early reptiles by conducting an extensive phylogenetic revision of Paleozoic reptiles, informed by new computed X-ray micro tomography (μCT) data, including high-resolution synchrotron phase-contrast μCT that enables us to collect and analyze previously unobservable anatomy. Our phylogenetic dataset improves on previous datasets by broadly sampling all putative stem-reptiles from the Late Carboniferous (Pennsylvanian) through the explosive radiation of the reptile crown in the Triassic, and by incorporating unsampled character-rich systems (e.g., neurocranium and palatoquadrate) in key taxa. Our new tomography data includes putative stem-reptiles such as the varanopid

*Heleosaurus*^40^, the araeoscelidian *Araeoscelis*^41^, the millerettids *Milleropsis*^8^ and *Milleretta*^9^, and neodiapsids, including *Youngina*^42^ and the *Endothiodon* Assemblage Zone (EAZ) younginiform^43^ SAM-PK-K7710, as well as other taxa across early amniote phylogeny (Table 1). We placed a special focus on millerettids, which share previously underappreciated similarities with crown reptiles^44–46^ (in modern cladistic frameworks) and which substantiate our understanding of anatomical transformations along the reptile stem lineage (Figs. 2–3, Videos S1–3).

**Table 1.**
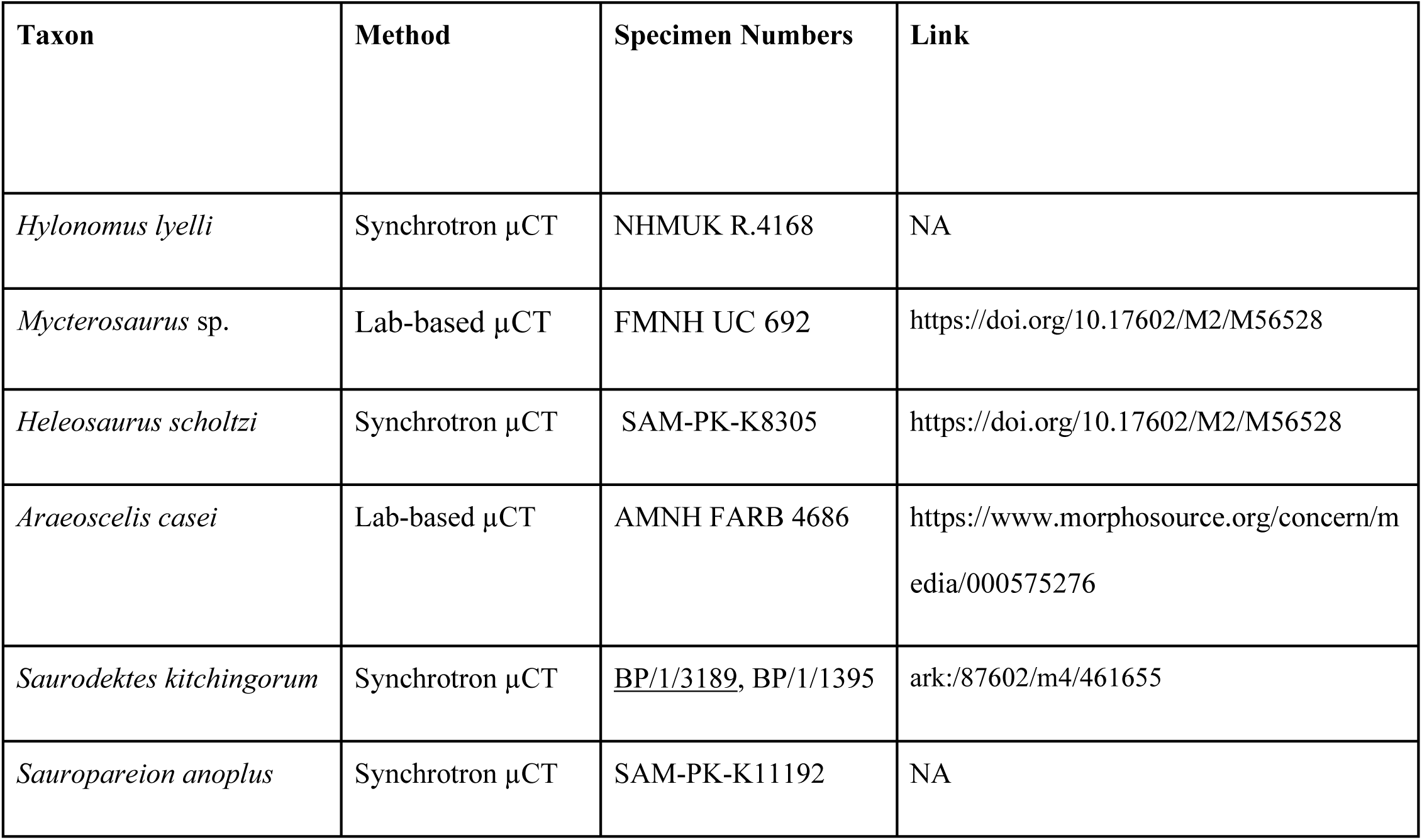
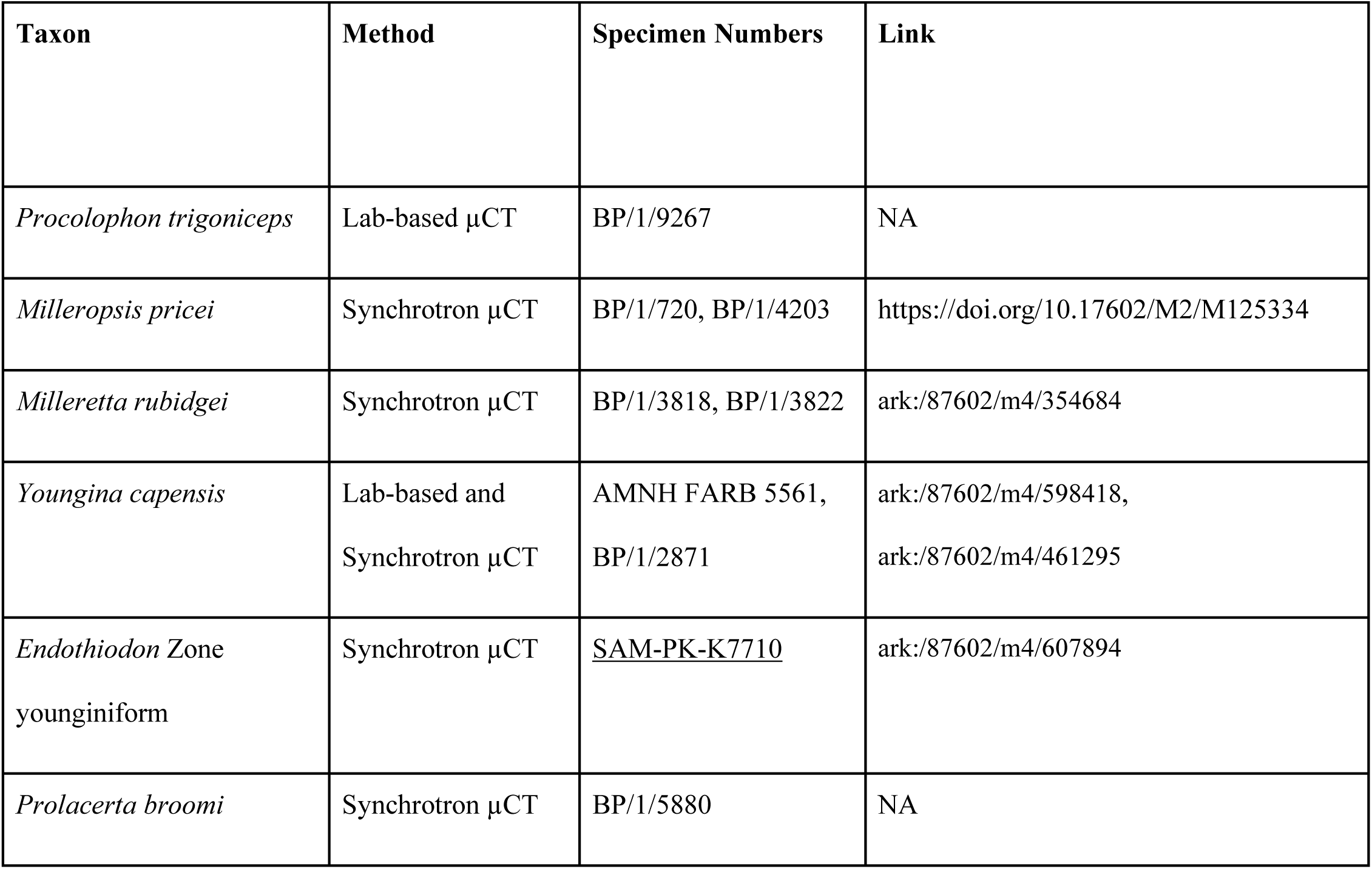
Scan data of Paleozoic and Triassic amniotes that are accessible and under current study by one or more of the authors. Links are provided when appropriate.

## Methods

### Computed Tomography

BP/1/720 (*Milleropsis*), BP/1/3822 (*Milleretta*), SAM-PK-K7710 (a non saurian-neodiapsid), and other early amniotes (Table 1) were imaged at the ID19 beamline of the European Synchrotron Radiation Facility (Grenoble, France) using propagation phase-contrast synchrotron X-ray microtomography. The imaged material of BP/1/720 was of a nodule containing three skulls: individuals II, VI, and IX^8^. BP/1/4203, a fully articulated specimen of *Milleropsis*, was imaged onsite at the University of the Witwatersrand. The holotypes of *Youngina capensis* (AMNH FARB 5561) and *Araeoscelis gracilis* (AMNH FARB 4686) were imaged onsite at the American Museum of Natural History. The μCT data were processed and segmented using Dragonfly 2022.268 (https://dragonfly.comet.tech/)^46^.

### Phylogenetic Analysis

We evaluated the phylogenetic relationships of Paleozoic-Triassic reptiles by creating a new phylogenetic dataset using substantial new anatomical data from our scans of millerettids, neodiapsids, and other Permo-Triassic amniotes (Table 1). Our matrix (total characters = 647) includes comprehensive rescoring of skeletal anatomy informative for higher-level reptile relationships, particularly for millerettid and non-saurian neodiapsids (Table 1).

We expanded on previous datasets^4,26^ of early amniote relationships by including 259 additional characters not used in previous analyses of the phylogeny of stem-reptiles, which substantially improved representation of the neurocranium and postcranial skeleton of early amniotes. We attempted to incorporate all potentially relevant phylogenetic characters based on a comprehensive review of existing phylogenetic matrices that sample stem-amniote^31^, parareptile^47^ and crown reptile relationships^2,3^, surveys of the descriptive literature, and observations of the early amniote neurocranium and postcranium in collections. These include features that were previously inaccessible in many taxa for which CT data were not available, including the detailed morphology of the orbit, tympanic recess, palatoquadrate, braincase and mandible. We illustrate a fraction of this anatomical variation in these regions using *Captorhinus*, *Heleosaurus*, *Saurodektes*, *Milleropsis*, and the EAZ younginiform as representative taxa (Figs. 4–8).

**Figure 4.**
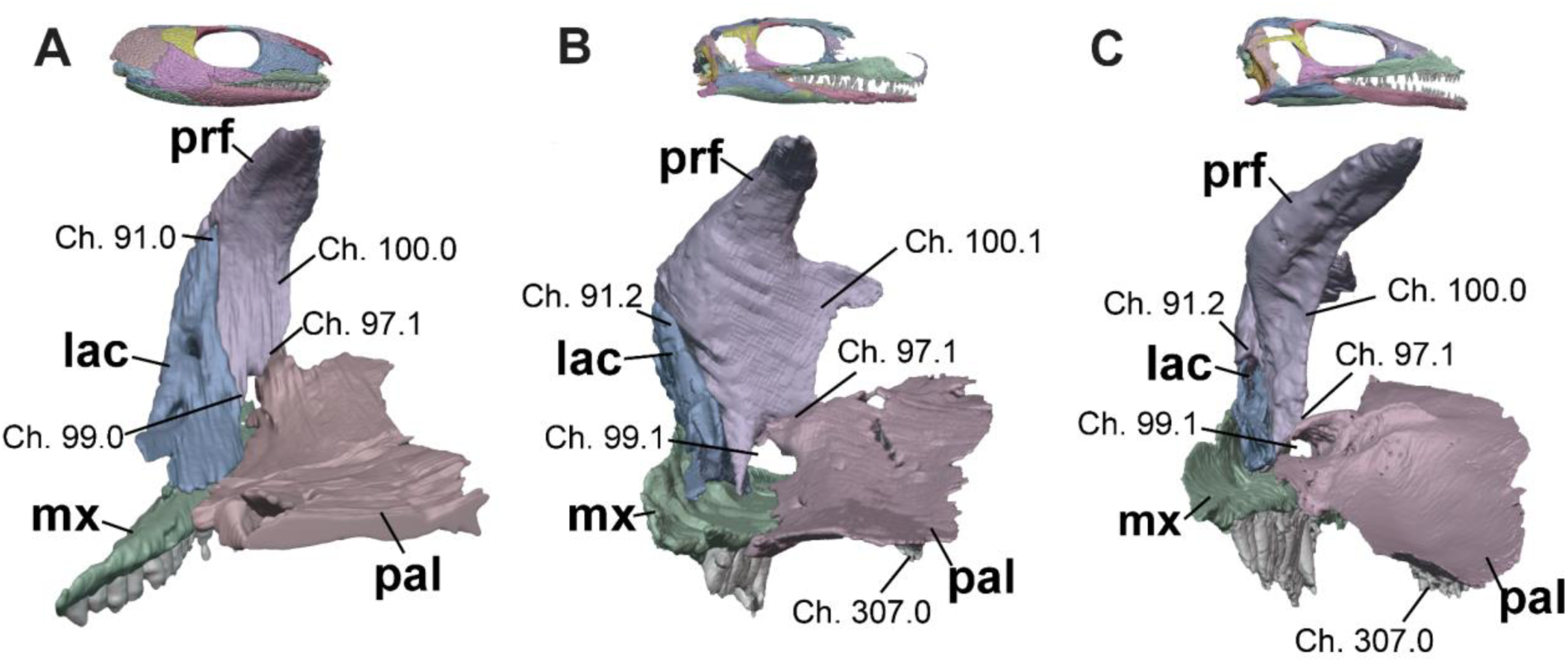
Anterior orbital walls in posterior view of **A**, *Captorhinus* (OMNH 44816)^91^, **B**, *Milleropsis* (BP/1/720), and **C**, EAZ younginiform (SAM-PK-K7710) demonstrating select characters informative to early amniote evolution, including: the contribution of the lacrimal to the orbit (**Ch. 91**, mediolaterally broad in near-amniotes and early amniotes), prefrontal-palatine contact* (**Ch. 97**, present in most taxa within this sampling), the presence of a foramen orbitonasale between the prefrontal and palatine* **(Ch. 99.1,** a contribution to this foramen by the lacrimal is present in captorhinids), the width of the ventral process of the prefrontal*, (**Ch. 100**, wide in edaphosaurids, procolophonians, millerettids, and rhynchosaurs), and the presence of palatine dentition (**Ch. 307**, absent in some procolophonians and archosauromorphs). * Features considered autapomorphies of Parareptilia in the literature but in fact widespread within and around Amniota. Abbreviations: lac, lacrimal; mx, maxilla; pal, palatine; and prf, prefrontal.

**Figure 8.**
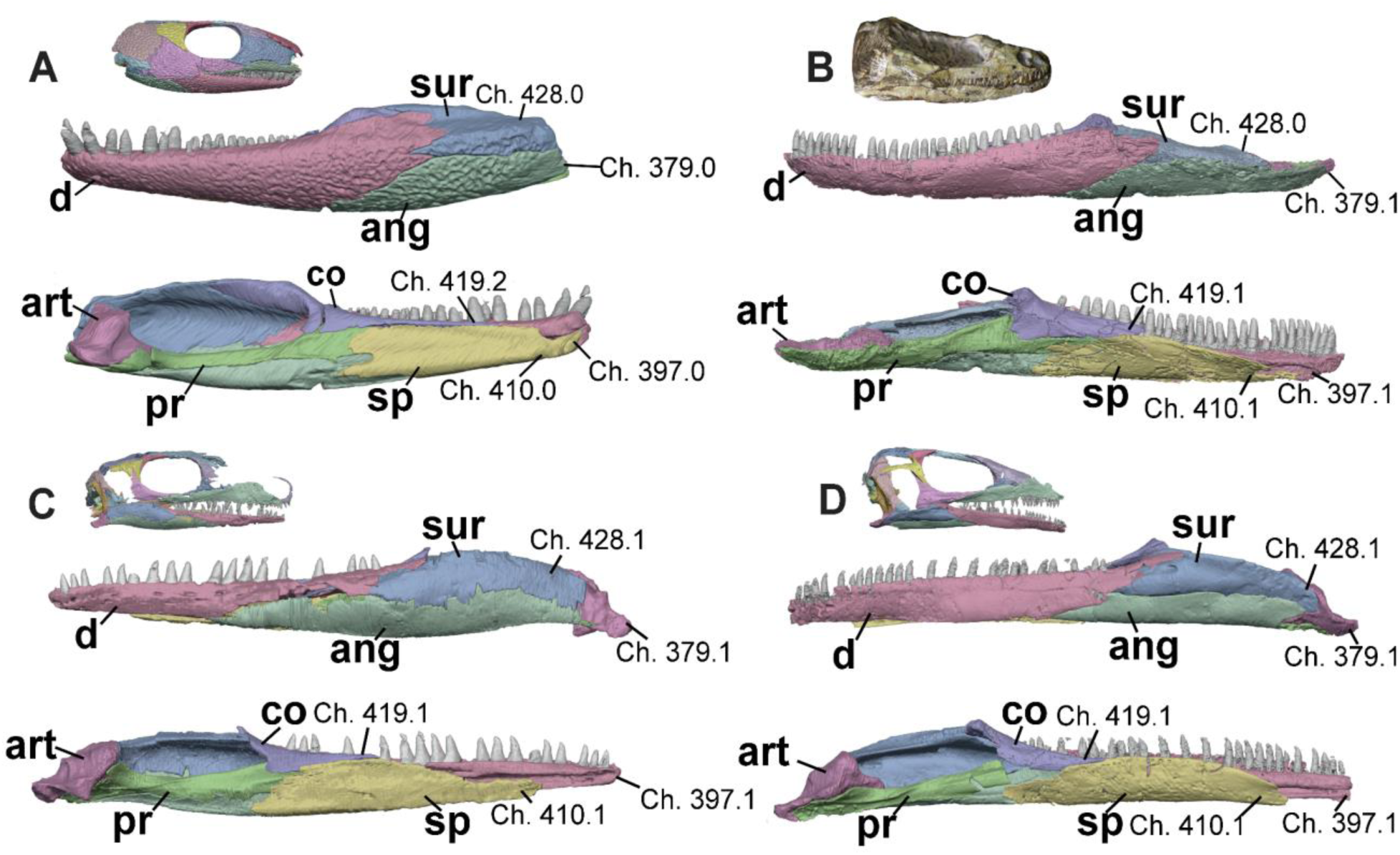
Left mandibles of **A**, *Captorhinus* (OMNH 44816)^91^, **B,** *Saurodektes* (BP/1/1395), **C**, *Milleropsis* (BP/1/720), and **D**, EAZ younginiform (SAM-PK-K7710) in lateral (upper) and medial (lower) views, emphasizing the morphology of the symphysis and articular region. Select characters include: the presence of a retroarticular process (**Ch. 379,** variably absent in this sample), the pathway of the Meckelian canal through the anterior mandible (**Ch. 397**, piercing the dentary in total-group reptiles), the splenial contribution to the mandibular symphysis (**Ch. 410,** absent in most stem-reptiles but present in other amniotes and convergently in pareiasaurs and rhynchosaurs within this sample), the length of the coronoid (**Ch. 419**, extremely long in captorhinids, which may be the result of a fusion of two coronoids), and the presence of a posterior surangular foramen (**Ch. 428**, present in early parapleurotans). Abbreviations: ang, angular; art, articular; co, coronoid; d, dentary; pr, prearticular; sa, surangular; and sp, splenial.

We also improved the taxonomic sampling of this matrix by adding 68 taxa typically not included in analyses of reptile origins, including 13 recumbirostrans, 14 parareptiles, and 22 neodiapsids including crown reptiles bringing the total to 167 operational taxonomic units (OTUs). Notably, our dataset is the first to sample all adequately known millerettids, including *Broomia perplexa*, *Milleropsis pricei*, *Millerosaurus nuffieldi*, and *Milleretta rubidgei*, with all known early parareptile groups and the early members of Neodiapsida. We did not include candidate stem turtles such as *Eunotosaurus africanus*^47^ in our analyses because resolving key uncertainties in this area requires additional focused study. Instead, our determination of the reptile crown group relies on the assumption that turtles are the sister to archosaurs among living groups, which is well-supported by numerous molecular phylogenetic studies^48–49^. The full character and taxon list (with annotations on their use in previous analyses), optimizations, and rationale for their inclusion are provided in the SI Appendix along with the character-taxon matrix.

### Parsimony Analysis

Parsimony analysis was carried out in TNT v1.6^51^ using driven-search implementations of the “new technology” heuristic strategies with sectorial, drift and tree fusing algorithms following the methods of^4^. *Whatcheeria deltae* was set as the outgroup taxon, and a total of 80 characters were ordered. The resulting 48 most parsimonious trees (MPTs) of 4751 steps were then subjected to a final round of tree bisection and reconnection branch swapping, with a 50% collapsing rule. Branch support was calculated using Bremer support and bootstrap resampling analyses, with 1,000 pseudoreplicates for absolute and group present/contradicted frequencies, also calculated in TNT v.1.6^52^ (Fig. S3). To reconstruct the ACCTRAN/DELTRAN optimizations for each character, we conducted a heuristic search with tree bisection and reconnection, with 1,000 random additional replicates in PAUP* 4.0a169^50^, resulting in 48 MPTs (identical to those found in TNT) with 4751 steps. The ACCTRAN/DELTRAN optimizations for each new character are provided in SI Appendix 8.

We ran an additional parsimony analysis following the methods above (41 MPTs of branch length = 4710) in which seven characters (118, 122, 133, 134, 155, 172, 177) regarding temporal fenestration and emargination were removed to assess the influence of patterns of temporal fenestration on amniote relationships (Fig. S3). We also computed a series of constraint analyses (analyses C1–C6) under maximum parsimony in PAUP* 4.0a169^52^ to assess differences in tree topology, length, and resolution when imposing backbone constraints mimicking previous phylogenetic hypotheses of early reptile evolution^2–6^. Analysis C1 constrains a monophyletic Eureptilia including Captorhinidae, Araeoscelidia, and Neodiapsida following the hypotheses of^5–7^. Analysis C2 specifically excludes Millerettidae from Eureptilia. Analysis C3 constrains the monophyly of Parareptilia encompassing millerettids,bolosaurids, acleistorhinids, and procolophonians to the exclusion of all eureptiles^5,6,50^. Analysis C4 constrains araeoscelidians as the sister taxon to Neodiapsida so they become diapsids according to the apomorphy-based definition of Diapsida^53^, and analysis C5 excludes Millerettidae from Diapsida. Lastly, analysis C6 constrains Recumbirostra within the reptile total-group following several recent hypotheses of a reptilian placement of this group^31,54^.

### Bayesian Inference

Bayesian phylogenetic inference was carried out in MrBayes 3.2.7a using a fossilized birth–death (FBD) tree prior^55–56^. The ages of all OTUs were specified using uniform distributions spanning the minimum and maximum first occurrence dates of each OTU. The setting for the tree age prior was set to offsetexponential (x,y), where x is the oldest fossil in the taxon list, *Whatcheeria deltae* (347.6 Ma), and y the earliest putative date of the hypothetical common ancestor, in this case, the age of *Tiktaalik* (372 Ma). Two runs, each consisting of four chains at a temperature of 09, were performed for 100 million generations, sampling every 10,000 generations with a burn-in of 50%. We performed a second, non-time-calibrated Bayesian analysis using the MkV model with a ‘standard’ datatype, coding set at ‘variable’ and with rates set to a gamma distribution, which allows the MkV substitution model to allow variation in the evolutionary rates among characters. The analysis was run for 100 million generations, sampling every 10,000^th^ generation, with a burn-in of the first 10,000 sampled trees (50%). The effective sample size for tree length and shape of gamma distribution was greater than 3,900 and the average potential scale reduction factor was 1.01 or less on all parameters, indicating convergence. Our analytical scripts and resulting tree files are available at https://osf.io/9jac3/.

### Stratigraphic Fit

To compare the stratigraphic fit of our new phylogenetic hypotheses to those of previous hypotheses for early reptile relationships, we computed the modified Manhattan stratigraphic measure (MSM*)^57^ and gap excess ratio (GER*)^58^ stratigraphic congruence analyses. These measurements use the sum of the ghost ranges (or MIG, minimum implied gap) relative to the optimal and/or G_min_ (in the case of MSM*) and/or maximally suboptimal fits to stratigraphy (GER*). These measures account for stratigraphic uncertainty by using a resampling technique over 1000 iterations that incorporate possible ranges of a given taxon using the minimum (LAD) and maximum ages (FAD)^54^.

To further investigate the stratigraphic consistency of our phylogenetic hypothesis relative to more ‘traditional’ hypotheses of early amniote relationships, we calculated the MSM*Diff (the differences in MSM* values) using the 48 MPTs from our parsimony analysis and 62 MPTs of a constrained version of our dataset that enforces the monophyly of Eureptilia, Parareptilia, and Diapsida. This constrained version therefore approximates the four stem-reptile ghost lineages in prevailing phylogenetic hypotheses of early reptiles over the last three decades^5–7^. Metrics for stratigraphic fit to phylogeny are not comparable across different taxon sets, and for this reason we could not directly compare our preferred phylogenetic hypothesis to those of previous workers^57^. However, our constraint trees used here approximate the general relationships hypothesized in previous work. For comparative purposes, we calculated measures of stratigraphic fit (but not MSM*Diff due to differences in taxonomic sampling in these matrices and ours) for two^24,5^ recent matrices on early reptile evolution.

### Ancestral-State Reconstruction

Ancestral-state reconstruction for the presence and morphological variation of the tympanic fossa and upper and lower temporal fenestrae were carried out in R using the phytools package^60–61^ and its dependencies using the function *fitMk*, which fits a continuous time-reversible Markov model to simulate character history. We estimated the ancestral states of tympanic emarginations and temporal fenestration in four discrete analyses: testing the presence and absence of a tympanic emargination (two characters), the anatomy and location of the tympanic fossa (six characters), the presence and morphology of a lower temporal opening (five characters), and the presence or absence of an upper temporal fenestra (two characters). When the anatomy of tympanic emarginations or temporal fenestration is unknown in certain taxa (e.g., taxa in which the skull is absent or only partially preserved), the taxa were treated as uncertain following the methods of^60–61^. For each analysis, we fit two different models of evolution onto our time-calibrated FBD tree (Fig. S4): “ER”, the equal rates model, and “ARD”, the all-rates-different model, and compare these two models using the ‘ANOVA’ function. The posterior probabilities at each ancestral node in our FBD tree were calculated and plotted for ancestral-state reconstruction of tympanic emargination (Fig. 11) and temporal fenestration (Fig. 12).

**Figure 11.**
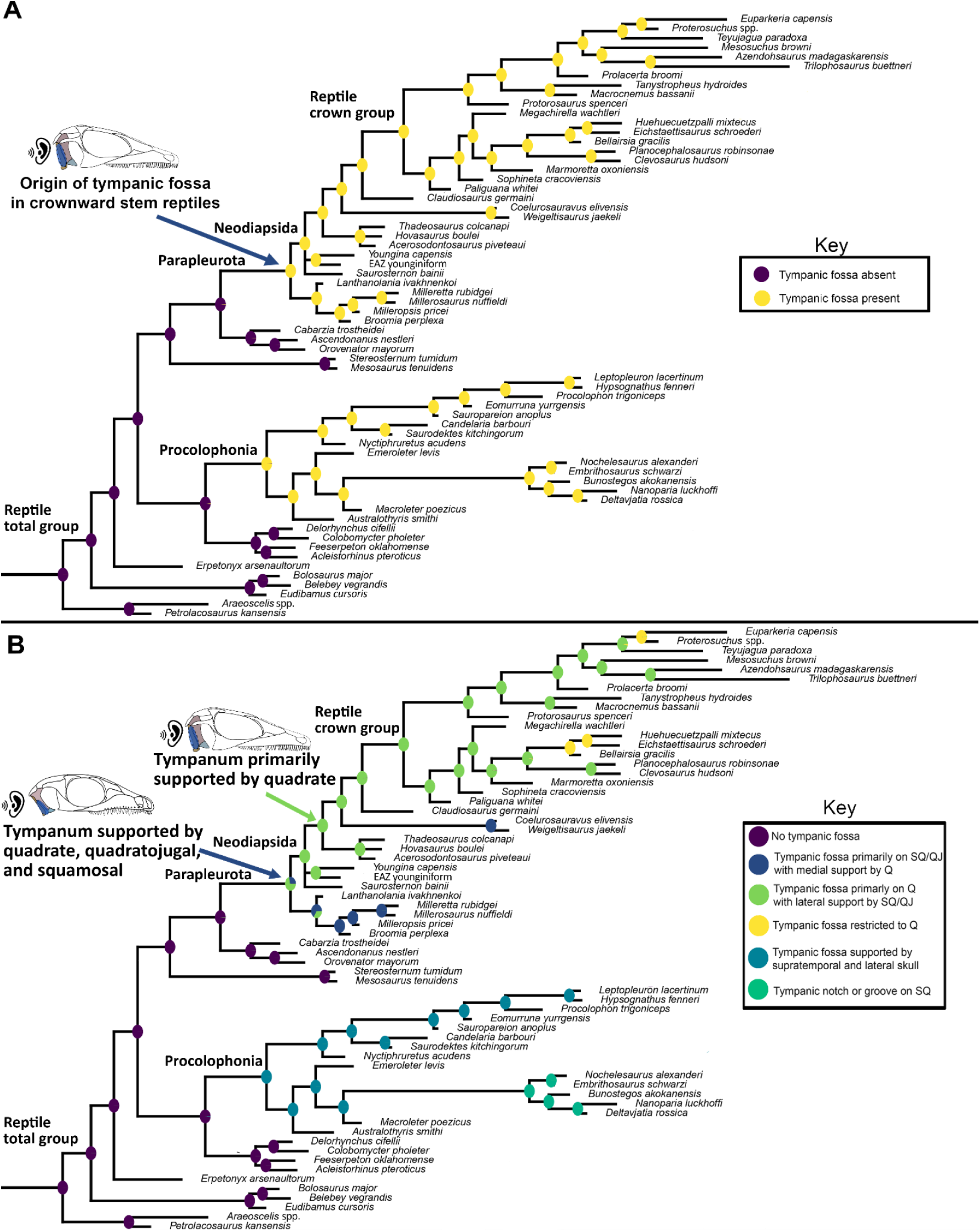
Ancestral-state reconstruction estimates for **A**) the presence or absence of a tympanic emargination; and **B)** the anatomy of the tympanic emargination. A tympanic emargination is reconstructed as having evolved convergently in parapleurotans (supported on the quadrate/squamosal/quadratojugal) and in procolophonians (squamosal/quadratojugal/supratemporal), with a stepwise increase in the contribution of the quadrate to the tympanic emargination leading towards crown reptiles.

**Figure 12.**
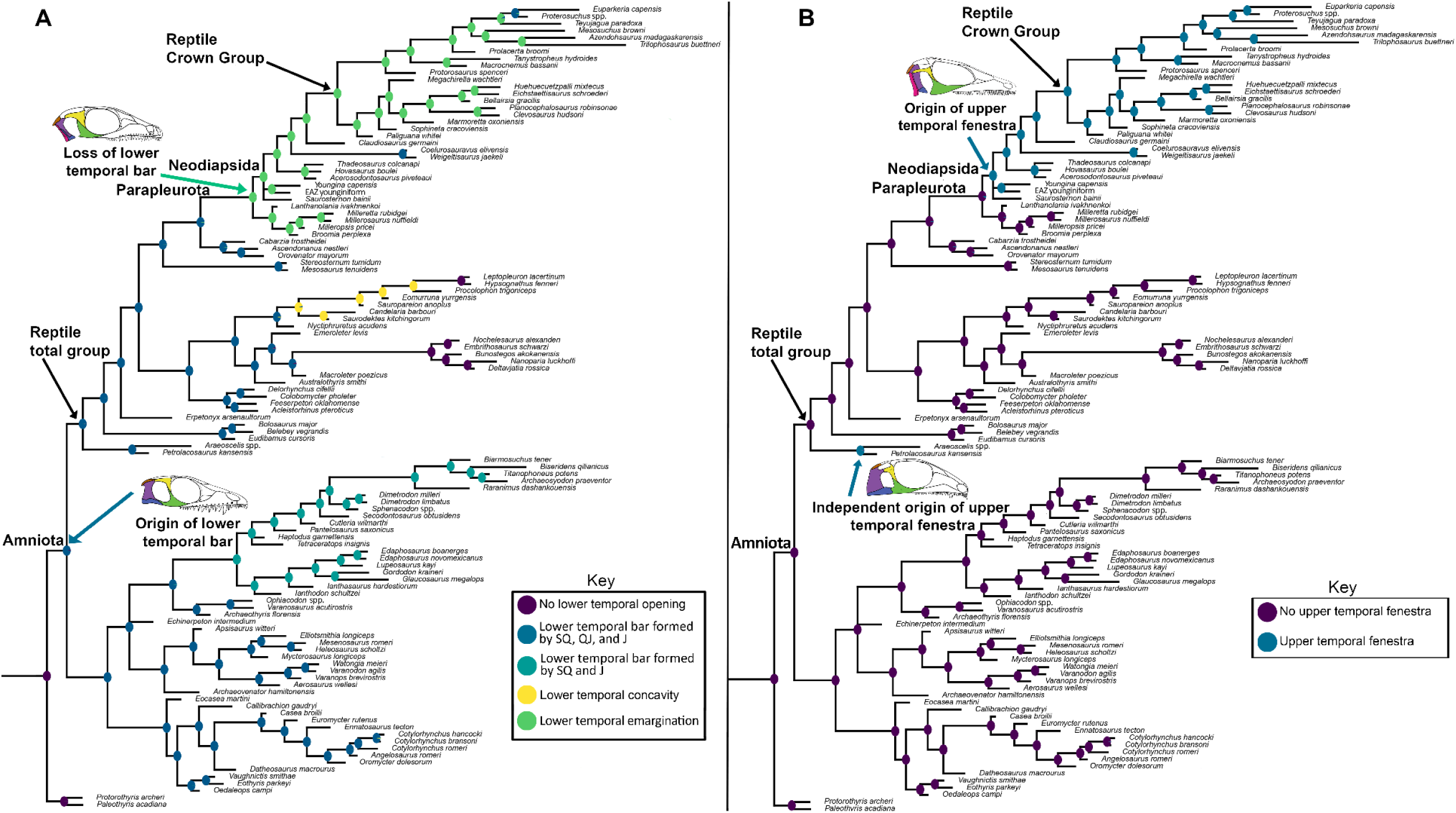
Ancestral-state reconstruction estimates for **A**) the presence and morphology of the lower temporal fenestra/opening and **B)** the presence of the upper temporal fenestra. The amniote common ancestor is reconstructed as fenestrated, with the ‘synapsid’ or a single lower temporal fenestration as the plesiomorphic condition. Modifications to this lower temporal condition occur independently in araeoscelidians + bolosaurids, procolophonians, and parapleurotans. In parapleurotans the lower temporal bar is open, forming an emargination (the reconstructed ancestral condition of the reptile crown group). An upper temporal fenestra is reconstructed as having evolved independently in araeoscelidians like *Petrolacosaurus* and in neodiapsids.

## Results

### Phylogenetic analyses

Maximum parsimony analysis and Bayesian inference (Fig. 9, Figs. S3–S5) place *Milleropsis* and other members of Millerettidae as the immediate sister group to Neodiapsida (Bremer support = 5, posterior probability = 1). We propose the name Parapleurota nom. cl. nov for Millerettidae + Neodiapsida, which is supported by 21 unambiguous autapomorphies absent in traditional ‘eureptiles’ or ‘early diapsids’ including: the presence of a tympanic recess on the posterolateral surface of the skull, the loss of a lacrimal contribution to the naris, the presence of a quadrate foramen formed only by the quadrate and quadratojugal, lateral exposure of the quadrate, the presence of both anterior and posterior surangular foramina, the absence of an ossified sphenethmoid, laterally directed caudal ribs, two lateral elements of gastralia per row or fewer, a midline gastral element, and a proximal expansion of the fifth metatarsal. We define Parapleurota as all taxa sharing a more recent common ancestor with *Milleretta rubidgei* Broom [1938]^63^ and *Youngina capensis* Broom 1914^64^ than with *Petrolacosaurus kansensis* Lane 1945^65^, *Orovenator mayorum* Reisz et al. 2011^66^, *Procolophon trigoniceps* Owen 1876^67^, or *Mesosaurus tenuidens* Gervais 1865^68^.

**Figure 9.**
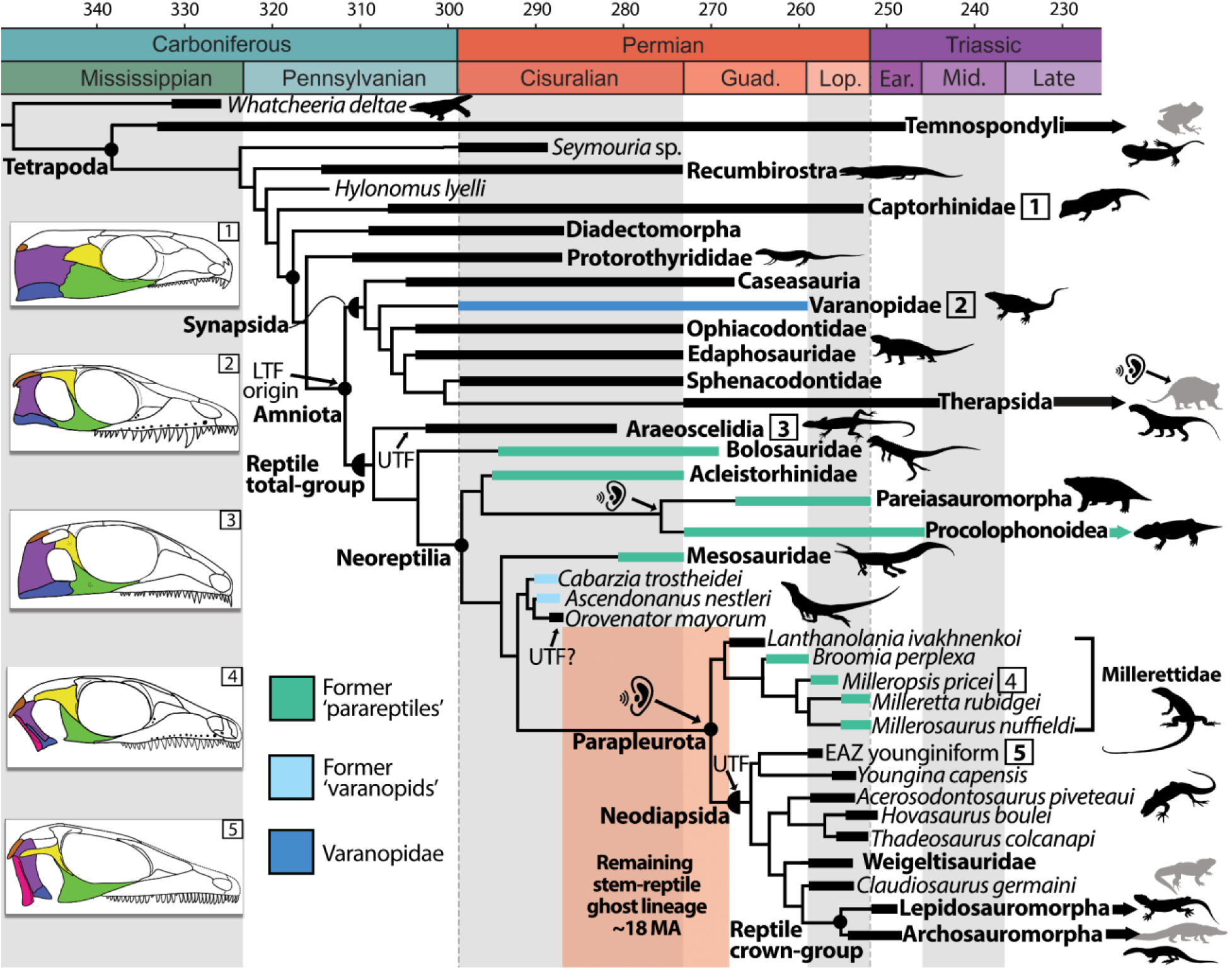
Simplified cladogram using parsimony and the maximum clade credibility tree recovered from Bayesian analysis using the time-calibrated FBD model on Paleozoic and Triassic tetrapods; emphasizing reptile relationships and patterns of temporal fenestration. Branch and skull coloring matches that of Fig. 1, showing the polyphyly of Varanopidae and traditional Parareptilia. The ear symbol denotes the hypothesized independent origins of tympanic hearing in Paleozoic amniotes, including a single origin ancestral to the reptile crown (see Figures 11 and 13). The last amniote common ancestor is hypothesized to have possessed a lower temporal fenestra (LTF). UTF indicates the presence of an upper temporal fenestra, which may have evolved up to three times in the reptile total-group, including once in Neodiapsida (see Figure 12). Skull reconstructions: 1, *Captorhinus*; 2, *Heleosaurus*; 3, *Petrolacosaurus*; 4, *Milleropsis*; and 5, EAZ younginiform. Silhouettes are available from Phylopic (www.phylopic.org) under CC BY 3.0 licenses or within public domain. Black silhouettes represent extinct taxa, whereas grey silhouettes represent extant members of crown Tetrapoda. We elect to place Lissamphibia within Temnospondyli^108^, but note that lissamphibians have been hypothesized as closer to amniotes than to temnospondyls^109–111^, a hypothesis that is not tested here. The reptile crown-group labelled here reflects molecular hypotheses of reptile evolution that support a close relationship between turtles and archosaurs, forming the clade Archelosauria^48–49^.

The enigmatic ‘neodiapsid’ *Lanthanolania*^38^ is found to be the basalmost millerettid (posterior probability = .98), supported by features including the presence of an anterior maxillary foramen (Ch. 57), an enclosed vidian canal (Ch. 260), and flat ventral plate of the parabasisphenoid which lacks the deep median fossa present in non-saurian neodiapsids such as Youngina (Ch. 265). The early amniotes *Orovenator*^37^ and *Ascendonanus*^62^, previously interpreted as close relatives to Neodiapsida or Varanopidae, are found crownward of Ankyramorpha (posterior probability = 1) as outgroups to Parapleurota (Figs. S3,4). We find *Saurosternon* and ‘younginiforms’ as the earliest-diverging neodiapsids in our parsimony (Fig. S3) and MkV analyses (Fig. S5), with tangasaurids, weigeltisaurids, and *Claudiosaurus* in successively more crownward positions (Fig. 9). To reconcile previous definitions of Neodiapsida with ongoing shifts in early reptile phylogenetics, we redefine Neodiapsida) nom. cl. conv here as all taxa sharing a more recent common ancestor with *Youngina capensis* Broom 1914^64^, *Claudiosaurus germaini* Carroll 1981^69^, *Hovasaurus boulei* Piveteau 1926, and *Lacerta agilis* Linnaeus 1758 than with *Orovenator mayorum* Reisz et al. 2011^66^, *Milleretta rubidgei* Broom [1938]^63^, *Procolophon trigonicep*s Owen 1876^67^, *Bolosaurus striatus* Cope 1878^70^, *Mesosaurus tenuidens* Gervais 1865^68^, and *Petrolacosaurus kansensis* Lane 1945^65^. The nomenclatural acts for Parapleurota and Neodiapsida within this manuscript are not considered final, and ultimately these definitions will be registered with Phylocode at time of publication in another journal.

The monophyly of the traditional groupings ‘Parareptilia’, ‘Eureptilia’, and ‘Diapsida’ is not supported in any of our analyses, with an additional 81 additional steps required in our parsimony analysis to force the monophyly of ‘Parareptilia’, 105 for ‘Eureptilia’ and 64 for ‘Diapsida’ (Fig. 10). When the monophyly of both ‘Eureptilia’ and ‘Diapsida’ was enforced in our constraint analyses, Millerettidae consistently emerges as sister to Neodiapsida, demonstrating robust support for the novel group Parapleurota even under alternative hypotheses of stem reptile interrelationships (Fig. 10). Our tomography data reveals the presence of classical ‘parareptilian’ features in former ‘eureptiles’ including araeoscelidians, neodiapsids, and early-diverging members of the reptile crown, such as prefrontal-palatine contact and a foramen orbitonasale, suggesting that these features are instead symplesiomorphies of the reptile total-group (Fig. 4). Other features previously hypothesized to support the monophyly of ‘Parareptilia’ or ‘Eureptilia’ are widespread among early amniotes (e.g., swollen dorsal vertebrae), were incorrectly optimized due to the low taxon sampling of previous studies (e.g., the presence or absence of a supraglenoid foramen), or misinterpreted for millerettids (e.g., the supposed lacrimal contribution to the naris, Figs. 2–3).

**Figure 10.**
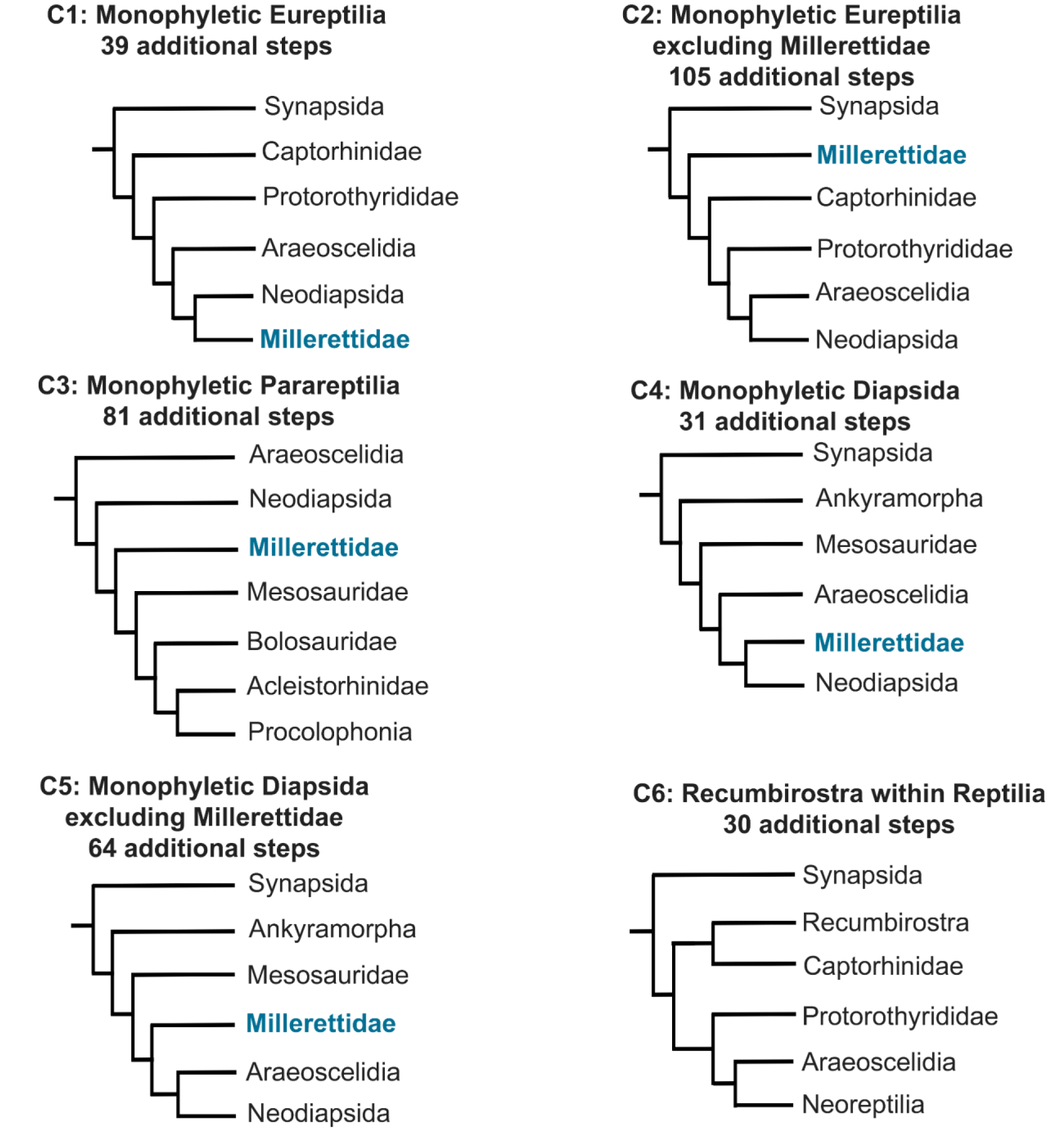
Simplified topologies of strict consensus trees returned by Maximum Parsimony analyses testing alternative hypotheses of early reptile evolution. Trees show cost of constraining the monophyly of Eureptilia (C1, 39 additional steps); Eureptilia excluding Millerettidae (C2, 105 additional steps); Parareptilia (C3, 81 additional steps); Diapsida (C4, 31 additional steps); Diapsida excluding Millerettidae (C5, 64 additional steps) as well as the placement of Recumbirostra as reptiles (C6, 30 additional steps). Neoreptilia in C6 includes ankyramorph “parareptiles,” neodiapsids, and millerettids.

Araeoscelidians, conventionally interpreted as close relatives of Neodiapsida^5–6^, are instead supported here as the sister group to all other stem-reptiles (posterior probability =.98, Bremer support = 3) (Fig. 9, Fig. S3–S5), corroborating recent work^4,19,22^. Bolosaurid ‘parareptiles’^29^ excluding *Erpetonyx* are placed immediately crownward of Araeoscelidia with strong support (posterior probability = .92, Bremer support = 2), although we note that this phylogenetic hypothesis merits further investigation. The monophyly of Ankyramorpha^6^, a group of former ‘parareptiles’ consisting of Acleistorhinidae and Procolophonia, is strongly supported (posterior probability = 1, Bremer support = 3), and ankyramorphs are placed immediately crownward of Bolosauridae (Fig. 9, Fig. S3,S5). *Lanthanosuchus*, which was originally allied with Olson’s ‘Parareptilia’ when it included seymouriamorphs and most recently has been found as an ankyramorph parareptile^6^, is found far outside of crown Amniota as an early-branching tetrapod (Figs. S3–5), although its exact placement among tetrapods necessitates additional early tetrapod taxonomic and character sampling. Mesosauridae^71–73^ are placed another node crownward of Ankyramorpha, albeit with moderate support (posterior probability = 0.91).

Facilitated by our synchrotron tomography data of *Heleosaurus*, we demonstrate that Varanopidae — a contentious early amniote group that has recently been found on the reptile stem lineage,^4^ but more often placed as members of the mammalian total group (Synapsida)^5,6,24^ — are indeed synapsids. We find varanopids specifically as the earliest-diverging eupelycosaurs (posterior probability = 0.98) supported by 14 unambiguous synapomorphies. A synapsid placement of Varanopidae is supported by synapsid-like palatal and braincase features of *Heleosaurus* revealed in our scan data, and which were previously never incorporated as characters in phylogenetic analyses. These features include the presence of a dorsally positioned trigeminal notch (Ch. 233, Fig. 7B), a sheet-like shaft of the epipterygoid (Ch. 293, Fig. 6B), a floccular fossa that is enclosed by the supraoccipital (Fig. 7B), and the pathway of the abducens nerve (CN VI) through the dorsum sellae (Ch. 278). Other amniote plesiomorphies documented in *Heleosaurus*, such as an epipterygoid contribution to the basicranial articulation (Ch. 288, Fig. 6B) or the presence of two coronoid ossifications (Ch. 416), are incompatible with a placement of Varanopidae among more crownward stem reptiles (contra^4^). Previous hypotheses of varanopid relationships relied on externally visible features such as temporal fenestration^5,6^, the presence of caniniform teeth, and postcranial features such as gracile limb proportions^4^ that show greater homoplasy among early amniotes, generating uncertainties that had fueled recent debates about whether varanopids were synapsids or stem-reptiles^4^.

**Figure 6.**
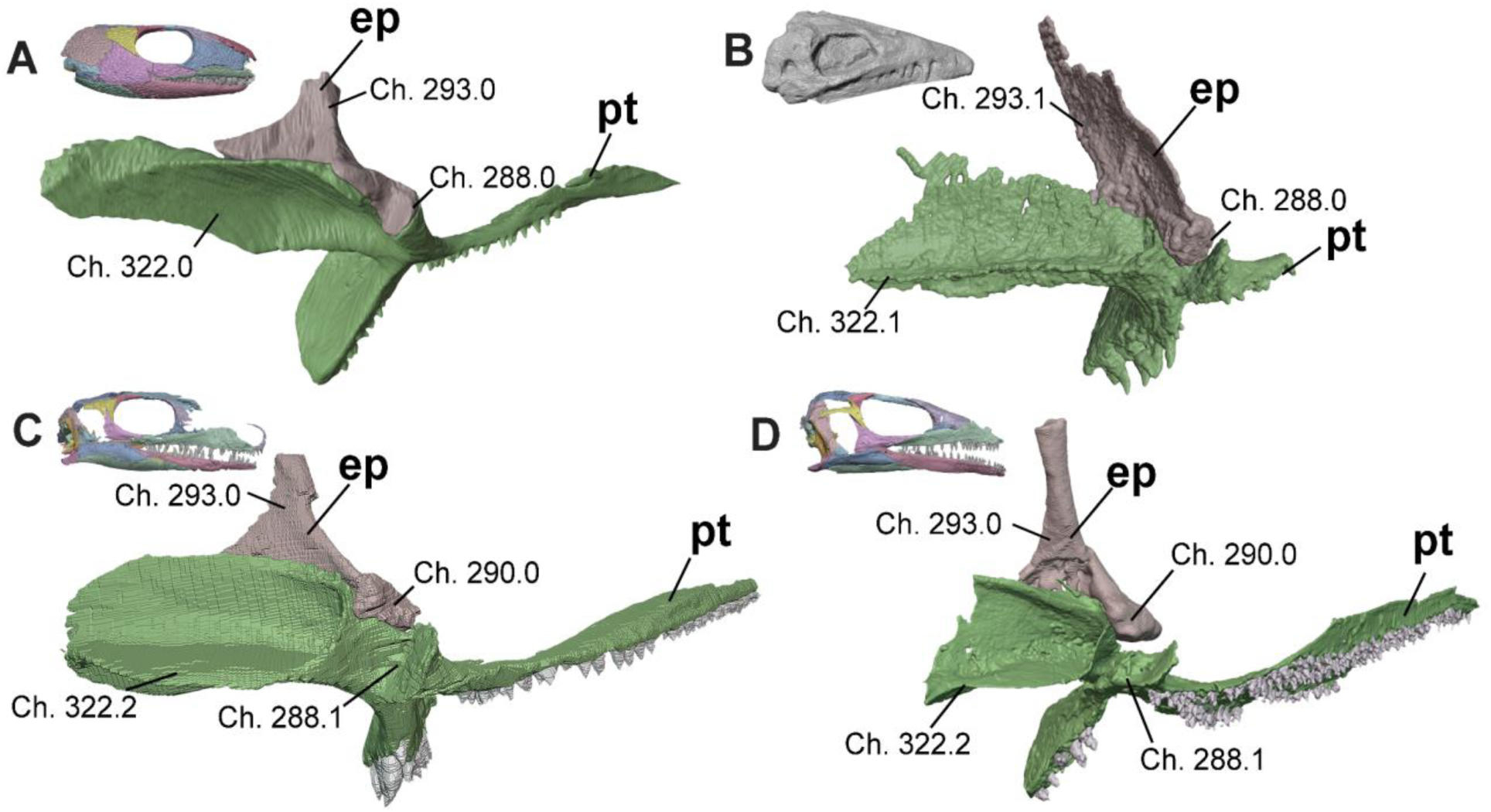
Basicranial region of **A**, *Captorhinus* (OMNH 44816)^91^, **B,** *Heleosaurus* (SAM-PK-K8305), **C**, *Milleropsis* (BP/1/720), and **D**, EAZ younginiform (SAM-PK-K7710) in oblique posterior view demonstrating the reduction and eventual absence of an epipterygoid contribution to the basal articulation seen in Parapleurota. Select characters from our dataset include: the presence and development of the tympanic flange of the pterygoid (**Ch. 322,** present in early crown amniotes), contribution of the epipterygoid to the basicranial articulation (**Ch. 288**, lost in Parapleurota), the presence of a medial lip of the epipterygoid (**Ch. 290,** lost in some lepidosauromorphs), and the morphology of the dorsal shaft of the epipterygoid (**Ch. 293,** sheetlike in synapsids including Varanopidae). Abbreviations: ep, epipterygoid; pt, pterygoid.

**Figure 7.**
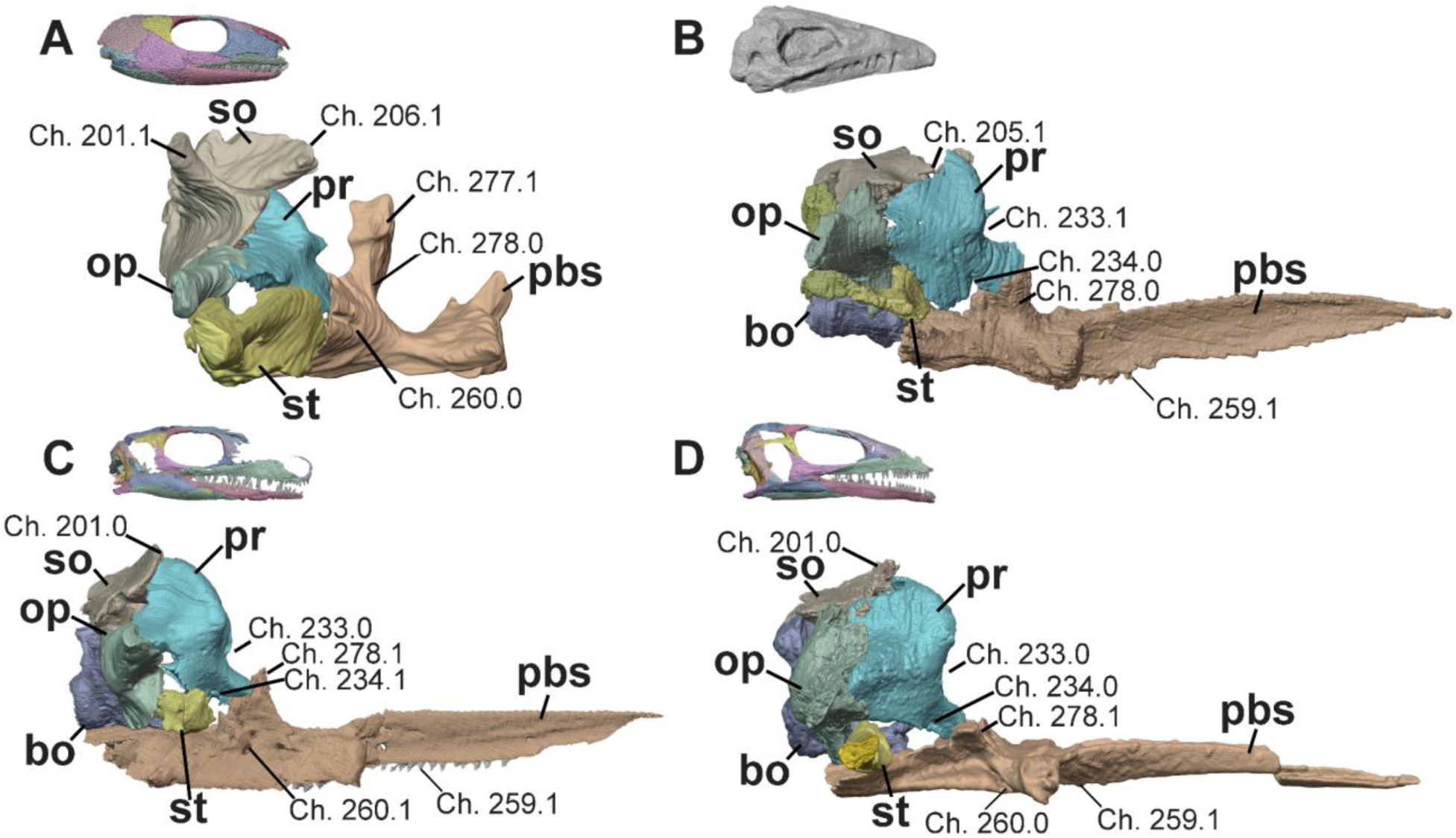
Braincase of **A**, *Captorhinus* (OMNH 44816)^91^, **B,** *Heleosaurus* (SAM-PK-K8305), **C**, *Milleropsis* (BP/1/720), and **D**, EAZ younginiform (SAM-PK-K7710) in right lateral views, focusing on the reduction of the stapes relative to the otic capsules and the pathways of the cranial nerves through the braincase. Select characters from our dataset include: the presence of lateral ascending processes of the supraoccipital (**Ch. 201**, present in captorhinids and some recumbirostrans), a supraoccipital contribution to the floccular fossa (**Ch. 205**, present in synapsids including Varanopidae), the median ascending process of the supraoccipital (**Ch. 206,** present in captorhinids and some recumbirostrans), the position of the trigeminal nerve relative to the ventral process of the prootic (**Ch. 233**, located posterodorsal to the ventral process of the prootic in synapsids including varanopids), the pathway of the facial nerve (VII; **Ch. 234**, a notch in millerettids), cultriform process dentition (**Ch. 259**, absent in most crown reptiles within this sampling), the location of the vidian canal or groove (**Ch. 260**, within the lateral wall of the braincase in procolophonians, millerettids, and some lepidosauromorphs), the height of the dorsum sellae (**Ch. 277**, tall in captorhinids and the recumbirostran *Pantylus*), and the pathway of the abducens nerve (VI) through the parabasisphenoid (**Ch. 278,** through the clinoid processes and prootic articulation in parapleurotans). Abbreviations: bo, basioccipital; op, opisthotic; pbs, parabasisphenoid, pr, prootic; so, supraoccipital; and st, stapes.

We find evidence for the placement of nominal early ‘eureptiles’ such as captorhinids and protorothyridids outside of crown Amniota, corroborating older hypotheses^15–16^ of the ‘Captorhinomorpha’, ‘Romeriidae’, and ‘Cotylosauria’ as ancestral to fenestrated amniotes^74–76^. Protorothyrididae is found here to form the sister-group to Amniota (posterior probability = .76, Fig. S5), whereas Captorhinidae is placed rootward (posterior probability = 1, Fig. 3, Figs. S3– 5), a position that remains stable even when characters relating to temporal fenestration are removed (Fig. S3B). The ‘eureptile’ *Hylonomus*, previously suggested to be a stem amniote^15–16^ or the earliest stem reptile^34^, and therefore an important node age constraint in molecular phylogenetic studies of vertebrate evolution as the earliest crown amniote, is found outside of crown Amniota (posterior probability = 0.85). Pre-cladistic studies similarly interpreted *Hylonomus* as being outside of the amniote crown, either as a ‘cotylosaur’^74–75^or protorothyridid^76^. All of the features previously hypothesized to place *Hylonomus* within Amniota (e.g., the presence of a transverse flange of the pterygoid^15^ (Ch. 313) or astragalus^7^ (Ch. 620)) or the reptile total group (e.g., absence of postorbital-supratemporal contact^34^ (Ch. 128) or slender limbs^7^ (Ch. 602)), are also variably present in the stem amniote groups Seymouriamorpha, Diadectomorpha, and Recumbirostra^31^ as well as early synapsids^77^ and are here interpreted as symplesiomorphies of Amniota. These character distributions corroborate recent studies^25, 31^ demonstrating the widespread presence of classical amniote or reptile features among stem and crown amniotes, including synapsids (Table S2).

Recumbirostra are found here outside of crown Amniota, crownward of seymouriamorphs, and 30 additional steps are required to place them with the reptile total-group (Fig. 9). Previous features suggesting a stem-reptile placement of Recumbirostra,e.g., a crista interfenestralis of the opisthotic (Ch. 224) or a single ossification of the supraoccipital (Ch. 198), are documented in early-diverging synapsids, including varanopids (Fig. 7B) and ophiacodontids^77^. The placement of Recumbirostra remains stable even when characters related to the presence of temporal fenestration are removed (Fig. S1). More detailed anatomical descriptions using 3D imaging of fossil taxa including ‘microsaurs’, protorothyridids, *Hylonomus*, and the earliest synapsids may help resolve uncertainties in the evolutionary origin of amniotes, which merits further investigation.

### Stratigraphic Fit

The results of our stratigraphic-fit analyses demonstrate that our phylogenetic hypothesis of millerettids as close relatives of Neodiapsida is significantly more congruent with expectations from a stratigraphic null model than previous hypotheses of early reptile evolution by at least 27% (GER) and 28% (MSM*). Traditional phylogenetic hypotheses yielding a monophyletic Eureptilia, Parareptilia, and Diapsida are less congruent with stratigraphy (therefore implying longer ghost lineages) than hypotheses finding no support for these groups^24^, including this study. The differences in MSM* values (MSM*Diff) between our phylogenetic hypothesis and our constraint analysis are summarized in Table S4. MSM*Diffs between these two topologies were positive in 100% of comparisons (Fig. S9). On average, the closure of the expansive ghost lineage for Millerettidae and three of the four major ghost lineages (‘Eureptilia’, ‘Diapsida’, and Neodiapsida ghost lineage B, Fig. 1) among the reptile stem improved stratigraphic fit by at least 28%, although a ghost lineage along the parapleurotan stem (equivalent to ‘Neodiapsid ghost lineage A’, Fig. 1) still stretches from the middle early Permian (Artinskian)^37^ to the appearance of millerettids in the beginning of the middle Permian (Roadian)^38^ (Fig. 9).

### Ancestral-State Reconstruction

Our ancestral-state reconstructions mapped anatomical features related to tympanic emargination and temporal fenestration using two models of character state evolution. In each instance, the equal rates (ER) model, which assumes a single rate of evolution for all branches within our topology, was best supported, outperforming the all-rates-different model (ARD) according to Akaike’s information criterion.

Our findings suggest that the tympanic fossa of Parapleurota originated on the quadrate, squamosal, and quadratojugal and subsequently migrated to be primarily supported by the quadrate (Fig. 11B). The ancestral-state estimates concerning the presence of a tympanic emargination provide robust evidence that the tympanic fossa originated twice during early reptile evolution: once within Parapleurota (including the reptile crown) and once within Procolophonia (Fig. 11A). The deep-time origins of this structure are supported by the shared presence of a tympanic fossa on the quadrate, squamosal, and quadratojugal in millerettids and early-diverging neodiapsids, including SAM-PK-K7710 (Figs. 5D, 11), *Youngina* (pers. obs. BP/1/375), and *Hovasaurus* (pers. obs. MNHN.F.MAP373).

**Figure 5.**
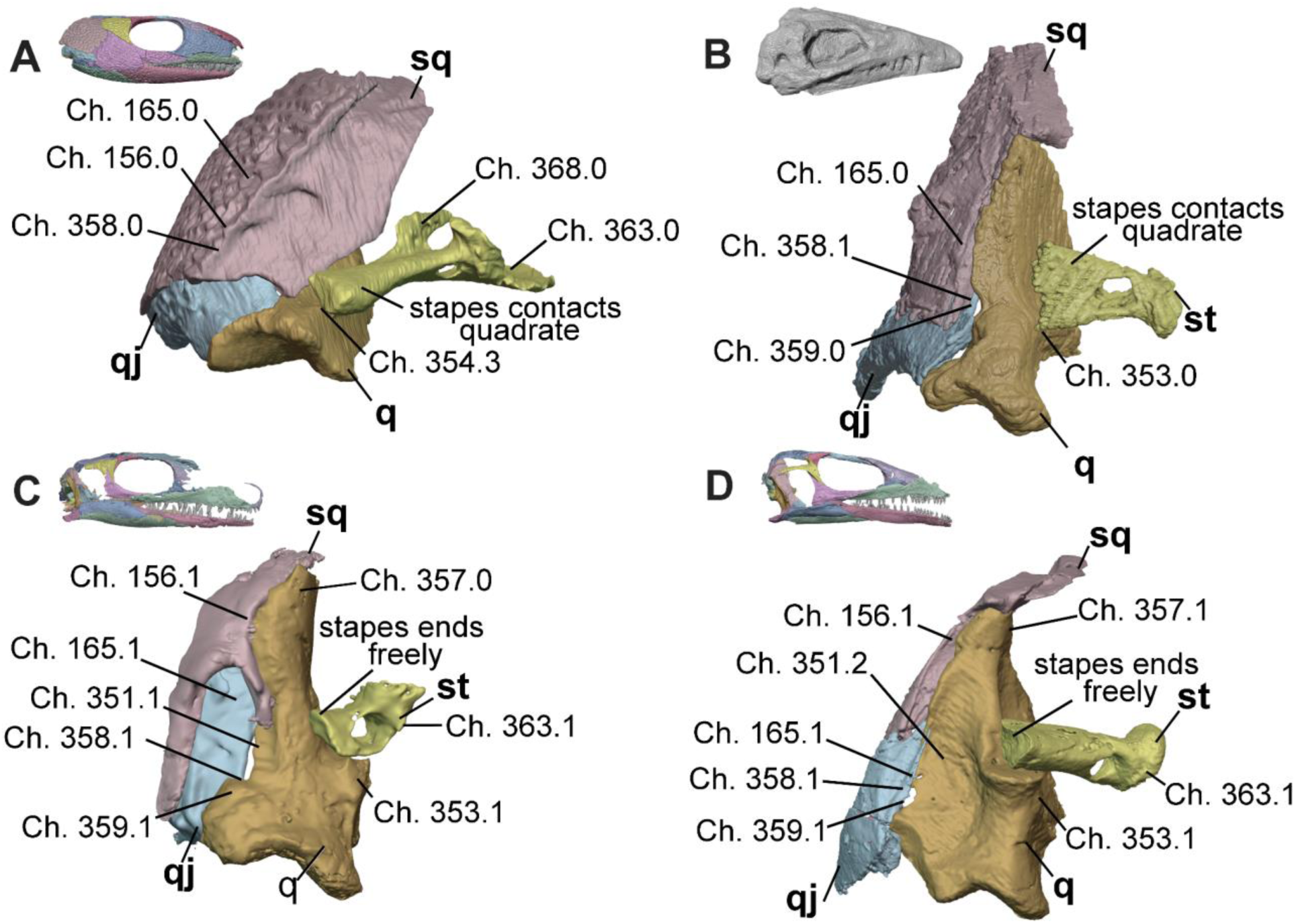
Tympanic and middle ear region of **A**, *Captorhinus* (OMNH 44816)^91^, **B,** *Heleosaurus* (SAM-PK-K8305), **C**, *Milleropsis* (BP/1/720), and **D**, EAZ younginiform (SAM-PK-K7710) in posterior views demonstrating stepwise reductions in the size and anatomy of the squamosal and stapes and the eventual appearance of a tympanic emargination. Select characters from our dataset include the occipital shelf of the squamosal (**Ch. 156**, lost in Parapleurota), presence or absence of a tympanic emargination (**Ch. 165**, present in Parapleurota and convergently in Procolophonia) and, if present, its presence on the quadrate (**Ch. 351**, present in Neodiapsida, although millerettids bear a small contribution here), the relationship of the quadrate with the stapes (**Ch. 353**, early amniotes bear a stapedial groove that osseously receives the stapes whereas early parapleurotans bear a stapedial boss that cartilaginously supports the ventral surface of the stapedial shaft), cephalic condyle of the quadrate (**Ch. 357,** present in some neodiapsids and early crown reptiles), presence (**Ch. 358**, present in all early amniotes) and location (**Ch. 359,** located between the quadratojugal and quadrate in neodiapsids) of the quadrate foramen, the presence of the dorsal process of the stapes (**Ch. 368,** lost in Parapleurota), and size of stapedial footplate (**Ch. 363**). Abbreviations: q, quadrate; qj, quadratojugal; sq, squamosal; and st, stapes.

Our ancestral-state reconstruction predicts that the last common ancestor of amniotes bore a lower temporal bar framing the lower boundary of a lower temporal fenestra with subsequent closure in pareiasaurs, the araeoscelidian *Araeoscelis*, and in adult individuals of *Milleretta*. The lower temporal bar was lost in the last common ancestor of Parapleurota, resulting in the presence of a lower temporal emargination (Fig. 12A). These models also suggest that the lower temporal opening of procolophonoids, here termed a ‘lower temporal concavity’, is not strictly homologous with the ‘open’ lower temporal openings of Parapleurota, as both features evolved independently from taxa that possessed a closed lower temporal bar (Fig. 12A). The important phylogenetic implication of this finding is that the lower temporal emargination is present both in early neodiapsids and in nominal parareptiles, and so can no longer be interpreted as a synapomorphy supporting a monophyletic Parareptilia^6^.

Our ancestral-state reconstructions suggest that the upper temporal fenestra had two independent origins in reptiles, once in Late Carboniferous Araeoscelidia at the base of the reptile total-group and again in Neodiapsida in the late Permian, with the upper temporal fenestra being absent in all other stem-reptiles except *Orovenator*^37^, possibly *Lanthanolania*^38^, and in juvenile millerettids (Fig. 3), although the former two taxa have incomplete preservation of the temporal region (Fig. 12B). These findings corroborate interpretations from recent phylogenetic analyses which also find araeoscelidians including the famously diapsid *Petrolacosaurus* as either stem-amniotes^24,25^ or the earliest-diverging group of stem-reptiles^4,28^, rejecting the hypothesis that araeoscelidians are ‘eureptiles’ closely related to Neodiapsida.

## Discussion

### Millerettids as crownward stem-reptiles

Our phylogenetic results place various groups of middle and late Permian fossil reptiles, including millerettids, within the crownward portion of the reptile stem lineage as close relatives of Neodiapsida (Fig. 9). The placement of Millerettidae as sister to Neodiapsida (forming Parapleurota) is robustly supported in our analyses using parsimony, Bayesian FBD and Mkv-only, and when analyses are constrained to represent alternative hypotheses of early reptile evolution (Fig. 10). This relationship is further corroborated by the relatively continuous stratigraphic record of the middle and late Permian Karoo Basin, which preserves a succession of millerettid and neodiapsid stem-reptiles^78^. Other anatomical observations are consistent with a sister-relationship between Millerettidae and Neodiapsida, but are not optimized as unambiguous synapomorphies due to the uncertain scoring of these characters in close relatives of Parapleurota (e.g., *Orovenator*), including: a quadratojugal foramen with no contribution by the squamosal (Ch. 359, Fig. 5), no contribution of the epipterygoid to the basal articulation (Ch. 288, Fig. 6), the pathway of CN VI through the clinoid processes rather than the dorsum sellae (Ch. 278, Fig. 7), and a posteriorly directed ilium (Ch. 592).

Our results differ substantially from the otherwise prevailing phylogenetic paradigm in which millerettids were hypothesized as parareptiles^5–7^, and therefore only distant relatives of Neodiapsida and crown reptiles. These studies were hampered by the lack of tomographic data on key taxa, which resulted in the critical undersampling of cranial characters, especially for the braincase, palatoquadrate, and palate (Figs. 4–8). Furthermore, early cladistic analyses of early reptile origins^4–6^ lacked a robust sampling of non-saurian neodiapsids (Fig. S1) and included only one representative of Millerettidae: the later-branching and latest-occurring *Milleretta rubidgei*, which closes its lateral temporal fenestra during ontogeny^9,45^. Our high-resolution tomographic data reveal that earlier-branching representatives of Millerettidae^44^ present seemingly derived features such as the lack of contact between the postorbital and supratemporal and a gastral basket consisting of a midline element and two lateral elements, which are now interpreted as the plesiomorphic condition for Millerettidae (Fig. S3). Both of these features were previously interpreted as synapomorphies of Eureptilia^7^ and Neodiapsida, respectively. Another significant factor in the sister-placement of Millerettidae and Neodiapsida are unrecognized plesiomorphies in the early neodiapsid skull revealed by our tomographic data, such as the presence of multiple rows of dentition on the transverse flange of the pterygoid and the presence of dentition on the ventral plate on the parabasisphenoid in SAM-PK-K7710. The loss of this palatal dentition was previously interpreted as a neodiapsid autapomorphy^3^, but our results instead suggest that these features were lost later in neodiapsid evolution.

Our finding of robust character support for millerettids as sister to Neodiapsida solidifies a fundamental shift in how paleontologists can understand Paleozoic reptile evolution. Paleozoic stem reptiles have long been recognized as having explored a substantial range of habitat, dietary, and locomotor spaces in the Permian (bipedal bolosaurids^96^; large-bodied herbivorous pareiasaurs^80^ and their burrowing procolophonoid^81^ relatives; semiaquatic mesosaurids^72^ and tangasaurids^22^; the arboreal *Ascendonanus*^59^; and gliding weigeltisaurids^23^). Our phylogenetic hypothesis underscores that this ecomorphological diversity in the Permian not only precedes more extreme radiations in the Triassic, but in fact represents a succession of outgroups to the reptile crown that documents the evolutionary origin of crown reptile traits along an extensive stem lineage, akin to the established trends of evolutionary experimentation and innovation along the mammalian stem lineage during the same interval^82^.

### Improving the stratigraphic fit of the early reptile fossil record

The placement of the middle and late Permian Millerettidae as the sister group to Neodiapsida substantially improves the stratigraphic fit of the fossil record of crownward stem-reptiles by 28% compared to traditional phylogenetic hypotheses of reptile relationships (Fig. S11, Table S4). This phylogenetic position for Millerettidae reduces their hypothesized ghost lineage extending down to the Late Carboniferous when this group is interpreted as branching among basal parareptiles (shortened by ∼35 Ma). Furthermore, the placement of millerettids as sister to Neodiapsida improves our understanding of the evolution of neodiapsid outgroups from the middle Permian (Roadian) onward (∼10 MA; Neodiapsida ghost lineage B of Figure 1). Our phylogenetic hypothesis is congruent with the temporally sequential appearances of millerettid and neodiapsid stem-reptiles in the well-sampled and continuous stratigraphic sections of the Karoo Basin in South Africa^83^. However, the long-recognized ghost lineage (Neodiapsida ghost lineage A) extending from *Orovenator*^37^ to the earliest parapleurotan, *Lanthanolania*^38^, remains an issue, as this gap aligns with known biases in the biogeography of the fossil record where early Permian rocks preserve equatorial ecosystems but middle to late Permian rocks preferentially preserve high latitudes^84^. We suggest further fieldwork in Kungurian-to Roadian-age formations as the most likely solution.

Our phylogenetic hypothesis proposes that the groups ‘Eureptilia’ and ‘Parareptilia’^5^, which are common textbook examples of early reptile diversification, are in fact polyphyletic assemblages of distantly related taxa. This result removes the need for a Carboniferous ‘Eureptilia’ ghost lineage^2^ (Fig. 1), and we find support for only a short ghost lineage (∼4 Ma) between the earliest stem-mammals and stem-reptiles. Furthermore, our placement of the ‘eureptile’ *Hylonomus* (∼318 Ma) as a stem amniote raises doubts about its suitability as a minimum-age calibration^85^ for the amniote crown group and indicates a need for restudy of the anatomy of this important early taxon. Similarly aged putative amniotes such as *Protoclepsydrops*^15^, identified as synapsid without justification— are similarly unsuited for minimum-age calibration^85^. Future molecular calibrations for the amniote crown should use more confidently assigned members such as the synapsids *Melanedaphodon*^86^ (∼307 Ma) and *Archaeothyris* (∼306 Ma) in the Moscovian.

The polyphyly of Parareptilia distributes various groups of former ‘parareptiles’ as successively crownward outgroups of Parapleurota and greatly contributes to the improvement of our hypothesis to the stratigraphic fit of the early reptile fossil record (Fig. 3). In particular, the breaking apart of Parareptilia shortens the ghost lineage for Neodiapsida by closing the ‘Diapsida’ ghost lineage (Fig. 1, ∼15 Ma) that extended from *Petrolacosaurus* in the Late Carboniferous (Gzhelian^35^) until the appearance of *Orovenator* in the early Permian (Artinskian^37^) (Fig. 3). This ‘Diapsida’ ghost lineage is now filled by the successive divergences of the Ankyramorpha, Mesosauridae, and *Ascendonanus* shortly after the origin of the reptile total-group (Fig. 9), although the evolutionary succession and phylogenetic relationships of these groups are ripe for further investigation. Ankyramorpha contains many of the taxa previously referred to Parareptilia and has a near-uninterrupted fossil record throughout the end-Paleozoic, in part due to their cosmopolitan distribution overcoming biases in the rock record, standing in contrast with nearly all other early reptile groups^87^.

### Evolution of Temporal Fenestration in Amniota

Temporal fenestration morphologies have been pivotal in hypotheses for early amniote evolution and classification^10–11, 88–89^. These propose that distinct patterns of fenestration evolved independently on the mammal and reptile stem lineages (the ‘synapsid’ and ‘diapsid’ patterns) from a common ancestor that lacked fenestrae. Such historical schemes have rarely fit neatly into phylogenetic topologies, which generally find multiple additional origins of temporal fenestration within Amniota^4–6^. Our findings suggest that taxa lacking temporal fenestrae such as Protorothyrididae and Captorhinidae (but see^24–25^) are better interpreted as stem amniotes, a phylogenetic result that is found even when excluding characters related to temporal fenestration (Fig. S3). We find that a lower temporal fenestra (LTF) closed by the squamosal, quadratojugal, and jugal was present in the most recent common ancestor of the amniote crown groups, but that the subsequent evolution of temporal fenestration, and the arrangement and contacts round the fenestrae, were remarkably homoplastic (Fig. 12). For example, early synapsids possess an anteroventral process of the squamosal that excludes the quadratojugal from the LTF^4^, whereas later synapsids exclude the quadratojugal from the lower temporal bar entirely (Fig. 12). The LTF was also frequently modified among Paleozoic stem reptiles, including its closure Araeoscelidia^41^, Pareiasauria^81^ and mature individuals of *Milleretta* (Fig. 3) and the loss of the lower temporal bar (resulting in a lower temporal emargination) in the common ancestor of Parapleurota. Later groups of Mesozoic reptiles continued to make substantial modifications to the ancestral LTF with closure in Pantestudines^91^ and the reformation of a lower temporal bar (resulting in a lower temporal fenestra) in Archosauria and *Sphenodon*^90^.

Our phylogenetic topology also challenges the conventional view of homology between the doubly fenestrated ‘diapsid’ conditions present in the araeoscelidian *Petrolacosaurus*^36^, in *Orovenator*^37^, and in neodiapsids. Ancestral-state estimation instead suggests that an upper temporal fenestra (UTF) evolved independently in these three groups (Fig. 12; and see^4,24–28^).

This observation is congruent with the absence of the diapsid condition in the only other adequately known araeoscelidian, *Araeoscelis* (which lacks a lower temporal fenestra^41^), its absence in various neoreptiles, and the notably different fenestral architecture of araeoscelidians compared to both *Orovenator*^37^ and neodiapsids^19^ The usage of Diapsida^34^for a groupcomprising araeoscelidians and neodiapsidsis therefore unsupported, and we suggest avoiding the use of ‘Diapsida’ entirely.

Interestingly, millerettids such as *Milleretta rubidgei*^45^ and *Milleropsis pricei*^44^ possess a unique ventral flange of the parietal that closes an unossified gap between the parietal, postorbital, and squamosal during ontogeny, a feature that has been hypothesized to represent the secondary closure of an upper temporal fenestra in this group^8^. This high degree of plasticity of temporal architecture among early reptiles may result from the high functional relevance of fenestrae to jaw kinematics^90–91^. These findings necessitate future research into the homology of the upper temporal fenestra in the last common ancestor of modern reptiles.

### Tympanic hearing was initiated before the origin of the reptile crown group

Our phylogeny and novel anatomical observations of millerettids (Fig. 2) and SAM-PK-K7710 (a non-saurian neodiapsid with a tympanic emargination on the quadrate) show that tympanic hearing first evolved among early parapleurotans in the middle Permian (Fig. 13) and was later inherited by the common ancestor of crown reptiles. Although bolosaurid ‘parareptiles’ have been recently implicated in the origins of tympanic hearing^27^, this conclusion resulted from misidentification of the robust bolosaurid stapes as an opisthotic^27^. Our results are congruent with Laurin’s hypothesis^93^ and recent developmental work on crown reptiles^19^ that suggest a single origin of crown reptile tympanic hearing but differs from some previous work^20^ that proposed up to four separate (Triassic) origins of tympanic hearing among crown reptiles – once within Lepidosauromorpha^94^, once within Testudines^95^, and at least twice within Archosauromorpha, including a single shared origin for modern Archosauria^96^. Previous anatomical studies often used a narrow concept of osteological correlates for the presence of a tympanum, restricted to the observation of both an emargination and tympanic crests on the quadrate specifically, giving rise to the hypothesis of multiple origins of the tympanum within the reptile crown group^96^. We observe that the large tympanic emargination of the quadrate of crown reptiles (Ch. 351) has antecedents on the squamosal, quadratojugal, and quadrate seen in the tympanic fossae and crests in the posterior cranium of millerettids and early neodiapsids (Ch. 165, Fig. 5). Similar traits also evolved independently among procolophonians such as *Macroleter*^97^ that bear a large tympanic fossa on the squamosal and quadratojugal, but which differ in their extension onto the skull roof (Ch. 185) (Fig. 13).

**Figure 13.**
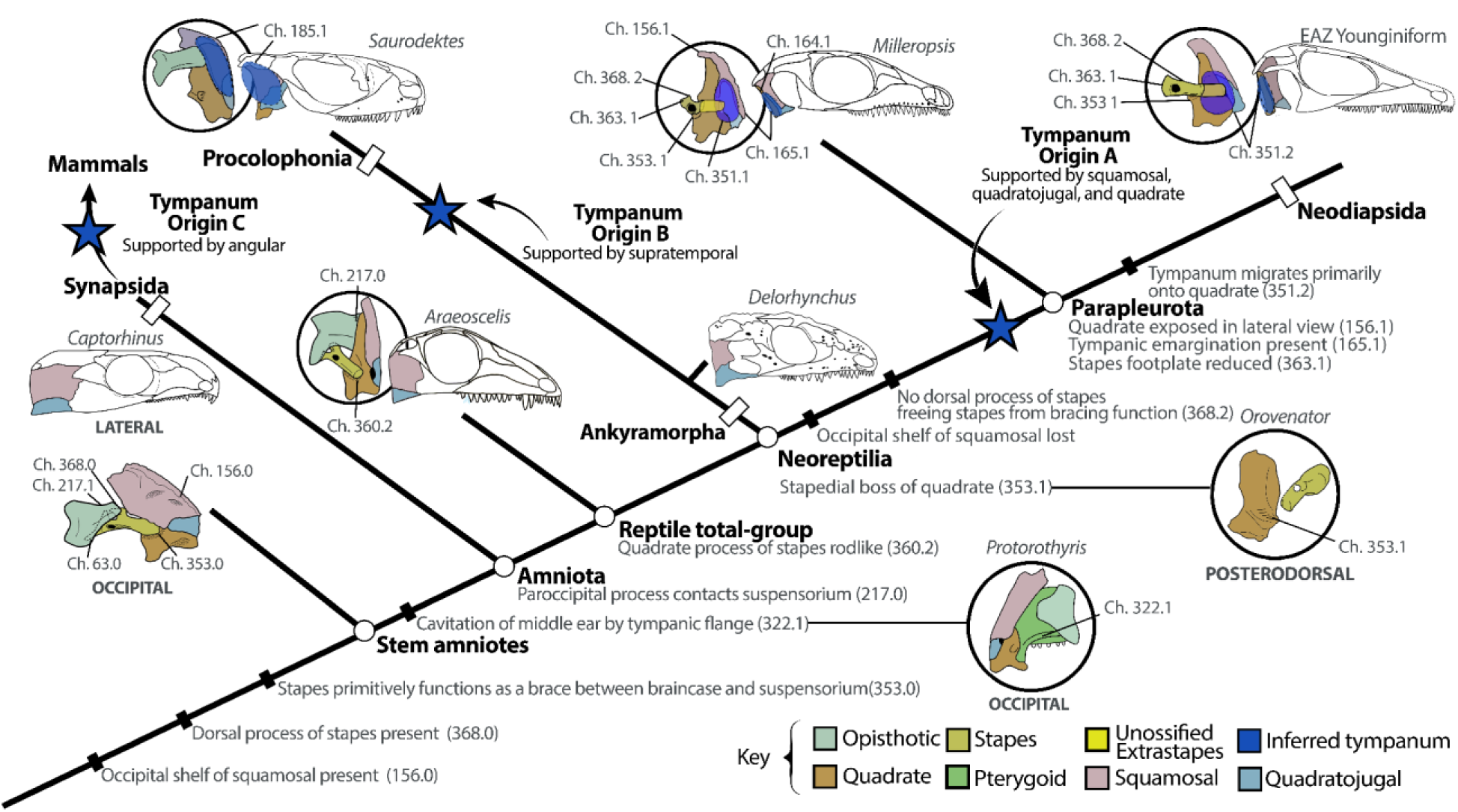
Evolution of amniote hearing using sensory characters of the middle ear from this study, demonstrating the stepwise reduction of the stapes, reduction in the squamosal and quadratojugal, and corresponding exposure of the quadrate in lateral view. Additionally, three independent origins of tympanic hearing among amniotes (in Synapsida, Procolophonia, and Parapleurota) are indicated by blue stars. Skull reconstructions are from this study or modified from references: *Captorhinus*^91^, *Araeoscelis*^112^, Procolophonia (*Saurodektes*^113^), and Parapleurota (*Milleropsis* and the EAZ younginiform, this study).

Our phylogeny and ancestral-state reconstruction (Fig. 11) provide evidence that a tympanum was likely already present among stem-reptiles before the origin of Parapleurota and was housed jointly between on the quadrate and squamosal/quadratojugal, as seen in *Milleropsis* and other millerettids (Figs 2, 11A, 13). The tympanum then migrated onto the quadrate (with at most a smaller contribution by the squamosal and quadratojugal) in Neodiapsida (Fig. 5D), possibly due to the evolutionary reduction of the squamosal (Ch. 156, Fig. 11B, 13) and its incorporation into the margin of the upper temporal fenestra^3^. In these early neodiapsids (e.g., SAM-PK-K7710, Fig. S2), the posterior margin of the quadrate remained straight, suggesting that the presence of a tympanic fossa on the quadrate is independent from the morphology of its posterior shaft. Furthermore, the tympanic fossa of early parapleurotans was smaller relative to the stapedial footplate compared to that of crown reptiles, indicating a lower area ratio (4:1 in *Milleropsis* and 7:1 in SAM-PK-K7710), and therefore provided poorer sound transmission than most crown reptiles, yet still was likely capable of amplifying sound. Observations and dissections of the middle ear of both living and fossil reptiles, which unequivocally demonstrate the presence of a tympanic membrane attachment onto the squamosal and quadratojugal in lizards, birds^98^, crocodilians^99^, and turtles^100^, are consistent with our proposal that the tympanic emargination of the quadrate has antecedents on other elements of the posterior skull (Figs. 9, 11).

Our observations of the stapes provide further evidence for the evolution of the reptile middle ear, which we extensively codify in our phylogenetic dataset (Fig. 5, SI Appendix). Previous studies used simplified concepts of stapedial variation, generally encoding it as a dichotomy between ‘robust’ and ‘rod-like’ morphologies^5–7^. We use a more detailed characterization of variation in stapedial morphology (SI Appendix 8, Chs. 360–370) and find evidence for the acquisition of potential auditory function of the stapes earlier in reptile evolution, by the middle Permian (Fig. 13). Stepwise reductions in the size of the stapedial footplate (Ch. 363) and columella (Ch. 366) throughout the Permian, paired with the losses of its dorsal process (Ch. 368) and stapedial foramen (Ch. 361), indicate that the stapes was free from a role as a brace for the skull by the late Permian. In contrast to earlier-diverging reptiles, the stapes of parapleurotans ends freely and is directed towards the tympanic fossa, suggesting that the stapes of millerettids and early neodiapsids may have acted as a proto-amplifier of sound (Fig. S2). The posteromedial placement of the stapedial boss in parapleurotans, which is located further posteriorly than in other neoreptiles, may have facilitated the freeing of the stapes. The increasing length of the stapedial shaft seen in neodiapsids (SAM-PK-K7710; Fig. S2) may have been influenced by the anterior shift of the otic capsules relative to the jaw joint in this group compared to millerettids. This may have enabled the stapes of these taxa to transduce sounds at higher frequencies, although this merits biomechanical investigation (Fig. 13). These results, corroborated by several recent developmental studies establishing the homology of the tympanic membrane^19^ and extracolumella^101^ in crown reptiles, demonstrate that complex neurosensory innovations including enhanced auditory capabilities were present in reptiles prior to the End-Permian Mass Extinction, predating the ecomorphological diversification of crown reptiles in the Triassic^2^.

## Conclusion

Our synchrotron tomography data allows new insights and interpretations into the anatomy of important stem reptile fossils, particularly for the Millerettidae. Our resulting phylogenetic hypotheses indicate the polyphyly of Eureptilia and Parareptilia and the sister relationship between Millerettidae and Neodiapsida challenge existing paradigms of early reptile evolution and allows for a reinterpretation of anatomical transformations during the evolutionary assembly of crown reptile anatomy in the Permian. This includes new hypotheses for the stepwise origins of tympanic hearing in Parapleurota and fenestration patterns across stem-reptiles. Future work may resolve additional questions regarding the evolutionary diversification of both stem and crown reptiles, including fieldwork^102^ and detailed studies on the anatomy of Carboniferous stem amniotes, will be central to our understanding of the early evolution of the reptile crown group and the rapid diversification of morphologically disparate reptile groups in the Triassic (e.g., Ichthyosauromorpha, Pantestudines, Sauropterygia, etc.).

## Data, scripts, code, and supplementary information availability

All computed tomography data and three-dimensional models for taxa reported in this paper are available at MorphoSource and are listed above in Table 1. Our phylogenetic scripts, including the matrix and full analytical settings, are available at: https://osf.io/9jac3/

## Acknowledgments

We acknowledge the European Synchrotron Radiation Facility for the provision of beam time on ID19 for proposals ES873 and LS3248 and Gideon Chinamatira and Kudakwashe Jakata for CT scanning at Wits. We thank Sifelani Jirah and Bernhard Zipfel (ESI), Zaituna Erasmus (Iziko), Rose Prevec (Albany Museum), Jennifer Botha and Elize Butler (formerly of National Museum, Bloemfontein), Carl Mehling (AMNH), Katherine Anderson (UWBM), Amanda Millhouse (NMNH), and William Simpson (FMNH), for collections access at their respective institutions. We are grateful for Cy Marchant’s assistance with the skull reconstruction of SAM-PK-K7710 and Ben Creisler for discussion on nomenclature. We thank the Willi Hennig Society for making TNT freely available. Lastly, we thank David Marjanović, Valentin Buffa, and Michel Laurin for their thoughtful, thorough, and encouraging reviews of this manuscript.

Author Contributions

XAJ designed and performed research, analyzed data, and wrote the paper. RBJB performed research, analyzed data, and wrote the paper. DPF designed research and wrote the paper. CB contributed data. VF and KD analyzed data and imaged specimens. TG and EG analyzed data. JC and BRP designed and performed research and wrote the paper.

## Funding

Grants from NRF AOP (#136516, 118794), the DSI-NRFCentre of Excellence in Palaeosciences, the BLM for computing, the AMNH Collection Study Grant Program, Biological Sciences Research Committee (ISU), and the ISU Graduate School for funding. This work was partially supported by NSF/GSA Graduate Student Geoscience Grant # 13090-21, which is funded by NSF Award # 1949901.

## Conflict of interest disclosure

The authors declare they have no conflict of interest relating to the content of this article.

## Index

**Figure S1.** BP/1/4203, articulated skeleton (A,B) and reconstruction (C,D) of *Milleropsis pricei* in dorsal (A,C) and ventral (B,D) views.

**Fig S2.** Tympanic fossa of SAM-PK-7710 (EAZ younginiform) (**A–B)**, *Milleropsis* (BP/1/720) (**C–D**), and *Milleretta* (BP/1/3822) (**E–F)** in posterior and posterolateral views, respectively. Abbreviations: q, quadrate; qj, quadratojugal; sb, stapedial boss; sq, squamosal; stp, stapes; and tc, tympanic crests.

**Figure S3.** Strict consensus phylogenetic tree of parsimony analysis of all characters within the matrix (A) and with temporal fenestration characters removed (B). Absolute (left) and present/contradicted (right) group bootstrap frequencies) and Bremer support values (below the branches)

**Figure S4.** Majority rule consensus tree from Bayesian analysis using the time-calibrated FBD model including nodes with posterior probability < 0.5)

**Figure S5.** Majority rule consensus tree from Bayesian analysis using the maximum clade credibility tree (Mkv) including nodes with posterior probability < 0.5).

**Video S1.** Digital rendering of *Milleropsis* BP/1/720, with the three skulls oriented horizontally.

**Video S2.** Digital rendering of *Milleropsis* BP/1/720, with the three skulls oriented vertically.

**Video S3**. Digital rendering of the articulated skeleton of *Milleropsis* BP/1/4203.

## Notes

https://www.morphosource.org/concern/media/000125334

https://osf.io/9jac3/

### Competing Interest Statement

The authors have declared no competing interest.

### Summary of Updates

I've removed line numbering and the title page for recomendation in a PCI journal

