## Supplementary figures and images for "Evolutionary assembly of crown reptile anatomy clarified by late Paleozoic relatives of Neodiapsida"

### Fig. S1

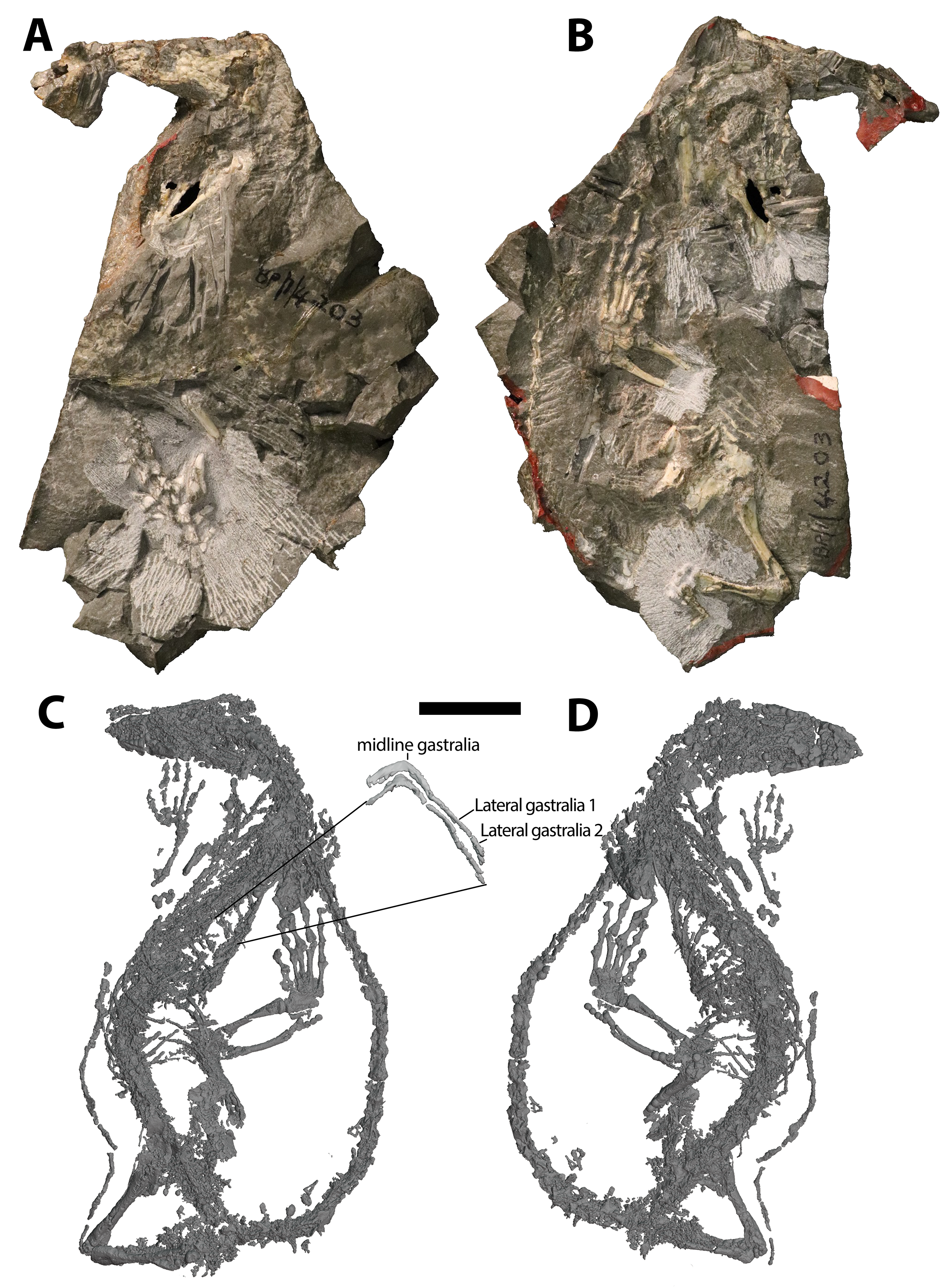

### Fig. S2

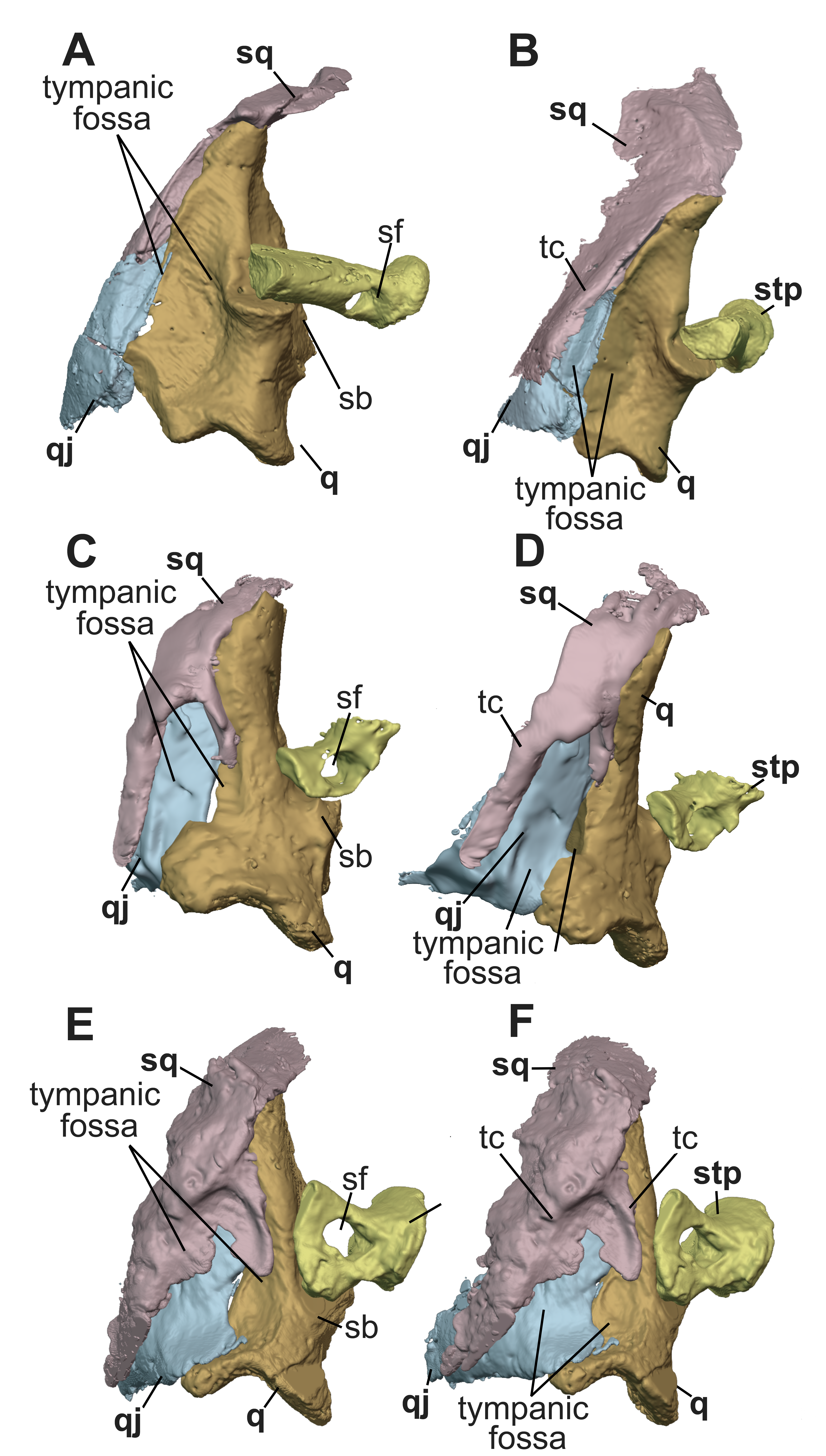

### Fig. S3

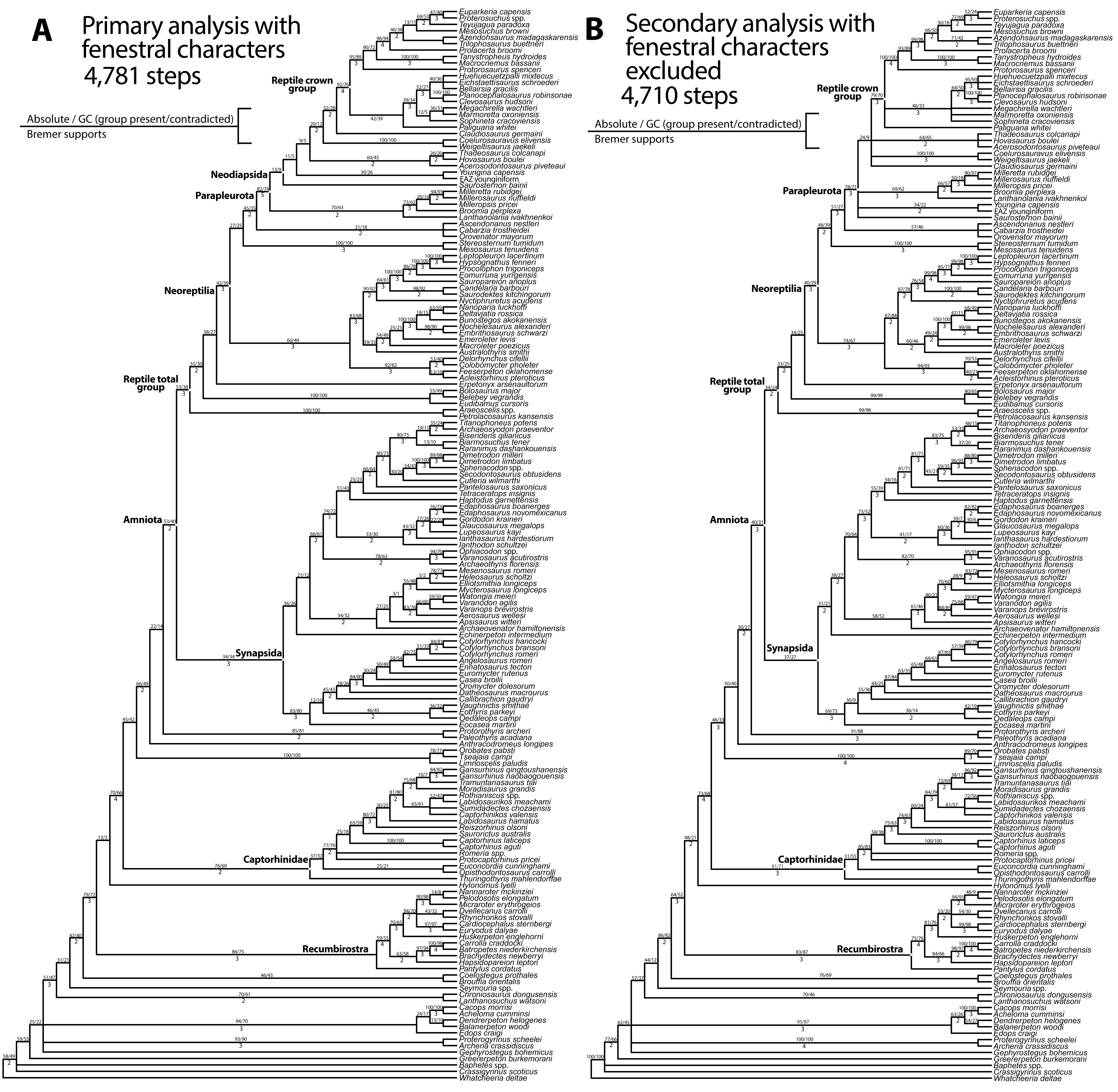

### Fig. S4

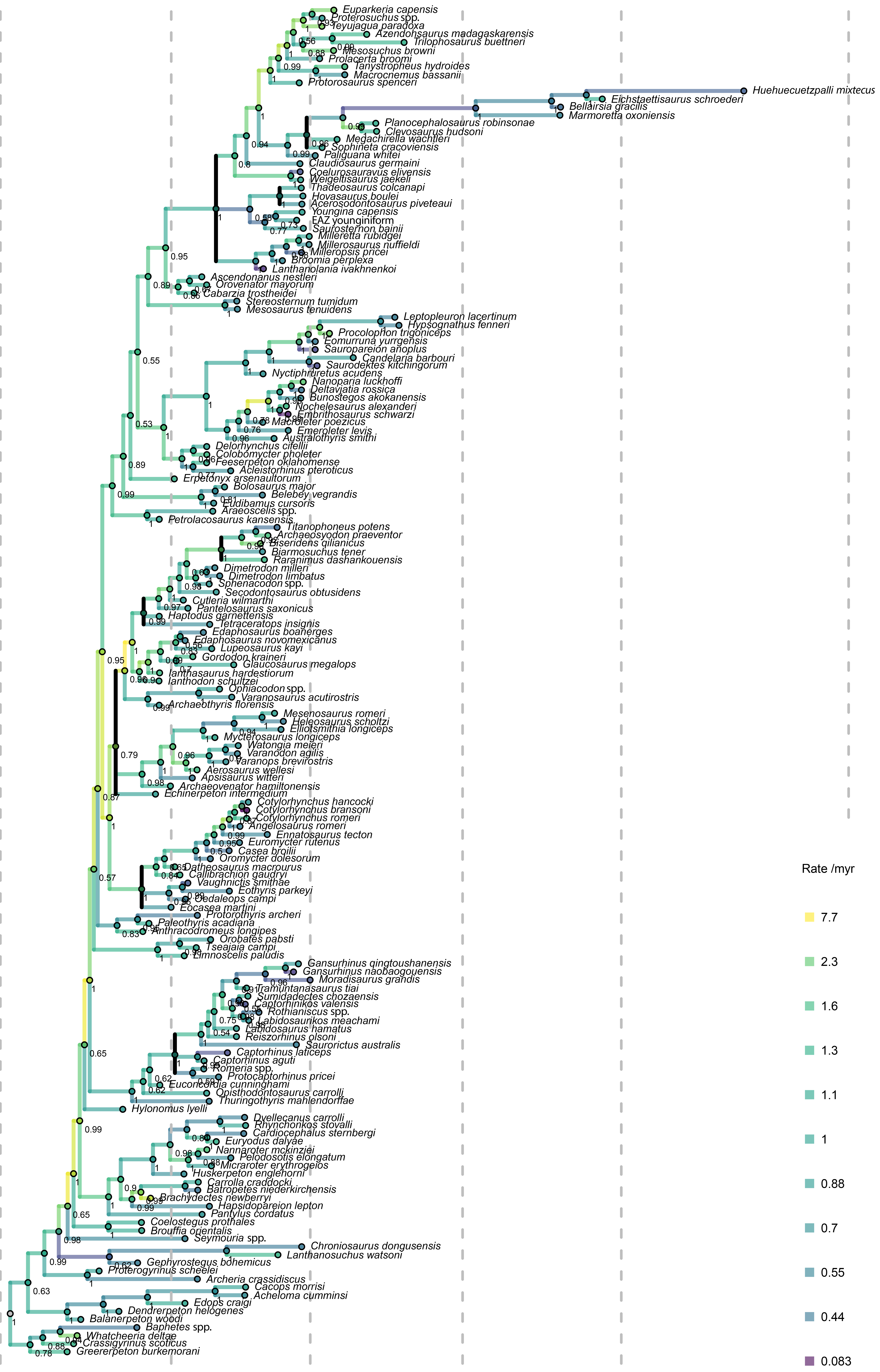

### Fig. S5

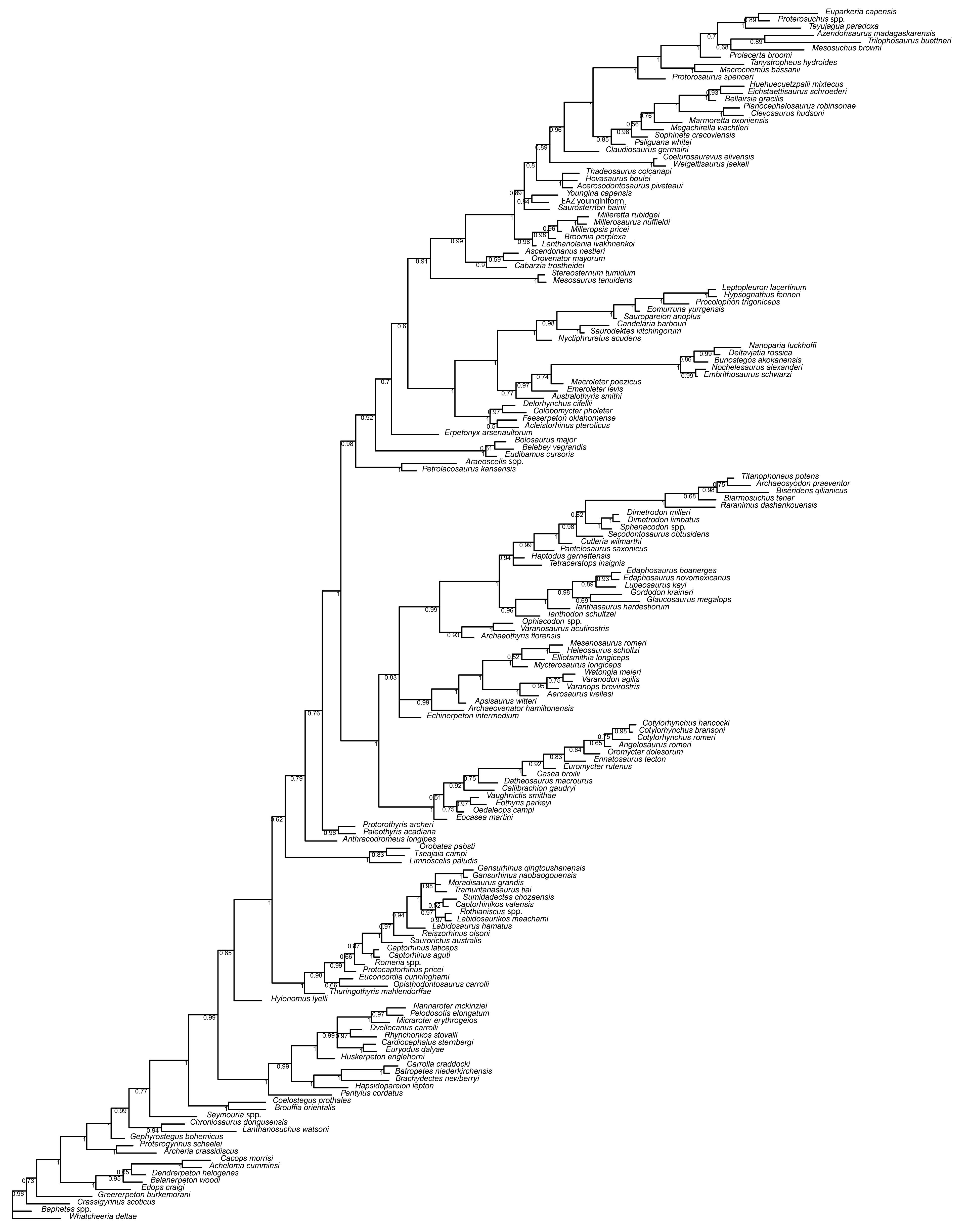
