## Supplementary material for "Evolutionary assembly of crown reptile anatomy clarified by late Paleozoic relatives of Neodiapsida": Supp Data

TABLE OF CONTENTS

1. **Systematic Paleontology, including definitions and diagnoses for Parapleurota and Neodiapsida.** Page 2.
2. **Brief description of the material**. Pages 3-6.
3. **Autapomorphies for select clades under different optimizations.** Pages 7-12.
4. **Autapomorphies of Parareptilia according to previous studies.** Pages 13-17.
5. **Features classically linking *Hylonomus* and Amniota.** Pages 18-21.
6. **Additional Steps required to support monophyly of Parareptilia, Eureptilia, and Diapsida, and the inclusion of Recumbirostra in Reptilia.** Pages 22-26.
7. **Code for analysis of stratigraphic fit.** Pages 27-29.
8. **Character additions and modifications.** Pages 30-196.
9. **Specimens under study.** Pages 197-208.
10. **Supporting video captions.** Page 208.
11. **Supporting references.** Page 209-223.

### <u>1. Systematic Palaeontology</u>

**PARAPLEUROTA** (this study)

**Etymology:** 'Parapleuros' and 'ota', derived from the Greek terms for "at or in the side of anything" and "ears", in reference to the presence of a tympanum on the posterolateral surface of the skull in these taxa, in stark contrast to the procolophonian 'otic notch' that extends onto the skull roof.

**Diagnosis**: Early parapleurotans are small-bodied Permian reptiles diagnosed by the following features: Postfrontal bearing a distinct posterodorsally directed process, loss of squamosal contact with the posteroventral margin of jugal, presence of a tympanum on posterior surface of squamosal, quadratojugal, and quadrate, anteroventrally directed basipterygoid processes, the loss of an ossified sphenethmoid, loss of squamosal contribution to quadrate foramen, basioccipital lacking a contribution to the fenestra ovalis, subequal heights of labial and lingual walls of alveolar groove of the maxilla, possessing fewer than 20 dorsal vertebrae, the loss of atlantal ribs, bearing fewer than two lateral gastralia per row, radius between .68 and .82 length of humerus, humerus with intermediate torsion, a perforating foramen of the tarsus located at midlength of proximal tarsals, and the proximal expansion of fifth metatarsal that articulates with both distal tarsal V and IV.

**NEODIAPSIDA** Benton, 1985

**Revised definition:** All taxa closer to *Youngina capensis* Broom 1914, *Claudiosaurus germaini* Carroll 1981, and *Lacerta agilis* Linnaeus 1758 than to *Orovenator mayorum* Reisz et al. 2011, *Milleretta rubidgei* Broom [1938], *Procolophon trigoniceps* Owen 1876, *Bolosaurus striatus* Cope 1978, *Mesosaurus tenuidens* Gervais 1865, and *Petrolacosaurus kansensis* Lane 1945.

**Rationale:** The clade Neodiapsida (Benton 1985) has two definitions in use today, the node-based definition of Laurin (1991) "The most recent common ancestor of younginiforms and living diapsids, and all its descendants" that references younginiforms, a non-monophyletic group of non-saurian diapsids; and the stem-based definition of Reisz et al. (2011) "*Youngina capensis* Broom, 1914 and all species more closely related to it than to *Petrolacosaurus kansensis* Lane, 1945," which is broadly equivalent to Neoreptilia of Ford and Benson (2019), and would require that several unrelated 'parareptile' groups be referred to Neodiapsida. Our proposed strict definition specifies the exclusion of several reptile taxa in hopes that, even if a taxonomic reshuffling were to occur in later studies, the usage of Neodiapsida the closest immediate relatives of the reptile crown group of the latest Permian, will be maintained.

### 2. Brief description of the material.

**Systematic paleontology.** Amniota Haeckel, 1866; Sauropsida Huxley, 1864; Neoreptilia Ford and Benson, 2019; Parapleurota, this study; Millerettidae Camp, 1945, *Milleropsis* Gow, 1972; *Milleropsis pricei* [Watson, 1957].

**Material:** BP/1/720 (holotype), a nodule containing at least nine partially articulated individuals based primarily on the number of exposed lower jaws and maxillae (Fig. 2, Videos S1-2). BP/1/4203 (referred specimen), nearly complete, articulated skeleton (Fig. S1, Video S3).

**Locality and Horizon:** The *Milleropsis* specimens come from the Middle *Cistecephalus* Assemblage Zone at Swaelkrans, Murraysburg, South Africa (BP/1/720) and Nieu Bethesda, Graaff-Reinet, South Africa (BP/1/4203).

*Milleretta* Broom 1948; *Milleretta rubidgei* [Broom, 1938].

**Material:** BP/1/3822 (referred specimen), a nearly complete skull of a juvenile individual lacking the dorsal surface of the antorbital snout (Fig. 3).

**Locality and Horizon:** Upper *Cistecephalus* Assemblage Zone of the Karoo Basin from Doornplass, Eastern Cape, South Africa.

Detailed anatomical descriptions on the cranial anatomy of these two taxa are published inRoyal Society Open Science (Jenkins et al. 2025a) and the Zoological Journal of the Linnean Society (Jenkins et al. 2025b). We provide a brief summary of some important findings below.

**Skull.** The cranial anatomy of millerettids is remarkably similar to that of neodiapsids (Gow 1974), including crown reptiles, sharing key saurian traits that are absent in traditional 'eureptiles' such as captorhinids (Heaton 1979) and araeoscelidians (Vaughn 1995, Reisz 1981). Many of these similarities were obfuscated by the extremely modified and derived anatomy of *Milleretta rubidgei*, which bears cranial osteoderms that obscure sutures of the skull roof and antorbital skull. The anteroposteriorly reduced lacrimal of both *Milleropsis* and *Milleretta* does not reach the external naris, a feature shared with neodiapsids (Laurin and Reisz 1991, Pritchard et al. 2021)(Figs. 2–3). This contrasts with the elongated lacrimal of early amniotes including captorhinids and araeoscelidians such as *Petrolacosaurus* (Reisz 1981) in which the lacrimal forms nearly the entire posterior margin of the external nares. The millerettid skull lacks an upper temporal fenestra (UTF), a synapomorphy of Neodiapsida. Instead, the parietal possesses a ventrolateral flange that rests in a groove in the postorbital and squamosal that develops during ontogeny, such that juvenile individuals bear an unossified gap in this region. This feature was previously interpreted as a secondary closure of the UTF in the 'Millerosauria', a clade proposed to include millerettids and mesenosaurine varanopids (Watson 1957).

*Milleropsis* possesses a lower temporal emargination framed by the jugal, postorbital, squamosal, and quadratojugal (Fig. 2), whereas this opening is secondarily closed through ontogeny in *Milleretta* (Fig. 3). The jugal of *Milleropsis* possesses a posteriorly directed spur that does not form a lower temporal bar, similar to the middle Permian reptile *Lanthanolania* (Modesto and Reisz 2004) and early crown reptiles. The squamosal of millerettids lacks an occipital shelf, exposing the quadrate in lateral view, like neodiapsids (Laurin and Reisz 1991), but unlike earlier 'eureptiles' such as protorothyridids in which the quadrate is hidden by this broad shelf of the squamosal (Carroll 1969). The squamosal of millerettids does not contact the quadrate ramus of the pterygoid, a feature shared with neodiapsids such as SAM-PK-K7710 and *Prolacerta* (BP/1/5880) but in stark contrast to typical ‘parareptiles’ such as procolophonids, and other early amniotes. The quadrate of millerettid stem reptiles lacks the stapedial recess seen in many early amniotes such as captorhinids (Heaton 1979) early synapsids (Romer and Price 1940), and araeoscelidian reptiles (Peabody 1952), and instead bears a distinct stapedial boss that likely ligamentously or cartilaginous extension of the stapes, as in neodiapsids like SAM-PK-K7710, *Claudiosaurus* (Carroll 1981) and *Acerosodontosaurus* (pers obv. MNHN.F.MAP359a) (Fig. 4). The posterolateral surfaces of the squamosal, quadratojugal, and quadrate in both *Milleropsis* and *Milleretta* bear a tympanic fossa that is confluent with ventrolateral fossa on quadrate medially. These are similar to the tympanic fossae/emarginations that span the quadratojugal and quadrate in early-diverging neodiapsids such as *Thadeosaurus* and SAM-PK-K7710 (Fig. 4) and the squamosal, quadratojugal and quadrate in early saurians, but contrast with procolophonian reptiles that bear a tympanic emargination that extends onto the supratemporal (e.g., *Saurodektes*, BP/1/3189) . The quadrate foramen in both taxa is formed by the quadratojugal and quadrate, similar to crown reptiles but unlike early amniotes in which the squamosal contributes to this foramen (Fig. 5).

**Palate.** The palates of millerettid and crownward stem reptiles are remarkably similar (Figs. 2–3). The quadrate ramus of the pterygoid bears strongly developed tympanic flanges contributing to the cavitation of the middle ear (Fig. 6), a feature present in saurians and other neoreptiles[35] but poorly developed in araeoscelidians (AMNH FARB 4686) and varanopids such as *Heleosaurus* (SAM-PK-K8305) and absent in captorhinids (Heaton 1979) and recumbirostran ‘microsaurs’ (Szostakiwskyj et al. 2015). As in crown reptiles, the basipterygoid recess of *Milleropsis* and *Milleretta* is formed entirely by the pterygoid (Fig. 6), differing markedly from earlier-branching amniotes such as araeoscelidians (Reisz 1981) the varanopid *Heleosaurus* (SAM-PK-K8305), and ‘parareptiles’ such as *Delorhynchus* (Reisz and MacDougall 2014) in which the epipterygoid contributes to the basipterygoid recess. This derived anatomical arrangement is an important adaptation that may have promoted the appearance of the kinetic palatoquadrate seen in extant saurians, whereas the plesiomorphic condition restricts movement at this joint (SI Appendix, Fig. S3).

The ectopterygoid of millerettids possesses a posterolateral process that is emarginated by the adductor fossa, as in *Orovenator* (Ford and Benson 2018) and neodiapsids (Hunt et al. 2023), but unlike araeoscelidians, acleistorhinids, and recumbirostrans in which the pterygoid excludes the ectopterygoid from the adductor fossa. It is completely edentulous, unlike the denticulate ectopterygoids of earlier-diverging reptiles such as *Petrolacosaurus* (Reisz 1981), *Orovenator* (Ford and Benson 2018) or acleistorhinid ‘parareptiles’ (Reisz and MacDougall 2014). The vomers bear enlarged vomerine ‘fangs’ anteriorly (Gow 1972). The millerettid palatine bears a dorsal eminence that contacts the prefrontal and contributes to the foramen orbitonasale, a trait previously considered diagnostic of ‘Parareptilia’ (Laurin and Reisz 1991) but that we find to be widespread among stem-reptiles including the EAZ younginiform (SAM-PK-K7710, Fig. 5) and the archosauromorph *Prolacerta* (BP/1/5880). The pterygoid of *Milleropsis* and *Milleretta* bears three rows of dentition; one on the transverse flange, one that extends anterolaterally onto the palatine, and another on the palatal process. The dentition on the transverse flange consists of a single large row with two rows of smaller teeth anteriorly, contrasting with neodiapsids other than the EAZ younginiform, which also bears two rows of teeth on the transverse flange of the pterygoid. A large suborbital foramen is present between the maxilla, ectopterygoid, and palatine, although it is smaller than the suborbital fenestra of neodiapsids.

***Braincase.*** The millerettid stapes is extremely simplified relative to earlier-diverging reptiles, lacking a dorsal process or blade-like quadrate process, similar to all neodiapsids but in contrast to earlier diverging taxa such as captorhinids, araeoscelidians, and many 'parareptiles'. The distal end of the stapes ends freely, and is visible in lateral view where it is directed towards the tympanic fossa. This indicates that the stapes of millerettids and neodiapsids lost the bracing function between the suspensorium and braincase that is present in early amniotes (Fig. 5). The parabasisphenoid of millerettids bears dentition on its ventral plate, a feature once considered to be absent in Neodiapsida that is indeed present in SAM-PK-K7710 and *Youngina* (Hunt et al. 2023). The abducens nerve (CN VI) in millerettids travels through the clinoid process, features we document in the early neodiapsid SAM-PK-K7710 and the archosauromorph *Prolacerta* BP/1/5880 that are absent in more basally-diverging reptiles such as *Araeoscelis* (AMNH FARB 4686, Table S1). The paroccipital process of the opisthotic makes lateral contact with the quadrate. The prootic bears a distinct floccular fossa, the earliest reported occurrence of this feature in a non-crown reptile (Fig. 7). However, the prootic of millerettids lacks a crista prootica and an anterior inferior process, in contrast to saurians but as in many non-saurian reptiles, including *Araeoscelis* (AMNH FARB 4686) and *Youngina* (BP/1/2871). The ventral process of the prootic frames the opening for the trigeminal nerve, CN V, and bears a ventral notch for the facial nerve, CN VII. The medial wall of the prootic is well-ossified for the pathways of the semicircular canals (Figs. 2–3). The semicircular canals are well-developed and arcuate. The vestibule and lagenar recess are smaller than in captorhinids (Bazzana et al. 2022) and are extremely similar to the endosseous labyrinth of both neodiapsids like *Youngina* and saurians such as the archosauromorph *Prolacerta*. A sphenethmoid is absent in all known millerettids, a feature shared with Neodiapsida, including the reptile crown.

***Lower Jaw.*** The lower jaw of *Milleropsis* is extremely slender and bears 23 alveoli, whereas that of *Milleretta* is shorter and bears 15 tooth positions (SI Appendix, Fig. S2). The implantation of the marginal dentition is subthecodont, and the lingual and labial walls have similar dorsal extents. A single, edentulous coronoid is present in millerettids, in contrast to the two coronoids present in early amniotes such as the varanopid *Heleosaurus* (SAM-PK-K9305) and the araeoscelidian *Araeoscelis* (AMNH FARB 4686) and the denticulated coronoids of some ankyramorphan 'parareptiles' (Haridy et al. 2017). The mandibular symphysis is formed solely by the dentaries, which exclude the splenial from the symphysial facet. The Meckelian canal pierces the anterior end of the dentary, similar to non-saurian neodiapsids scanned here but in contrast to most crown reptiles (Fig. 8). The surangular of *Milleropsis* bears both an anterior and posterior foramen (a feature previously only reported in neodiapsids, e.g., Ezcurra 2016) and a dorsomedial shelf. The angular is exposed broadly in lateral view. The retroarticular process is well-developed and upturned dorsally. A postsplenial is absent, as in all amniotes except possibly *Petrolacosaurus*[31].

***Postcranial skeleton.*** Synchrotron tomography of a complete, articulated postcranium of *Milleropsis* (BP/1/4203; Fig. S1, Video S3), reveals a postcranial skeleton remarkably similar to neodiapsids[6]. The lateral surface of the neural arches of dorsal vertebrae are weakly concave and not swollen, unlike the condition in many 'parareptiles' or diadectomorphs (Sumida 1989). The femora are relatively gracile compared to early amniotes, lacking both fourth and lesser trochanters and bearing reduced intertrochanteric and intercondylar fossae. The caudal ribs are fused to the centrum and are laterally directed, increasing the surface area for the origination of the caudofemoralis musculature, also present in neodiapsids including the reptile crown group (Currie 1981), but are unlike the hook-like caudal ribs that are only sutured to the centrum of early amniotes, such as the owenettid *Saurodektes kitchingorum* (Reisz and Scott 2002). Gastralia are abundant and consist of a midline element and two lateral elements per row (Fig. S1), a feature uniquely shared with neodiapsids such as *Claudiosaurus* (Carroll 1981) but unlike earlier diverging reptiles including mesosaurids and araeoscelidians that possess upwards of four rows and no midline element. A single coracoid is present, as in neodiapsids but unlike the two coracoids of early ‘eureptiles’ such as captorhinids and araeoscelidians and ‘parareptiles’ such as procolophonians (Ford and Benson 2019) The posterior process of the ilium is well-developed and posteriorly directed. A distinct fifth distal tarsal of the pes is absent and possibly fused with DT4, and the unhooked 5th metatarsal bears a distinct proximal expansion for articulation with the fourth distal tarsal, similar to the outer process of MtV seen in neodiapsids (Pritchard et al. 2015).

### 3. Autapomorphies of select clades in this study under different optimization regimes

U = unambiguous, D = deltran, A = acctran

**Autapomorphies of Neodiapsida (*Saurosternon* and other neodiapsids)**

U. Caudal ribs or transverse processes present on 13 or more caudal vertebrae (495: 0 → 3)
U. Mineralized sternum (535: 0 → 1).
U. Pisiform much larger than fourth distal carpal (563: 0 → 1)

**Synapomorphies of Younginidae and more crownward neodiapsids**
U. Anterior centrodiapophyseal lamina present on dorsal vertebrae (Ch. 471: 0 → 1).

**Autapomorphies of 'Tangasauridae' + Weigeltisauridae + *Claudiosaurus* + crown reptiles**
U. Frontal less than 1.5 times parietal length (109: 1→ 0).
U. Parabasisphenoid lacks dentition on ventral plate (257: 0 → 1)
U. Parabasisphenoid lacks dentition on cultriform process (259: 0 → 1).
U. MC IV is shorter than MC III (568: 0 → 1).
U. MC V greater than .65 the length of MC IV (569: 1 → 0).
U. Acetabulum not invaded by bone on the posterodorsal margin (596: 0 → 1)

**Synapomorphies of Weigeltisauridae + *Claudiosaurus* + the reptile crown group**
U. Supratemporal fossa present (6: 0 → 1).
U. Lingual wall of maxilla extends ventrally less than the labial wall (80: 0 → 1).
U. Facet for proatlas absent on exoccipital (241: 0 → 1).
U. Stapedial foramen absent (361: 0 → 1)
U. Stapes elongate, columella more than 4x as long as the height of the footplate (366: 0 → 1)
U. Ventral surface of dorsal vertebrae rounded (463: 1 → 0)
U. Chevrons taper at their distal extremity (492: 1 → 2)
U. Ventromedial plate of clavicle forms a highly obtuse angle with the shaft (503: 0 to 1)
U. Sternum no longer mineralized (535: 1 to 0).

**Synapomorphies of *Claudiosaurus* + the reptile crown group**
U. External nares located close to midline (85: 0 → 1)
U. Parietal with broad lateral embayments (177: 1 → 2)
U. Postparietals absent (208: 0 → 1)
U. Neural spines of caudal vertebrae moderately tall (491: 0 → 1)
U. Scapula lacks a distinct groove for a cleithrum (523: 0 → 1)
U. Tibioastragalar joint flat (618: 1 → 0)
U. MTIV long, at least 40% of digit IV (638: 0 → 1)

**Autapomorphies of the reptile crown-group**
U. Posterior margin of quadrate concave (351: 0 → 1)
U. Loss of stapedial boss of quadrate (353: 1 → 2)
U. Surangular lacks an anterodorsal process (431: 1 → 0)
U. Radiale not wider than long (558: 0 → 1)
U. Width of carpus is shorter than length of MCIV (560: 0 → 1)
U. DT4 bears a prominent posterior process (631: 0 → 1)
U. Fifth distal tarsal absent (633: 0 → 1)
U. MTV reduced in length relative to MTIV (639: 0 → 1)
U. MTV hooked (643: 0 → 1)

**Autapomorphies of Parapleurota (millerettids and neodiapsids from this study)**
U. Lacrimal does not contribute to external naris (89: 1→ 2).
U. Posterodorsal process of postfrontal (119: 1 → 0)
U. Quadratojugal does not contact the posteroventral margin of jugal (155: 0 → 1).
U. Squamosal superficial contact with tabular absent (160: 1 → 0).
U. Quadrate exposed in lateral view via a slender descending process of SQ (164: 0 → 1)
U. Tympanic fossa on posterior surface of QJ, and SQ (165: 0 → 1)
U. Loss of sphenethmoid (283:1→ 0)
U. Small tympanic fossa on quadrate (351: 0 → 1).
U. Lingual wall of dentary subequal in height to labial wall (403: 1 → 0)
U. Splenial contacts posterior coronoid (413. 0 → 1).
U. Anterior surangular foramen present (427: 0 → 1)
U. Posterior surangular foramen present (428: 0 → 1)
U. Fewer than 20 dorsal vertebrae (443: 0 → 1).
U. Atlantal centrum separate from axial intercentrum (447: 1 → 0).
U. Low caudal neural spines (491. 0 → 1).
U. Fewer than two lateral gastralia per row (399: 2→ 3)
U. Midline gastralia element present (501: 0 → 1)
U. Hindlimb longer than 90% of anteroposterior length of trunk (603: 0 → 1)
U. Fourth distal tarsal articulates with MTV (633: 0 → 1)
U. Metatarsals overlap (637: 0 → 1).
U. Proximal expansion of fifth metatarsal (644: 0→ 1)
A. Basioccipital loses contribution to fenestra ovalis (249: 0 → 1)
A. Loss of squamosal contribution to quadrate foramen (359:0→ 1)
A. CN VI pierces clinoid processes (278: 0 → 1).

**Autapomorphies of Millerettidae**
U. Ventral plate of parabasiphenoid flat (264: 1 → 0),
A. Dorsal process of QJ expanded and is nearly as tall as skull height (149: 0 → 1).
A. Parietal with ventral flange contributing to lateral surface of skull (180: 0 → 1).
A. Anterior vomerine buttress bearing enlarged vomerine teeth (305: 0 → 1).
A. Distal end of stapedial columella larger than footplate (460:0 → 1)
A. CN VII present as a notch within prootic. (234: 0 → 1).
A. Exoccipital fused to basioccipital. (245 → 0 to 1).
D. Row of 3-6 teeth on crista ventrolateralis of parabasisphenoid (259:1 → 2)
D. Lateral tooth row of vomer absent (304: 0 → 1)

**Synapomorphies of Parapleurota + (*Orovenator* + *Ascendonanus* + *Cabarzia*)**
U. Base of teeth not covered by cementum or bone of attachment (26: 1 → 2)
U. Row of supralabial foramina present on maxilla (56: 0 → 1)
U. Lingual wall of maxilla subequal in height to labial wall (80: 1 → 0)
U. Lacrimal contribution to orbit small or absent (91: 1 → 2)
U. Lateral expansion of prefrontal (101: 0 → 1)

U. Posterodorsal process of squamosal present (159: 0 → 1)
U. Slender supratemporal in squamosal or parietal trough (181: 0 → 1)
U. Carotid arteries within lateral wall of braincase (260: 0 → 1)
U. Teeth present on ventral plate of parabasisphenoid(257: 0 → 1)
U. Palatal process of pterygoid extends anterior to palatine (320: 0 → 1)
U. Arcuate flange of pterygoid well-developed (322: 1 → 0)
U. Anteriormost dentary teeth oriented vertically (399: 0 → 1)
U. Loss of posteroventromedial process of coronoid (420: 1 → 0)
U. Medial expansion of the surangular dorsal surface present (429: 0 → 1)
U. Caudal transverse processes on five or fewer vertebra (495: 1 → 2)
U. Loss of clavicle-clavicle contact (505:0 → 1)
U. Short MCV relative to MCIV (569: 0 → 1)
U. MTIV much longer than MT1 (636: 0 → 1)
A. Loss of epipterygoid contribution to basal articulation (288: 0 → 1)
D. Strut-like ascending process present on medial surface of maxilla (69. 0 → 1).

**Synapomorphies of Mesosauridae + *Orovenator* + Parapleurota**
U. Marginal dentition moderately recurved (8: 0→ 1)
U. Occipital shelf of squamosal poorly developed exposing quadrate posteriorly (156: 0 → 1)
U. Single row of teeth of transverse flange (315: 1 → 2)
U. Stapes footplate reduced relative to otic capsules (363: 0 → 1)
U. Surangular-dentary suture longer than angular-dentary suture (400: 0 → 1)
U. Single coracoid ossification (534: 1 → 0)
U. Femoral condyles extend equally distally (604: 0 → 1)
U. Distal adductor crest of femur poorly developed (604: 0 → 1)
A. Squamosal does not contact pterygoid (162: 1 → 0)
A. Supraglenoid foramen of scapula absent (519: 0 → 1)

**Autapomorphies of Neoreptilia**
U. Stapedial boss of quadrate present (353: 0 → 2)
U. Poorly developed capitellum and trochlea of humerus (539: 0 → 1)
U. Entepicondylar foramen of humerus present (549: 0 → 1)
U. Olecranon process absent or low (560: 2 → 0)
U. Overlapping metatarsals (637: 0 → 1)
A. Large anterior maxillary foramen (57: 0 → 1)
A. Caniniform teeth absent (62: 1 → 0).
A. Loss of ventromedial process of lacrimal (92: 1 → 0)
A. Supratemporal exceeds tabular posteroventrally (184: 0 → 1)
A. Presence of retroarticular process of mandible (379: 0 → 1)
A. Cnemial crest of tibia low or absent (619: 1 → 0)

D. Holocephalous cervical ribs (459: 1 → 0)
D. Single-headed dorsal ribs (474: 0 → 1)
D. Sacral ribs subequal in size (486: 0 → 1)

**Autapomorphies of Reptile Total-Group**
U. Prefrontal-palatal contact (97: 0 → 1)
U. Skull roof lacks fossae for epaxial muscular insertion (193: 1 → 0)
U. Meckelian canal pierces anterior end of dentary (397:0 → 1)
U. Caudal neural spines moderately tall (491: 0 → 1)
U. Convex anterior margin of scapula (519: 0 → 1)
U. Transversely narrow supinator process of humerus (553: 0 → 1)
U. Single pedal centrale (Ch. 634: 0 → 1)
A. Suborbital foramen present (345: 0 → 1)
A. Stapes rod-like (361: 1 → 0).

**Autapomorphies of Synapsida**
U. Maxillary canal possesses well-defined external nasal canals (70: 0 → 1)
U. Supraoccital bears a process framing the floccular fossa (205: 0 → 1)
U. Opisthotic fused with prootic in mature individuals (221: 0 → 1)
U. Otic trough of opisthotic present (223: 0 → 1)
U. Ventral end of PSC separated from utricular region by bone (226: 2 → 1)
U. CN V located dorsally on prootic and framed by an infratrigeminal process (233: 0 → 1)
U. Suture between basioccipital and parabasisphenoid non-overlapping (248: 0 → 1)
U. Posteroventral emargination on main body of parasphenoid (263: 0 → 1)
U. Braincase sutured between otic capsules and sphenoid in mature individuals (273: 0 → 1)
U. Columella of epipterygoid sheetlike (293: 0 → 1)
U. Palatine only weakly incised by choana (309: 0 → 2)
U. Medially projecting flange of prearticular ventral to adductor fossa (436: 0 → 1)
U. Distal head of humerus wide compared to length of shaft (445: 1 → 0)
U. Ventral surface of humeral entepicondyle flat or weakly convex (447: 1 → 0)
U. Pubis longer than acetabulum (582: 0 → 1)
U. Femora short and broad (602: 1 → 0)
U. Fourth trochanter present (603: 0 → 1)
A. Alveolar shelf of maxilla taller than mediolateral width of tooth (23: 0 → 1)
A. Four or fewer premaxillary teeth (30: 0 → 1)
A. Squamosal does not contact posterodorsal margin of jugal (151: 0 → 1)
A. Postparietals fused as a single median element (209: 0 → 1)

**Autapomorphies of Amniota**
U. Lacrimal with moderate contribution to the orbital margin (91: 0 → 1)
U. Posterior process of postorbital narrow (130: 0 → 1)
U. Jugal possesses a free posterior margin (133: 0 → 1)
U. Squamosal does not contact posterodorsal margin of jugal (151: 0 → 1)
U. Low lingual wall of dentary (403: 1 → 0)
U. Dorsal neural arches fused with centrum (477: 1 → 2)
A.Postorbital extends more than half temporal midlength (129: 0 → 1)
A. Paroccipital process contacts suspensorium laterally in mature individuals (217: 1 → 0)

A. Palatal process of pterygoid terminates posterior to anterior margin of palatine (320: 1 → 0)
A. Neural spines of anterior dorsal vertebrae sub-triangular (465: 0 → 1)

**Synapomorphies of Amniota + Protorothyrididae**
U. Squamosal contacts posteroventral margin of jugal (152: 0 → 1)
U. MC IV longer than MC III (568: 0 → 1)
U. Calcaneum much longer than wide (624: 0 → 1)
A. Posterior cervical and anterior dorsal ribs not expanded distally (462:1 → 0)

**Synapomorphies of Amniota + Protorothyrididae + *Anthracodromeus***
U. Postparietals entirely on the occipital surface of skull (211: 2 → 3).
U. 21 or more dorsal vertebrae (443: 1 → 0)
U. Distal head width less than .40 total humus length (545: 0 → 1)
U. Femur long and slender (602: 0 → 1)

**Synapomorphies of Amniota + Protorothyridae + Diadectomorpha**
U. Prefrontal-palatine contact absent (97: 1 → 0)
U. Frontal 1.5-2.5 times parietal length (109: 0 → 1)
U. Anterior process of frontal longer than posterior process (112: 0 → 1)
U. Asymmetric vomerine processes (298: 0 → 1)
U. Ridge or keel present on ventral surface of angular (421: 0 → 1)
U. Ectepicondylar foramen not enclosed (552: 1 → 2)
A. Posterior cervical and anterior dorsal ribs not expanded distally (462:1 → 0)

U. Maxillary caniniform teeth at least 1.5 times the size of surrounding dentition (64: 0 → 1)
U. Prefrontal does not contact postfrontal (97: 0 → 1)
U. Supratemporal slender and in parietal trough (184: 0 → 1)
U. Paroccipital process ends freely (217: 0 → 1)
U. Ectopterygoid makes lateral contact with jugal (338: 0 → 1)
U. Pterygoid ramus of quadrate anteroposteriorly short (355: 0 → 1)
U. Single coronoid in mandible (417: 1 → 2)
U. Coronoid dentition absent (419: 0 → 1)
U. Manual unguals taper distally (576: 0 → 1)

**Synapomorphies of Amniota + Protorothyridae + Diadectomorpha + Captorhinidae**
U. Bilateral embayment of skull roof (5: 0→ 1)
U. Tabular small and wedge-shaped (190: 0 → 1)
U. Postparietals with narrow contribution to skull roof (211: 1→ 2)
U. Medial tooth row of vomer present (303: 0 → 1)
U. Discrete anterolateral tooth row ‘t2’ present on pterygoid (317: 0 → 1)
U. Atlanta centrum fused to axial intercentrum (447: 0 → 1)
U. Two sacral vertebrae (489: 0 → 1)
U. Medial expansion on dorsal surface of iliac blade (591: 0 → 1)
U. Postacetabular process extends behind iliac body (593: 0 → 1)

U. Proximal phalanges similar in length to metatarsals (646: 0 → 1)
U. Pedal unguals much longer than non-ungual phalanges (647: 0 → 1)

**Synapomorphies of Amniota + Protorothyridae + Captorhinidae + Diadectomorpha + *Hylonomus***
U. Postparietals contribute to posttemporal fenestra (212: 0 → 1)
U. Transverse flange of pterygoid extends ventral to cheek in lateral view (328: 0 → 1)
U. Astragalus present (620: 0 → 1)
D. Arcuate flange of pterygoid present (322: 0→ 1)
A. Postorbital contact with parietal (127: 0 → 1)

**4. Autapomorphies of Parareptilia according to previous studies.**

Previous studies have found several autapomorphies for ‘Parareptilia’, including 14 given by Laurin and Reisz (1995), which were reduced to seven by deBraga and Reisz (1996). Many of these characters have been subsequently demonstrated to be present in other early amniote taxa (e.g., Carroll 1989,

Lee 2013, Spindler et al. 2018, 2019, Ford and Benson 2019). We find that none of the following characters are valid as unambiguous autapomorphies unique to ‘Parareptilia’, but are better interpreted as autapomorphies for smaller groups of ‘parareptiles’, notably Ankyramorpha, a clade consisting of procolophonians and acleistorhinids (deBraga and Reisz 1996). We describe the occurrence of these characters in other early tetrapods, notably other early reptiles, or their absence in certain ‘parareptiles’ below. These characters are included in our dataset, except for the presence of a foramen orbitonasale, a character that is invariably linked with the presence of a broad prefrontal-palatal contact within this current taxon sampling (Lee 1993).

**A) Prefrontal-palatine contact present.**

Prefrontal-palatine contact is a noted autapomorphy of ‘Parareptilia’, first acknowledged in a cladistic context by Laurin and Reisz (1995, Ch. 6), although this study correctly noted that this feature has a wider distribution among early tetrapods, being present in captorhinids, diadectids, and *Seymouria*. Prefrontal-palatine contact is also present in recumbirostrans (Gee et al. 2019a), edaphosaurids (Modesto 1995), varanopids (Berman and Reisz 1992), and the early reptile *Orovenator* (Ford and Benson 2018). Indeed, the ventral process of the prefrontal in the neodiapsid *Youngina* (pers. obv. synchrotron scans of BP/1/2871) makes broad contact with an ascending process of the palatine. This contact is also present in *Araeoscelis* (pers. obv. scans of AMNH FARB 4686), which possesses a broad medial flange of the prefrontal. Prefrontal-palatine contact is present in early saurians including rhynchocephalians (e.g., *Clevosaurus*, Fraser 1982) and rhynchosaurs (Dilkes 1998). Misscorings of this trait as ‘absent’ in previous ‘parareptile’ matrices (e.g., MacDougall et al. 2019, Cisneros et al. 2020) were likely driven by the absence of scan data in many Paleozoic reptiles at the time - the contact between the prefrontal and palatine is often obscured by matrix or by the maxilla and lacrimal in lateral view, and the ventral process of the prefrontal can be extremely narrow. Prefrontal-palatine contact is therefore probably a reptile, or even amniote, plesiomorphy.

**B) Foramen orbito-nasale enclosed between prefrontal, lacrimal and palatine.**

The foramen orbitonasale which pierces the prefrontal, lacrimal, and palatine, is noted as a autapomorphy of Parareptilia by Laurin and Reisz (1995, Ch. 10) and subsequent studies, although the contribution of the lacrimal to this foramen is absent in most non-bolosaurid or procolophonoid ‘parareptiles’ such as *Milleropsis* (BP/1/720). A foramen orbitonasale (or a foramen topologically similar) pierces the antorbital wall or the medial margin of the lacrimal in captorhinids (Heaton 1979), edaphosaurids (Modesto 1995), and the neodiapsid *Youngina* (pers. obv. synchrotron scans of BP/1/2871), where it is present between the dorsal process of the palatine and the prefrontal as in millerettids. Lee (1995) also noted that within the matrices of Laurin and Reisz (1995), the presence of an orbitonasal foramen is non-independent with the presence of a broad prefrontal-palatal contact, he advocated for the removal of this character in amniote phylogenies. A similar foramen is expressed in rhynchosaurs (Gentill and Ezcurra 2022), which also possess a broad prefrontal-palatine contact, as well as *Prolacerta*, which bears a more narrow prefrontal-palatine contact (Sobral 2023). However, we note that not all taxa with a broad prefrontal-palatine contact bear a foramen orbitonasale, such as recumbirostrans (e.g., *Hapsidopareion*, Jenkins et al. 2025).

**C) Anterior lateral maxillary foramen distinctly larger than other foramina**

An enlarged foramen on the maxilla was found to be an autapomorphy of Parareptilia by both Laurin and Reisz (1995, Ch. 20) and deBraga and Reisz (1996). An enlarged anterior lateral maxillary foramen is widespread in early amniotes, being present in the early varanopid *Archaeovenator* (Reisz and Dilkes 1993) and mesenosaurines (Botha-Brink and Modesto 2009), the captorhinid *Saurorictus* (Modesto and Smith 2001), the early reptile *Orovenator* (Ford and Benson 2018), and saurians (e.g., *Prolacerta*, BP/1/588). However, this foramen is absent in the earliest-diverging 'parareptilies', the mesosaurids (Modesto 2006), and its widespread occurrence across Amniota (and in particular, reptiles) indicates that the phylogenetic utility of this feature (at least at broader taxonomic levels) is less than previously interpreted, although it is still informative for smaller taxonomic groups in which this feature can be highly variable (e.g., Archosauromorpha).

**D) Anterior process of jugal reaching anterior orbital margin.**

An elongate, anterior process of the jugal reaching the anterior orbital margin was found by deBraga and Reisz (1996, Ch. 11) to be an ambiguous autapomorphy of Parareptilia. Under their optimization, a jugal reaching the anterior orbital margin was an autapomorphy of the clade, with reversals present in millerettids and *Acleistorhinus* (deBraga and Reisz 1996). A jugal reaching the anterior orbital margin is a feature of procolophonians and mesosaurids only within 'Parareptilia', with the jugal failing to reach the level of the anterior orbit in millerettids (e.g., *Milleropsis,* BP/1/720), acleistorhinids (e.g., *Delorhynchus*, Reisz et al. 2013), and bolosaurids (e.g., *Belebey*, Reisz et. al 2007). This character is extremely informative for other taxonomic groups (e.g., Captorhinidae) and for assessing broader amniote relationships, but does not diagnose 'Parareptilia' here or in other recent studies.

**E) Quadratojugal expanded dorsally**

Noted as an autapomorphy of 'Parareptilia' in Laurin and Reisz (1995, Ch. 28). We find the quadratojugal of millerettids (e.g., *Milleretta,* BP/1/3822) to be dorsally expanded, although this is not visible in lateral view, a autapomorphy of the clade which is also present in the putative stem-turtle *Eunotosaurus* (Bever et al. 2015). The quadratojugal of early procolophonoids (e.g., *Saurodektes,* Reisz and Scott 2002) is moderately expanded, although it is no taller than the jugal, unlike millerettids in which the quadratojugal is nearly as tall as the entire temporal region. In contrast, the quadratojugal is low in all other 'parareptiles' (and early amniotes more generally), including bolosaurids (e.g., *Belebey*, Reisz et al. 2007), mesosaurids (e.g., *Mesosaurus*, Modesto 2006), acleistorhinids (e.g., *Acleistorhinus*, deBraga and Reisz 1996), and pareiasauromorphs (e.g., *Macroleter*, Tsuji 2007).

**F) Temporal emargination bordered by quadratojugal and squamosal**

A lower temporal emargination was found by Laurin and Reisz (1995, Ch. 31) as an autapomorphy of Parareptilia. A temporal emargination, that is, a lower temporal fenestra that is not bounded by a lower temporal bar, is present in the millerettids '*Millerosaurus' nuffieldi* and *Milleropsis* (BP/1/720), the putative stem-turtle *Eunotosaurus*, and some procolophonoids (e.g., *Procolophon*, Carroll and Lindsay 1985), although this emargination is architecturally very different between millerettids and procolophonids. In procolophonids, this emargination is expressed as a concavity on the ventral surface of the quadratojugal and jugal, which remain in contact. In *Milleropsis*, the quadratojugal and jugal are not in contact, and this emargination is expressed on the ventral surface of the postorbital and squamosal. A lower temporal fenestra, that is, an emargination framed by a lower temporal bar, is present in all other 'parareptiles' (e.g., *Belebey,* Reisz et al. 2009), except mesosaurids (although see

Laurin and Piñeiro 2017, who argue for the presence of an LTF in *Mesosaurus*), and pareiasaurs, which secondarily close this opening (Lee 1993).

**G) Postfrontal-Supratemporal contact.**

Contact between the postfrontal and supratemporal was listed as an autapomorphy of Parareptilia by deBraga and Reisz (1996, Ch. 19), despite the occurrence of this contact being restricted to *Acleistorhinus* (deBraga and Reisz 1996) and the owenettid *Saurodektes* (Reisz and Scott 2002) within their sampling. The presence or absence of this contact does not seem particularly informative due to its absence in other acleistorhinids and procolophonoids, suggesting these features may be autapomorphic for these two taxa.

**H) Jaw articulation located slightly anterior to the occiput.**

A jaw articulation located anterior to the occiput was considered to be an autapomorphy of ‘Parareptilia’ by Laurin and Reisz (1995, Ch. 36). However, this morphology is only present in ankyramorphs and the millerettid *Milleretta* (BP/1/3822) among classical ‘parareptiles’. The jaw articulation is located level with or posterior to the occiput in all other ‘parareptiles’ including *Milleropsis* (BP/1/720), mesosaurids (Modesto 2006), and bolosaurids (Reisz et al. 2007).

**I) Ectopterygoid small.**

A small ectopterygoid was found as a autapomorphy of ‘Parareptilia’ by Laurin and Reisz (1995, Ch. 47), although it is here found to be a autapomorphy of Amniota + Protorothyrididae + Captorhinidae + Diadectomorpha. The ectopterygoid of early tetrapods, and many early reptiliomorphs (e.g., *Seymouria*, Klembara et al. 2005), is roughly equal to (and sometimes longer than) the palatine, in contrast to the reduced ectopterygoid of all unambiguous near-amniotes.

**J) Edentulous ectopterygoid.**

An ectopterygoid lacking dentition was found by Laurin and Reisz (1995, Ch. 38) as a autapomorphy of ‘Parareptilia’, although the acleistorhinid ‘parareptile’ *Delorhynchus* possesses denticles on the ectopterygoid (Reisz et al. 2014). Furthermore, ectopterygoid dentition is lost in neodiapsids (e.g., *Youngina*, Hunt et al. 2024), indicating an edentulous ectopterygoid is widespread in middle and late Permian reptiles.

**K) Ectopterygoid relationship to the transverse flange.**

An ectopterygoid that contributes to the transverse flange laterally is noted by deBraga and Reisz (1996, Ch. 33) as a autapomorphy of ‘Parareptilia’, although they correctly note in the supplemental data that this is a feature unique to ankyramorphs within their sampling. The ectopterygoid is located distal to the transverse flange in other parareptiles (e.g., the millerettid *Milleropsis* or the bolosaurid *Belebey,* Reisz et al. 2007), and is thus not diagnostic of ‘Parareptilia’.

**L) Paroccipital process sutured to squamosal and supratemporal**

A paroccipital process contacting the squamosal and supratemporal laterally was considered an autapomorphy of ‘Parareptilia’ by Laurin and Reisz (1995, Ch. 57). However, the paroccipital process of millerettids contacts the quadrate laterally— a feature millerettids share with neodiapsids among Paleozoic reptiles (Gow 1975). Further, the paroccipital process does not make contact with the lateral skull in mesosaurids. This feature has been noted to be ontogenetically variable (Modesto 2006) and mesosaurids exhibit delayed ossification (Núñez Demarco al. 2022), which may explain the absence of paroccipital processes-suspensorium contact in these taxa.

**M) Reduced posttemporal fenestra.**

The presence of a reduced posttemporal fenestra was supported as a autapomorphy of 'Parareptilia' by deBraga and Reisz (1996, Ch. 46), likely based on the line drawings of Gow (1972) which reconstruct the posttemporal fenestrae of millerettids as reduced, although this is not the case (e.g., *Milleropsis*, BP/1/720 or *Milleretta* BP/1/3822). Furthermore, the posttemporal fenestra is not reduced in procolophonoids (e.g., *Procolophon*, Carroll and Lindsay 1985), suggesting the reduced posttemporal fenestra is not diagnostic of a larger group of 'parareptiles'. A reduced posttemporal fenestra is also widespread elsewhere in Amniota, notably in synapsids.

**N) Stapedial dorsal process unossified.**

The loss of a dorsal process of the stapes was interpreted as an autapomorphy of 'Parareptilia' by Laurin and Reisz (1995, Ch. 67), but the dorsal process was later reported in acleistorhinids (deBraga and Reisz 1996). A dorsal process of the stapes is indeed present in taxa formerly interpreted as 'parareptiles', including *Mesosaurus* (Modesto 2006), acleistorhinids (e.g., *Feeserpeton*, MacDougall et al. 2019), and the early ankyramorph *Australothyris* (Modesto et al. 2009), suggesting that this process is plesiomorphically present in Ankyramorpha. In contrast, the dorsal process of the stapes is lost in procolophonomorphs (e.g., *Nyctiphruretus*), the early reptile *Orovenator* (Ford and Benson 2018), millerettids (Gow 1972), and neodiapsids (Carroll 1981).

**O) Sacral ribs with narrow distal contact**

The sacral ribs of 'parareptiles' were noted by Laurin and Reisz (1995, Ch. 89) as possessing a narrow distal contact, such that the space between the ribs is anteroposteriorly reduced. However, the sacral ribs of millerettids are not in contact (this study, also correctly noted by Spindler et al. 2019) and the space between the sacral ribs in mesosaurids is quite large (Modesto 2010).

**P) Supraglenoid foramen of the scapulocoracoid absent**

The loss of a supraglenoid foramen was interpreted as an autapomorphy of 'Parareptilia' by Laurin and Reisz (1995, Ch. 97) at the time of their writing. However, the supraglenoid foramen has been documented in procolophonoids (e.g., *Eomurruna,* Hamley et al. 2021) and the acleistorhinid *Delorhynchus* (Reisz et al. 2014) since that initial study. The supraglenoid foramen is absent in mesosaurids (Modesto 2010), millerettids (Gow 1972), the synapsid *Heleosaurus* (Botha-Brink and Modesto 2009), and neodiapsids (Gow 1975), indicating that the loss of this feature is not diagnostic of a larger 'Parareptilia', although it is certainly informative to narrower amniote clades.

**Q) Ch. 102. Ectepicondylar groove and foramen present.**

An ectepicondylar groove was considered by Laurin and Reisz (1995, Ch. 102) as a feature of 'Parareptilia'. The presence of an ectepicondylar groove and foramen of the humerus is the plesiomorphic condition in Amniota, and is present in varanopid synapsids (Botha-Brink and Modesto 2009) and non-'parareptilian' reptiles including the araeoscelidian *Araeoscelis* (Vaughn 1955), and some neodiapsids (e.g., *Acerosodontosaurus*, Currie 1981). Therefore, this character does not diagnose a clade of ‘parareptiles’, and an ontogenetic component of this character whereby the supinator process develops until it reaches and fuses with the ectepicondyle, thereby creating an entepicondylar foramen, cannot be ruled out (e.g., *Thadeosaurus*, Currie and Carroll 1984).

**R) Ch. 108. Iliac blade dorsally expanded**

The dorsal expansion of the iliac blade was found as an autapomorphy of ‘Parareptilia’ by Laurin and Reisz (1995, Ch. 108). We note that the iliac blade of ankyramorphan ‘parareptiles’ is not dorsally expanded, being held vertically above the acetabulum (e.g., *Saurodektes kitchingorum*, BP/1/4195), whereas the ilium of mesosaurids (e.g., *Stereosternum*, Modesto 1999) and bolosaurids is posterodorsally expanded, as in other early amniotes (Berman et al. 2021). In millerettids, the ilium is expanded posteriorly (this study), indicating that ilium morphology is extremely plastic in Paleozoic reptiles, and a single character state does not support a monophyletic ‘Parareptilia’, although it is informative for defining smaller clades of Paleozoic reptiles such as paraeiasauromorphs.

**S) Iliac blade fan-shaped lacking a posterior process**

In the analysis of deBraga and Reisz (1996, Ch. 58), ‘Parareptilia’ was diagnosed by the presence of a fan-shaped ilium lacking a posterior process, however, this feature is restricted to ankyramorphans in this study. Former ‘parareptiles’ such as millerettids (this study), mesosaurids (Modesto 2010), and bolosaurids (Berman et al. 2021) possess a discrete posterior process of the ilium.

### 5. Features classically linking *Hylonomus* and Amniota

#### Pre-cladistics

*Hylonomus* was originally described by Dawson (1863) as an early reptile and he erected 'Microsauria' to unite this taxon with similar, small-bodied reptile-like taxa, although later (Dawson 1882, 1896) he referred to *Hylonomus* and other 'microsaurs' as amphibians. This trend was followed by others for much of the next century until Romer (1950) suggested that *Hylonomus* may be a 'reptile' unrelated to 'microsaurs'. Carroll (1964) described more material of *Hylonomus*, and argued in support of its 'reptile' affinities, a sentiment echoed in later papers (Carroll 1969, Carroll 1982) and by other authors.

It is important to note, however, that the term 'reptile' used by these works was broadly equivalent to the our current understanding of the amniote total-group, and referred to many, distantly related taxa such as captorhinomorphs (captorhinids and protorothyridids), 'pelycosaurs', and total-group reptiles. For instance, Carroll (1969, p. 472) states "*We can be quite certain that romeriids and pelycosaurs had a close common ancestor which, in all probability, could be classified as a romeriid. Presumably all Mesozoic and Cenozoic amniotes evolved initially from this single family which had already achieved reptilian status in both morphology and physiology.*" Despite this, for quite some time phylogenetic analyses for both molecular and fossil divergences have used *Hylonomus* to estimate the earliest appearance of Reptilia, taking Carroll's usage of 'Reptilia' at face value.

We describe some of the anatomical features linking *Hylonomus* with 'reptiles' noted by Carroll (e.g., 1964, 1980) and other studies such as Heaton and Reisz (1986) and Müller and Reisz (2006), as well as their broad occurrence among stem-amniotes and small-bodied early synapsids below. We do note here, however, that many specimens referred to *Hylonomus* bear extremely disparate morphology (e.g., the two ilium and parasphenoid morphotypes described by Carroll 1964) and existing specimens are mostly disarticulated and a reassessment of material referred to this genus is therefore necessary.

1) **Postorbital-supratemporal contact absent.** The absence of postorbital-supratemporal contact is reported as a feature linking *Hylonomus* with eureptiles by Müller and Reisz (2006). We note here that a <u>supratemporal is unknown in *Hylonomus*</u>, and the only known postorbital is disarticulated from the rest of the skull in NHMUK 12016, <u>which is a disarticulated specimen lacking a supratemporal, parietal, and postfrontal</u>. This postorbital is a rectangular element, lacking the distinct posterior process present in many near and crown amniotes, including protorothyridids, synapsids, conventional 'parareptiles', and neodiapsids, but similar to the short postorbital present in many basal stem-amniotes such as recumbirostrans and seymouriamorphs. Although this postorbital is disarticulated, if indeed referable to *Hylonomus*, probably, this element did not contact the supratemporal posteriorly. This feature (absence of postorbital-supratemporal contact) has been found as an autapomorphy of Eureptilia by Müller and Reisz (2006), and is indeed absent in protorothyridids (Clark and Carroll 1973), early captorhinids (Heaton 1979), and araeoscelidians (Reisz 1981). The short postorbital in early reptiliomorphs such as *Seymouria* (Laurin 1995) and some diadectomorphs (*Tseajaia*, Moss 1972) does not make contact with the supratemporal either, and this condition is the plesiomorphic condition for near-amniotes. The posterior process of the postorbital develops during ontogeny in the millerettid *Milleretta*, in which this contact is absent in juvenile individuals.

The absence (or fusion) of a supratemporal in many 'microsaurs' (which also bear a similarly reduced postorbital) makes this feature impossible to score.

2) **Otic notch absent**. The absence of an 'otic notch', a term historically used to refer to the spiracular notch of early tetrapodomorphs, but also the tympanic emargination present in a variety of amniotes (although this latter usage has fallen out of favor (e.g., *Macroleter*, reviewed in part by Müller and Tsuji 2007)), is described by Carroll (1964) and subsequent studies as a feature uniting *Hylonomus* with captorhinomorphs. However, an 'otic notch' (regardless of whether it is a spiracular notch or tympanic emargination) is also absent in 'microsaurs' such as recumbirostrans, synapsids, and most early reptiles, and is therefore not a feature exclusive to Carroll's 'Captorhinomorpha'. An 'otic notch' is present in *Seymouria* and some diadectomorphs (e.g., *Orobates*, Berman et al. 2004). The loss of an otic notch is not found as an autapomorphy of Reptilia in any cladistic analysis, and is instead lost earlier in amniote evolution (e. g., Müller and Reisz 2006, Ford and Benson 2019).

3) **Transverse flange of the pterygoid.** The transverse flange of the pterygoid was described as a feature distinguishing *Hylonomus* and other 'reptiles' from Microsauria (Carroll 1964), although Carroll (1969) and other studies subsequently identified the development of this feature as widespread in Reptiliomorpha, including Recumbirostra and Synapsida (Mann et al. 2022). A transverse flange of the pterygoid is best interpreted as a feature ancestral to crownward-stem amniotes, which is in agreement with Carroll's usage of 'Reptilia' encompassing all near-amniotes.

4) **Tympanic flange of the pterygoid.** Carroll (1964) described the presence of a tympanic flange on the quadrate ramus of the pterygoid as a feature uniting *Hylonomus* with early reptiles such as *Petrolacosaurus*. A tympanic flange of the pterygoid is widespread around the origin of amniotes, and is present in diadectomorphs (Romer 1946, Berman et al. 2010), most synapsids (Romer and Price 1940), and nearly all 'reptiles' barring captorhinids (Heaton 1979). The presence of this feature certainly distinguishes *Hylonomus* from 'microsaurs' and suggests a crownward position of *Hylonomus* along the amniote stem (in agreement with Carroll's usage of 'Reptilia') but does not support the placement of this taxon within modern usages of Reptilia.

5) **Posterior cheek in line with the occiput.** Used in support of a *Hylonomus* placement among protorothyridid 'stem-reptiles', a grade then thought to be ancestral to crown Amniota in its modern usage by Carroll's 'The Origin of Reptiles' (Carroll 1991). This feature contrasts *Hylonomus* with many 'microsaurs' described by Carroll (namely groups now considered part of Recumbirostra), but it has been noted that the anterior placement of the cheek relative to the occiput in this clade is a plausible burrowing adaptation (Szostakiwskyj et al. 2015, Pardo et al. 2017). An occiput that is in line with the posterior margin of the cheek is a feature *Hylonomus* shares with protorothyridids and non-moradisaurine captorhinids among traditional 'eureptiles', but also the 'microsaur' *Pantylus* (Romer 1968), *Seymouria* (Laurin 1995), caseasaurs (Reisz et al. 2009), and small-bodied varanopids (e.g., *Mycterosaurus*, Berman and Reisz 1982), and cannot be taken to be diagnostic of total-group Reptilia in today's usage nor Protorothyridae of Carroll (1991).

6) **Narrow iliac blade.** A narrow iliac blade was found as an autapomorphy of *Hylonomus* within Eureptilia by (Müller and Reisz 2006), although this is the plesiomorphic condition within Tetrapoda, in particular in taxa with a small body size such as 'microsaurs' (Carroll and Gaskill 1978) and even early-diverging synapsids (e.g., *Echinerpeton,* Reisz 1971). Furthermore, Müller

and Reisz (2006) offered no quantifiable measure to score this feature, hence scoring it is not reproducible. Notably, material referred to *Hylonomus* document at least two different types of iliac blade morphology, with the ilium of the holotype specimen being much more broad distally (Carroll 1964, fig. 1) compared to referred specimen NMC 10048 (Carroll 1964, figs. 8-9). Additionally, neodiapsid stem-reptiles bear quite broad ilia (e.g., *Hovasaurus*, Currie 1981) that are much broader distally than documented in *Hylonomus* and various 'early eureptiles'. Therefore this character cannot be used to place *Hylonomus* within Reptilia to the exclusion of all other amniotes.

7) **Astragalus**. An astragalus ossification was argued by Carroll (1964) as evidence *Hylonomus* is a 'reptile' or amniote in his usage. However, recent work has demonstrated that an astragalus ossification is widespread among stem-amniotes crownward of Diadectomorpha and cannot be interpreted as a character placing *Hylonomus* within Reptilia (Heaton and Reisz 1986).
8) **Number of pedal centralia.** Carroll (1964) identifies an additional feature that *Hylonomus* was thought to share exclusively with total-group reptiles: the presence of a single pedal centrale, one fewer than present in most synapsids. However, the pes preserved in NMC 10048, a referred specimen to *Hylonomus* is only partially articulated, and we identify a disarticulated medial pedal centrale in this specimen. This observation is corroborated by Carroll (1991, fig. 9) who reconstructed the tarsus of *Hylonomus* based on this specimen with two pedal centralia. Furthermore, since the Carroll's work, several synapsids have been revealed to possess a single pedal centrale including some caseasaurs (e.g., *Martensius*, Berman et al. 2020) and varanopids (e.g., *Mesenosaurus*, Spindler et al. 2019). Additionally, the 'peanut-shaped' pedal centrale of captorhinids has been previously proposed to be a fusion of the medial and lateral centralia of stem amniotes (e.g., O'Keefe et al. 2005).

| **Feature** | *Hylonomus* | *Seymouria* | Microsauria | Captorhinidae | Diadectomorpha | Primitive Caseasauria | Primitive Varanopidae |
|---|---|---|---|---|---|---|---|

| | | | | | | | |
|---|---|---|---|---|---|---|---|
| No Postorbital-Supratemporal contact | ✓ | ✓ | ? | ✓ | ✓X | X | X |
| No otic notch | ✓ | X | ✓ | ✓ | X | ✓ | ✓ |
| Transverse flange of pterygoid | ✓ | ✓ | ✓ | ✓ | ✓ | ✓ | ✓ |
| Tympanic flange of pterygoid | ✓ | X | X | X | ✓ | ✓ | ✓ |
| Posterior cheek in line with occiput | ✓ | ✓ | X | ✓ | ✓X | ✓ | ✓ |
| Narrow Iliac blade | ✓ | ✓ | ✓ | ✓ | X | ✓ | ✓ |
| Lightly built limbs | ✓ | X | ✓ | ✓ | X | ✓ | ✓ |
| Astragalus | ✓ | X | X ✓ | ✓ | ✓ | ✓ | ✓ |
| One pedal centrale | X | X | X ✓ | X ✓ | X | ✓ | ✓ |

**Table S2.** Features previously suggested as supporting a placement of *Hylonomus* within Amniota and their widespread occurrence along the amniote stem and Synapsida.

### 6. Additional Steps required to support monophyly of Parareptilia, Eureptilia, Diapsida, and the inclusion of Recumbirostra within reptiles

Additional parsimony analyses were run in PAUP* 4.0a169 which included topological constraints to test alternative hypotheses for amniote relationships, including 'traditional' classification schemes. In all cases, the clade of focus was constrained to find the simplest (i.e. least amount of steps) tree to make an alternative class scheme possible. The phylogenetic hypotheses tested by this method were:

**(I)** A monophyletic Eureptilia including Captorhinidae, Araeoscelidia, and Neodiapsida.
**(II)** A monophyletic Eureptilia excluding the Millerettidae
**(III)** A monophyletic Parareptilia including Mesosauridae, Millerettidae, Bolosauridae, Acleistorhinidae, and Procolophonia.
**(IV)** A monophyletic Diapsida placing Araeoscelidia as sister to Neodiapsida
**(V)** A monophyletic Diapsida excluding Millerettidae
**(VI)** Recumbirostra within Reptilia

| Taxon | Tree Length | Additional Steps | Most Parsimonious Trees |
|---|---|---|---|
| **Monophyletic Eureptilia** | 4790 | 39 | 102 |
| **Monophyletic Eureptilia excluding Millerettidae** | 4856 | 105 | 184 |
| **Monophyletic Parareptilia** | 4832 | 81 | 97 |
| **Monophyletic Diapsida** | 4784 | 33 | 96 |
| **Monophyletic Diapsida excluding Millerettidae** | 4815 | 64 | 133 |
| **Recumbirostra within Reptilia** | 4781 | 30 | 112 |

**Table S3.** Constrained amniote relationships resulting in the monophyly of clades not supported by this study, listing tree length, additional steps, and the number of most parsimonious trees found by parsimony analysis when these constraints are used.

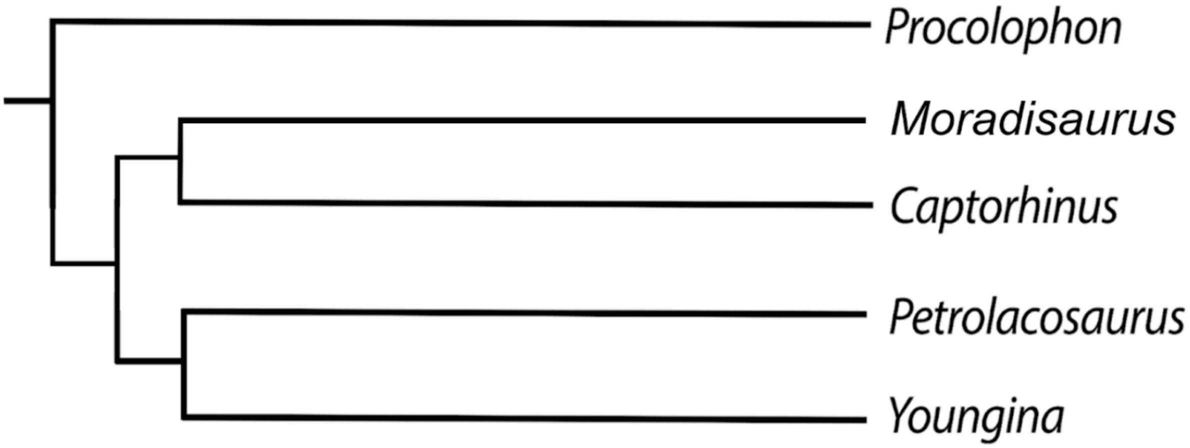

**Fig S6. Backbone topology used for the constraint analysis testing a monophyletic Eureptilia following the definition of the clade provided by Tsuji and Müller (2009).**

### Testing a monophyletic Eureptilia

The enforcement of a monophyletic 'Eureptilia', that is, a clade including, captorhinids, araeoscelidians, and neodiapsid reptiles to the exclusion of traditional 'parareptiles' following the definition of the clade by Tsuji and Müller (2009). This was examined using a constraint analysis in PAUP* 4.0a169. The backbone topology here enforced a clade that consisting of Captorhinidae, Araeoscelidia, and Neodiapsida respectively to the exclusion of *Procolophon*, the 'parareptile' specifier for the definition of Eureptilia used by Tsuji and Müller 2009. The resulting tree was 4790 steps, or 39 additional steps compared to our base analysis. When a monophyletic 'Eureptilia' is enforced, the monophyly of 'Parareptilia' remains unsupported. Millerettidae is still placed as sister to all Neodiapsida, therefore Parapleurota remains strongly supported regardless of the position of other 'parareptiles'.

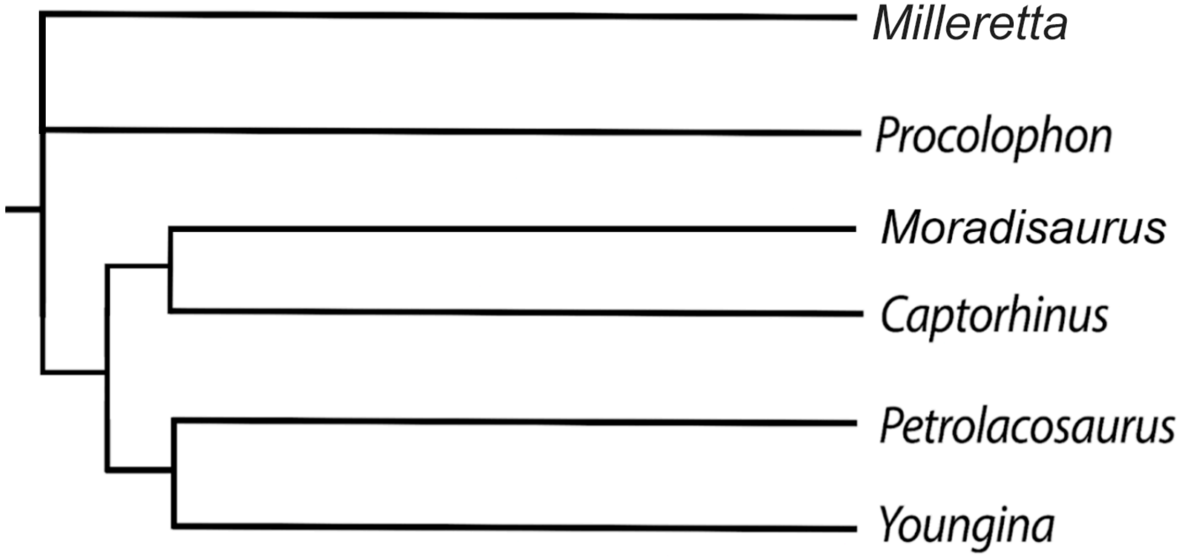


**Fig S7. Backbone topology used for the constraint analysis testing a monophyletic Eureptilia wherein all former 'parareptiles' are excluded from the group.**

### Testing a monophyletic Eureptilia excluding Millerettidae

We also performed a constraint analysis that mirrors most conventional phylogenetic analyses (e.g., MacDougall et al. 2019) wherein there is a Eureptilia-Parareptilia dichotomy, where we specifically exclude Millerettidae from Eureptilia. This constraint analysis is similar to the above, with the exception that *Milleretta rubidgei* is forced outside of a clade consisting of Captorhinidae, Araeoscelidia, and Neodiapsida. When a more conventional Eureptilia is enforced, the simplest tree is 4856 steps, or 105 additional steps compared to our base analysis. When this constraint is enforced, Millerettidae are placed as sister to Eureptilia and do not form a clade with other 'parareptiles'.

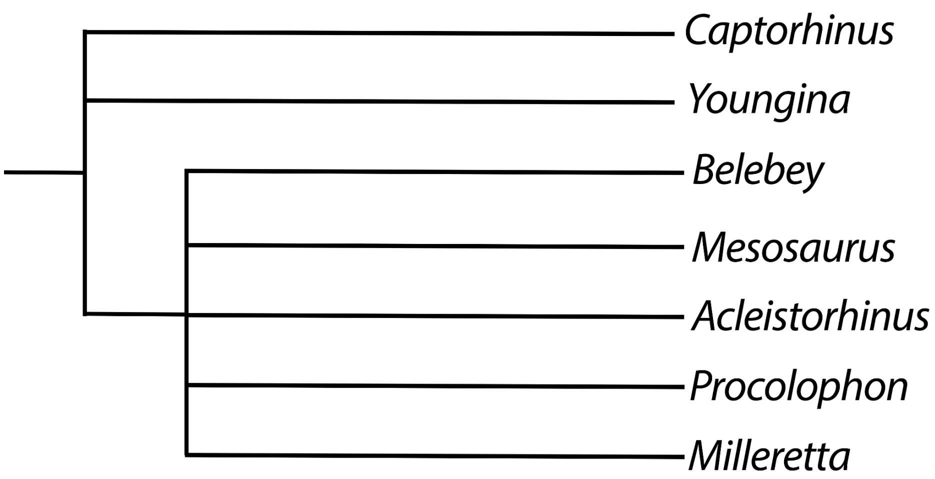


**Fig S6. Backbone topology used for the constraint analysis testing a monophyletic Parareptilia.**

Testing a monophyletic Parareptilia

We also performed a constraint analysis forcing the monophyly of 'Parareptilia', that is, a group of reptiles including the mesosaurs, millerettids, bolosaurids, acleistorhinids, and procolophonians. Our backbone topology enforced a relationship between all Millerettidae, Mesosauridae, Bolosauridae, and Acleistorhinidae, and Procolophonia that excludes both *Captorhinus* and *Youngina*, again following the definitions of 'Eureptilia' and 'Parareptilia' proposed by Tsuji and Müller (2009). The simplest tree was 4832 steps, or 81 additional steps when compared to our base analysis. When this topology was constrained, amniote relationships are broadly similar to our base analysis, that is, recumbirostrans, captorhinids, protorothyridids, and diadectomorphs are placed along the amniote stem. Under this constraint, however, a monophyletic 'Parareptilia' is still crownward of araeoscelidians, yielding a similar topology for Neoreptilia to that of Ford and Benson (2019).

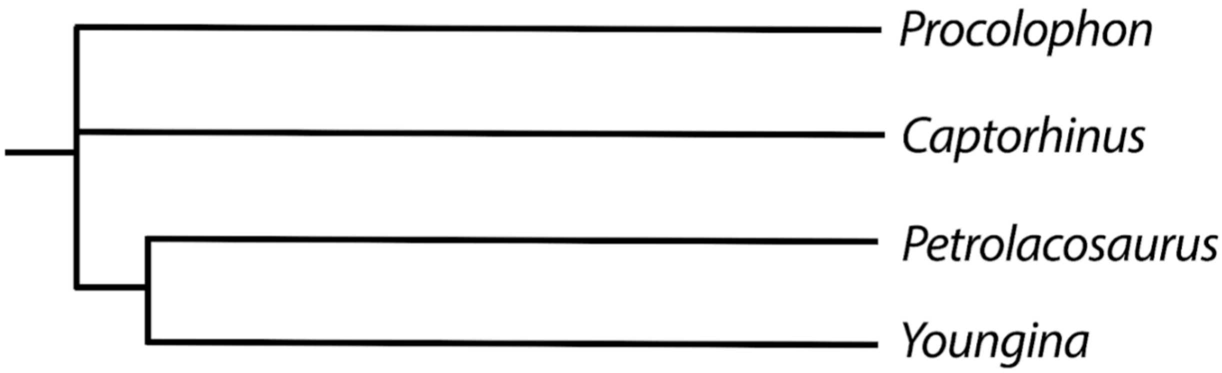


**Fig S8. Backbone topology used for the constraint analysis testing a monophyletic Diapsida**

Testing a monophyletic Diapsida

The enforcement of a monophyletic 'Diapsida', that is, a clade consisting of Araeoscelidea and Neodiapsida, was investigated using constraint analysis. As the phylogenetic definition of Diapsida provided by the PhyloCode does not explicitly restrict the placement of this group, we performed a constraint analysis that forced the relationship araeoscelidians (namely *Petrolacosaurus*) and neodiapsids (namely *Youngina*) to the exclusion of *Captorhinus* and *Procolophon*, the two early reptile species generally used in other phylogenetic definitions (e.g., Tsuji and Müller (2009) or Ford and Benson (2019)). Under this constraint, Millerettidae is placed within Diapsida as sister to Neodiapsida with araeoscelidians as the earliest diverging diapsids. Therefore, the name Parapleurota remains applicable even under alternative hypotheses of reptile relationships.

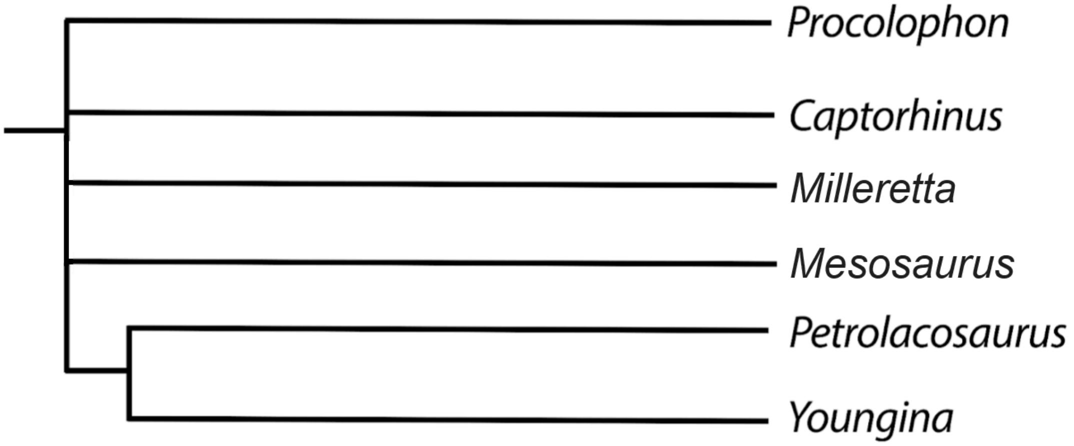


**Fig S9. Backbone topology used for the constraint analysis testing a monophyletic Diapsida excluding Millerettidae.**

Testing a monophyletic Diapsida excluding Millerettidae

Constraining a monophyletic 'Diapsida' mirroring conventional phylogenetic analyses (MacDougall et al. 2019) required the forcing of araeoscelidians to be placed in a clade sister to Neodiapsida to the exclusion of all eureptiles and parareptiles. This backbone topology enforced only a relationship between *Youngina* and *Petrolacosaurus* to the exclusion of other 'eureptiles' (in this instance, *Captorhinus*) and 'parareptiles' (represented by *Procolophon, Mesosaurus, and Milleretta*). The simplest tree found here was 4815 steps, or 64 additional steps relative to our base analysis. There is no support for a monophyletic 'Parareptilia' when Diapsida is constrained.

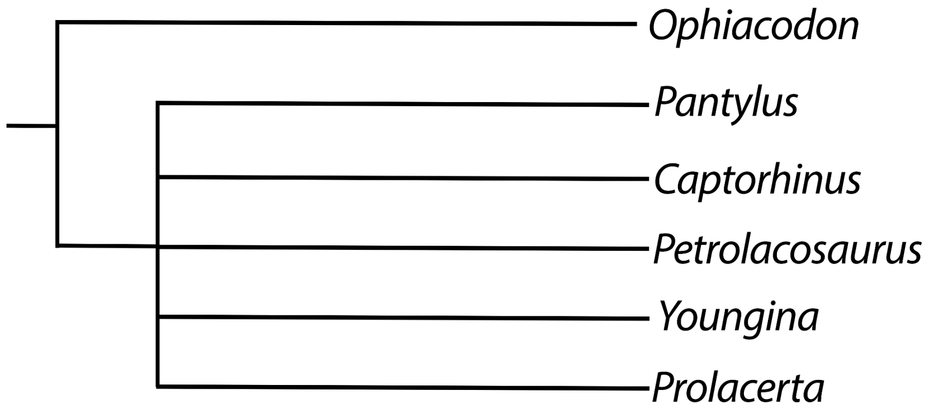


**Fig S10. Backbone topology used for testing a placement of Recumbirostra within Reptilia.**

Testing the Placement of Recumbirostra within Reptilia

Constraining Recumbirostra to be placed within Reptilia was quite complicated. Forcing a topological constraint with the earliest-diverging reptiles in this matrix, the araeoscelidians, resulted in trees in which Recumbirostra + Araeoscelidia were placed as stem-amniotes. To restrict Recumbirostra into

Reptilia, we had to apply additional constraints using captorhinids (*Captorhinus*), araeoscelidians (*Petrolacosaurus*), neodiapsids (*Youngina*), and the reptile crown (*Prolacerta*) as reference taxa to the exclusion of the synapsid *Ophiacodon*. When recumbirostrans are forced into Reptilia, the shortest resulting tree is 4781 steps, or 30 additional steps relative to our base analysis. Enforcing these constraints also resulted in the placement of captorhinids and protorothyridids within Reptilia and diadectomorphs within Synapsida, perhaps corroborating views by Pardo (2017) or Mann et al. (2022) that captorhinids (and other stem amniotes more broadly) share widespread similarities with recumbirostrans. Otherwise, the relationships of early reptiles are identical to our original topology.

### 7. Measures of Stratigraphic fit

#### Data Files

The following data file is a script designed to work in conjunction in R v. 3.3.0, following the framework provided by Bell and Lloyd (2015) for MSM* and GER* and Pol and Norell (2003) for MSM*Diff. Explanatory notes are provided:

```
#load in required packages for stratigraphic fit analyses
library(geoscale)
library(strap)
library(ape)
#read age file of FADs and LADs associated with our Bayesian script in format required by Bell and Lloyd (2015)
Milleropsis.ages <- read.table("Milleropsis_dates.txt", header=T)
#read tree files of 12 MPTs of parsimony analysis
Milleropsis.tree <- read.tree("MilleropsisMPT.tre")
#Time scale the MPTs using attached tree and age files
Milleropsis.ts.tree <- DatePhylo(Milleropsis.tree[[1]], Milleropsis.ages)
#Assessing stratigraphic congruence, using measures including MSM and GER
 X <- StratPhyloCongruence(Milleropsis.tree, Milleropsis.ages, hard=FALSE,randomly.sample.ages=TRUE,
fix.topology=TRUE, fix.outgroup=TRUE, outgroup.taxon = "Whatcheeria_deltae")
```

```
#View results within R
X$input.tree.results
#export results of 12 MPTs to a CSV titled Milleropsis_strat_congruence
write.table(X$input.tree.results, "Milleropsis_strat_congruence.csv", sep=",")
#Export results of resampling
write.table(X$samp.permutation.results, "Milleropsis_strat_resampled.csv", sep=",")
#view resampled results
X$samp.permutation.results
# view input trees most congruent with stratigraphy using GER
best.input.GER <- X$input.trees[[which.max(X$input.tree.results[, "GER"])]]
# view input trees most congruent with stratigraphy using MSM*
best.input.MSM <- X$input.trees[[which.max(X$input.tree.results[, "MSM*"])]]
#HISTOGRAMS OF GER MEASURES FOR INPUT VS RESAMPLED USING A SIGNIFICANCE CUTOFF
pdf("Milleropsis_strat_fit_GER.pdf", width=10, height=7)
par(mfrow=c(2, 1))
cutoff <- qnorm(0.95, mean(X$samp.permutation.results[, "GER"]),sd(X$samp.permutation.results[, "GER"]))
hist(X$input.tree.results[, "GER"], xlim=c(.4, 1), xlab="GER",
 main="Input trees", col="black")
lines(x=c(cutoff, cutoff), y=c(0, 1000), lty=2)
 hist(X$rand.permutations[, "GER"], xlim=c(.4, 1), xlab="GER",
 main="Randomly generated trees", col="black")
 lines(x=c(cutoff, cutoff), y=c(0, 1000), lty=2)
 dev.off()
 #HISTOGRAMS OF MSM* MEASURES FOR INPUT VS RESAMPLED USING A SIGNIFICANCE CUTOFF
pdf("Milleropsis_strat_fit_MSM.pdf", width=10, height=7)
par(mfrow=c(2, 1))
cutoff <- qnorm(0.95, mean(X$samp.permutation.results[, "MSM*"]),sd(X$samp.permutation.results[, "MSM*"]))
print(cutoff)
hist(X$input.tree.results[, "GER"], xlim=c(0, 1), xlab="MSM*",
 main="Input trees", col="black")
lines(x=c(cutoff, cutoff), y=c(0, 1000), lty=2)
hist(X$rand.permutations[, "MSM*"], xlim=c(0, 1), xlab="MSM*",
 main="Randomly generated trees", col="black")
lines(x=c(cutoff, cutoff), y=c(0, 1000), lty=2)
dev.off()
```

| **Tree** | **GER** | **MSM*** | ***est.p.* GER** | ***est.p.* MSM*** |
|---|---|---|---|---|
| **This study** | 0.8674823 | 0.15994 | >.00001 | >.00001 |
| **Constraint** | 0.68649 | 0.12483 | >.00001 | >.00001 |
| **Cisneros et al. 2020** | 0.59187 | 0.13623 | 0.00031 | >.00001 |
| **Simões et al. 2022** | 0.82394 | 0.14992 | >.00001 | >.00001 |

**Table S4.** Summary of stratigraphic fit analyses for GER and MSM*. The constraint analysis is a modified version of our dataset in which the monophyly of ‘Parareptilia’ and ‘Eureptilia’ are enforced. For comparability, we provide stratigraphic fit measurements for two recent studies of early reptile relationships, but MSM*Diff values were not calculated here due to differing taxonomic sampling.

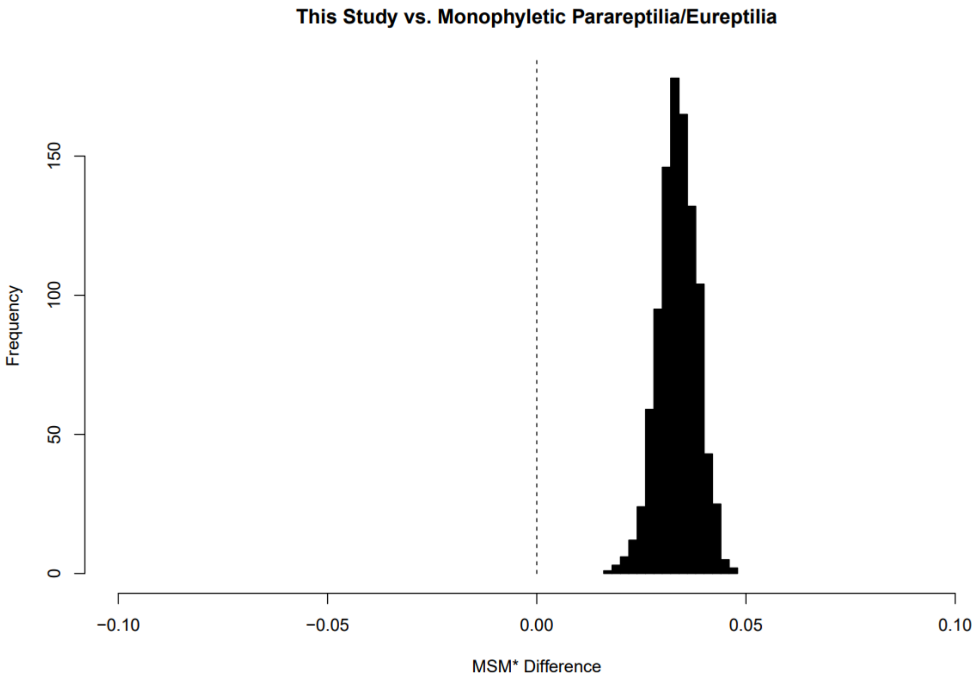


**Fig. S11.** Difference in Manhattan Stratigraphic Measure* replicate values (MSM*Diff) between pairwise topologies under our phylogenetic hypothesis and the constraint yielding a monophyletic Parareptilia, Eureptilia, and a placement of all classical varanopids within Synapsida. Dashed line indicates an MSM*Diff of 0.

**Discussion**

One of the more frequently discussed criticisms of a eureptilian placement of Neodiapsida are several ghost lineages present on the reptile stem according to prevailing phylogenetic hypotheses (Ezcurra et al. 2014). These include the 'eureptile' ghost lineage extending from the appearance of the supposed stem-reptile *Hylonomus* (~318 Ma) to the next appearing 'eureptile' *Petrolacosaurus* (~307-302 Ma), a 'diapsid' ghost lineage from *Petrolacosaurus* to the possible 'diapsid' *Orovenator* (~289 Ma), and two 'neodiapsid' ghost lineages extending from *Orovenator* to *Lanthanolania* and from *Lanthanolania* to the 'younginiform' specimens from the *Endothiodon* Assemblage Zone of the South African Karoo Basin (Late Permian ~252 Ma). Although two taxa (*Orovenator*, 289 mya; and *Lanthanolania,* 268 Ma) were both interpreted to partially fill these ghost lineages as early neodiapsids, the anatomical support for them as 'neodiapsids' has been low, considering the lack of preserves temporal region in both taxa (Modesto and Reisz 2004, Ford and Benson 2018). These ghost lineages for Eureptilia and Neodiapsida in particular have puzzled paleontologists for a quarter of a century, in the absence of reptile taxa that appear anatomically 'transitional' between the remarkably primitive araeoscelidians and derived neodiapsids (Benton 1985). Furthermore, the enigmatic 'eureptile' *Hylonomus* (318 Ma), well predates the existence of the sister taxon to Eureptilia – the Parareptilia – by at least 12 million years, given the oldest putative members of this clade are Stephanian or younger (Fig. 1). Parareptilia similarly possess ghost lineages extending from the Carboniferous to middle Permian, as the Millerettidae, hypothesized to be among the earliest-diverging parareptiles, first occur in the latest middle Permian (MacDougall and Brocklehurst 2019). Fragmentary taxa hypothesized to fill these 'reptile gaps'- such as the putative early reptile *Westlothiana* and the possibly anatomically diapsid *Orovenator* (Reisz et al. 2011) have been demonstrated to lack derived reptile affinities, instead representing an earlier radiation of reptiles (Ford and Benson 2019) or even a non-amniote ‘lepospondyl’ (in the case of the former) (Ruta and Coates 2003, Ruta & Coates 2007, Marjanović & Laurin 2019).

Our phylogenetic hypothesis closes or shortens these various ghost lineages for stem-reptiles. The ghost lineage for ‘Eureptilia’ is non-existent, with the monophyly of this clade being unsupported (Fig. 3). By placing Millerettidae and other taxa (e.g., *Orovenator* and various neoreptiles) as successive outgroups to Neodiapsida, the ghost-lineage for ‘Parareptilia’ is no longer supported here, as the monophyly of this clade is also dubious. The ghost lineage for the neodiapsid stem (neodiapsid Ghost Lineage A) remains 18 million years, extending from the Artinskian (early Permian) through the Roadian (earliest middle Permian), although we do not find support for either *Orovenator* or *Lanthanolania* as neodiapsids (Ezcurra et al. 2014). Our placement of *Hylonomus*– a taxon by some estimates more than ten million years older than the next unambiguous amniotes (in this case, the synapsids *Melanedaphodon*, *Echinerpeton* and *Archaeothyris*)– outside of crown Amniota reduces ghost lineages for Amniota and Reptilia by at least 10 to 12 million years, respectively. Our results – demonstrating the polyphyly of both ‘Eureptilia’ and ‘Parareptilia’ creating a grade of Permian reptiles – necessitate the absence of ghost lineages created by the existence of a dichotomy between these early reptile clades (Ezcurra et al. 2014). The measures of congruence provided above demonstrate that our phylogenetic topology – perhaps unsurprisingly given the scaffolding detailed above – is more stratigraphically congruent than competing hypotheses of reptile evolution.

### 8. Character Additions

The character list used in this phylogenetic analysis has been compiled from a comprehensive list of characters from published sources as well as novel observations here. An emphasis was placed on characters or observations regarding ‘parareptile’, neodiapsid, synapsid, captorhinid, recumbirostran, and early tetrapod morphology: Olson 1947, Benton 1985, Heaton and Reisz 1986, Gauthier et al. 1988, Estes and Gauthier 1988, Laurin 1991, Reisz et al. 1992, Lee 1993, Laurin 1993, Juul 1994, Dodick and Modesto 1995, Lee 1995, Laurin and Reisz 1995, Modesto 1995, Berman et al. 1995, Gower and Sennikov 1996, deBraga and Rieppel 1997, Lee 1997, Sidor and Hopson 1998, Reisz et al. 1998, Dilkes 1998, Modesto 1999, Reisz and Dilkes 2003, Modesto and Sues 2004, Müller et al. 2006, Müller and Reisz 2006, Tsuji 2006, Maddin et al. 2006, Müller and Tsuji 2007, Reisz et al. 2007, Ruta and Coates 2007, Anderson et al. 2008, Maddin et al. 2008, Cisneros 2008, Reisz et al. 2009, Botha-Brink and Modesto 2009, Callier et al. 2009, Sumida et al. 2010, Reisz et al. 2010, Reisz et al. 2011, Nesbitt 2011, Fröbisch et al. 2011, Benson 2012, Clack et al. 2012, Huttenlocker et al. 2013, Modesto et al. 2014, Motani et al. 2014, MacDougall and Reisz 2014, Reisz and Fröbisch 2014, Ezcurra et al. 2014, Klembara et al. 2014, Pritchard et al. 2015, Modesto et al. 2015, Ezcurra 2016, Brocklehurst et al. 2016, Laurin and Piñeiro 2017, Pardo et al. 2017, Pritchard and Nesbitt 2017, Simões et al. 2018, Spindler et al. 2018, Modesto et al. 2018, MacDougall et al. 2017, Pritchard et al. 2018, Ford and Benson 2019, Klembara et al. 2019, Pritchard and Sues 2019, Spindler et al. 2019,

Marjanović and Laurin 2019, Cisneros et al. 2020, Van den Brandt 2020, Ford et al. 2021, Griffiths et al. 2021, Pritchard et al. 2021, Brocklehurst and Benson 2022, Simões et al. 2022, and Mann et al. 2022.

We provide figures for all novel characters added to this dataset and select characters from previous datasets unless characters have been figured in the above pre-existing literature. When figures are provided below, specimen numbers of specimens imaged are placed below their respective images (scan or otherwise).

**Ch. 1. Skull, antorbital-postorbital length ratio: postorbital part longer than antorbital part (0), antorbital part of skull (anterior margin of orbit to anteriormost extremity of skull) equal to or longer than the postorbital part (posterior margin of orbit to posteriormost extremity of skull) (1).**

Modified from Brocklehurst et al. (2022, Ch. 1).

**Ch. 2. Skull, position of nasal in antorbital region: nasal largely restricted to dorsal surface of skull (0), nasal is restricted to the lateral surface of snout and is vertically oriented (1)**

Modified from Brocklehurst et al. (2022, Ch. 2).

**Ch. 3. Skull, extensive dermal sculpturing of skull roof: present (0), absent or weakly developed, lacking depth or being restricted to specific bones (1).**

Modified from Brocklehurst et al. (2022, Ch. 3).

**Ch. 4. composed of ridges and pits arranged in a honeycomb pattern (0), an irregular distribution of rounded pits (1), anastomosing narrow grooves and low ridges (2), deep pits with prominently raised knobs (3) raised bumps as a result of osteodermal incrustation (4).**

Reworked from Brocklehurst et al. (2022, Ch. 4).

Rationale: Added State 4, which concerns the derived state of millerettid stem reptiles which bear cranial osteoderms that are fused with underlying bones.

**Ch. 5. Skull, posterior margin in dorsal view: roughly straight (0), with a single, median embayment (1), embayed bilaterally (2).**

Modified from Brocklehurst et al. (2022, Ch. 5).

**Ch. 6. Skull, dorsal surface of the temporal region: flat or weakly developed, lateral surface of parietal is dorsoventrally thin (0), "supratemporal" fossa present, such that parietals/squamosals are deeply embayed and concave (1).**

Modified from Laurin (1991, Ch. G2), Muller (2004, Ch. 13), Reisz et al. (2010, Ch. 30), Nesbitt (2011, Ch. 59), Ezcurra et al. (2014, Ch. 29), Pritchard et al. (2015, Ch.20), Ezcurra (2016, Ch. 8).

The supratemporal fossa, or a ventral deflection of the skull roof for the adductor musculature, sometimes surrounding the endocranial cavity, is present in crocopod archosauromorphs (e.g., *Euparkeria,* SAM-PK-5867), non-rhynchocephalian lepidosauromorphs (e.g., *Marmoretta*, NMS G1992.47), and therapsids in this sampling (Sidor and Hopson 1998). In contrast, the skull roof elements of most early amniotes are flat and thin with no supratemporal fossa present (e.g., *Saurodektes*, BP/1/4195).

Unambiguous: Weigeltisauridae+ Tangasauridae + *Claudiosaurus* + *Acerosodontosaurus* + Sauria (0 to 1). DELTRAN: Therapsida (0 to 1), Squamata (1 to 0). ACCTRAN: Dinocephalia + *Biseridens* (0 to 1), Rhychocephalia (1 to 0).

**Ch. 7. Skull, skull roof, lateral line sulci: present (0), absent (1).**

Modified from Anderson et al. (2008, Ch. 64), Huttenlocker et al. (2013, Ch. 64).

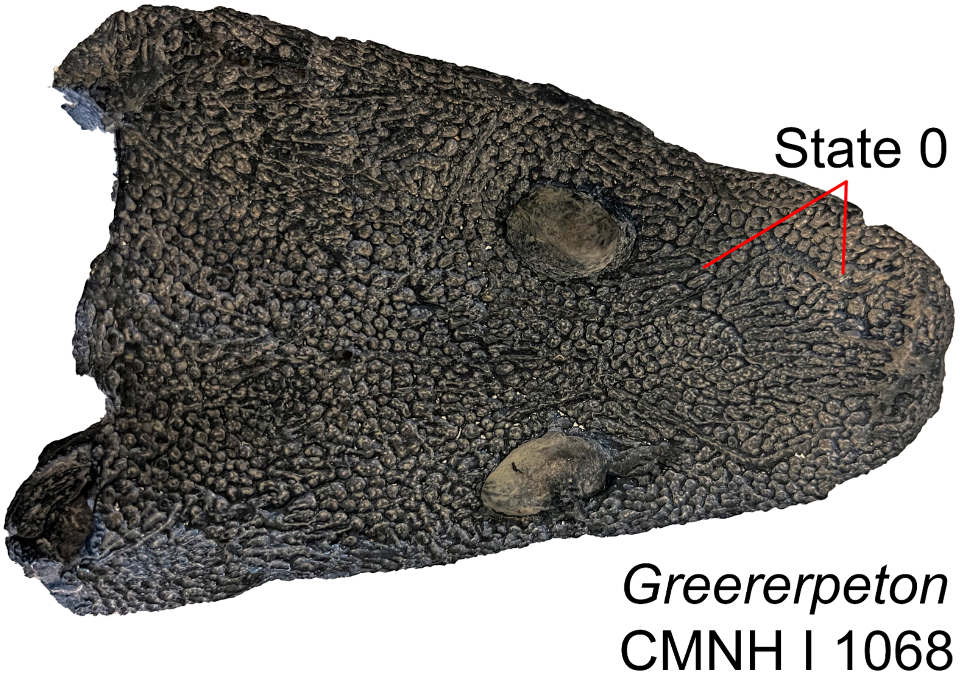


**Fig. S12.** Skull of *Greererpeton* specimen CMNH L I 1068 in dorsal view demonstrating the presence of lateral line sulci (Ch. 7, State 0).

Lateral line sulci are present in early tetrapods such as *Whatcheeria deltae* (FM PR 1634), embolomeres' (e.g., *Archeria crassidiscus*, AMNH FARB 4550), and some temnospondyls (Schoch 1999) and marks the egress of the lateral line system in these taxa. Lateral line sulci are absent (State 1) in all other taxa within this sampling.

UNAMBIGUOUS: Temnospondyli + Reptiliomorpha + *Gephyrostegus* (0 to 1).

**Ch. 8. Premaxillary and anterior dentary teeth, distal recurvature of marginal teeth: present (0), absent (1)**

Modified from Brocklehurst et al. (2022, Ch. 6).

**Ch. 9. Teeth, distal recurvature of marginal teeth, if present: slight to moderate (0), strongly recurved, apex approximately 70-90 degrees from vertical (1).**

Modified from Brocklehurst et al. (2022, Ch. 7).

**Ch. 10. Teeth, marginal dentition, cutting edges ( carinae) on the mesial and distal surfaces: absent or only present distally (0), present throughout the length of the crown (1).**

Modified from Brocklehurst et al. (2022, Ch. 8).

**Ch. 11. Premaxillary and anterior dentary teeth, serrations on crown of marginal teeth: absent (0), present (1).**

Modified from Brocklehurst et al. (2022, Ch. 9).

**Ch. 12. Premaxillary and anterior dentary teeth, lateral compression of marginal dentition only apically or nowhere (0), over two-thirds of tooth (1).**

Modified from Brocklehurst et al. (2022, Ch. 10).

**Ch. 13. Premaxillary and anterior dentary dentition, shoulder or heel on lingual surface: absent (0), present (1).**

Modified from Brocklehurst et al. (2022, Ch. 298).

**Ch. 14. Marginal dentition, number of mesiodistal cusps: one (0), two (1), three (2), more than three (3). ORDERED.**

Modified from Brocklehurst et al. (2022, Ch. 11).

**Ch. 15. Maxillary and posterior dentary teeth, distal recurvature of marginal teeth: absent (0), present (1).**

Modified from Brocklehurst et al. (2022, Ch. 363).

**Ch. 16. Maxillary and posterior dentary teeth, lateral compression of marginal dentition: only apically or nowhere (0), over two-thirds of tooth (1).**

**Ch. 17. Maxillary and posterior dentary teeth, serrations on crown of marginal teeth: absent (0), present (1).**

Modified from Brocklehurst et al. (2022, Ch. 365).

**Ch. 18. Maxillary and posterior dentary dentition lingual heel : absent (0), present (1).**

Modified from Brocklehurst et al. (2022, Ch. 298).

**Ch. 19. Maxilla and dentary teeth, infolding or plicidentine at the base of teeth: absent (0), present (1).**

Modified from Motani et al. (2014, Ch. 53).

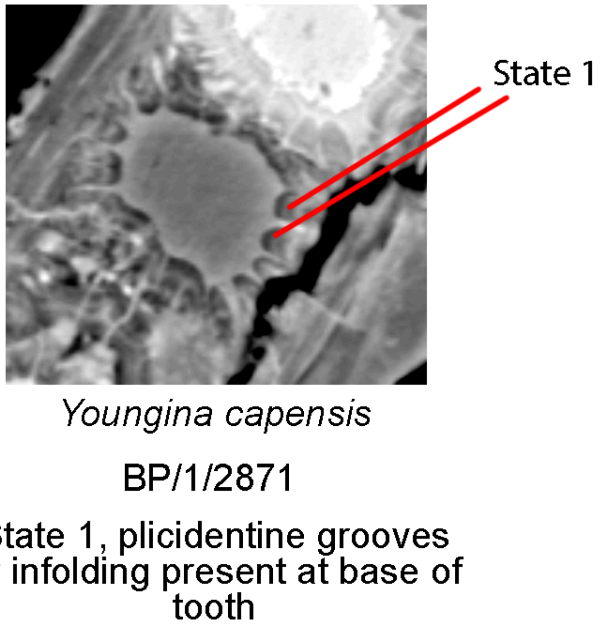


**Fig. S13.** Slice data through the dentary of *Youngina* specimen BP/1/2871 demonstrating the presence of plicidentine infolding at the base of each tooth, previously unreported in a neodiapsid (Ch. 19, State 1).

The presence of plicidentine or 'labyrinthine infolding' (State 1), a pattern of dentine infolding around the pulp cavity of a tooth, is present in many tetrapods including caseasaurs (e.g., *Oedaleops*, UCMP V 35758) 'parareptiles' such as millerettids (e.g., *Milleretta rubidgei* BP/1/3822, Gow 1972) and the acleistorhinid *Colobomycter pholeter* (MacDougall et al. 2014). We score this character only in taxa in which histological or scan data is available, as many taxa possess plicidentine that is not visible in lateral view (e.g., *Youngina capensis*, Hunt et al. 2023).

UNAMBIGUOUS: Moradisaurinae + *Labidosaurus* (1 to 0). DELTRAN: *Claudiosaurus* + Sauria (1 to 0), *Hapsidopareion* (1 to 0), Araeoscelidia (1 to 0). ACCTRAN: *Llistrofus* + Chthonosauria (1 to 0), Bolosauridae (1 to 0), Weigeltisauridae+ Tangasauridae + *Claudiosaurus* + *Acerosodontosaurus* + Sauria (1 to 0)

**Ch. 20. Maxilla and dentary teeth, if infolding or plicidentine present, extent: restricted to the base or root of teeth (0), plicidentine continues through more than a third of the tooth crown, sometimes approaching the tip (1). Inapplicable (-) in taxa lacking plicidentine infolding (Ch. 19).**

Modified from Motani et al. (2014, Ch. 53) and from MacDougall et al. (2014), who detail this anatomy in acleistorhinids.

Plicidentine is restricted to the base or root of teeth in most amniotes sampled here (e.g., *Delorhynchus cifellii,* OMNH 72363 or *Dimetrodon limbatus*, Brink et al. 2014). However, plicidentine continues towards the tip of the tooth in temnospondyls (e.g., *Acheloma*, OMNH 79318, Gee et al. 2019b) and several early or stem amniotes (e.g., *Carbonodraco lundi*, CM 23055 or *Erpetonyx arsenaultorum* ROM 55402).

Unambiguous: *Erpetonyx* (0 to 1). DELTRAN: *Hylonomus lyelli* (0 to 1), Diadectomorpha + Protorothyrididae + Amniota + Recumbirostra (1 to 0). ACCTRAN: Amniota + Protorothyrididae + Diadectomorpha + Recumbirostra + Captorhinidae + *Lanthanosuchus* + *Chroniosaurus* (1 to 0).

**Ch. 21. Posterior marginal dentition, labiolingually expanded accessory cusps: absent (0), present (1).**

Modified from Cisneros 2008 (Ch. 27).

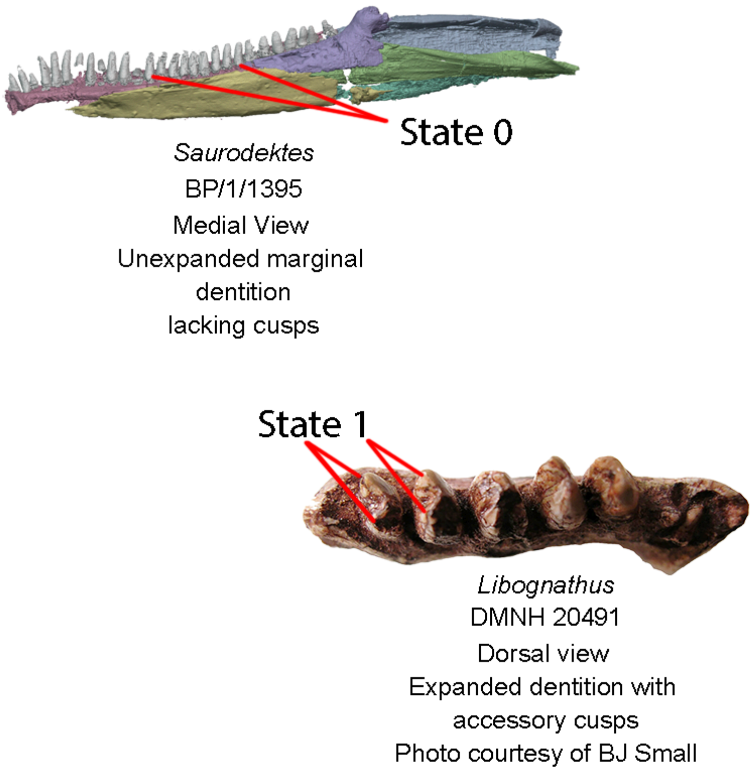


**Fig. S14.** Top left, segmentation of mandible of *Saurodektes kitchingorum* (BP/1/3895, this study) demonstrating (Ch. 21, State 0), or the lack of accessory cusps on its conical dentition. Bottom right, dentary of *Libognathus sheddi* (DMNH 20491) in dorsal view, demonstrating (Ch. 21, State 1), the presence of labiolingual accessory cusps on its marginal dentition.

State 1 is present in procolophonids (e.g., *Sauropareion*, SAM-PK-11192), but absent in early-diverging procolophonoids including owenettids (e.g., *Saurodektes,* Reisz and Scott 2002). A similar expansion of the marginal dentition is present in the archosauromorph *Trilophosaurus* (Heckert et al. 2006) in this taxon sampling.

UNAMBIGUOUS: Leptopleuroninae + *Procolophon* + *Eomurruna* (0 to 1), *Trilophosaurus buettneri* (0 to 1).

**Ch. 22. Posterior marginal dentition, dorsoventral height of roots of non-caniniform teeth: roots absent or low, less than a third of the height of the crown (0), roots moderately developed, equal to or more than a third the height of the tooth crown (1), roots well-developed, taller than the tooth crown (2). ORDERED.**

New.

Most terminal taxa within this matrix possess little to no development of a tooth root within the alveolar shelves (State 0, pers. obv. RM 12016a, a specimen referred to *Hylonomus*). In many synapsids, (e.g., *Varanosaurus,* AMNH 4174), the roots are moderately developed, with bases of teeth enclosed within the alveolar shelves and bone of attachment (State 1). State 2 is considered an autapomorphyy of bolosaurids (Reisz et al. 2007).

UNAMBIGUOUS: Archosauromorpha (0 to 1). DELTRAN: Gymnarthridae (1 to 0), Recumbirostra (0 to 1), Caseidae (0 to 1), Varanopinae (0 to 1), Eupelycosauria (0 to 1), Bolosauridae (0 to 2). ACCTRAN: Recumbirostra (0 to 1), Eupelycosauria + Varanopidae (0 to 1), Ophiacodontidae (1 to 0), Bolosauridae (0 to 2), Pareiasauromorpha (0 to 1), Mesosauridae (0 to 1).

**Ch. 23. Maxilla, alveolar shelf, height relative to width of non-caniniform tooth base: alveolar shelf thin, not taller than width of non-caniniform marginal tooth bases (0), alveolar shelf tall, dorsoventrally taller than the anteroposterior width of the non-caniniform tooth bases (1). (-) in taxa possessing teeth fused to the apex of the crista dorsalis (Ch. 28).**

New.

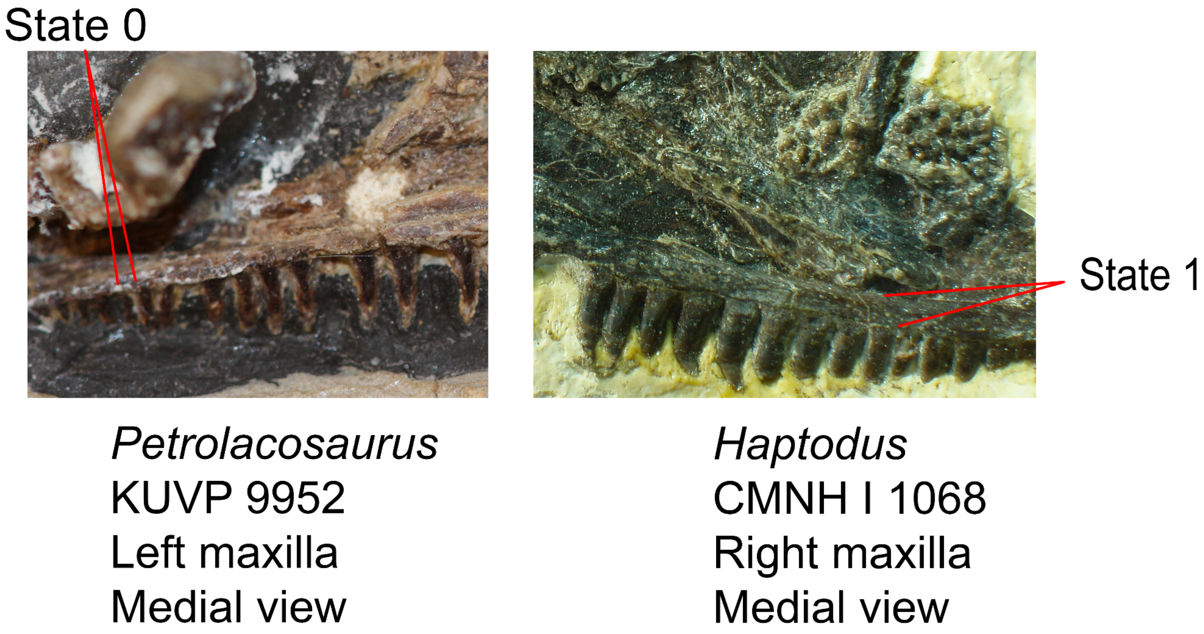


**Fig. S15.** The maxillae of *Petrolacosaurus* (KUVP 9952) and *Haptodus* (CNMH I 1068) in medial views. *Petrolacosaurus* exhibits an extremely dorsoventrally thin alveolar shelf (Ch. 23, State 0) in the posterior marginal dentition, as the alveolar shelf thins posteriorly whereas the alveolar shelf in *Haptodus* is dorsoventrally thickened and similar in size to the base of the marginal dentition (Ch. 23, State 1).

The alveolar shelves of the maxilla are dorsoventrally thin relative to the width of tooth bases in non-synapsid taxa within this sampling. In Synapsida, the alveolar shelf is dorsoventrally thickened relative to the base of the teeth in medial view, although this character can also be observed when the lateral surface of the maxilla has worn away (e.g., *Cutleria,* USNM 22099).

UNAMBIGUOUS: *Opisthodontosaurus carrolli* (0 to 1), *Araeoscelis* + Bolosauridae, 0 to 1, Pareiasauria + *Emeroleter* (0 to 1). DELTRAN: *Crassigyrinus scoticus* (0 to 1), Caseidae (0 to 1), Synapsida (0 to 1), Crocopoda (0 to 1), *Ascendonanus* + *Cabarzia* (0 to 1). ACCTRAN: *Crassigyrinus* + *Greererpeton* (0 to 1), CASEIDAE (0 to 1), Eupelycosauria + Varanopidae (0 to 1), Archosauromorpha (0 to 1), Parapleurota + *Orovenator* + *Ascendonanus* + *Cabarzi*a (1 to 0), Mesosauridae + *Ascendonanus* + *Orovenator* + *Cabarzia* + Parapleuroata (0 to 1).

**Ch. 24. Posterior marginal dentition, replacement teeth: absent (0), present (1).**

Modified from Pinna (1986, Ch. 26), Ford et al. (2021, Ch. 206), Simões et al (2022, Ch. 210).

Replacement teeth are absent in rhynchocephalian lepidosauromorphs within the current sampling (e.g., *Clevosaurus hudsoni*, NHMUK PV R36832).

UNAMBIGUOUS: Rhynchocephalia (0 to 1). DELTRAN: Squamata (1 to 0).

**Ch. 25. Posterior marginal dentition, replacement teeth, position in relation to functional teeth: lingual (0), posterolingual (1). Inapplicable (-) in taxa lacking replacement teeth (Ch. 24).**

Modified from Pinna (1986, Ch. 26), Ford et al. (2021, Ch. 207), Simões et al (2022, Ch. 211).

In many tetrapods possessing multiple tooth rows, tooth replacement occurs posterolingual to the functional tooth (State 1). This includes gymnarthrid recumbirostrans (Bolt and Demar 1983), captorhinids (Leblanc and Reisz 2014) and rhynchosaurs (e.g., *Mesosuchus*, SAM-PK-6536).

UNAMBIGUOUS: Captorhinidae (0 to 1), *Biseridens qilianicus* (0 to 1), *Mesosuchus browni* (0 to 1). DELTRAN: Gymnarthridae + *Nannaroter* + Rhynchonkidae + *Huskerpeton* (0 to 1) ACCTRAN: Recumbirostra (0 to 1)

**Ch. 26. Posterior marginal dentition, base, bone of attachment in lingual view: base of tooth in lingual view is not covered by cementum or 'bone of attachment' (0), base enclosed by woven "bone of attachment" or cementum (1).**

New.

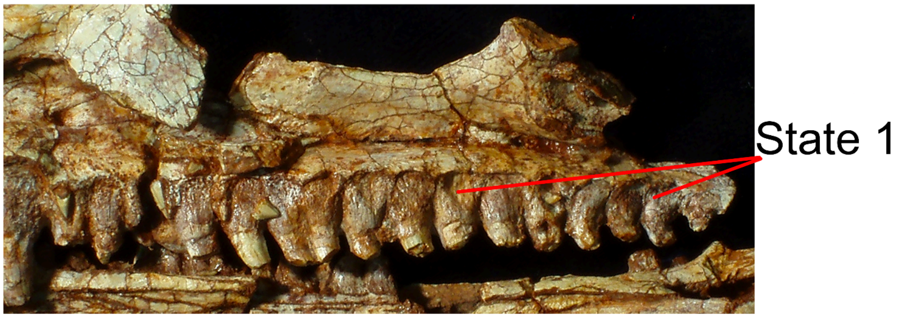


*Varanops*
TMM 43628-1
Right maxilla
Medial view

**Fig. S16.** Maxilla of *Varanops brevirostris* (TMM 43628-1) demonstrating (Ch. 26, State 1), or the presence of cementum or bone of attachment covering the base of the marginal dentition in lingual view.

The tooth bases of many tetrapods (e.g.,*Oromycter*, FMNH PR 2881) are covered by 'bone of attachment' or cementum in medial view (State 1).

UNAMBIGUOUS: *Varanodon agilis* (1 to 0), *Petrolacosaurus kansensis* (1 to 0). DELTRAN:, *Owenetta kitchingorum* (1 to 0), *Nyctiphruretus acudens* (1 to 0), *Orovenator mayorum* (1 to 0) *Paliguana whitei* (0 to 1), Archosauromorpha (0 to 1), *Paleothyris acadiana* (1 to 0). ACCTRAN: Tanystropheidae + Crocopoda (0 to 1), Mesenosaurinae (0 to 1), Mesosauridae + *Ascendonanus* + *Orovenator* + *Cabarzia* + Parapleurota (1 to 0), Protorothyrididae (1 to 0).

**Ch. 27. Posterior marginal dentition, bone of attachment, if present: tooth base partially enclosed by woven bone that does not form paradental plates between the lingual/medial surfaces of the tooth bases/sockets (0), tooth base enclosed by well-defined bone of attachment forming paradental plates (1).**

New.

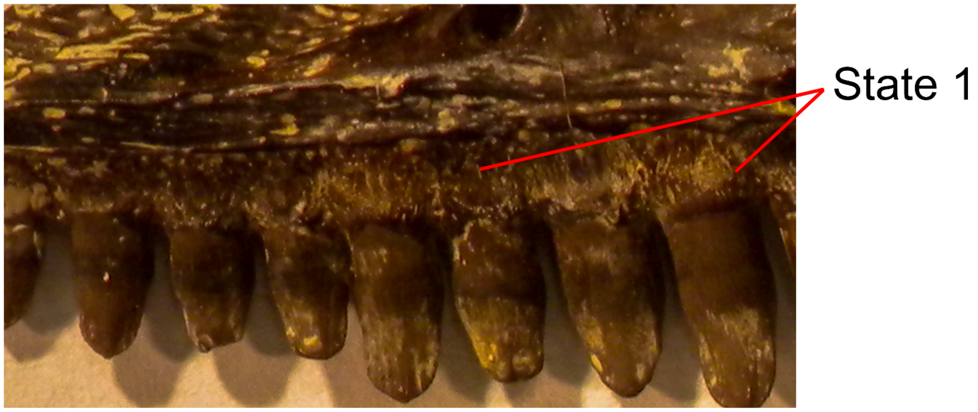


*Oromycter*
FMNH PR 2281
Left maxilla
Medial view

**Fig. S17.** Maxilla of *Oromycter* (FMNH PR 2281) in medial view demonstrating (Ch. 27, State 1), or the presence of cementum or bone of attachment forming paradental plates between the base of each tooth.

In most tetrapods, bone of attachment is restricted to the tooth root and does not form a consistent lamina between teeth or 'paradental plates' (State 0, e.g., *Varanops*, TMM 43628-1). However, in some archosauriforms (e.g., *Proterosuchus*, NMQR 1484), therapsids (Sidor and Hopson 1998), and caseid synapsids (*Oromocyter*, FMNH PR 2881), the bone of attachment forms a discrete paradental plate in medial view (State 1).

UNAMBIGUOUS: Ankyramorpha (0 to 1), Archosauromorpha (0 to 1). DELTRAN: *Crassigyrinus scoticus* (1 to 0), Caseidae (0 to 1), Sphenacodontoidea (0 to 1), *Belebey vegrandis* (0 to 1), *Tseajaia* + *Orobates* (1 to 0), Temnospondyli (1 to 0). ACCTRAN: Caseidae (0 to 1), Sphenacodontoidea + *Haptodus* + *Cutleria* + *Tetraceratops* (0 to 1), Araeoscelidia (0 to 1), *Seymouria* spp. (0 to 1), Temnospondyli + Reptiliomorpha (1 to 0).

**Ch. 28. Posterior marginal dentition, the position of tooth base relative to crista dorsalis (apex of labial wall of dentary): lingual (0), apical or traditionally 'acrodont' (1).**

Modified from Gauthier 1988 (Ch. 75), Simões et al. 2022 (Ch. 213).

The large majority of taxa in this sampling possess tooth bases that are located lingual to the crista dorsalis (State 0). State 1 is an autapomorphy of captorhinids (Leblanc and Reisz 2015), procolophonians (e.g., *Procolophon*, BP/1/5927), and rhynchocephalians (Fraser 1982).

UNAMBIGUOUS: *Pantylus cordatus* (0 to 1), Pareiasauria + *Emeroleter* (0 to 1), Procolophonoidea + *Nyctiphruretus* (0 to 1). DELTRAN: *Llistrofus* + Chthonosauria (0 to 1, Captorhinidae (0 to 1), Squamata (0 to 1). ACCTRAN: Gymnarthridae (0 to 1) Rhynchocephalia, (0 to 1), Captorhinidae (0 to 1).

**Ch. 29. Posterior marginal dentition, "complete" ankylosis to crista dorsalis (apex of labial wall) of dentary: absent (0), present (1).**

Modified from Simões 2018 (Ch.11), Simões et al. 2022 (Ch. 214).

Dentary dentition is completely ankylosed to the apex of the labial wall of the dentary in rhynchocephalians, several procolophonians (e.g., *Eomurruna,* Hamley et al. 2021), and the early-diverging captorhinid *Opisthodontosaurus* (Haridy et al. 2018).

UNAMBIGUOUS: *Opisthodontosaurus carrolli* (0 to 1). DELTRAN: *Eomurruna yurrgensis* (0 to 1), *Sauropareion anoplus* (0 to 1), Squamata (0 to 1), *Mesosuchus browni* (0 to 1). ACCTRAN: Leptopleuroninae + *Procolophon* + *Eomurruna* (1 to 0) Procolophonoidea + *Nyctiphruretus* (0 to 1), Rhynchocephalia (0 to 1), *Trilophosaurus* + *Mesosuchus* (0 to 1).

**Ch. 30. Premaxilla, number of teeth: five or more (0), four (1), three to one (2), absent (3). ORDERED.**

Modified from Brocklehurst et al. (2022, Ch. 12).

**Ch. 31. Premaxilla, orientation of ascending ( supranarial) process in respect to the marginal tooth row of the premaxilla: slopes anterodorsally, overhanging tooth row (0), subvertical at base, then slopes posterodorsally (1), slopes posterodorsally at an angle < 75° (2). ORDERED.**

Modified from Brocklehurst et al. (2022, Ch. 13).

**Ch. 32. Premaxilla, subnarial process excludes maxilla from margin of external naris or septomaxilla : absent (0), present (1).**

Modified from Brocklehurst et al. (2022, Ch. 14).

**Ch. 33. Premaxilla, posterior surface of subnarial process, if present: contacts the lacrimal only (0), contacts the nasal (1).**

Modified from Brocklehurst et al. (2022, Ch. 15).

**Ch. 34. Premaxilla, external suture with maxilla: suture between elements non-interdigitated (0), interdigitating suture present (1).**

Modified from Callier et al. (2009 Ch. 11), Clack et al. (2012 Ch. 11), Pardo et al. (2017 Ch. 266).

The maxilla of early tetrapods generally weakly overlaps the lateral surface of the premaxilla (e.g., *Youngina*, BP/1/2871) or makes a point contact with this element. In early temnospondyls (e.g., *Edops*, MCZ 1378), diadectomorphs (e.g., *Limnoscelis*, YPM 811), and some sphenacodontians (e.g., *Dimetrodon*, Romer and Price 1940) this contact takes the form of an interdigitating suture (State 1).

UNAMBIGUOUS: *Cacops morrisi* (1 to 0), Pareiasauria + *Emeroleter* (0 to 1) *Procolophon trigoniceps* (0 to 1), Temnospondyli (0 to 1). DELTRAN: *Crassigyrinus scoticus* (0 to 1), *Baphetes* spp. (0 to 1), Sphenacodontinae (0 to 1), Dinocephalia (0 to 1), *Mesosuchus browni* (0 to 1), Diadectomorpha + Protorothyrididae + Amniota (0 to 1). ACCTRAN: *Greererpeton burkemorani* (1 to 0), *Dimetrodon* (0 to 1), Dinocephalia + *Biseridens* (0 to 1), *Trilophosaurus* + *Mesosuchus* (0 to 1), Diadectomorpha (0 to 1).

**Ch. 35. Premaxilla, supranarial ( dorsal) process: short, ends anterior to posterior margin of external naris (0), elongate and posterior beyond the posterior margin of the external naris (1), supranarial process absent (2).**

Modified from Brocklehurst et al. (2022, Ch. 16).

**Ch. 36. Premaxilla, suture with nasal: single pronged or transversely flat (0), two-pronged (alary process present) (1), transversely broad or interdigitated suture (more than two prongs present) (2).**

Modified from Brocklehurst et al. (2022, Ch. 7).

Rationale: Added State 2, which concerns the transversely broad and often interdigitated suture between the dorsal process of the premaxilla and the nasal seen in early tetrapods and temnospondyls (e.g., *Edops*, Romer and Witter 1942).

**Ch. 37. Premaxilla, supranarial process, dorsal fontanelle: absent (0), present, medial margin of supranarial process is concave, forming a fontanelle with contralateral premaxilla (1).**

Modified from Callier (2009 Ch. 88), Clack et al. (2012, Ch. 88), Pardo et al. (2017 Ch. 341).

Tetrapodomorphs and early tetrapods, including some temnospondyls (e.g., *Dendrepeton*, NSM 987 GF 99.1) possess State 1.

UNAMBIGUOUS: *Crassigyrinus* + *Greererpeton* (0 to 1), *Dendrerpeton* + *Balanerpeton* +*Acheloma* + *Cacops* (0 to 1), *Chroniosaurus dongusensis* (0 to 1).

**Ch. 38. Premaxilla, slope of alveolar margin relative to ventral margin of maxilla: slopes anterodorsally (0), parallel (1), slightly anteroventral, less than 10° from horizontal (2), moderately anteroventral, 10-20° from horizontal (3), strongly anteroventral, 30-45° from horizontal (4), very strongly anteroventral > 45° from horizontal (5). ORDERED.**

Modified from Brocklehurst et al. (2022, Ch. 18).

**Ch. 39. Premaxilla, narial shelf: more or less sharp edge between lateral (= sculptured, if sculpture is present) surface of skull and ventral wall of naris (0), rounded ventral narial shelf that transitions smoothly into ventral edge of skull (1).**

Modified from Brocklehurst et al. (2022, Ch. 19).

**Ch. 40. Premaxilla, tooth size compared to marginal maxillary teeth: small or subequal to (non-caniniform) maxillary teeth (0), non-caniniform premaxillary teeth larger (1).**

Modified from Brocklehurst et al. (2022, Ch. 20).

**Ch. 41. Premaxilla, tooth size: isodont (0), anterior teeth larger than posterior teeth (1).**

Modified from Brocklehurst et al. (2022, Ch. 21).

**Ch. 42. Premaxilla, palatal process or medial flange: shorter than maxillary process or absent (0), well-developed, extends posterior to the level of the maxillary process (1).**

Modified from Reisz et al. (1992 Ch. 4,) Laurin (1993 Ch. 3), Sidor and Hopson (1998 Ch. 3) Frobisch et al. (2011, Ch. 2).

A well-developed posterior or palatal process of the premaxillae which extends posteriorly further than the maxillary process of the maxilla (State 1) is present in the early tetrapodomorph *Whatcheeria* (FM PR 1634)*,* mesosaurids (e.g., *Mesosaurus* SAM-PK-K8381), and procolophonians (e.g., *Procolophon*, Carroll and Lindsay 1985).

UNAMBIGUOUS: Procolophonoidea + *Nyctiphruretus* (0 to 1), *Crassigyrinus* + *Greererpeton* + *Baphetes* + Temnospondyli + Reptiliomorpha (1 to 0). DELTRAN: *Deltavjatia rossica* (0 to 1), Pareiasauria (0 to 1), *Mesosaurus tenuidens* (0 to 1). ACCTRAN: *Bunostegos akokanensis* (1 to 0), Pareiasauromorpha + *Australothyris* (0 to 1), Mesosauridae (0 to 1).

**Ch. 43. Premaxilla, palatal surface, intermaxillary fossa: absent (0), present (1). (-) in taxa lacking a palatal process of the premaxilla (Ch. 42)**

Modified from Anderson et al. (2009, Ch. 95), Huttenlocker et al. (2013, Ch. 95), Pardo et al. (2017, Ch. 198).

An intermaxillary fossa or fenestra between the midline contact of the palatal processes of the premaxillae (State 1) is present in early tetrapodomorphs (e.g., *Crassigyrinus,* Cowdenbeath skull of Panchen 1985) and early-diverging temnospondyls (e.g., *Dendrerpeton,* NSM 987 GF 99.1).

UNAMBIGUOUS: *Dendrerpeton* + *Balanerpeton* + *Acheloma* + *Cacops* (0 to 1). DELTRAN: *Baphetes* spp. (1 to 0), Temnospondyli + Reptiliomorpha (1 to 0). ACCTRAN: *Crassigyrinus* + *Greererpeton* (0 to 1).

**Ch. 44. Premaxilla, palatal surface, intermaxillary fossa, if present: paired (0), unpaired (1). Inapplicable (-) in taxa lacking an intermaxillary fossa (Ch. 43).**

Modified from Pardo et al. (2017, Ch. 199).

When an intermaxillary fossa of the premaxilla is present, it is paired in early tetrapodomorphs (e.g., *Greererpeton*, CMNH 11079) and unpaired in temnospondyls (e.g., *Cacops*, OMNH 53073).

DELTRAN: *Dendrerpeton* + *Balanerpeton* + *Acheloma* + *Cacops* (0 to 1). ACCTRAN: Temnospondyli + Reptiliomorpha (0 to 1)

**Ch. 45. Premaxilla, contribution to choana or internal nares: absent or weak, premaxilla contributes to the choana via weak point contact (0), present, premaxilla deeply emarginated such that it forms the anterior, medial, lateral and/or medial margins of choana (1).**

This study.

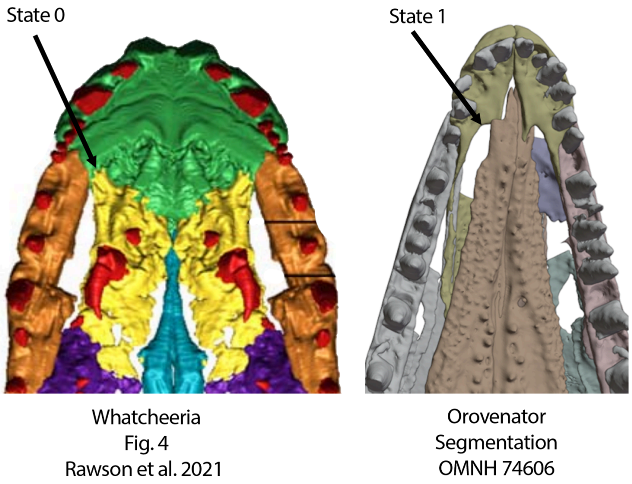


**Fig. S18.** Left, antorbital skull in palatal view of *Whatcheeria* (Modified from fig. 4 of Rawson et al. 2021) demonstrating the absence of a contribution of the premaxilla to the internal naris or choana (Ch. 45, State 0). Right, anterior palate of *Orovenator* demonstrating a broad contribution of the premaxilla to the choana (Ch. 45, State 1).

The premaxilla of early tetrapods is excluded from (or weakly contributes to) the internal nares (e.g., *Whatcheeria* FM PR 1634) (State 0). All known seymouriamorphs (Laurin 1996), diadectomorphs (e.g., *Limnoscelis,* YPM 811), recumbirostrans (e.g., *Hapsidopareion*, FMNH 2303), and ‘conventional’ early amniotes possess a broad contribution of the premaxilla to the internal naris (State 1).

UNAMBIGUOUS: *Lanthanosuchus* (1 to 0), Temnospondyli (1 to 0). DELTRAN: *Mesosaurus tenuidens* (1 to 0), Archosauriformes (1 to 0), Embolomeri (0 to 1). ACCTRAN: Allokotosauria + *Mesosuchus* (1 to 0), Mesosauridae (1 to 0),Temnospondyli + Reptiliomorpha (0 to 1).

**Ch. 46. Septomaxilla, morphology in lateral view: forms the posteroventral margin of the external naris (0), a rectangular sheet of bone shaped as a conical funnel (1), an anteroposteriorly broad sheet of bone that fully or partially partitions the external naris (2), a pillar of bone, partitioning the naris or forming the posterior margin of the external naris (3).**

Modified from Brocklehurst et al. (2022, Ch. 22).

**Ch. 47. Maxilla, premaxillary process, dorsal margin, narial shelf: absent, anterior portion of the maxilla is flat or convex (0), present, anterior portion of maxilla possesses an anteriorly directed narial shelf (1)**

Modified from Reisz et al. (1998, Ch. 2), Maddin et al. (2006, Ch. 6), Reisz et al. (2009 Ch. 5), Benson (2012, Ch. 15).

In most tetrapods, the premaxillary process of the maxilla is unexpanded at its contribution to the external narial margin (e.g., *Milleropsis,* BP/1/720) (State 0). In contrast, the dorsal margin of the premaxillary process of the maxilla in ankyramorph reptiles (e.g., *Acleistorhinus* FM UR 1038) and caseasaurian synapsids (e.g., *Oromycter*, FMNH PR 2281*)* possesses an anterodorsally-oriented narial shelf or fossa at the external naris (State 1).

UNAMBIGUOUS: Caseidae (0 to 1), Pareiasauria + *Emeroleter* (0 to 1). DELTRAN: Procolophonia (0 to 1), *Deltavjatia* + *Bunostegos* (0 to 1). ACCTRAN: Owenettidae (1 to 0), Ankyramorpha + *Erpetonyx* (0 to 1).

**Ch. 48. Maxilla, postnarial fossa or depression: absent, lateral surface maxilla is convex or flat in lateral view (0), present, narial shelf expanded as a lateral fossa posterior to external nares (1). Inapplicable (-) in taxa lacking a narial shelf of the maxilla (Ch. 46).**

Modified from Modesto et al. (2001, Ch. 5), Cisneros (2008, Ch. 5).

Among taxa that possess narial shelf of the maxilla, in non-leptopleuronine procolophonoids the narial shelf is expanded as a lateral fossa posterior to the external nares and is broadly visible in lateral view (e.g., *Coletta seca,* GHG 228).

UNAMBIGUOUS: Leptopleuroninae (1 to 0). DELTRAN: Procolophonoidea + *Nyctiphruretus* (0 to 1), *Nyctiphruretus acudens* (0 to 1). ACCTRAN: Leptopleuroninae (1 to 0), Procolophonia (0 to 1).

**Ch. 49. Maxilla, preorbital ascending process: convex dorsal margin that does not divide lacrimal (0), divides lacrimal in lateral view (1).**

Modified from Brocklehurst et al. (2022, Ch. 23).

**Ch. 50. Maxilla, preorbital ascending process, maximum dorsoventral height in respect to the height of the maxilla at the anterior orbital margin: less than 1.5 times the height (0), between 1.5 and twice the height (1), greater than twice the height (2). ORDERED.**

Modified from Brocklehurst et al. (2022, Ch. 24).

**Ch. 51. Maxilla, position of apex of preorbital ascending process: process absent or very low (0), tallest point located centrally between the orbit and external naris (1), tallest point located anteriorly, immediately posterior to the external naris (2), tallest point located anterior to the external naris (3), tallest point located immediately anterior to orbit (4). ORDERED.**

Modified from Brocklehurst et al. (2022, Ch. 25).

**Ch. 52. Maxilla, lacrimal facet at base of dorsal process: absent (0), present, distinct dorsoventral ridge present on ascending process divides anterior and posterior depressions (1).**

Modified from Brocklehurst et al. (2022, Ch. 302).

**Ch. 53. Maxilla and lacrimal, antorbital fenestra: absent, maxilla remains in tight articulation with lacrimal or other antorbital elements dorsally (0) present, gap or fenestra present between maxilla, lacrimal, or other antorbital elements (1).**

Modified from Juul (1994), Dilkes (1998, Ch. 5), Nesbitt (2011, Ch. 136), Dilkes and Arcucci (2012, ch. 2), Pritchard et al. (2015, Ch. 13), Ezcurra (2016, Ch. 13).

State 1 is an autapomorphy of all archosauriformes within this sampling (Nesbitt 2011).

UNAMBIGUOUS: Archosauriformes (0 to 1).

**Ch. 54. Maxilla, teeth, interdental plates: contact both labial and lingual walls of alveolar groove, fully separating adjacent teeth (0), do not contact labial and lingual walls of alveolar groove or are absent (1)**

Modified from Pritchard et al. (2015, Ch. 96) and Ford et al. (2021, Ch. 378).

Interdental plates fully separating adjacent alveoli are present in derived sphenacodontians (e.g., *Secodontosaurus*, FMNH UC 754) and archosauriformes within this sampling (State 1).

UNAMBIGUOUS: Caseidae (1 to 0), Procolophonoidea + *Nyctiphruretus* (1 to 0), Archosauromorpha (1 to 0). DELTRAN: Therapsida (1 to 0), Bolosauridae (1 to 0), *Deltavjatia rossica* (1 to 0). ACCTRAN: Sphenacodontoidea + *Haptodus* + *Cutleria* + *Tetraceratops* (1 to 0), Bolosauridae (1 to 0), Pareiasauria + *Emeroleter* (1 to 0).

**Ch. 55. Maxilla, contact with prefrontal: absent, separated by the anterior extension of the lacrimal or nasal (0), present, lacrimal/nasal is reduced anteriorly allowing contact (1).**

Modified from Brocklehurst et al. (2022, Ch. 26).

**Ch. 56. Maxilla, row of lateral ( supralabial) maxillary foramina parallel to ventral margin of maxilla: absent (0), present (1).**

Modified from Brocklehurst et al. (2022, Ch. 27).

**Ch. 57. Maxilla, anterior lateral ( supralabial) maxillary foramen: equal in size to other maxillary foramina, if present (0), much larger than other foramina or a single large foramen (1).**

Modified from Brocklehurst et al. (2022, Ch. 28).

**Ch. 58. Maxilla, number of tooth rows: single row of marginal teeth only (0), 2-3 (1), four or more (2). ORDERED.**

Modified from Brocklehurst et al. (2022, Ch. 29).

**Ch. 59. Maxilla, if more than one tooth row present, size of tooth field: <40 (0), 40 or more (1).**

Modified from Brocklehurst et al. (2022, Ch. 30).

**Ch. 60. Maxilla, if more than 3 tooth rows present, double tooth row extending far anterior to more posterior tooth rows: absent (0), present, double row of teeth extends far anteriorly on tooth row (1). (-) in taxa lacking more than three maxillary tooth rows.**

Modified from Modesto et al. (2014, Ch. 11) and Reisz et al. (2011, Ch. 11).

A double row of teeth extends anterior relative to more posterior maxillary tooth rows in the moradisaurine captorhinids *Labidosaurikos* and *Rothianiscus* (Dodick and Modesto 1995).

UNAMBIGUOUS: *Labidosaurikos meachami* + *Rothianiscus* spp. (0 to 1).

**Ch. 61. Maxilla, marginal tooth number (count) or number of tooth stations: 35 or more (0), 28-34 (1), 12-27 (2), 8-11 (3), 1-7 (4). ORDERED.**

Modified from Brocklehurst et al. (2022, Ch. 30).

**Ch. 62. Maxilla, caniniform teeth: absent (0), present (1).**

Modified from Brocklehurst et al. (2022, Ch. 31).

**Ch. 63. Maxilla, caniniform teeth number (count): more than two caniniform teeth present (caniniform region) (0), one or two caniniform teeth present (1).**

Modified from Brocklehurst et al. (2022, Ch. 32).

**Ch. 64. Maxilla, caniniform teeth, size: slightly larger than the adjacent maxillary teeth (0), at least 1.5 times as large as adjacent maxillary teeth (1).**

Modified from Brocklehurst et al. (2022, Ch. 33).

**Ch. 65. Maxilla, secondary caniniform region: absent (0), present (1).**

Modified from Brocklehurst et al. (2022, Ch. 34).

**Ch. 66. Maxilla, number of pre-caniniform maxillary teeth: five or less (0), six or more (1).**

Modified from Brocklehurst et al. (2022, Ch. 35).

**Ch. 67. Maxilla, lateral supracanine buttress: absent or slight swelling (0), distinct lateral swelling or dorsally oriented ridge on the maxilla, dorsal to the caniniform dentition (1).**

Modified from Brocklehurst et al. (2022, Ch. 36).

**Ch. 68. Maxilla, supracanine buttress on medial surface: absent, alveolar shelf has uniform or gradually tapering dorsoventral height (0), present, subtriangular dorsal thickening of anterior alveolar shelf (1).**

Modified from Brocklehurst et al. (2022, Ch. 37).

**Ch. 69. Maxilla, morphology of dorsal portion of supracanine buttress, if present: anteroposteriorly broad region of thickened bone (0), narrow, strut-like ascending process extends posterodorsally from supracanine buttress (1).**

Modified from Brocklehurst et al. (2022, Ch. 38).

**Ch. 70. Maxilla, external nasal canal: maxillary canal ends blindly into a conical cavity or no dorsal development (0), maxillary canal opens externally into numerous foramina forming an 'external nasal canal' (1).**

New, based on observations stated in Benoit et al. (2021).

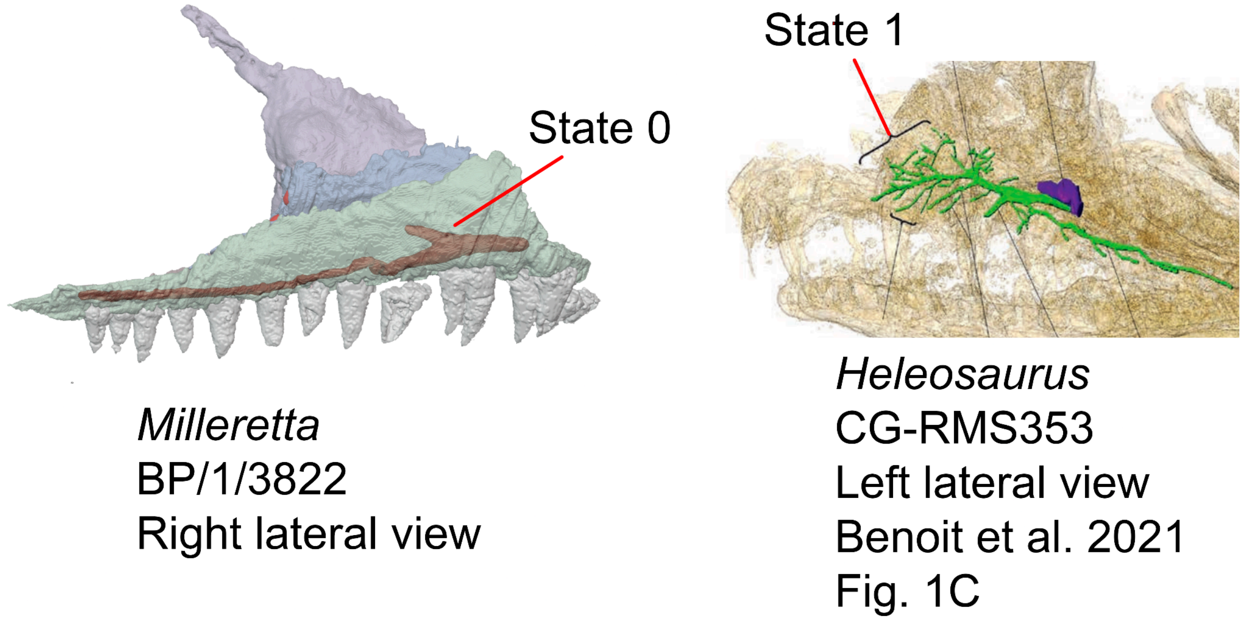


**Fig. S19.** Left, right antorbital skull of *Milleretta* (BP/1/3822, this study) demonstrating the presence of a relatively simple external nasal canal (Ch. 70, State 0) and right, left antorbital skull of *Heleosaurus* CM-RMS353 (Modified from Benoit et al. 2021, fig. 1c) demonstrating the presence of a complex external nasal canal with many foramina (Ch. 70, State 1).

Among early tetrapods, the maxillary canals of non-synapsid taxa such as the captorhinid *Captorhinus* (pers. Obv. OMNH 44816), the araeoscelidian *Araeoscelis* (pers. obv scans of AMNH FARB 4686), *Orovenator* (Benoit et al. 2021), the millerettid *Milleropsis* (pers. obv. scans of BP/1/720), and the archosauromorph *Prolacerta* (Benoit et al. 2021) is a simple structure, extending anteroposteriorly with a slight posterodorsal tube-like development termed a conical cavity (State 0). In contrast, the maxillary canal of early synapsids, including varanopids (Benoit et al. 2021, this study), caseasaurs such as *Cotylorhynchus*, and the edaphosaurid *Edaphosaurus* (pers. obv FMNH UC 674) is complex, opening dorsally into numerous foramina termed the external nasal canal (State 1, Benoit et al. 2021).

UNAMBIGUOUS: Synapsida (0 to 1).

**Ch. 71. Maxilla, diastema present between anteriormost dentition and rest of tooth row: diastema absent (0), large diastema present between first two maxillary teeth and more posterior teeth (1).**

Modified from Liu (2023, Ch. 79).

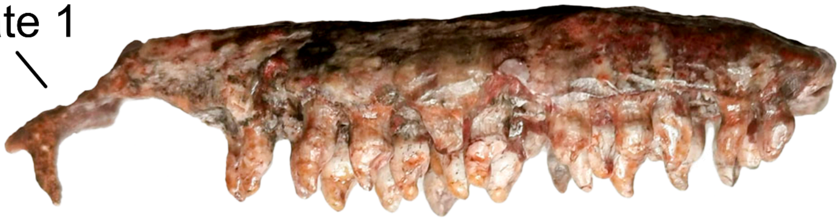


**Fig. S20.** Left maxilla of *Gansurhinus qingtoushanensis* (IVPP V15989) in medial view, demonstrating (Ch. 71, State 1), the presence of a diastema between the first maxillary tooth position and those more posterior to it. The rostral tip of this maxilla has been damaged (Liu 2023).

Captorhinids in the genus *Gansurhinus* possess a large diastema between the first two maxillary teeth and more posteriormost teeth (Lui et al. 2023). This character specifically refers to a diastema or gap between maxillary teeth, and not a diastema between the premaxillary and maxillary tooth rows.

DELTRAN: *Gansurhinus qingtoushanensis* + *Gansurhinus naobaogouensis* (0 to 1). ACCTRAN: *Gansurhinus qingtoushanensis* + *Gansurhinus naobaogouensis* + *Tramuntanasaurus tiai* (0 to 1).

**Ch. 72. Maxilla, orientation of lateral antorbital surface: vertical or slopes weakly dorsomedially (0), slopes dorsolaterally, overhanging tooth row or triturating surface (1).**

Modified from Brocklehurst et al. (2022, Ch. 39).

**Ch. 73. Maxilla, orientation of lateral surface of suborbital ramus: vertical, with no lateral overhang of tooth row (0), dorsolaterally inclined, overhangs tooth row (1).**

Modified from Brocklehurst et al. (2022, Ch. 40).

**Ch. 74. Maxilla, outline in dorsal or ventral view: relatively straight or mildly convex (0), posterior end flexed laterally resulting in concave outline seen in dorsal/ventral view (1).**

Modified from Brocklehurst et al. (2022, Ch. 41).

**Ch. 75. Maxilla, posterior extension: maxilla terminates posterior to the posterior orbital rim (0), anterior to or level with the posterior orbital rim but posterior to anterior orbital rim (1), anterior to anterior orbital rim (2). ORDERED.**

Modified from Brocklehurst et al. (2022, Ch. 42).

**Ch. 76. Maxilla, posterior extent of dentition if posterior extension level with or posterior to posterior orbital rim: continues posterior to the posterior orbital rim (0), terminates ventral to the posterior orbital rim (1), anterior to the posterior orbital rim exclusively (2). ORDERED.**

Modified from Brocklehurst et al. (2022, Ch.43).

**Ch. 77. Maxilla, ventral surface: straight or weakly convex (0), strongly convex with prominent precanine step anteriorly (1), pronounced convexity but no 'precanine step' anteriorly (2). ORDERED.**

Modified from Brocklehurst et al. (2022, Ch. 44).

**Ch. 78. Maxillary and dentary alveolar ridges: straight (0), twisted (helical), distal teeth inclined laterally (1).**

Modified from Brocklehurst et al. (2022, Ch. 304).

**Ch. 79. Maxilla and lacrimal, anteroventrally directed ridge or swelling: absent (0), present, anteroventrally oriented ridge or swelling extending from lacrimal to maxilla (1).**

Modified from Van den Brandt (2020).

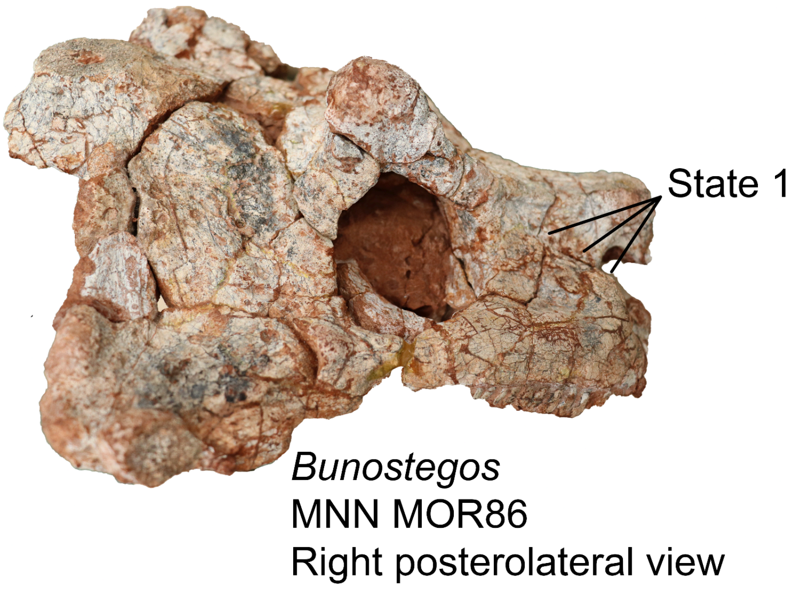


**Fig. S21.** Skull of *Bunostegos* (MNNN MOR86) in lateral view, demonstrating (Ch. 79, State 1) or the presence of a distinct anteroventrally trending ridge or swelling that extends from the lacrimal onto the maxilla and approaches the naris.

Pareiasaurs possess an anteroventrally directed ridge on the lateral surface of the lacrimal and maxilla, sometimes creating a fossa between these elements and the nasals dorsally (State 1, e.g., *Nochelesaurus*, SAM-PK-K6239*)*.

UNAMBIGUOUS: Pareiasauria (0 to 1).

**Ch. 80. Maxilla, teeth, asymmetry of dorsoventral height of labial and lingual walls of alveolar groove: lingual walls subequal in height to labial wall (0), lingual wall absent or low, less than half the height of the labial wall (1).**

Modified from Ford et al. (2021 Ch. 378).

UNAMBIGUOUS: *Opisthodontosaurus carrolli* (1 to 0), *Haptodus garnettensis* (1 to 0), Procolophonoidea + *Nyctiphruretus* (1 to 0). DELTRAN: Bolosauridae (1 to 0) *Orovenator* + *Ascendonanus* (1 to 0), Tanystropheidae + Crocopoda (1 to 0), Tangasauridae (0 to 1), ACCTRAN: Bolosauridae (1 to 0), Crocopoda (1 to 0), Weigeltisauridae+ Tangasauridae + *Claudiosaurus* + Sauria (0 to 1), Mesenosaurinae(0 to 1).

**Ch. 81. Maxilla, teeth, dorsoventral height of lingual wall of alveolar groove, when asymmetry is present: low, less than half the height of the labial wall (0), absent (1). Inapplicable in taxa lacking asymmetry between the labial and lingual wall of alveolar groove (Ch. 79).**

Modified from Ford et al. (2021 Ch. 379).

DELTRAN: *Crassigyrinus scoticus* (1 to 0), *Edaphosaurus boanerges* (0 to 1), Lepidosauromorpha (0 to 1), Weigeltisauridae + *Claudiosaurus* + Sauria (0 to 1), *Ascendonanus* + *Cabarzia* (0 to 1), Amniota + Protorothyrididae + *Anthracodromeus* (1 to 0). ACCTRAN: *Crassigyrinus* + *Greererpeton* (1 to 0), Edaphosauridae + *Ianthodon* (0 to 1), Lepidosauria + *Marmoretta* + *Megachirella* (0 to 1), Weigeltisauridae (0 to 1), Mesosauridae (0 to 1), Amniota + Protorothyrididae + *Anthracodromeus* + *Hylonomus* + *Thuringothyris* (1 to 0).

**Ch. 82. Marginal teeth, bulbous base giving a teardrop-shaped outline: absent (0), present (1).**

Modified from Brocklehurst et al. (2022, Ch. 300).

**Ch. 83. External naris, posterodorsal expansion: absent (0), present (1).**

Modified from Brocklehurst et al. (2022, Ch. 45).

**Ch. 84. External naris, morphology of posterodorsal expansion if present: pinched between nasal and maxilla (0), greatly enlarged, between nasal and lacrimal (1).**

Modified from Brocklehurst et al. (2022, Ch. 46).

**Ch. 85. External naris, position: marginal (minimal distance between nares >0.35 snout width at same level) (0), close to midline (1).**

Modified from Brocklehurst et al. (2022, Ch. 47).
**Ch. 86. Nasal, length: distinctly shorter than the frontal (0), approximately equal to the frontal (1), longer than frontal (2).**

Modified from Brocklehurst et al. (2022, Ch. 48).

**Ch. 87. Nasal, contribution to external naris "external narial shelf": forms posterodorsal margin only (0), extends anteroventrally as a blade-like process ("external narial shelf") bearing a lateral fossa with rim (1).**

Modified from Brocklehurst et al. (2022, Ch. 49).

**Ch. 88. Lacrimal: present (0), absent as a distinct ossification (1)**

Modified from Estes (1988, Ch. 28).

The lacrimal is absent in many lepidosauromorphs within this sampling (e.g., *Eichstaettisaurus*, BSPG 1937).

DELTRAN: *Belliarsia* + Squamata (0 to 1), Squamata (0 to 1).

**Ch. 89. Lacrimal, contribution to external naris: broad (0), narrow (1), absent (2). ORDERED.**

Modified from Brocklehurst et al. (2022, Ch. 50).

**Ch. 90. Lacrimals, anterolateral margin, fossa or depression forming narial rim: absent (0), present (1).**

Modified from Brocklehurst et al. (2022, Ch. 305).

**Ch. 91. Lacrimal, contribution to orbital margin as seen in lateral view: large, greater than 20% or more of the orbital circumference (0), moderate, between 8-16% of the orbital circumference (1), small less than 5% of the orbital circumference or absent, restricted by a prominent ventral process of the prefrontal (2). ORDERED.**

Modified from Brocklehurst et al. (2022, Ch. 52).

**Ch. 92. Lacrimal, ventromedial process contributing to antorbital buttress: absent (0). present medial process of lacrimal contributes to junction of prefrontal and palatine at the anterior orbital margin (1). (-) in taxa lacking an antorbital buttress (Ch. 101).**

Modified from Reisz et al. (2007).

The lacrimal contributes to the antorbital buttress via a medial process in bolosaurids (e.g., *Belebey*, SGU 104/B-2020) and derived procolophonoids within this study (e.g., *Procolophon*, NHMUK PV R. 1949).

DELTRAN: Reptilia (1 to 0), *Tseajaia* + *Orobates* (0 to 1). ACCTRAN: Neoreptilia (1 to 0), Reptiliomorpha (0 to 1).

**Ch. 93. Lacrimal, duct: opens on posterior (orbital rim) surface of lacrimal (0), opens laterally near posterior edge of lacrimal (1), opens laterally on concave surface of lacrimal (2), opens on a notch between the lateral and posterior (orbital) surface of the lacrimal (3).**

Modified from Brocklehurst et al. (2022, Ch. 53).

**Ch. 94. Lacrimal, suture with jugal in lateral view: well developed, greater than 50% of the minimum dorsoventral height of the suborbital bar (0), small, less than 50% of the minimum dorsoventral height of the suborbital bar (1), absent (2). ORDERED.**

Modified from Brocklehurst et al. (2022, Ch. 55).

**Ch. 95. Prefrontal, contact with postfrontal or postorbital, if present: contact of prefrontal and postfrontal/postorbital excludes the frontal from the dorsal margin of the orbit (0), prefrontal and postfrontal/postorbital separated by the frontal along the dorsal margin of the orbit (1).**

Modified from Brocklehurst et al. (2022, Ch. 56).

**Ch. 96. Prefrontal, medial process in dorsal view: absent, suture with frontal relatively straight (0), present, prefrontal medially expanded constricting the anterior process of the frontal in dorsal view (1).**

Modified from Brocklehurst et al. (2022, Ch. 57).

**Ch. 97. Prefrontal, palatal contact: absent (0), present (1).**

Modified from Brocklehurst et al. (2022, Ch. 58).

**Ch. 98. Prefrontal, contact with palatine, if present, sutural contact: absent, prefrontal-palatal contact is weak or abutting (0), sutured (1). (-) Inapplicable in taxa lacking prefrontal-palatal contact (Ch. 97).**

Modified from Laurin and Reisz (1995 Ch. 5), MacDougall et al. (2019, Ch. 12), Cisneros et al. (2020, Ch. 12)

The prefrontal forms an interdigitating suture with the palatine in procolophonians (State 1, *Embrithosaurus*, SAM-PK-8034).

UNAMBIGUOUS: Procolophonoidea + *Nyctiphruretus* (0 to 1), Owenettidae (0 to 1). DELTRAN: *Acheloma* + *Cacops* (0 to 1), *Belebey vegrandis* (0 to 1). ACCTRAN: Bolosauridae (0 to 1), Temnospondyli (0 to 1).

**Ch. 100. Prefrontal, foramen orbitonasale along prefrontal palatal contact: absent, or foramen between lacrimal and palatine (0), present (1).**

Modified from Laurin and Reisz (1995, Ch. 10).

**Ch. 101. Prefrontal, suture with nasal: parasagittal, at least in its caudal third (0), anterolateral (1), transverse, forming right angle to parasagittal plane (1).**

Modified from Brocklehurst et al. (2022, Ch. 60).

**Ch. 102. Prefrontal, lateral surface: approximately flat or convex (0), concave (1).**

Modified from Brocklehurst et al. (2022, Ch. 61).

**Ch. 103. Prefrontal, lateral expansion overhangs the lateral surface of the skull excluding supraorbital shelf or 'horns': absent, lateral surface of the prefrontal is flat and bears no lateral expansion or swelling (0), present, prefrontal swells laterally overhanging the lacrimal and maxillary tooth row (1).**

This study.

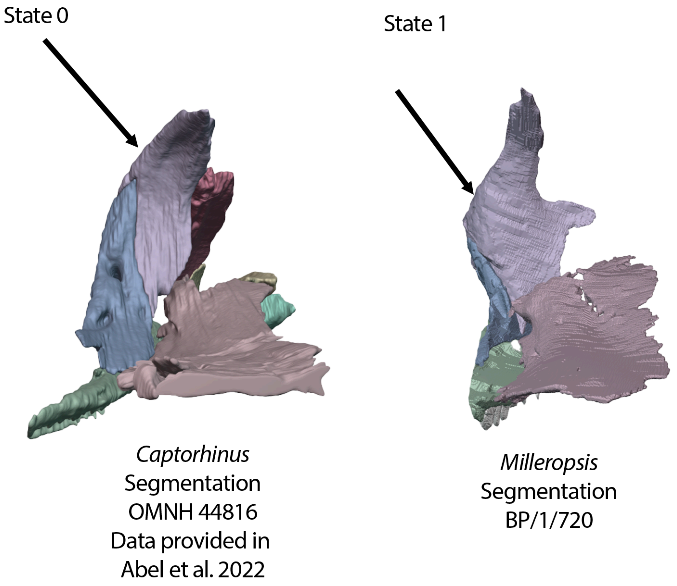


**Fig. S22.** Left, anterior orbit wall of *Captorhinus* (OMNH 44816) in posterior view, demonstrating, or the absence of a lateral expansion of the prefrontal (Ch. 103, State 0). Right, anterior orbit wall of *Milleropsis pricei* (BP/1/720)

in posterior view demonstrating a swollen or laterally expanded prefrontal that overhangs the lacrimal and maxilla (Ch. 103, State 1).

The prefrontal of millerettids (e.g., *Milleretta*, BP/1/3822) swells laterally, such that it overhangs the lacrimal and maxillary tooth row laterally (State 1).

DELTRAN: *Eothyris parkeyi* (0 to 1), *Orovenator* + *Ascendonanus* + *Cabarzia* (0 to 1), *Milleretta* + *Millerosaurus* + *Milleropsis* + *Broomia* (0 to 1), *Mesosuchus browni* (0 to 1), *Weigeltisaurus jaekeli* (0 to 1). ACCTRAN: *Thuringothyris* + Captorhinidae (0 to 1) *Mesosuchus browni* (0 to 1), *Weigeltisaurus jaekeli* (0 to 1), Millerettidae (0 to 1).

**Ch. 104. Prefrontal, sharp angle between the dorsal and the lateral surface: absent (0), present (1).**

Modified from Brocklehurst et al. (2022, Ch. 62).

**Ch. 105: Prefrontal, anterior process, contribution to external naris: absent, anterior process of prefrontal excluded from nares by the nasal, lacrimal, or maxilla (0), present, anterior process of prefrontal contributes to external naris (1).**

Modified from Pardo et al. (2017, Ch. 25).

**Ch. 106. Circumorbital bones, raised orbitotemporal crest along orbital margin: absent (0), present (1).**

Modified from Anderson et al. (2008, Ch. 44), Huttenlocker et al. (2013, Ch. 44), Pardo et al. (2017, Ch. 40).

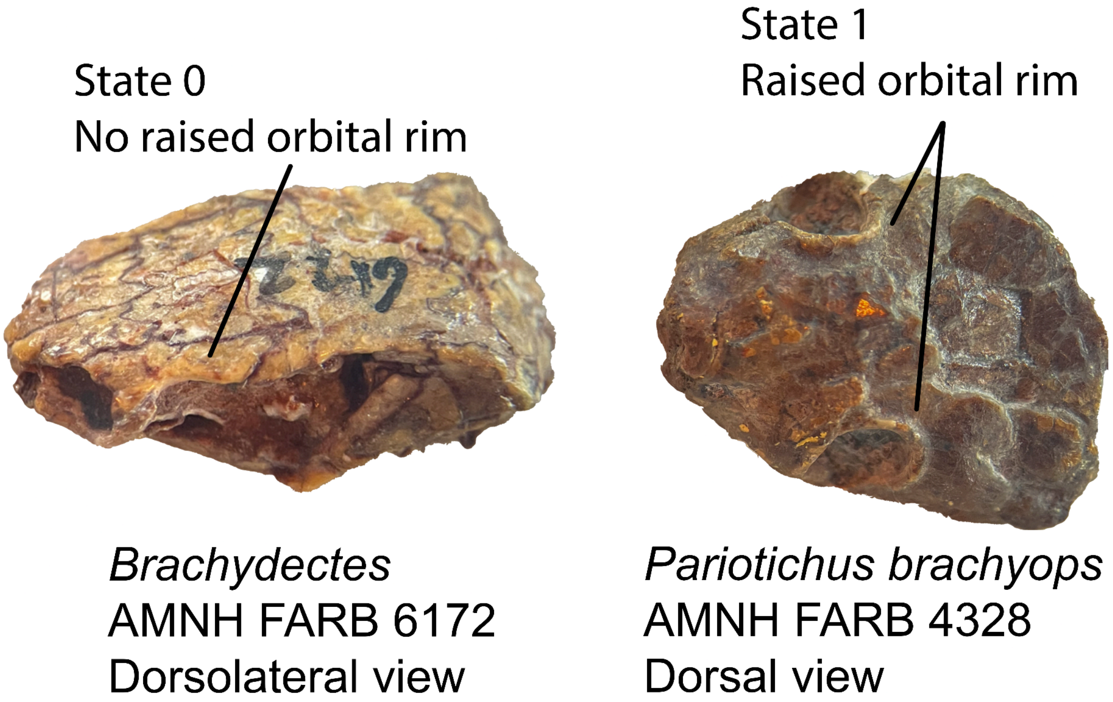


**Fig. S23.** Left, skull of *Brachydectes* (AMNH FARB 6172) in dorsolateral view, lacking a raised orbital rim (Ch, 106, State 0). Right, skull of *Pariotichus* (AMNH FARB 4328) in dorsal view which bears a distinct orbital ridge on the circumorbital elements (Ch, 106, State 1).

A distinct orbitotemporal crest (a raised ridge along the orbital and sometimes temporal margins) is present on the circumorbital bones (prefrontal, frontal, postfrontal) in several recumbirostrans (e.g., *Rhynchonkos*, FM-UR 1039) and procolophonoids (e.g., *Saurodektes*, BP/1/4195) in this sampling (State 1).

DELTRAN. Rhynchonkidae + *Huskerpeton* (0 to 1), Leptopleuroninae (0 to 1), *Nyctiphruretus acudens* (0 to 1), *Mesenosaurus romeri* (0 to 1), *Elliotsmithia longiceps* (0 to 1). Gymnarthridae + *Nannaroter* + Rhynchonkidae + *Huskerpeton* (0 to 1). Procolophonoidea + *Nyctiphruretus* (1 to 0), Procolophonia (0 to 1), *Heleosaurus scholtzi* (1 to 0), Mesenosaurinae (0 to 1).

**Ch. 107. Frontal, contribution to orbital margin, when present: narrow (0), broad, > 1/4 of maximum orbital length (1).**

Modified from Brocklehurst et al. (2022, Ch. 63).

**Ch. 108. Frontal, contribution to orbital margin, if present, formed by lateral lappet: absent (0), present, lateral extension of bone projects from the main body of frontal, contributing to orbital margin (1).**

Modified from Brocklehurst et al. (2022, Ch. 64).

**Ch. 109. Frontal, anteroposterior length at the midline: less than 1.5 times parietal length (0), 1.5-2.5 times parietal length (1), greater than 2.5 times parietal length (2). ORDERED.**

Modified from Brocklehurst et al. (2022, Ch. 65).

**Ch. 110. Frontal, mediolateral width: frontal anteroposteriorly longer than mediolaterally wide (0), frontal broader than long (1).**

Reworked from Brocklehurst et al. (2022, Ch. 297).

Rationale: The original description presents quantitative boundaries, but does not explain how to calculate them because frontal length on its own is not enough info to compute a ratio. We specify width here.

**Ch. 111. Frontal, posterolateral process: absent, frontoparietal suture forming right angle to parasagittal plane (0), short, only partially separating parietal from dorsolateral margin of postfrontal or forming obtuse posterolateral angle to parasagittal plane (1), long, narrow, frontoparietal suture forming acute posterolateral angle with parasagittal plane, substantially or fully separating parietal from dorsomedial margin of postfrontal (2). ORDERED.**

Modified from Brocklehurst et al. (2022, Ch. 66).

**Ch. 112. Frontal, anterior process length: short, not longer than the posterior process (0), longer than posterior process (1), very long, forming at least 2/3 of the anteroposterior length of bone (2). ORDERED.**

Modified from Brocklehurst et al. (2022, Ch. 67).

**Ch. 113. Frontal, anterior process, width: width equal to that of posterior process (0), narrower than posterior process (1).**

Modified from Brocklehurst et al. (2022, Ch. 307).

**Ch. 114. Frontal, fusion to contralateral frontal: unfused, suture patent (0), fused in midline, no clear suture patent (1).**

Modified from Benton (1985, Ch. Y1), Gauthier (1986), Pritchard et al. (2015, Ch. 14), Ezcurra (2016, Ch. 112).

The paired frontals are fused into a single, median element (State 1) in lepidosauromorphs (e.g., *Huehuecuetzpalil* IGM 7389) within this sampling.

ACCTRAN: Lepidosauromopha (0 to 1).

**Ch. 115. Frontal, ventral surface, descending flange articulating with the sphenethmoid or orbitosphenoid: absent (0), present, descending flange of frontal braces the sphenethmoid or orbitosphenoid dorsolaterally (1).**

Modified from Pardo et al. (2017, Ch. 195) and Huttenlocker et al. (2013, Ch. 223).

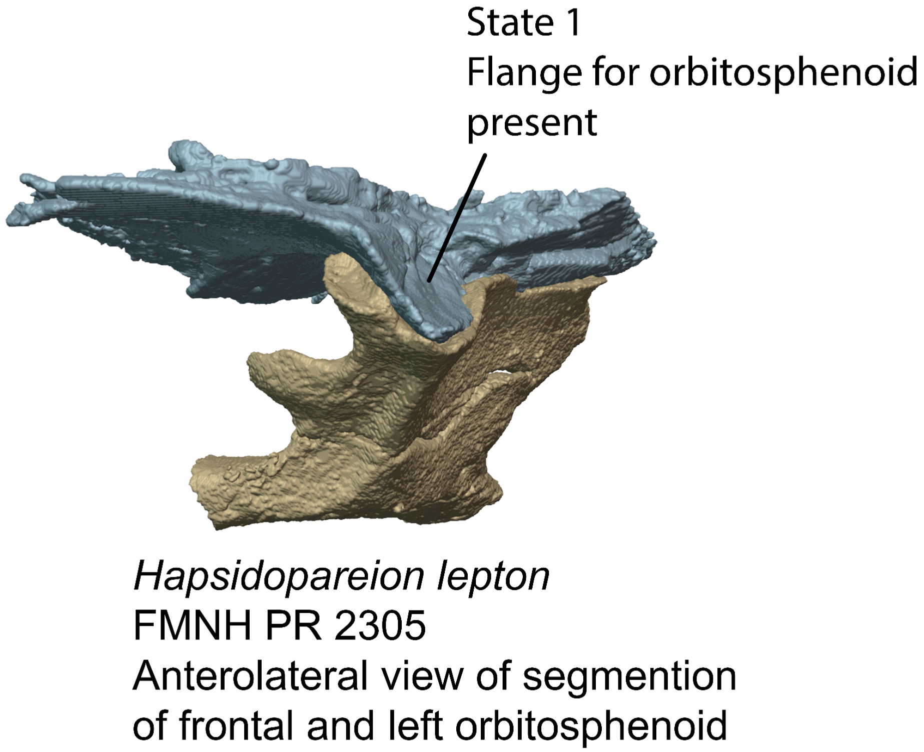


**Fig. S24.** Frontal and orbitosphenoid of *Hapsidopareion lepton* (FMNH PR 2305) in anterior view, demonstrating the descending flange of the frontal that articulates with the orbitosphenoid ventrally (Ch. 115, State 1).

UNAMBIGUOUS: *Brachydectes newberryi* (1 to 0), Recumbirostra (0 to 1).

**Ch. 116. Palpebral, ossification: absent or unossified (0), present, single large plate of bone above orbit (1)**

Modified from Huttenlocker et al. (2013, Ch. 49), Pardo et al. 2017 Ch. 45).

A palpebral, or a distinct, free-floating element near the dorsal orbital margin, is present in recumbirostrans (e.g., *Llistrofus*, OMNH 79031) within this sampling.

UNAMBIGUOUS: *Brachydectes newberryi* (1 to 0), Recumbirostra (0 to 1). DELTRAN: *Euryodus dalyae* (1 to 0), *Nannaroter mckinzei* (1 to 0) . ACCTRAN: *Cardiocephalus sternbergi* (0 to 1), Gymnarthridae + *Nannaroter* (1 to 0).

**Ch. 117. Postfrontal, lateral margin: narrow, contributing to the concave rim of the dorsal orbital margin (0), laterally expanded, forming a supraorbital shelf (1).**

Modified from Spindler et al. (2019 Ch. 43).

The lateral margin of the postfrontal is expanded laterally forming a supraorbital platform or shelf (State 1) in non-varanopine synapsids within this sampling (e.g., *Varanosaurus*, BSPHM 190).

DELTRAN: *Ennatosaurus tecton* (1 to 0), Caseidae (0 to 1), Eupelycosauria + Varanopidae (0 to 1), *Trilophosaurus buettneri* (0 to 1), *Orobates pabsti* (0 to 1), *Limnoscelis paludis* (0 to 1). ACCTRAN: *Ennatosaurus tecton* (1 to 0), Eupelycosauria (0 to 1), *Trilophosaurus buettneri* (0 to 1), *Tseajaia campi* (1 to 0), Diadectomorpha (0 to 1).

**Ch. 118. Postfrontal, free posterior margin: absent (0), present (1).**

Modified from Laurin (1991: A1,B3), Dilkes (1998 Ch. 22).

The postfrontal of early saurians, such as lepidosauromorphs (e.g., *Clevosaurus*, Fraser 1988) and early archosauromorphs (e.g., *Macrocnemus*, PIMUZ T 2477) possesses a free posterior margin where it contributes to the upper temporal fenestra (State 1).

DELTRAN: *Huehuecuetzpalli mixtecus* (1 to 0), *Trilophosaurus buettneri* (0 to 1), Tanystropheidae + Crocopoda (1 to 0), *Protorosaurus speneri* (1 to 0), *Claudiosaurus germaini* (1 to 0).

**Ch. 119. Postfrontal, posterodorsal process: present (0), absent, posterior margin of postfrontal flat or convex (1).**

Modified from Simões (2018 Ch. 47), Simões et al. (2022 Ch. 64).

UNAMBIGUOUS: Recumbirostra + Diadectomorpha + Protorothyrididae + Amniota + (1 to 0), Parapleurota (1 → 0).

**Ch. 120. Postorbital-postfrontal contact: overall trend approximately straight (0), incised by postorbital (1).**

Modified from Brocklehurst et al. (2022, Ch. 309).

**Ch. 121. Postorbital, lateral tuberosity at orbital margin: absent (0), present (1).**

Modified from Brocklehurst et al. (2022, Ch. 68).

**Ch. 122. Postorbital, free dorsal margin: absent (0), present (1). (-) in taxa lacking a free lateral margin of the parietal for the current sampling.**

Modified from Brocklehurst et al. (2022, Ch. 69).

**Ch. 123. Postorbital-squamosal, upper temporal bar: positioned distinctly ventral to the level of the dorsal border of the orbit, broadly exposing the supratemporal fenestra in lateral view (0), positioned approximately aligned to the dorsal border of the orbit and the supratemporal fenestra is almost not exposed in lateral view (1). (-) in taxa lacking both an upper and lower temporal fenestra which frame a postorbital bar.**

Modified from Ezcurra (2016) Ch. 126.

In early neodiapsids such as the EAZ younginiform or *Claudiosaurus* (Carroll 1981), the postorbital bar is positioned relatively low, approximately level with the dorsoventral midheight of the orbit in lateral view (State 0). In Archosauriformes (e.g., *Euparkeria,* Sookias et al. 2020) and *Tanystropheus* within this sampling, the postorbital bar is located approximately level with the dorsal rim of the orbit (State 1).

UNAMBIGUOUS: *Tanystropheus hydroides* (0 to 1), *Mesosuchus* + Archosauriformes (0 to 1).

**Ch. 124. Postorbital, supratemporal fossa: absent (0), present, supratemporal fossa extends onto posterior surface of postorbital. Inapplicable (-) in taxa lacking a supratemporal fossa of the skull (Ch. 6).**

Modified from Nesbitt (2011, Ch. 144), Ezcurra (2016, Ch. 129).

The supratemporal fossa, when present, extends onto the posterior surface of the postorbital (State 1) in crocopod archosauromorphs (e.g., *Azendohsaurus* UA-7-20-99-653) in this sampling.

DELTRAN: *Biseridens qilianicus* (0 to 1), Crocopoda (0 to 1). ACCTRAN: *Biseridens qilianicus* (0 to 1), Allokotosauria + *Mesosuchus* + Archosauriformes (0 to 1).

**Ch. 125. Postorbital, dorsal and lateral surfaces: form smooth curve (or dorsal surface absent, postorbital not participating in skull roof) (0), sharply divided (meeting at edge) (1).**

Modified from Brocklehurst et al. (2022, Ch. 72 ).

**Ch. 126. Postorbital and jugal, medial orbital process ( deep, dorsoventrally tall medial flange): absent (0), present (1).**

Modified from Brocklehurst et al. (2022, Ch. 310).

**Ch. 127. Postorbital, contact with parietal: absent (0), present (1).**

Modified from Brocklehurst et al. (2022, Ch. 73).

**Ch. 128. Postorbital, posterior process, contact with supratemporal: absent (0), present (1).**

Modified from Brocklehurst et al. (2022, Ch. 69).

**Ch. 129. Postorbital, anteroposterior length of posterior process: short (0), long, extending more than half of temporal length (1).**

Modified from Brocklehurst et al. (2022, Ch. 70).

**Ch. 130. Postorbital, posterior process, transverse ( dorsoventral) width at midlength: broad, at least 25% of dorsoventral height of temporal region (0), narrow, no more than 20% of dorsoventral height of temporal region (1).**

Modified from Brocklehurst et al. (2022, Ch. 71).

**Ch. 131. Postorbital, posterior process, contact with squamosal: present, contacting squamosal posteriorly (0), absent, no contact between posterior portion of postorbital and anterior portion of squamosal (1).**

Modified from Pritchard et al. (2021. Ch. 339).

The posterior process of the postorbital does not make contact with the squamosal (State 1) in edaphosaurid synapsids (e.g., *Gordodon,* NMMNH P-70796) or weigeltisaurid neodiapsids (e.g., *Coelurosauravus*, Buffa et al. 2021) within this sampling.

UNAMBIGUOUS: Edaphosauridae (0 to 1), *Candelaria barbouri* (0 to 1), Weigeltisauridae (0 to 1).

**Ch. 132. Temporal ( = Postorbital) region, height of ventral temporal ( zygomatic) bar ( complete or partially complete): tall, occupying more than 20% of skull height (0), narrow, occupying less than 20% of postorbital skull height (1) inapplicable, ventral temporal bar absent (-).**

Modified from Brocklehurst et al. (2022, Ch. 74).

**Ch. 133. Jugal, free posterior margin: absent (0), present (1).**

Modified from Brocklehurst et al. (2022, Ch. 76).

**Ch. 134. Jugal, free concave ventral margin: absent (0), present (1).**

Modified from Brocklehurst et al. (2022, Ch. 77).

**Ch. 135. Jugal, contribution to ventral margin of the skull: present (0), absent (1).**

Modified from Brocklehurst et al. (2022, Ch. 78).

**Ch. 136. Jugal, if contribution to ventral margin of skull is absent: jugal excluded from ventral margin by maxilla-quadratojugal contact achieved by a long posterior extension of the maxilla (0), maxilla-quadratojugal contact achieved by long anterior extension of the quadratojugal (1).**

Modified from Brocklehurst et al. (2022, Ch. 79).

**Ch. 137. Jugal, suborbital ramus: does not extend to anterior orbital rim (0), extends at least to level of anterior orbital rim (1).**

Modified from Brocklehurst et al. (2022, Ch. 80).

**Ch. 138. Jugal, anteroposterior thickness of dorsal ramus ( forming postorbital bar): broad, temporal fenestra only weakly emarginates the jugal (0), narrow (1).**

Modified from Brocklehurst et al. (2022, Ch. 81).

**Ch. 139. Jugal, posterior ramus length: long, extending to the posterior end of temporal fenestra (or past temporal midlength in taxa that lack a temporal fenestra) (0), short, extending to, or less than, temporal midlength (1).**

Modified from Brocklehurst et al. (2022, Ch. 82).

**Ch. 140. Jugal, tuberous ornamentation on lateral surface: absent (0), present (1).**

Modified from Brocklehurst et al. (2022, Ch. 83).

**Ch. 141. Jugal, ventral contact with squamosal perforated by small elongate fenestra: absent (0), present, upper margin of fenestra enclosed by anteroventral extension of the squamosal (1).**

Modified from Brocklehurst et al. (2022, Ch. 84).

**Ch. 142. Jugal, if ectopterygoid absent, alary process: alary process absent (0), present, no higher than the midpoint of the suborbital process of the jugal and distinct from the orbital margin (1), present, positioned on the medial surface of the jugal and flush with orbital margin (2).**

Reworked from Brocklehurst et al. (2022, Ch. 150).

Rationale: The original framing of Ch. 150 concerned both the presence of the ectopterygoid and the morphology of the jugal, both features that vary independently in taxa within this matrix. We therefore split Ch. 150 accordingly.

**Ch. 143. Quadratojugal: present (0), absent as a discrete ossification (1).**

Modified from Gauthier (1988b Ch. 12)

The quadratojugal is absent as a distinct ossification (State 1) in lepidosauromorphs (e.g., *Bellairsia,* Tałanda et al. 2022) and several recumbirostrans (e.g., *Brachydectes*, KUVP 49541) within this sampling.

DELTRAN: *Brachydectes newberryi* (0 to 1), *Dvellecanus carrolli* (0 to 1), *Rhynchonkos stovalli* (0 to 1), Lepidosauria (0 to 1). ACCTRAN: Chthonosauria (0 to 1), Rhynchonkidae (0 to 1), *Bellairsia*+ Squamata (0 to 1).

**Ch. 144. Quadratojugal, contact with posterodorsal margin of jugal: absent (0), present (1).**

Modified from Brocklehurst et al. (2022, Ch. 86).

**Ch. 145. Quadratojugal, contact with posteroventral margin of jugal: present (0), absent (1).**

Modified from Brocklehurst et al. (2022, Ch. 85).

**Ch. 146. Quadratojugal, shape of anterior margin of anteroventral process: square- tipped (0), acuminate (tapering to a point) (1).**

Modified from Brocklehurst et al. (2022, Ch. 87).

**Ch. 147. Quadratojugal, lateral surface: smooth or mildly convex (0), lateral surface possesses at least one laterally-directed spine-like process of bone (1). (-) in taxa lacking a quadratojugal (Ch. 143).**

Modified from Modesto et al. (2002 Ch. 14), Cisneros (2008,Ch. 13).

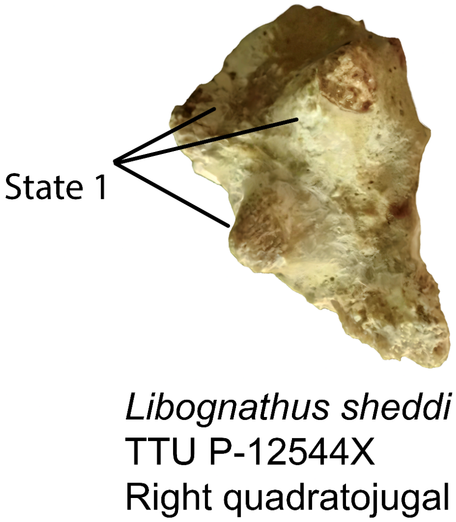


**Fig. S25.** Right quadratojugal of *Libognathus sheddi* (TTU P-12544X) which bears three quadratojugal spines (Ch. 147, State 1).

The quadratojugal of derived procolophonoids (e.g., *Hypsognathus*, YPM 55831) possess spine-like projections on the lateral surface that may have been covered in a keratinous sheath (State 1).

UNAMBIGUOUS: Leptopleuroninae + *Procolophon* (0 to 1).

**Ch. 148. Quadratojugal, ventral process or flange: absent (0), present (1). (-) in taxa lacking a quadratojugal (Ch. 143)**

Modified from Van den Brandt (2020, Ch.44).

The quadratojugal of pareiasaurs (e.g., *Deltavjatia,* Tsuji 2013) possesses a ventral flange which projects far ventral to both the maxillary and dentary tooth rows in lateral view, sometimes covering the jaw articulation (State 1).

UNAMBIGUOUS: Pareiasauria + *Emerolete*r (0 to 1), Leptopleuroninae (0 to 1).

**Ch. 149. Quadratojugal, dorsal process, dorsoventral height: short or absent, equal to or less than .5 times the total height of skull (0), tall, approximately .7 the dorsoventral height of skull in temporal region (1). (-) in taxa lacking a quadratojugal (Ch. 143).**

This study.

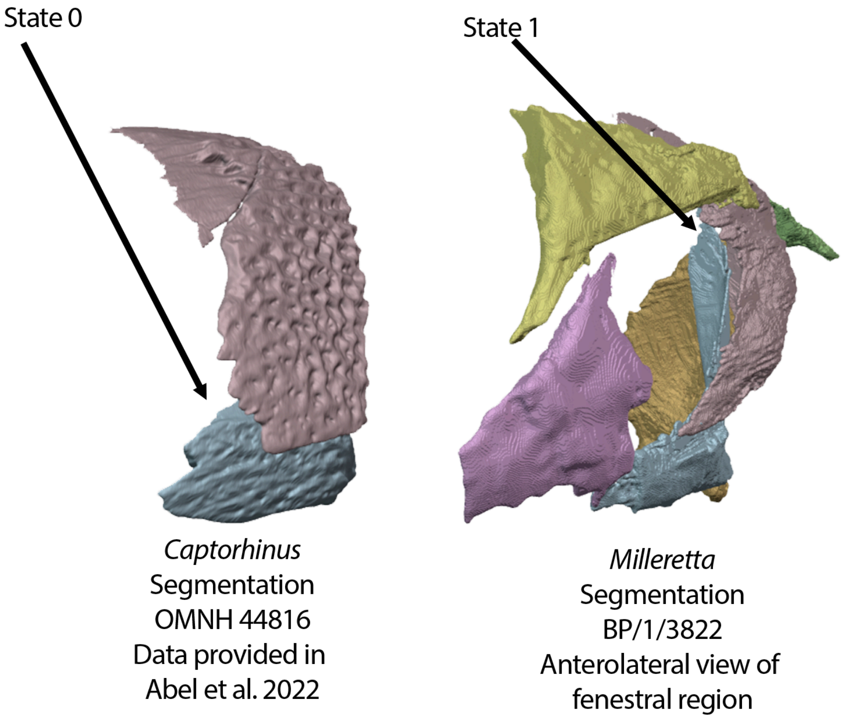


**Fig. S26. Left,** left squamosal and quadratojugal of *Captorhinus* (OMNH 44816) in lateral view demonstrating the plesiomorphic condition for Amniota; a dorsoventrally low quadratojugal (Ch. 149, State 0). Right, left temporal region of *Milleretta* (BP/1/3822, this study) demonstrating the derived condition of a dorsoventrally expanded dorsal process of the quadratojugal (Ch. 149, State 1).

The quadratojugal of most early tetrapods is short, lacking a dorsal development in lateral view (State 0, e.g., *Protorothyris*, MCZ 1532). Millerettids (e.g., *Milleretta,* BP/1/3822) and *Eunotosaurus* (CM777) possess a dorsoventrally tall quadratojugal that extends the entire dorsoventral height of the temporal region (State 1).

DELTRAN: *Hapsidopareion* (0 to 1). ACCTRAN: *Llistrofus* + Chthonosauria (0 to 1),Millerettidae (0 to 1).

**Ch. 150. Postorbital cheek: straight and unexpanded laterally relative to lateral margin of maxilla (0), moderately expanded laterally relative to lateral margin of maxilla (1).**

Modified from Brocklehurst et al. (2022, Ch. 88).

**Ch. 151. Squamosal, contacts posterodorsal margin of jugal: present (0), absent (1).**

Modified from Brocklehurst et al. (2022, Ch. 89).

**Ch. 152. Squamosal, contacts posteroventral margin of jugal: absent (0), present (1).**

Modified from Brocklehurst et al. (2022, Ch. 90).

**Ch. 153. Squamosal, anteroventral process: absent (0), present, anteroventral process extends from body of squamosal in lateral view (1).**

Modified from Simões et al. (2022, Ch. 57).

**Ch. 154. Squamosal, supratemporal fossa: absent (0), present, supratemporal fossa extends onto squamosal (1). Inapplicable (-) in taxa lacking a supratemporal fossa (Ch. 6).**

Modified from Nesbitt (2011, Ch. 55), Ezcurra (2016 Ch. 149).

The supratemporal fossa, when present, extends onto the anterior margin of the squamosal in allokotosaurian archosauromorphs within this sampling (e.g., *Trilophosaurus,* TMM 31025-140).

DELTRAN: Therapsida (0 to 1), *Trilophosaurus buettneri* (0 to 1), *Azendohsaurus madagaskarensis* (0 to 1). ACCTRAN: *Mesosuchus browni* (1 to 0), Allokotosauria + *Mesosuchus* (0 to 1).

**Ch. 155. Squamosal, anteroposterior breadth of post-fenestral portion on lateral surface of skull: narrow (0), broad, just over half the breadth of the temporal region, restricting the posterior extent of the temporal fenestra (1).**

Modified from Brocklehurst et al. (2022, Ch. 91).

**Ch. 156. Squamosal, occipital shelf ( flange): mediolaterally broad, contributes to occipital surface of skull, obscuring much of the dorsal ramus of the quadrate in occipital view (0), absent or mediolaterally very narrow dorsal ramus of quadrate exposed in occipital view (1).**

Modified from Brocklehurst et al. (2022, Ch. 92).

**Ch. 157. Squamosal, dorsal process: absent (0), present, squamosal possesses a discrete dorsal process which braces the parietal in lateral view (1).**

Modified from Simões et al. (2022, Ch. 58).

**Ch. 158. Squamosal, posterodorsal process: absent (0), present, a posterodorsal process of SQ roofs the quadrate in lateral view (1).**

Modified from Brocklehurst et al. (2022, Ch. 93).

**Ch. 159. Squamosal, contact with postparietal: absent (0), present (1). Inapplicable (-) in taxa lacking a postparietal (Ch. 208).**

Modified from Anderson et al. (2008, Ch. 56), Huttenlocker et al. (2013, Ch. 56).

The squamosal of some recumbirostrans (e.g., *Rhynchonkos*, FM-UR 1039) contacts the postparietals ventrally (State 1).

UNAMBIGUOUS: *Rhynchonkos stovalli* (0 to 1), *Reiszorhinus olson*i (0 to 1). DELTRAN: *Brachydectes newberryi* (0 to 1). ACCTRAN: Chthonosauria (0 to 1).

**Ch. 160. Squamosal, superficial contact with tabular: absent (0), present (1). Inapplicable (-) in taxa lacking a tabular (Ch. 189)**

Modified from Anderson et al. (2008, Ch. 10), Huttenlocker et al. (2013,Ch. 10).

The squamosal of early amniotes contacts the tabular dorsomedially (e.g., *Ophiacodon,* Romer and Price 1940). In neoreptiles, this contact is separated by the supratemporal or parietal in posterior view (e.g., *Youngina*, BP/1/2871).

UNAMBIGUOUS: *Greererpeton burkemoran*i (0 to 1), *Acheloma* + *Cacops* (0 to 1), Recumbirostra + Captorhinidae + Diadectomorpha + Protorothyrididae + Amniota (0 to 1), Parapleurota (0 to 1). DELTRAN: *Varanops* + *Varanodon* (1 to 0). Ankyramorpha + *Erpetonyx* (1 to 0). . ACCTRAN: Caseasauria (1 to 0), Ankyramorpha + *Erpetonyx* (1 to 0).

**Ch. 161. Squamosal, contribution to the posttemporal fenestra: absent (0), present (1).**

Modified from Brocklehurst et al. (2022, Ch. 94).

**Ch. 162. Squamosal, pterygoid ramus or contact: absent (0), present, squamosal contacts quadrate ramus of pterygoid (1).**

This study.

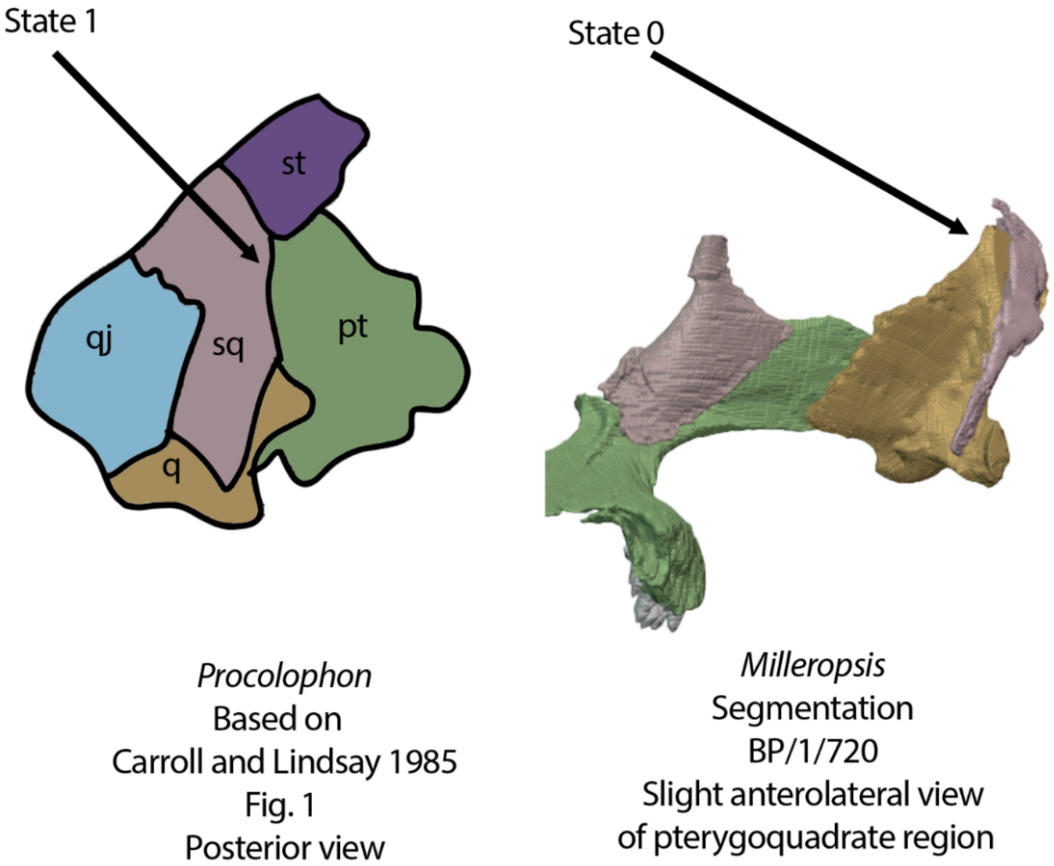


**Fig. S27.** Left, diagram of the left palatoquadrate region of *Procolophon* (based on Carroll and Lindsay 1985) demonstrating the contact between the squamosal and the quadrate ramus of the pterygoid (Ch. 162, State 0). Right, left palatoquadrate of *Milleropsis pricei* (BP/1/720, this study) demonstrating (Ch. 162, State 1) or the absence of squamosal-pterygoid contact.

The squamosal of early tetrapods (e.g., *Edops,* MCZ 1378) and early amniotes such as synapsids (Romer and Price 1940), protorothyridids (e.g., *Protorothyris,* MCZ 1532), and ankyramorphan neoreptiles (e.g., *Procolophon*, Carroll and Linday 1985) possesses a pterygoid ramus which contacts the quadrate ramus of the pterygoid (State 0). In contrast, the squamosal of parapleurotan (e.g., *Milleretta* BP/1/3822 or *Prolacerta* BP/1/5880) reptiles lacks a pterygoid ramus and thus does not contact the pterygoid ventrally (State 1).

DELTRAN: *Varanops* + *Varanodon* (1 to 0), Araeoscelidia (1 to 0), Pareiasauria + *Emeroleter* (1 to 0), Reptile total-group (0 to 1). ACCTRAN: *Huskerpeton englehorni* (0 to 1), Gymnarthridae + *Nannaroter* +

Rhynchonkidae + *Huskerpeton* (1 to 0), Eupelycosauria + Varanopidae (0 to 1), Pareiasauromorpha (1 to 0) Ankyramorpha + *Erpetonyx* (0 to 1).

**Ch. 163. Squamosal, lateral (=descending) process/flange, dorsoventral height: tall, at least 50% the dorsoventral height of the temporal region (0) short, less than 50% the dorsoventral height of the temporal region (1).**

Modified from Pritchard (2018, Ch. 80).

The squamosal of many archosauromorph reptiles (e.g., *Tanystropheus*, PIMUZ T 2790) possesses little to no ventral development in lateral view, such that the squamosal is no more than 50% the height of the temporal region in lateral view (State 1).

DELTRAN: *Huehuecuetzpalli mixtecus* (0 to 1), *Mesosuchus browni* (1 to 0), Allokotosauria + *Mesosuchus* + Archosauriformes (1 to 0), Archosauromorpha (0 to 1). ACCTRAN: *Bellairsia*+ Squamata (0 to 1), *Mesosuchus browni* (1 to 0), Archosauriformes (1 to 0), Tanystropheidae + Crocopoda (0 to 1).

**Ch. 164. Squamosal, lateral (=descending) process/flange, anteroposterior exposure: anteroposteriorly broad, covering quadrate entirely in lateral view (0), anteroposteriorly slender, partially exposing dorsal shaft of quadrate in lateral view (1).**

Modified from Pritchard (2018, Ch. 60).

The descending process of the squamosal in parapleurotan reptiles (e.g., *Milleropsis* BP/1/720 or *Youngina* BP/1/2871) is anteroposteriorly slender, such that the quadrate is exposed in lateral view (State 1).

UNAMBIGUOUS: Parapleurota (0 to 1). DELTRAN: *Claudiosaurus germaini* (1 to 0), Weigeltisauridae (1 to 0).

**Ch. 165. Squamosal/Quadratojugal, posterolateral portion, emargination: absent (0) present, posterolateral portion of SQ/QJ emarginated or concave (1) posterolateral portion of SQ possesses a groove or notch (2). ORDERED.**

This study.

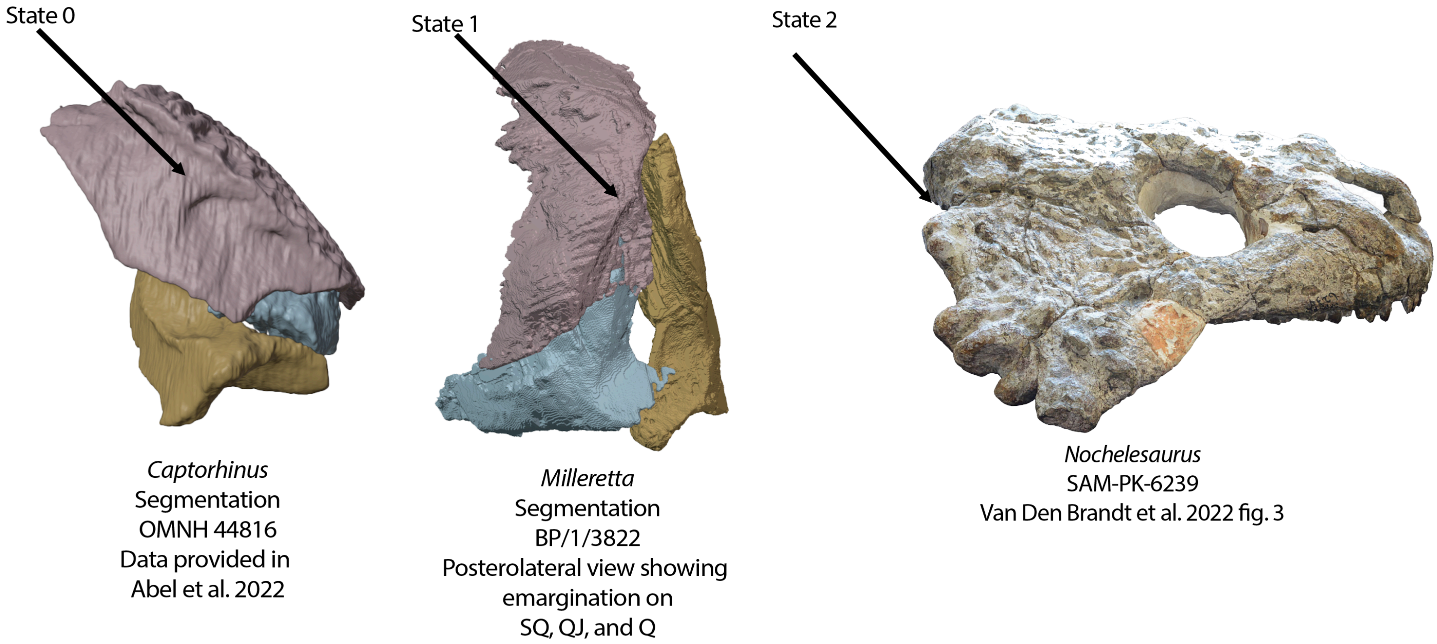


**Fig. S28.** Squamosal, quadratojugal, and quadrate of *Captorhinus* (OMNH 44816), *Milleretta* (BP/1/3822, this study) in occipital views and right, skull of *Nochelesaurus* (SAM-PK-6239) in lateral view. *Captorhinus* lacks a tympanic emargination (Ch. 166, State 0), *Milleretta* bears a tympanic emargination shared between the squamosal, quadratojugal, and quadrate (Ch. 165, State 1), and *Nochelesaurus* bears a notch in the squamosal that has been homologized with the tympanic recess of other procolophonians (Ch. 165, State 2).

The squamosals and/or quadratojugals of procolophonian ankyramorphs (e.g., *Sauropareion*, SAM-PK-K11192), millerettids (e.g., *Milleretta* BP/1/3822), early neodiapsids (*Rautiania*, PIN 5130/62), archosauromorphs (e.g., *Tanystropheus,* PIMUZ T 2790), rhynchocephalians (e.g., *Clevosaurus*, NHMUK PV R36832) possesses a posterolateral emargination likely supporting or contributing to a tympanum (State 0). Pareiasaurs possess a distinct groove marking their 'otic notch' (Van den Brandt et al. 2021).

UNAMBIGUOUS: *Greererpeton burkemorani* (1 to 0), Pareiasauria + *Emeroleter* (1 to 2) Ankyramorpha (0 to 1), Parapleurota (0 to 1). DELTRAN: *Marmoretta oxoniensis* (0 to 1), Squamata (1 to 0), Diadectomorpha (1 to 0). ACCTRAN: Recumbirostra + Diadectomorpha + Protorothyrididae + Amniota (1 to 0).

**Ch. 166. Squamosal/Quadratojugal, posterolateral portion, emargination when present: open ventrally (0), closed ventrally by tympanic crests (1).**

This study.

The posterolateral emargination or tympanum on the squamosal and quadratojugal is closed dorsally by tympanic crests but remains open ventrally (State 0) in most early reptiles possessing a squamosal or quadratojugal emargination (e.g., *Milleretta*. BP/1/3822). In contrast, the posterolateral emargination is closed ventrally by tympanic crests in procolophonian ankyramorphs (e.g., *Macroleter*, Tsuji 2006) in the sampling here (State 1).

DELTRAN: Ankyramorpha (0 to 1). ACCTRAN: Ankyramorpha + *Erpetonyx* (0 to 1).

**Ch. 167. Preopercular: present (0), absent (1).**

Derived from Clack et al. 2012, Pardo et al. 2019 Ch. 272

The preopercular bone is present (State 0) in the outgroup within this matrix, *Whatcheeria* FM PR 1634.

UNAMBIGUOUS: All taxa more derived than *Whatcheeria* (0 to 1).

**Ch. 168. Skull, posterior margin, if emargination(s) present, anteroposterior length: greater than 70% postorbital length of skull (0), less than 70% postorbital length of the skull (1). Inapplicable (-) in taxa lacking a posterolateral emargination of the skull e.g., 'otic notch' 'spiracular notch' 'tympanum'.**

This study.

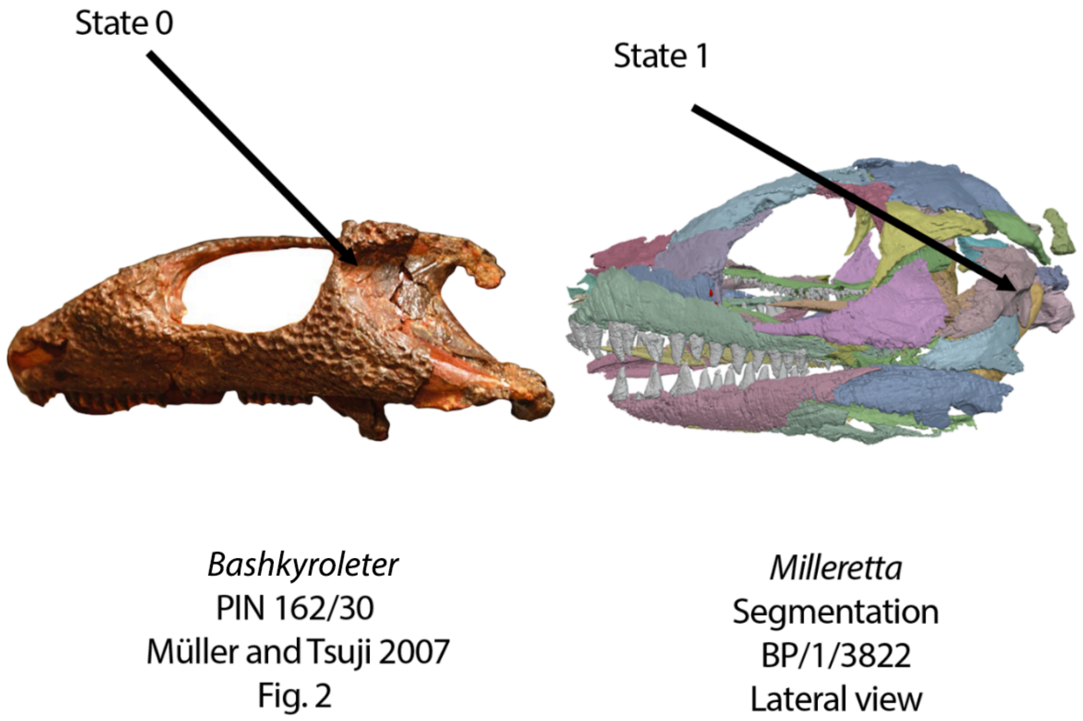


**Fig. S29.** Left, skull of *Bashkyroleter* (PIN 162/30) and right, *Milleretta* (BP/1/3822, this study) in lateral view. *Bashkyroleter* bears a anteroposteriorly broad tympanic emargination that is broadly exposed in lateral view, nearly reaching the level of the posterior orbital margin (Ch. 168, State 0) whereas the tympanum of *Milleretta* is restricted to the posterolateral surface of the skull (Ch. 168, State 1).

The ‘tympanum’, ‘spiracular notch’, ‘otic notch’ or posterior skull emarginations of select temnospondyls (e.g., *Cacops,* OMNH 53073) and early pareiasauromorphs (e.g., *Emeroleter* PIN 2212/14) are anteroposteriorly broad, strongly emarginating the skull anteroposteriorly in lateral view (State 1).

UNAMBIGUOUS: *Acheloma* + *Cacops* (1 to 0). DELTRAN: *Emeroleter levis* (1 to 0), *Macroleter poezicus* (1 to 0), *Seymouria* spp. (1 to 0), *Archeria crassidiscus* (1 to 0). ACCTRAN: Pareiasauria + *Emeroleter* (0 to 1), Pareiasauromorpha + *Australothyris* (1 to 0).

**Ch. 169. Skull, posterior margin, if emargination(s) present, dorsoventral height: greater than 3/5 the height of temporal region (0), less than or equal to 3/5 of dorsoventral height of occipital region (1). Inapplicable (-) in taxa lacking a posterolateral emargination of the skull 'otic notch' 'spiracular notch' 'tympanum'.**

This study.

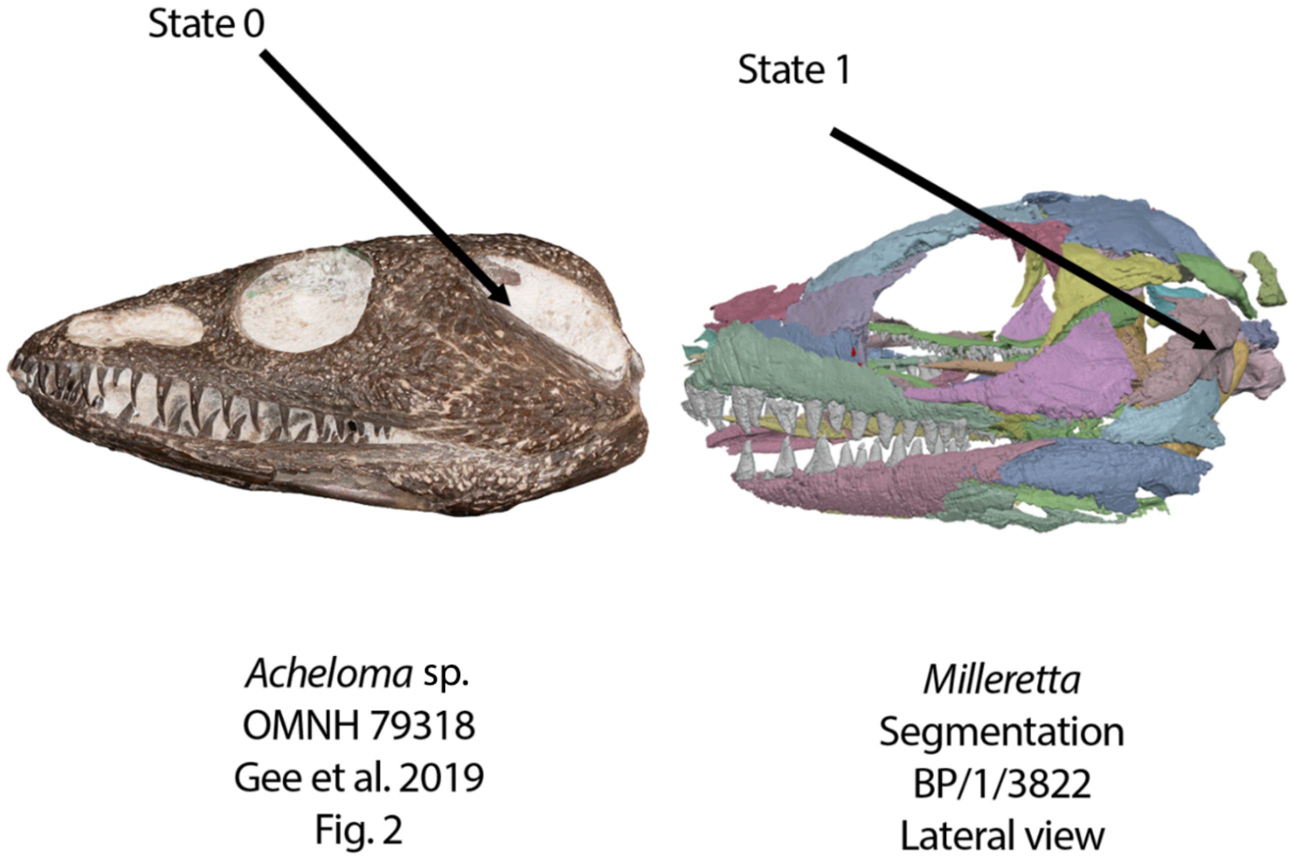


**Fig. S30.** Left, skull of *Acheloma* (OMNH 79318, modified from Gee et al. 2019b) and right, *Milleretta* (BP/1/3822, this study) in lateral view. *Acheloma* bears a dorsoventrally tall tympanic emargination that is nearly as tall as the entire postorbital skull (Ch. 169, State 0) whereas the tympanum of *Milleretta* is dorsoventrally short (Ch. 169, State 1).

The ‘tympanum’, ‘spiracular notch’, ‘otic notch’ or posterior skull emarginations of many early tetrapods (e.g., *Chroniosaurus*, PIN 3585/124) and procolophonians (e.g., *Procolophon* NHMUK PV R. 4087) are dorsoventrally tall, more than 3/5$^{th}$ the height of the occipital region in lateral or occipital view (State 1).

DELTRAN: Pareiasauria (0 to 1), Ankyramorpha (1 to 0), *Paliguana whitei* (1 to 0), *Orobates pabst*i (1 to 0), *Lanthanosuchus* + *Chroniosaurus* (1 to 0). ACCTRAN: Pareiasauria + *Emeroleter* (0 to 1), *Paliguana whitei* (1 to 0), Ankyramorpha + *Erpetonyx* (1 to 0).

**Ch. 170. Parietal, fusion with contralateral parietal: absent (0), present (1).**

Modified from Benton (1985 Ch. Y1), Gauthier (1986), Pritchard et al. (2015 Ch. 14), Ezcurra (2016 Ch. 112).

The fusion of the parietals into a single, median element (State 1) is present in several lepidosauromorphs (e.g., *Planocephalosaurus*, AUP 11070) within this taxonomic sampling.

DELTRAN: *Marmoretta oxoniensis* (0 to 1), *Planocephalosaurus robinsonae* (0 to 1), *Huehuecuetzpalli mixtecus* (0 to 1), *Bellairsia gracilis* (0 to 1), *Mesosuchus browni* (0 to 1). ACCTRAN: *Megachirella wachtleri* (1 to 0), *Clevosaurus hudson*i (1 to 0), *Eichstaettisaurus schroederi* (1 to 0), *Mesosuchus browni* (0 to 1), Lepidosauromorpha (0 to 1)

**Ch. 171.** Parietal, anterolateral process: absent (0), present, anterolateral process partially separates postfrontal and postorbital (or intertemporal, if present) (1).

This study.

Noted as an autapomorphy of Millerettidae by Romer (1956), the anterolateral process of the parietal is a subtriangular process of bone that partially separates contact between the postorbital and postfrontal in dorsal view (State 1). It is widespread among stem and crown amniotes, with a similar feature observed in some captorhinids (pers. obv. *Euconcordia*), ophiacodontids (e.g., *Varanosaurus*, BSPHM 190 XV 20), and mesosaurids (e.g., *Mesosaurus*, MCZ 4031a).

ACCTRAN: Millerettidae (0 to 1).

**Ch. 172. Parietal, free lateral margin: absent (0), present (1).**

Modified from Brocklehurst et al. (2022, Ch. 95).

**Ch. 173. Parietal contribution to the orbit: absent (0), present (1).**

Modified from Cisneros (2008, Ch. 10).

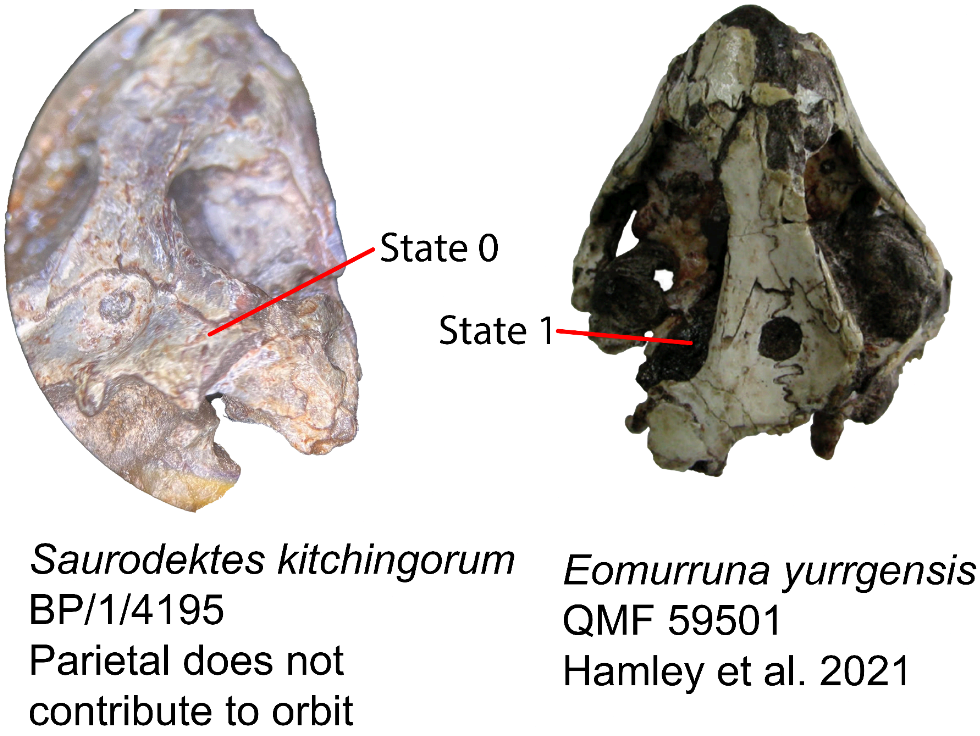


**Fig. S31.** Skulls of *Saurodektes* (BP/1/4195) and *Eomurrunna* (QMF 59501, modified from Hamley et al. 2021) in dorsal views. The parietal of *Saurodektes* does not contribute to the orbital margin (Ch. 173, State 0) whereas the parietal of *Eomurruna* does (Ch. 173, State 0).

The parietal contributes to the orbital margin (State 1) in taxa that possess enlarged orbitotemporal temporal fenestra or lose a discrete ossification of the postfrontal and/or postorbital, which includes the recumbirostran *Brachydectes* (e.g., KUVP 49541) and procolophonids within this sampling (e.g., *Hypsognathus*, YPM 55831).

UNAMBIGUOUS: *Brachydectes newberryi* (0 to 1), Procolophonidae (0 to 1).

**Ch. 174. Parietal, anteroposterior diameter of pineal foramen: large, more than 25% of mid-parietal length (0), small, less than 25% of mid-parietal length (1), absent (2). ORDERED.**

Modified from Brocklehurst et al. (2022, Ch. 96).

**Ch. 175. Parietal, pineal foramen location: anteriorly on parietal midline suture, such that the anterior rim of the foramen lies within the anterior third of the midline suture (0), midlength on parietal midline suture, such that the anterior and posterior rims of the foramen both lie within the middle third of the midline suture (1), posteriorly on parietal midline suture, such that the posterior rim of the foramen lies within the posterior third of the midline suture (2). ORDERED.**

Modified from Brocklehurst et al. (2022, Ch. 97).

**Ch. 176. Parietal, skull table, morphology of pineal region: pineal foramen flush with skull roof (0), surrounded by raised area forming a pineal ridge or boss (1), pineal foramen lies in a fossa or depression (2).**

Reworked from Brocklehurst et al. (2022, Ch. 311).

Rationale: Added a third state concerning the morphology present in many stem reptiles, such as owenettids (Reisz and Scott 2002).

**Ch. 177. Parietal, dorsoventrally narrow flange extending ventrally from the lateral margin of the parietal (ventrolateral flange): absent (0), present (2).**

Modified from Brocklehurst et al. (2022, Ch. 98).

**Ch. 178. Parietals, ventrolateral (posterolateral) processes, height: dorsoventrally low, considerably lower in height than the supraoccipital (0), posterolateral process of the parietal dorsoventrally deep, being plate-like in occipital view and subequal to the height of the supraoccipital (1). Inapplicable in taxa lacking a ventrolateral flange of the parietals (Ch. 178)**

Modified from Ezcurra (2016, Ch. 168)

The ventrolateral processes of the parietals, a feature present in many neodiapsids (see Ch. 173) are dorsoventrally taller relative to the height of the supraoccipital (State 1) in the crocopod archosauromorphs within this sampling (e.g., *Azendohsaurus* UA-7-20-99-653).

DELTRAN: Tanystropheidae + Crocopoda (0 to 1). ACCTRAN: Crocopoda (0 to 1).

**Ch. 179. Parietals, lateral frill, horns, or spikes: absent (0), present (1).**

Modified from Simões (2018, Ch.77), Pritchard et al. (2021 Ch. 59).

The lateral margins of the parietals, squamosals, and quadratojugals of weigeltisaurid neodiapsids (e.g., *Rautiania*, PIN 5130/62) possess a lateral development bearing raised tubercles, ‘horns’ or ‘spikes’ which form a posteriorly-oriented frill in lateral view (State 1).

UNAMBIGUOUS: Weigeltisauridae (0 to 1).

**Ch. 180. Parietal, lateral margin: restricted to skull table in lateral view (0), contributes to lateral surface of the skull dorsally (1) (-) in taxa lacking posterior circumorbital elements.**

This study, based on the observations of Watson (1957).

The lateral margin of the parietal in early reptiles (e.g., *Captorhinus* OMNH 44816, *Saurodektes* BP/1/4195) is restricted to the dorsal surface of the skull table (State 0). In millerettids and the putative stem-turtle *Eunotosaurus* (e.g., CM777), the parietal possesses a ventral deflection which contributes broadly to the lateral surface of the skull (State 1), interpreted as a secondary closure of the upper temporal fenestra by several studies (Watson 1957, Bever et al. 2015).

DELTRAN: Millerettidae (0 to 1), *Secodontosaurus obtusidens* (0 to 1). ACCTRAN: *Secodontosaurus obtusidens* (0 to 1).

**Ch. 181. Supratemporal: present (0), absent(1).**

Modified from Brocklehurst et al. (2022, Ch. 100).

**Ch. 182. Supratemporal, restricted to skull table (excluding emargination of spiracular notch): present (0), absent, supratemporal has occipital exposure (1).**

Modified from Pardo et al. (2017, Ch. 6).

**Ch. 183. Supratemporal, size: broad, mediolateral width is greater than 30% of width of posterior skull table from lateral margin of supratemporal to midline (0), slender, in parietal or parietal/squamosal trough, mediolateral width approximately 15% (1).**

Modified from Brocklehurst et al. (2022, Ch. 100).

**Ch. 184. Supratemporal, posteroventral extent: approximately equal to or less than that of the tabular (0), greatly exceeds that of the tabular (1).**

Modified from Brocklehurst et al. (2022, Ch. 101).

**Ch. 185. Supratemporal, posteroventral portion, overhangs suspensorium: absent (0), present, posteroventral portion of supratemporal emarginated such that it roofs or overhangs skull posteriorly (1). Inapplicable in taxa lacking a supratemporal (Ch. 181).**

Modified from Spindler et al. (2018, Ch. 54).

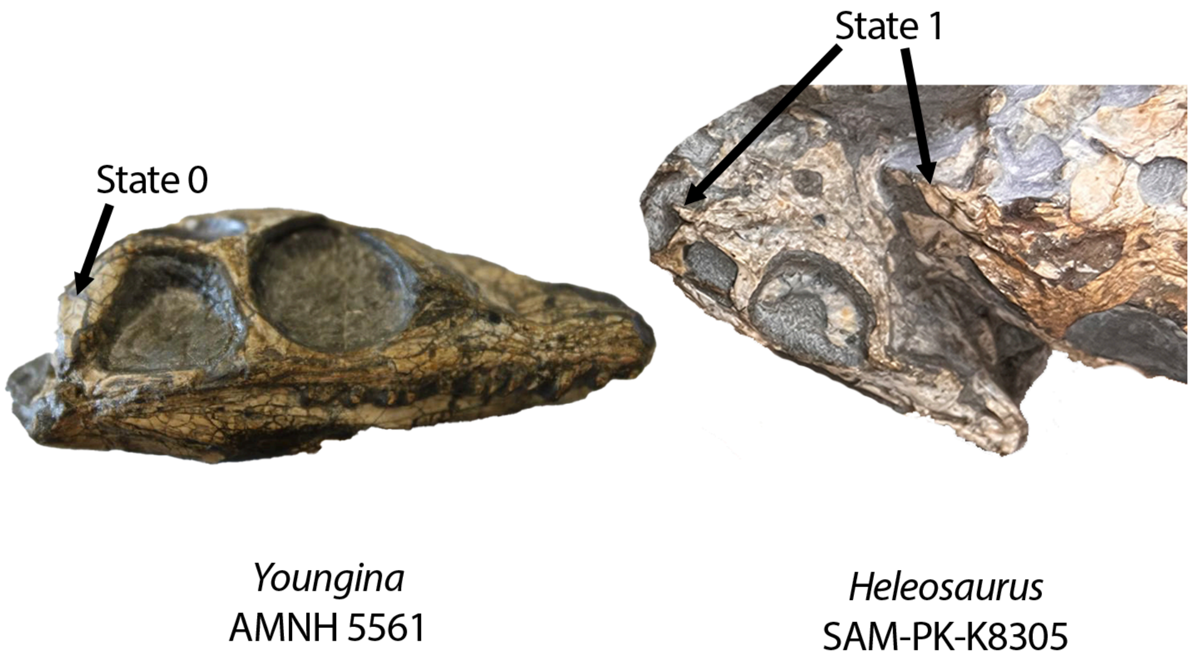


**Fig. S32.** Left, skull of *Youngina* (AMNH 5561) in right lateral view which bears a supratemporal that does not posteriorly overhang the skull (Ch. 185, State 0). Right, skull of *Heleosaurus* (SAM-PK-K8305) demonstrating (Ch. 185, State 1), or the presence of a supratemporal that overhangs the suspensorium.

A posterior projection of the supratemporal, interpreted to form the dorsal surface of the tympanum by (Spindler et al. 2018) is present (State 1) in *Ascendonanus* (Spindler et al. 2018) as well as the varanopid *Heleosaurus* (SAM-PK-K8305). The supratemporal of millerettids (e.g., *Milleretta*, BP/1/3822) and procolophonians (e.g., *Sauropareion*, SAM-PK-11192) also possess a posterior projection (State 1).

UNAMBIGUOUS: *Greererpeton burkemorani* (1 to 0), *Edops craigi* (1 to 0), Bradysauridae (1 to 0). DELTRAN: *Cotylorhynchus* (0 to 1), Pareiasauromorpha + *Australothyris*(0 to 1), Procolophonia (0 to 1), *Ascendonanus nestleri* (0 to 1), Mesenosaurinae (0 to 1), Millerettidae (0 to 1). ACCTRAN: Caseidae (1 to 0), Caseasauria (0 to 1) *Australothyris* (1 to 0), Ankyramorpha (0 to 1).

**Ch. 186. Supratemporal, posteroventral portion, if overhangs suspensorium: margin is smoothed, featureless (0), distinct dorsal crest frames fossa or emargination (1) . Inapplicable (-) in taxa lacking a supratemporal (Ch. 181).**

Modified from Spindler et al. (2018, Ch. 54).

The medial tympanic crest extends onto the ventral surface of the posteroventral development of the supratemporal in procolophonians (e.g., *Emeroleter* PIN 2212/14) within this sampling.

DELTRAN: *Dendrerpeton helogenes* (0 to 1), *Acheloma* + *Cacops* (0 to 1), Ankyramorpha (0 to 1). ACCTRAN: *Balanerpeton woodi* (1 to 0), Ankyramorpha + *Erpetonyx* (0 to 1), Temnospondyli (0 to 1).

**Ch. 187. Intertemporal: present (0), absent (1).**

Modified from Brocklehurst et al. (2022, Ch. 102).

**Ch. 188. Tabular: present (0), absent (1).**

Modified from Brocklehurst et al. (2022, Ch. 103).

**Ch. 189. Tabular, if present, contact with parietal: present (0), absent (1). Inapplicable (-) in taxa lacking a tabular (Ch. 188).**

Modified from Anderson et al. (2008, Ch. 52), Huttenlocker et al. (2013 Ch. 52), Pardo et al. (2017, Ch. 46).

The tabular does not contact the parietal in early tetrapodomorphs due to a combination of the large size of the postparietals (which are generally the same width as the parietals) and a posteromedial extension of the supratemporal (e.g., *Dendrerpeton* NSM 987 GF 99.1).

UNAMBIGUOUS: Temnospondyli (0 to 1), Temnospondyli + Reptiliomorpha (1 to 0). DELTRAN: *Cotylorhynchus romeri* (0 to 1). ACCTRAN: *Cotylorhynchus* + *Angelosaurus* (0 to 1).

**Ch. 190. Tabular, morphology: broad sub-rectangular sheet located dorsal to the fenestra (0), small and wedge-shaped, located dorsal to the post-temporal fenestra (1), long and narrow, displaced laterally, located dorsolateral to post- temporal fenestra (2), large L-shaped bone enclosing post-temporal fenestra dorsally and laterally (3), broad sheet with broad exposure on the skull roof (4).**

Modified from Brocklehurst et al. (2022, Ch. 104).

**Ch. 191. Tabular, posteroventral portion, emargination: absent (0), present, posteroventral portion of tabular emarginated, including the "tabular horn" structure of stem amniotes (1). Inapplicable (-) in taxa lacking a tabular (Ch. 188) and in taxa with an occipital exposure of the supratemporal (Ch. 185).**

This study.

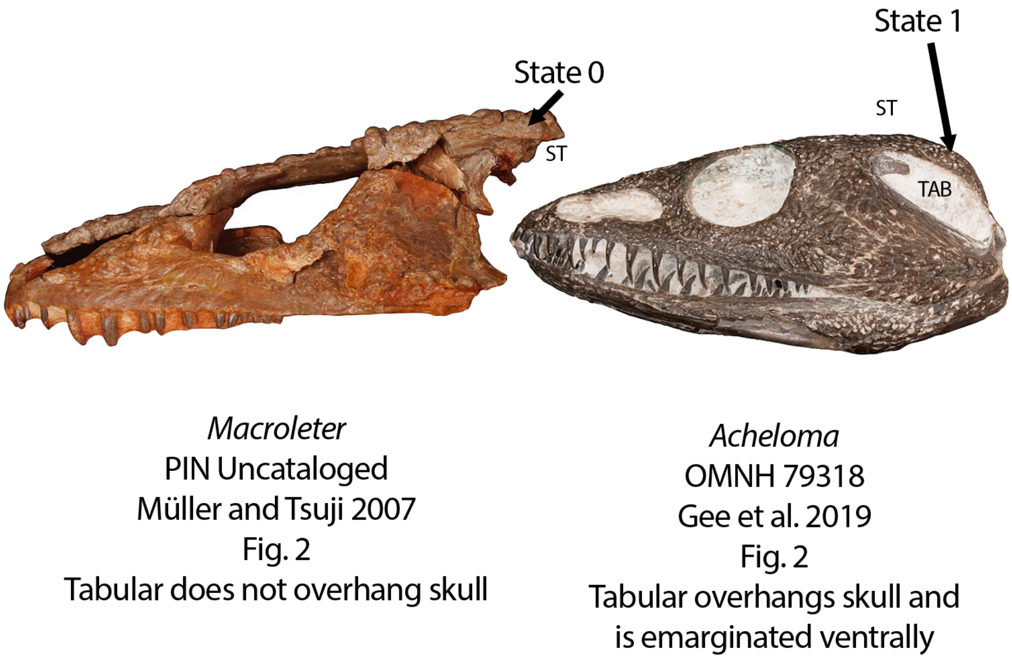


**Fig. S33.** The skulls of *Macroleter* (left, PIN uncatalogued, modified from Müller and Tsuji 2007) and *Acheloma* (right, OMNH 79318, modified from Gee et al. 2019b) in left lateral views. The tabular of *Macroleter* does not contribute to the tympanic emargination (Ch. 191, State 0), in contrast to *Acheloma*, in which the tabular bears a distinct ventrolaterally facing fossa for this emargination (Ch. 191, State 1).

Unlike early amniotes bearing an 'otic-notch' or tympanum-like structure which is restricted to the supratemporal among the temporal series (State 0 here) the 'otic' or 'spiracular' notch of early tetrapods can be present as a distinct emargination on the ventral surface of the tabular (State 1, e.g., *Balanerpeton,* UMZCT131).

UNAMBIGUOUS: *Greererpeton burkemorani* (1 to 0), Recumbirostra + Diadectomorpha + Protorothyrididae + Amniota (1 to 0).

**Ch. 192. Tabular, posteroventral portion, emargination, when present: margin is smooth or featureless (0), distinct lateral crest "tympanic crest" or frames emargination (1) Inapplicable (-) in taxa lacking a tabular or a tabular emargination (Ch. 189). (-) in taxa with an occipital exposure of the supratemporal (Ch. 187).**

This study.

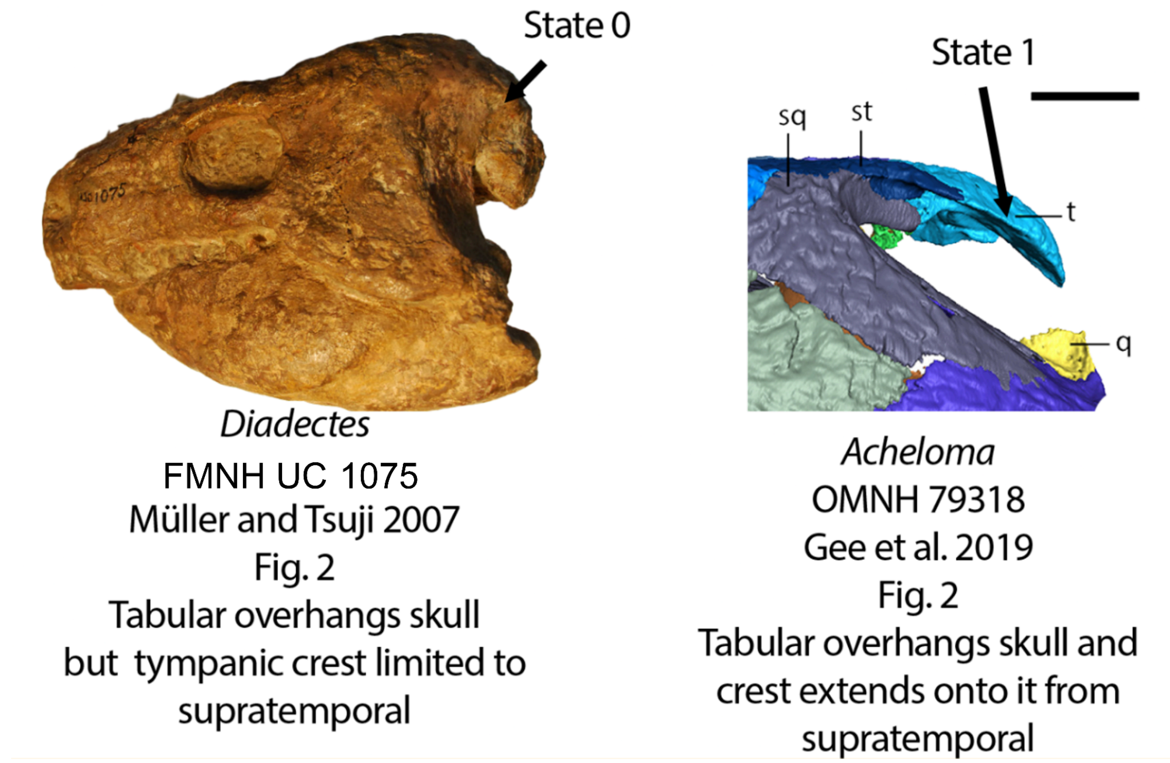


**Fig. S34.** Left, the skull of *Diadectes* (FMNH UC 1075) in which the tabular contribution to the tympanic emargination lacks a tympanic crest (Ch. 192, State 0) and right, *Acheloma* (OMNH 7938, modified from Gee et al. 2019b) which bears a distinct crest on the tabular (Ch. 192, State 1).

A 'tympanic' crest extends onto the tabular emargination (State 1) in temnospondyls within this matrix, best seen in lateral or ventrolateral view (e.g., *Acheloma*, OMNH 79318), indicating tympanic membrane or other soft tissue attachment.

UNAMBIGUOUS: Temnospondyli (0 to 1).

**Ch. 193. Tabular, occipital margin, fossae for epaxial musculature: absent, occipital surface of tabular flat or featureless (0), bears broad, shallow fossae for insertion of epaxial musculature (1). Inapplicable (-) in taxa lacking a tabular (Ch. 188).**

Modified from Pardo et al. (2017, Ch. 235).

The tabular of recumbirostran microsaurs bear distinct fossae for the insertion of the epaxial musculature on the occipital surface (State 1, e.g., *Llistrofus* OMNH 79031). This feature is also widespread in other early tetrapods including *Limnoscelis* (YPM 811) and synapsids including *Casea broilii* (Olson 1968) and can occur in tandem with posttemporal fenestra below.

DELTRAN: Amniota (1 to 0). ACCTRAN: Reptilia (1 to 0).

**Ch. 194. Tabular, posteromedial process that contacts the dorsolateral corner of the supraoccipital: absent (0), present (1).**

Modified from Brocklehurst et al. (2022, Ch. 105).

**Ch. 195. Temporal ( postorbital) roof orientation: subhorizontal or weakly inclined posteroventrally (0), strongly inclined posteroventrally, at least 45° (1).**

Modified from Brocklehurst et al. (2022, Ch. 301).

**Ch. 196. Occiput, posttemporal fenestra: absent or reduced, occipital region not perforated by a large fenestra or only a fossae or present (0), present, discrete, rounded fenestra located dorsally on occiput (1).**

Modified from deBraga and Rieppel (1997 Ch. 59), MacDougall et al. (2017, Ch. 95), Cisneros et al. (2020, Ch.93).

The occiput of generalized early tetrapodomorphs (e.g., *Archeria*, Holmes 1989) is plate-like and lacks rounded posttemporal fenestrae , although sometimes a small slit may be present (e.g., *Seymouria*, State 0). In early or near-amniotes (e.g., *Pantylus* MCZ 2040 or *Captorhinus* OMNH 44816), a rounded posttemporal fenestra pierces the occipital surface of the skull located dorsal to the paroccipital processes of the opisthotic.

DELTRAN: Amniota (0 to 1). ACCTRAN: Reptilia (1 to 0).

**Ch. 197. Synotic tectum, ossification (i.e. supraoccipital absent or unossified): no distinct ossification within the synotic tectum (0), 1 supraoccipital ossification present within synotic tectum (1).**

Modified from Pardo et al. (2017, Ch. 217).

In various early tetrapods, including temnospondyls (e.g., *Acheloma*, OMNH 79318 ) there is not a distinct ossification within the synotic tectum, instead, there is an unossified gap or a dorsal development of the exoccipital complex (State 0). Amniotes and stem-amniotes including recumbirostrans possess a distinct supraoccipital ossification within this region (State 1).

UNAMBIGUOUS: Reptiliomorpha (0 to 1).

**Ch. 198. Synotic tectum, if ossification present, supraoccipital: paired at some point in ontogeny (0), unpaired, discrete supraoccipital present (1). (-) in taxa lacking an ossification of the synotic tectum Ch. 198).**

Modified from Pardo et al. (2017, Ch. 217).

The supraoccipital is paired during ontogeny in many near-amniotes including *Seymouria* (Bazzana et al. 2019) and diadectomorphs (e.g., *Limnoscelis*, Berman et al. 1992). In most other near-amniotes, including the crown, a single ossification of the supraoccipital is present (e.g., *Captorhinus*, OMNH 44816).

DELTRAN: Gymnarthridae + *Nannaroter* + Rhynchonkidae + *Huskerpeton* (0 to 1), Amniota + Protorothyrididae + *Anthracodromeus* + *Hylonomus* + *Thuringothyris* + Captorhinidae (0 to 1), *Orobates pabsti* (0 to 1), Recumbirostra (0 to 1). ACCTRAN: *Hapsidopareion* (1 to 0), *Pantylus cordatus* (1 to 0), *Limnoscelis paludis* (1 to 0), Amniota + Protorothyrididae + Diadectomorpha + Recumbirostra + Captorhinidae + *Lanthanosuchus* + *Chroniosaurus* (0 to 1).

**Ch. 199. Synotic tectum, if ossification present, contribution to endosseous labyrinth: absent, synotic tectum does not contribute to semicircular canals (0), present, pathways of semicircular canals present within synotic tectum (1). (-) in taxa lacking an ossification of the synotic tectum Ch. 197).**

This study.

The synotic tectum contains the pathways of the anterior and posterior semicircular canals in nearly all reptiliomorphs in which element is known. In early-diverging reptiliomorphs such as *Seymouria* (Bazzana et al. 2019) and *Archeria* (Pardo 2013), the thin and platelike ossifications of the synotic tectum do not contribute to the endosseous labyrinth ventrally.

UNAMBIGUOUS: Reptiliomorpha excluding *Seymouria* (0 to 1).

**Ch. 200. Synotic tectum, endolymphatic fossae: paired fossae along the posteromedial surface of the synotic tectum for the endolymphatic sac present (0), absent (1).**

This study.

The synotic tectum, or supraoccipital, forms around the endolymphatic sac, leaving conspicuous paired fossa along the ventral surface of this element in early reptiliomorphs including *Seymouria* (Bazzana et al. 2019) and *Limnoscelis* (Klembara et al. 2021), but these fossae are unreported in recumbirostrans (e.g., *Hapsidopareion*, FMNH PR 2305) and all amniotes within this sampling excluding the pareiasauromorph *Emeroleter*, which was described by Bazzana-Adams et al. (2024) as bearing an analogous structure.

UNAMBIGUOUS: Recumbirostra (0 to 1), Captorhinidae +Protorothyrididae +Amniota (0 to 1). ACCTRAN: Recumbirostra + Diadectomorpha + Captorhinidae + Protorothyrididae + Amniota (0 to 1), Diadectomorpha (1 to 0).

**Ch. 201. Supraoccipital, lateral ascending process of supraoccipital: absent (0), present (1). Inapplicable (-) in taxa lacking a supraoccipital (Ch. 198).**

Modified from Pardo et al. (2017, Ch. 219).

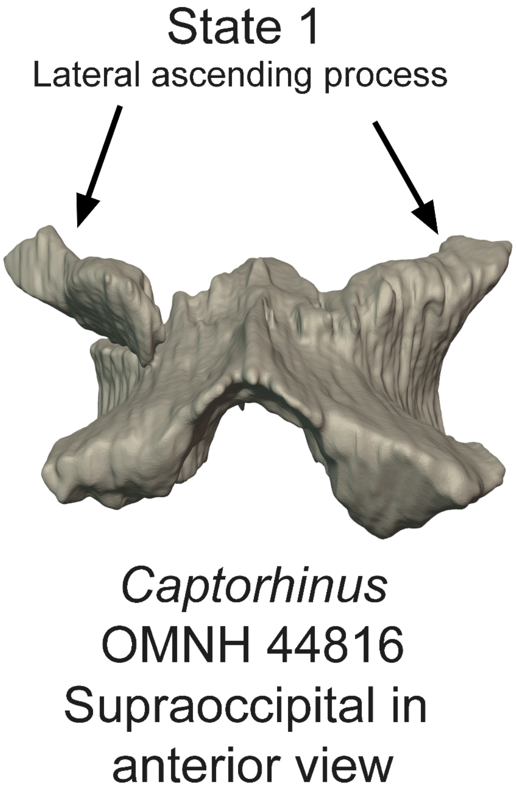


**Fig. S35.** The supraoccipital of *Captorhinus* (OMNH 44816) in anterior view, demonstrating the derived state (Ch. 201, State 1) or the presence of lateral ascending processes of the supraoccipital.

The supraoccipital of captorhinids (e.g., *Captorhinus*, OMNH 44816) and some recumbirostrans (e.g., *Huskerpeton*) possess discrete, laterally directed ascending processes which extend from the main body of the supraoccipital and underlie the postparietals ventrally. Lateral ascending processes of the supraoccipital are absent in synapsids and definitive crown reptiles within this sampling. We do not consider this homologous with the lateral expansion or processes present in some synapsids (e.g., *Aerosaurus*, UCMP 35762) and araeoscelidians (e.g., *Petrolacosaurus,* Reisz 1981), which do not extend dorsally or underlie the postparietals, but rather extend laterally over the posttemporal fenestrae.

DELTRAN: Gymnarthridae + *Nannaroter* + Rhynchonkidae + *Huskerpeton* (0 to 1), *Pantylus cordatus* (0 to 1), Captorhinidae (0 to 1). ACCTRAN: *Hapsidopareion* (1 to 0), Araeoscelidia (0 to 1), Captorhinidae (0 to 1), Recumbirostra (0 to 1), *Hapsidopareion* (1 to 0) Araeoscelidia (0 to 1), Captorhinidae (0 to 1), Recumbirostra (0 to 1).

**Ch. 202. Supraoccipital, fusion to opisthotics: unfused (0), fused (1). Inapplicable (-) in taxa lacking a supraoccipital. (Ch. 199).**

Modified from Simões et al. (2022, Ch. 131).

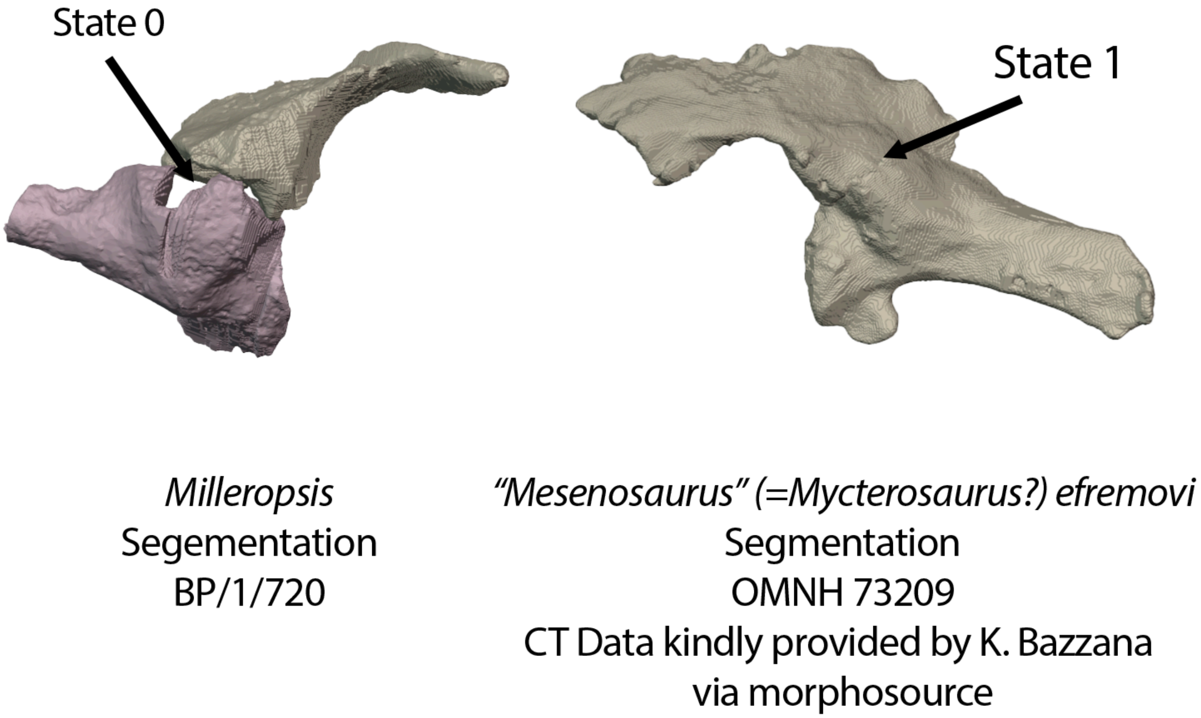


**Fig. S36.** Left, the unfused supraoccipital and opisthotic (Ch. 202, State 0) of *Milleropsis* (BP/1/720, this study) and right, the co-ossified supraoccipital and opisthotic (Ch. 202, State 1) of *M. efromovi* (OMNH 73209, segmentation from this study)

The supraoccipital is indistinguishably fused to the opisthotics in many synapsids including caseasaurs (e.g., *Euromycter*, Sigoneau-Russell and Russell 1974) and sphenacodontians (e.g., *Secodontosaurus*, AMNH 4007).

UNAMBIGUOUS: *Carrolla craddocki (*0 to 1), *Delorhynchus cifellii* (0 to 1, *Australothyris* (0 to 1), Pareiasauria (0 to 1). DELTRAN: Sphenacomorpha (0 to 1), Varanopinae (0 to 1), *Ophiacodon* spp. (0 to 1). ACCTRAN: Synapsida (0 to 1), Rhynchocephalia (0 to 1).

**Ch. 203. Supraoccipital, lateral expansion providing dorsal margin of posttemporal fenestra: absent (0), present (1).**

Modified from Brocklehurst et al. (2022, Ch. 107).

**Ch. 204. Supraoccipital, posterior surface: smooth or with a slight median ridge that grades smoothly into the posterodorsal surface of the supraoccipital (0), with a prominent vertical median ridge that rises abruptly relative to the posterodorsal surface of the supraoccipital (1).**

Modified from Brocklehurst et al. (2022, Ch. 106).

**Ch. 205. Supraoccipital, ossified medial wall for floccular or 'subarcuate' fossa: absent, floccular fossa not ossified medially (0), present, a medial flange descends from supraoccipital framing floccular fossa in bone medially (1). (-) in taxa lacking an ossification of the synotic tectum (Ch. 198).**

This study.

The supraoccipital of fully mature synapsid specimens such as varanopids (Bazzana et al. 2022) and *Dimetrodon* (Bazzana et al. 2023) possess a medial flange that frames the floccular fossa in bone. The supraoccipital lacks such a wall in all other taxa in this matrix. We note that this flange may develop during ontogeny as it is absent in juvenile specimens of *Heleosaurus* (individuals 2-4) but present in the larger (but still immature) Individual 1.

UNAMBIGUOUS: Synapsida (0 to 1).

**Ch. 206. Suproccipital, median ascending process incises the midline suture of the postparietals: absent (0), present, postparietals separated from the midline ventrally by the median ascending process of the supraoccipital (1).**

Modified from Brocklehurst et al. (2022, Ch. 110).

**Ch. 207. Supraoccipital, dorsal sinus between skull roof: absent (0), present (1). (-) in taxa lacking an ossification of the synotic tectum Ch. 197).**

Modified from Huttenlocker et al. (2013, Ch. 222) and Pardo et al. (2017, Ch. 244).

A dorsal sinus separates the supraoccipital from the skull roof (postparietals or parietals) in several early tetrapods within this study, including *Seymouria* (Bazzana et al. 2019) and non-chthonosaurian recumbirostrans (e.g., *Rhynchonkos*). In all other OTU's, the supraoccipital is in tight articulation with the skull roof dorsally and the occiput ventrally (State 0).

UNAMBIGUOUS: Gymnarthridae + Rhynchonkidae + *Huskerpeton* (0 → 1), *Seymouria* (0 to 1).

**Ch. 208. Postparietals: present (0), absent (1).**

Modified from Brocklehurst et al. (2022, Ch. 108).

**Ch. 209. Postparietals: unfused/paired (0), fused to form a midline element (1).**

Modified from Brocklehurst et al. (2022, Ch. 109).

**Ch. 210. Postparietals, size, transverse width of paired or single medial element: equal to or greater than the transverse width of one parietal at its widest point (0), less than the transverse width of one parietal (1).**

Modified from Brocklehurst et al. (2022, Ch. 111).

**Ch. 211. Postparietals, dorsal and occipital exposure: dorsally exposed parts of postparietals anteroposteriorly longer than mediolaterally wide (0), dorsal mediolateral width between 1-4 times the anteroposterior length (1), more than four times as wide (mediolaterally) as long (2) or absent, postparietals entirely on occipital surface of skull (3). ORDERED.**

Modified from Marjanovic and Laurin (2019, Ch. 40)

**Ch. 212. Postparietals, contribution to margin of post-temporal fenestra: absent (0), present (1).**

Modified from Brocklehurst et al. (2022, Ch. 112).

**Ch. 213. Postparietal(s) or parietals, ventral contact with exoccipital: absent, contact excluded by supraoccipital or opisthotics (0), present (1). (-) in taxa lacking the ossification of the postparietals (Ch. 208).**

Modified from Ruta and Coates (2007, Ch. 51) and Marjanovic and Laurin (2019, Ch. 135)

The postparietal or parietals of amniotes and diadectomorphs are restricted to contacting the supraoccipital ventrally (State 0). In early tetrapods (e.g., *Whatcheeria*, Rawsonet al. 2021), temnospondyls, and non-chthonosaurian recumbirostrans, the postparietals contact the exoccipitals ventrally, lateral to the synotic tectum and dorsomedial to the opisthotics (State 1).

UNAMBIGUOUS: Reptiliomorpha (1 → 0). ACCTRAN: Reptiliomorpha (1 → 0), Embolomerii (1 → 0), Gymnarthridae + Rhynchonkidae + *Huskerpeton* (0 → 1).

**Ch. 214. Postparietal(s) or parietals, ventral contact with opisthotic: absent, contact excluded by supraoccipital (0), present (1). (-) in taxa lacking ossification of the postparietals (Ch. 208).**

Modified from Ruta and Coates (2007, Ch. 51) and Marjanovic and Laurin (2019, Ch. 135)

Parietal or postparietal contact with the opisthotic ventrally near the margin of the foramen magnum is lost in all taxa closer to Amniota than Recumbirostra (State 0) by an expansion of the occipital exposure of the supraoccipital.

UNAMBIGUOUS: Captorhinidae + Diadectomorpha + Amniota (1 → 0).

**Ch. 215. Opisthotic, paroccipital process, posterior extension: not visible in lateral view (0), visible in lateral view (1).**

Modified from Brocklehurst et al. (2022, Ch. 113).

**Ch. 216. Opisthotic, morphology and orientation of paroccipital process: robust and rod- like, extends laterally or ventrolaterally (0), slender and rod-like, extends posteroventrolaterally (1), dorsoventrally broad sheet with short lateral extension (2), dorsoventrally broad sheet, extends laterally or dorsolaterally (3).**

**Ch. 217. Opisthotic, paroccipital process attachment: contacts skull laterally (0), ends freely (1).**

Modified from Brocklehurst et al. (2022, Ch. 114).

**Ch. 218. Opisthotic, if paroccipital process contacts skull laterally, bony contact: contact restricted to quadrate or squamosal, no contact with skull roof (0), contacts the supratemporal and squamosal (1), contacts the tabular (2).**

Reworked from Brocklehurst et al. (2022, Ch. 115).

Rationale: The original phrasing of this character concerned the contact between the paroccipital process and the tabular. However, the paroccipital process can also contact the quadrate, squamosal, or supratemporal, and we codify this here.

**Ch. 219. Opisthotic, placement of metotic foramen relative to hypoglossal foramina (CN XII): anterior to hypoglossal foramen (0), level with or posterior to hypoglossal foramen (1)**

Modified from Hamley et al (2021, Ch. 63).

The hypoglossal foramina are located level with or anterior to the metotic foramen only within procolophonids (e.g., *Eomurruna*, QMF 59501) within this sampling (State 1). Modified from Hamley et al (2021, Ch. 63).

DELTRAN: Procolophonidae (0 to 1), Procolophonoidea + *Nyctiphruretus* (0 to 1).

**Ch. 220. Opisthotic, paroccipital process, distal end: maintains shape or tapers distally (0), anteroposteriorly-flattened distally (1).**

Modified from Pritchard et al. (2015 Ch. 63, Ezcurra 2016 Ch. 215).

The paroccipital process of crocopod archosauromorphs (e.g., *Mesosuchus,* SAM-PK-6536) and varanopine synapsids (e.g., *Aerosaurus,* UCMP 35762*)* becomes anteroposteriorly-flatted in cross-section distally (State 1).

DELTRAN: Varanopinae (0 to 1), Dinocephalia + *Biseridens* (0 to 1), Tanystropheidae + Crocopoda (0 to 1). ACCTRAN: *Secodontosaurus obtusidens* (1 to 0), Sphenacodontoidea + *Haptodus* + *Cutleria* + *Tetraceratops* (0 to 1), Varanopinae + *Apsisaurus* (0 to 1), Crocopoda (0 to 1).

**Ch. 221. Opisthotic, fusion with prootic: absent, elements of the otic capsule remain discrete elements throughout ontogeny (0), present, opisthotic and prootic fused, forming single "otic" or "periotic" element in adults (1).**

This study.

The opisthotic is co-ossified with the prootic in early synapsids (e.g., *Haptodus*, ROM 43604) and brachystelechid recumbirostrans (Pardo and Anderson 2016) within the current sampling (State 1).

UNAMBIGUOUS: Brachystelechidae (0 to 1). DELTRAN: Pareiasauromorpha + *Australothyris* (0 to 1), *Clevosaurus hudsoni* (0 to 1), *Tanystropheus hydroide*s (0 to 1) *Mesosuchus browni* (0 to 1), Amniota + Protorothyrididae + Diadectomorpha + Recumbirostra + Captorhinidae (1 to 0). ACCTRAN: Owenettidae (0 to 1), Lepidosauria (0 to 1), Synapsida (0 to 1), *Limnoscelis paludis* (0 to 1), Reptiliomorpha (1 to 0).

**Ch. 222. Opisthotic, ventral ramus: plate-like and rectilinear (0), subcircular in cross section and thinning ventrally (1).**

Modified from Pritchard et al. (2017, Ch. 57), Ezcurra (201, Ch. 217).

Primitively in amniotes, the ventral ramus of the opisthotic is plate-like and rectilinear in cross-section (e.g., *Varanops*, Romer and Price 1940). In parapleurotan reptiles, the ventral ramus thins ventrally and becomes circular in cross-section (e.g., *Milleropsis,* BP/1/720 or *Youngina* BP/1/2871).

DELTRAN: Parapleurota + *Orovenator* + *Ascendonanus* + *Cabarzia* (0 to 1). ACCTRAN: Mesosauridae + *Ascendonanus* + *Orovenator* + *Cabarzia* + Parapleurota (0 to 1).

**Ch. 223. Opisthotic, ventral ramus, anteroventrally-facing "otic" trough: absent, ventral ramus of opisthotic is convex anteroventrolaterally (0), present, an anteroventral "trough" extends from the cochlear recess to the fenestra ovalis (1).**

Modified from Berman et al. (1992 Ch. 5), Pardo et al. (2017 Ch. 229), Klembara et al. (2021 Ch. 280).

In diadectomorphs (e.g., *Limnoscelis* YPM 811) and early synapsids including caseasaurs (e.g.,*Casea broilii*, Olson 1968) and varanopids (e.g., *Varanops* and *Heleosaurus*, this study), a trough on the ventral surface of the opisthotic extends posteriorly from the fenestra ovalis. This trough is absent in most other early tetrapods including *Seymouria* (OMNH 79342), recumbirostrans (Pardo 2013), captorhinids (e.g., *Captorhinus laticeps*, OMNH 44816) and all non-mesosaurid neoreptiles in which this region is known (e.g., *Milleropsis,* BP/1/720), although this region is not well-described historically, especially in two-dimensional specimens.

UNAMBIGUOUS: Synapsida (0 to 1). ACCTRAN: Diadectomorpha (0 to 1).

**Ch. 224. Opisthotic, ventral ramus, crista interfenestralis: absent, internal (medial) surface of ventral ramus of opisthotic lacks a medially-placed crest separating the metotic fossa from the vestibule (0), present, crista interfenestralis separates the metotic fossa from the vestibule (1)**

Modified from Pardo et al. (2017, Ch. 32).

The crista interfenestralis is a crest or ridge on the medial surface of the opisthotic that separates the metotic fossa from the vestibule, which implies an anterior path of perilymph circulation as (State 1, Pardo and Anderson 2016). It is present in recumbirostrans (e.g., *Rhynchonkos*, FM-UR 1039), captorhinids (e.g., *Captorhinus*, OMNH 44816), and all known early amniotes in which this feature is observable (e.g., *Mycterosaurus*, FMNH UC 692). In contrast, early tetrapods or reptiliomorphs lack such a division between the metotic fossa and vestibule, and it is probable that in these taxa that the perilymph circulation is located more posteriorly, as in extant lissamphibians (State 0).

DELTRAN: Reptiliomorpha excluding *Seymouria* (0 to 1). ACCTRAN: Reptiliomorpha (0 to 1).

**Ch. 225. Opisthotic, lagenar crest (*sensu* Heaton 1979), development: absent or incipient, opisthotic lacks a lagenar crest/ridge separating the lagena and scala tympani (0), present, a well-developed lagenar crest, separates the scala tympani from the lagena (1)**

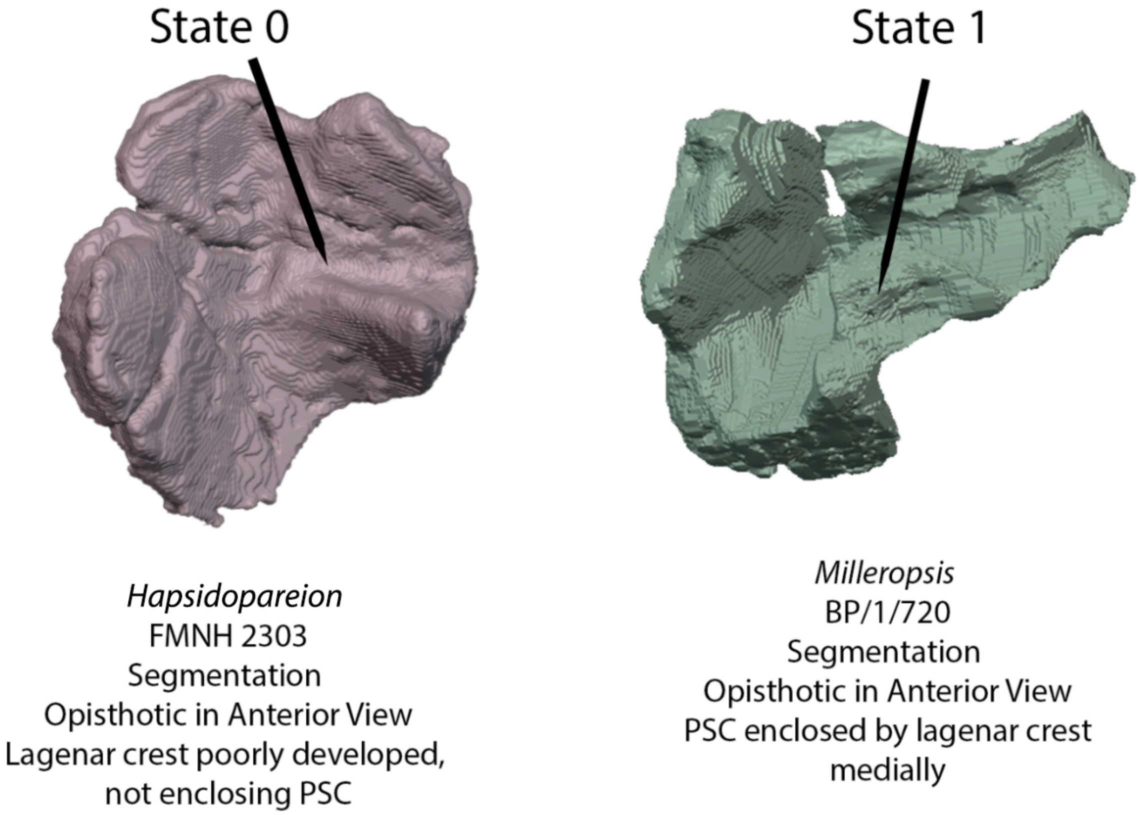


**Fig. S37.** Left, left opisthotic of *Hapsidopareion* (FMNH 2303) which bears a poorly developed lagenar crest (Ch. 225, State 0) and right, left opisthotic of *Milleropsis pricei* (BP/1/720, this study) which bears a strongly developed lagenar crest that canalized the medial portion of the posterior semicircular canal (Ch. 225, State 1).

The lagenar crest of the opisthotic separates the lagenar recess from scala tympani in stem and crown amniotes within this sampling (e.g., *Milleropsis*, BP/1/720). In non-pantylid recumbirostrans, (e.g., *Llistrofus*, Gee et al. 2019a), the lagenar crest is absent or extremely reduced, and the pathway of the posterior semicircular canal remains open to the lagenar recess, possibly related to the reduction in medial ossification of the prootic (State 0).

DELTRAN: Recumbirostra excluding *Pantylus* (1 to 0), Recumbirostra + Captorhinidae + Protorothyrididae + Diadectomorpha + Amniota ( 0 to 1). ACCTRAN: Reptiliomorpha (0 to 1).

**Ch. 226. Otic capsule, posterior semicircular canal: ventral end of the PSC confluent with utricular region, at least partially (0), ventral end separated from utricular region by ossification of the opisthotic (1)**

**Modified from Pardo et al. (2017 Ch. 253).**

The posterior semicircular canal is medially exposed to the utricular region in most early tetrapods within this sampling (e.g., *Dendrerpeton*, Robinson et al. 2005) or *Milleretta* BP/1/3822 (State 0). In many synapsids, including varanopids (e.g., *Mycterosaurus*, FMNH UC 692) the posterior semicircular canal is no longer exposed medially due to the development of an utricular crest of the opisthotic, fully canalizing the pathway of the posterior semicircular canal (State 1).

ACCTRAN: Synapsida (0 to 1).

**Ch. 227: Inner ear, ampullary fossa, definition: ampullary fossa present as distinct expanded fossae at the junction of the semicircular canals (0), absent, no distinct fossae/expansion at the junction of semicircular canals (1).**

This study.

Nearly all tetrapods within this sampling possess well-defined ampullary fossae for the semicircular canals (State 1). However, ampullary fossa are absent or undefined within the otic capsules of early reptiliomorphs such as *Seymouria* (Bazzana et al. 2021) and diadectomorphs (Klembara et al. 2021).

UNAMBIGUOUS: *Seymouria* (0 to 1), Limnoscelis (0 to 1).

**Ch. 228. Prootic, lateral surface: continuous and slightly convex, lacking crista prootica (0), low anterodorsally directed ridge extends onto articulation with opisthotic (1), distinct crest or 'crista prootica' present (2). ORDERED.**
Modified from Brocklehurst et al. (2022, Ch. 117).

We added a third state, state (0), concerning the low ridge expressed on the lateral surface of the prootic in millerettids (e.g., *Milleretta*, Jenkins et al., in press) and non-saurian neodiapsids such as *Youngina* (Gardner et al. (2010).

ACCTRAN: Parapleurota ( 0 to 1), Reptile crown group (1 to 2).

**Ch. 229. Prootic, anterior inferior process: absent, trigeminal foramen (cranial nerve V) open anteriorly (0), present, partially enclosing anterior margin of trigeminal foramen (cranial nerve V) (1).**

Modified from Brocklehurst et al. (2022, Ch. 118).

**Ch. 230. Prootic, medial surface of vestibule: unossified such that the vestibule and semicircular canals are not separated from utrical (0) partially ossified, path of ASC fully separated from utricle but LSC remains unossified (1), fully ossified, separating the vestibule semicircular canals from the endocranial cavity (2). ORDERED.**

This study.

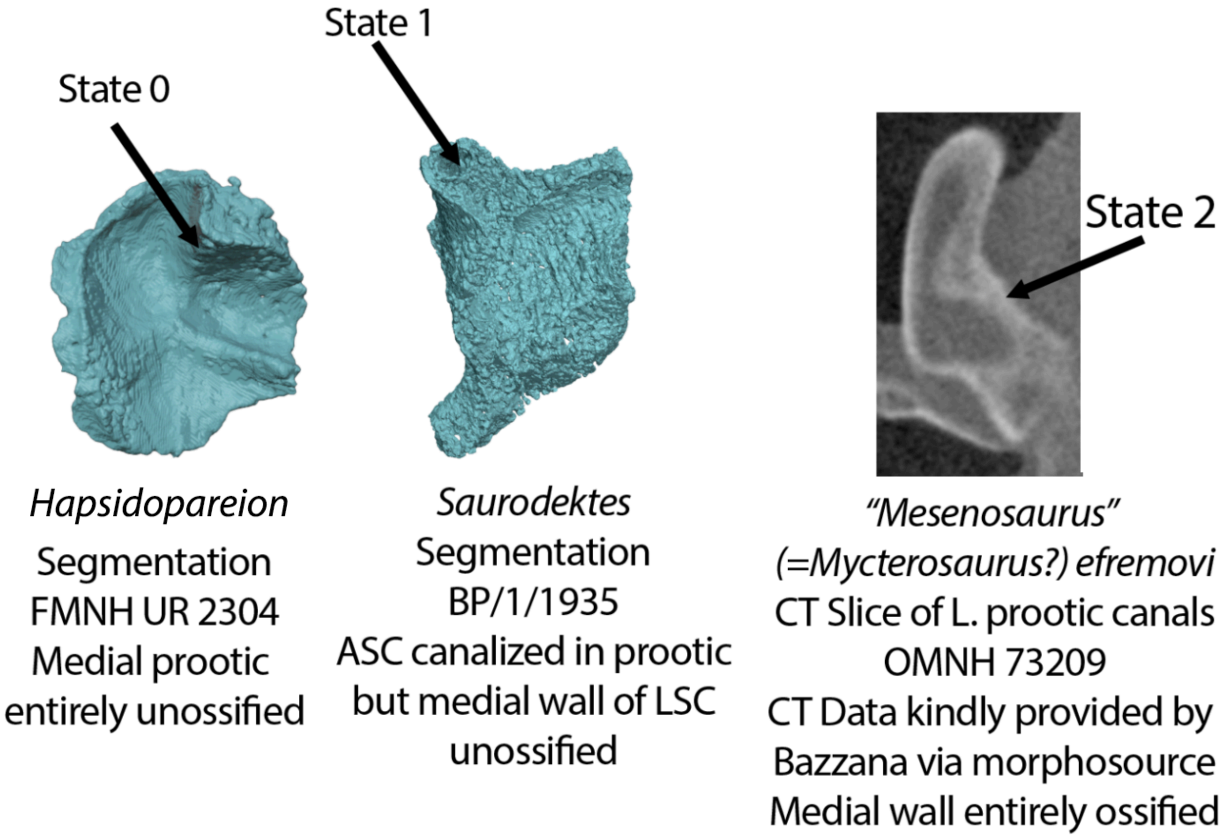


**Fig. S38.** Left, left prootic of *Hapsidopareion* (FMNH UR 2304) in medial view with an entirely unossified medial wall (State 0). Middle, left prootic of *Saurodektes* (BP/1/1935, this study) in which the medial wall of the prootic is ossified along the course of the anterior semicircular canal but open posteriorly (State 1) and right, slice data of *Mesenosaurus efremovi* (OMNH 73209) which bears an entirely ossified medial wall of the prootic (State 2).

In early tetrapods within this sampling, the medial surface of the prootic is poorly ossified such that the vestibule and semicircular canals are not separated from the utricular recess (State 0, e.g., *Dendrerpeton*, Robinson et al. 2005). In many Permian amniotes, the ASC is fully separated from the utricle although the LSC remains open ventrally (State 1,e.g., *Milleropsis*), whereas the ASC and LSC are fully separated from the utricle in non-caseasaurian synapsids in the sampling here (State 2, *Mycterosaurus*, FMNH UC 692).

DELTRAN: Eupelycosauria + Varanopidae (1 to 2), Pareiasauromorpha + *Australothyris*, (1 to 2), Diadectomorpha + Protorothyrididae + Amniota (0 to 1). ACCTRAN: Eupelycosauria (1 to 2), Amniota + Protorothyrididae + *Anthracodromeus* (0 to 1), Diadectomorpha (0 to 1).

**Ch. 231. Prootic, ossification of crista alaris: absent or incompletely ossified (0), present, anterodorsal surface of prootic bears an anterolaterally oriented rounded ridge for the anterior semicircular canal (1).**

**This study.**

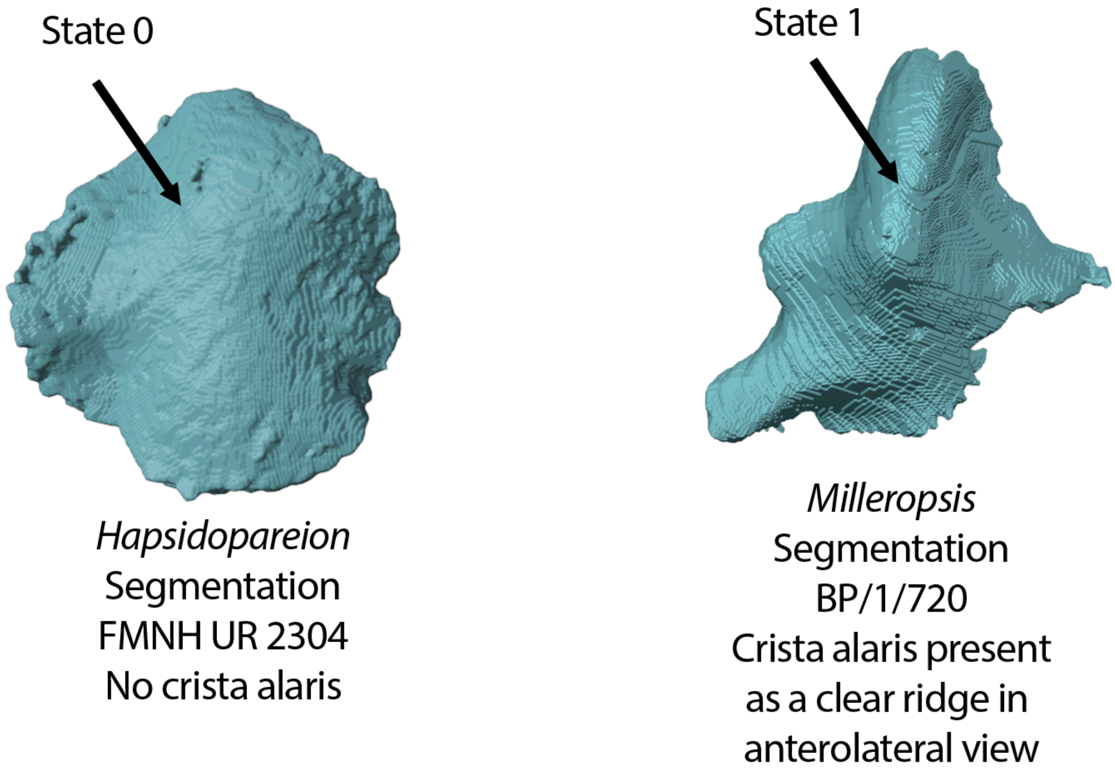


**Fig. S39.** Prootics of *Hapsidopareion* (FMNH UR 2304) and *Milleropsis* (BP/1/720) in anterolateral view. The prootic of *Hapsidopareion* bears a poorly developed crista alaris, being little more than a slight convexity (Ch. 231, State 0) whereas the crista alaris is present as a distinct bulge in *Milleropsis* (Ch. 231, State 1).

The crista alaris, a rounded ridge present on the anterodorsal surface of the prootic marking the pathway of the anterior semicircular canal is present (State 0) in most amniotes (e.g., *Ophiacodon*, Romer and Price 1940) and near-amniotes (e.g., *Captorhinus*, OMNH 44816). The prootic of recumbirostrans (e.g., *Hapsidopareion*, FMNH 2303) tends to be poorly ossified with the pathway of the semicircular canals not demarcated, such that the anterior surface of the prootic is flat or mildly convex (State 0), although this may be due to miniaturization in these taxa). State 0 is also reported in varanopine synapsids (Bazzana et al. 2022), in which the anterior semicircular canal is supposed to be restricted to the supraoccipital.

DELTRAN: Varanopinae (1 to 0) Captorhinidae+ Protorothyrididae+ Amniota (0 to 1), Diadectomorpha + Protorothyrididae + Amniota (0 to 1). ACCTRAN: Varanopinae + *Apsisaurus* (1 to 0).

**Ch. 232. Prootic, floccular fossa: absent, dorsomedial surface of prootic medial to ASC flat or convex (0), present, distinct concavity on medial surface of prootic for floccular or subarcuate fossa (1).**

**Modified from Klembara et al. (2020, Ch. 278).**

The prootic of all reptiliomorphs more crownard of *Seymouria* in this study (e.g., the recumbirostran *Pantylus*, Romer 1968) possess a floccular fossa on the dorsomedial surface, although it is absent in some derived recumbirostrans (e.g., *Hapsidopareion lepton,* FMNH 2305), probably a consequence of miniaturization (Maddin et al. 2011).

DELTRAN: *Pantylus cordatus* (0 to 1). ACCTRAN: Recumbirostra + Captorhinidae + Diadectomorpha + Protorothyrididae + Amniota (0 to 1), Recumbirostra excluding *Pantylus* (1 to 0).

**Ch. 233. Prootic, position of trigeminal nerve (CN V): anteroventrally, immediately posteromedial to prootic articulation with clinoid process of Parabasisphenoid and ventrally level with the prootic contribution to the fenestra ovalis (0), anterodorsally, CN V located on the dorsal surface of the prootic, immediately posterior to the alar process, if present, and dorsally elevated relative to fenestra ovalis (1), anterior surface of prootic lacks a discrete incisure for the trigeminal (2).**

**This study.**

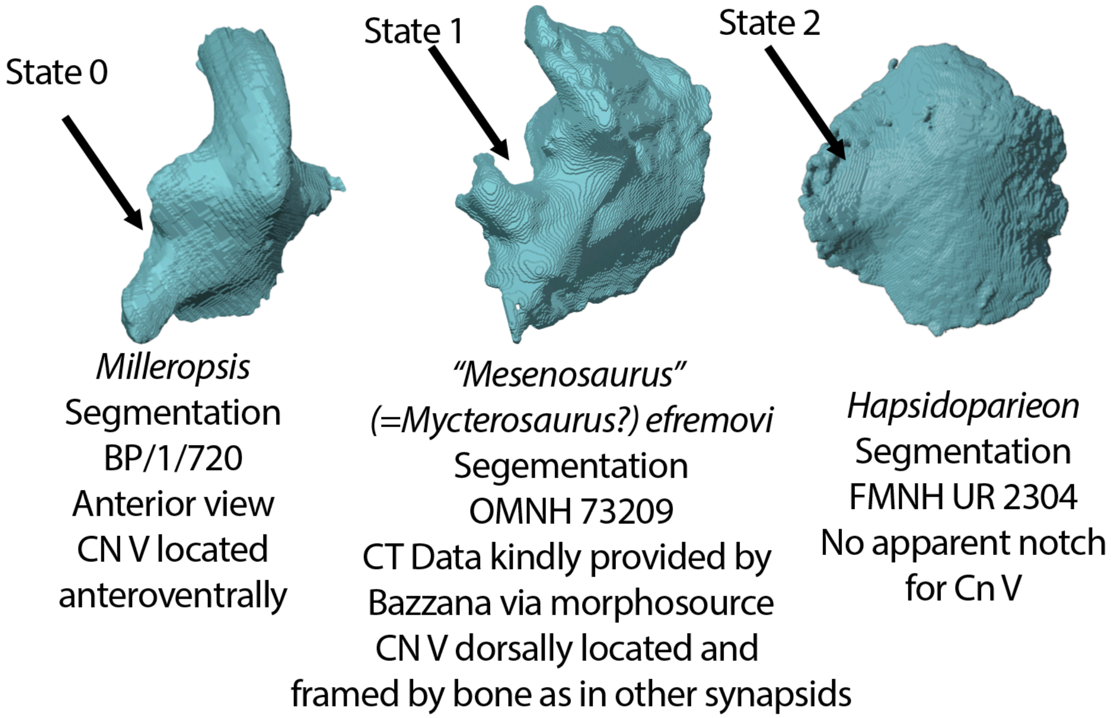


**Fig. S40.** Left, left prootic of *Milleropsis* (BP/1/720, this study) in anterior view, demonstrating an opening for CN V along its anteroventral surface (Ch. 233, State 0). Middle, left prootic of *Mesenosaurus efremovi* in anterior view, bearing a dorsally positioned opening for CN V (Ch. 233, State 1). Right, left prootic of *Hapsidopareion* (FMNH UR 2304) which lacks a distinct opening for CN V (Ch. 233, State 2).

The trigeminal notch or prootic incisure remains on the anteroventral surface of the prootic (State 0) in most sampled tetrapods within this matrix (e.g., *Youngina,* BP/1/2871 or *Milleropsis* BP/1/720). However, in recumbirostrans, the opening for CN V likely travels through an unossified gap between the anterior surface of the prootic (State 2) and the pleurosphenoids (e.g., *Hapsidopareion*, FMNH 2305). In early synapsids such as mesenosaurines (Botha-Brink and Modesto 2009) *Heleosaurus* (this study), the trigeminal notch is located on the dorsal surface of the prootic (State 1), and is elevated dorsally relative to the fenestra ovalis (e.g., *Dimetrodon* or *Ophiacodon*, see Romer and Price 1940).

UNAMBIGUOUS: *Edops craigi* (2 to 1), *Deltavjatia rossica* (0 to 1), Synapsida (0 to 1). DELTRAN: Amniota + Diadectomorpha + Protorothyrididae + Captorhinidae (2 to 0).

**Ch. 234. Prootic, morphology of facial nerve (CN VII): fully enclosed foramen (0), notch on ventral surface of inferior process (1).**

This study.

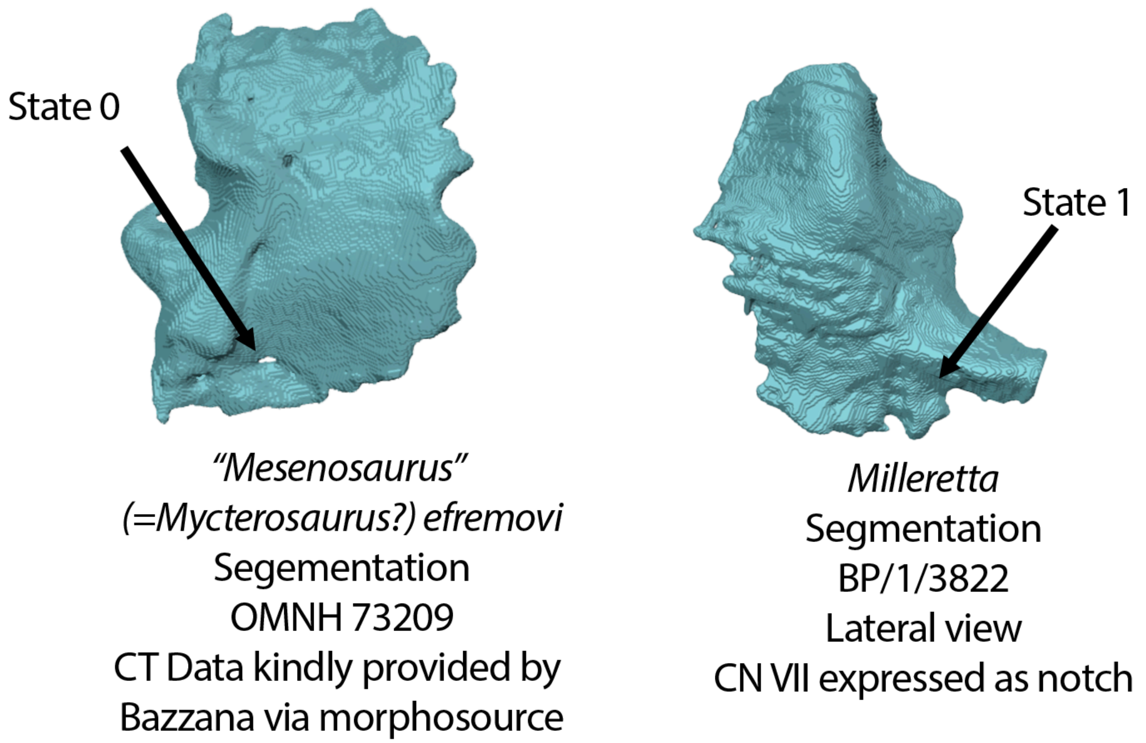


**Fig. S41** Left, left prootic of *Mesenosaurus efremovi* (OMNH 73209) in lateral view, bearing a foramen for CN VII along its ventral surface (Ch. 234, State 0). Right, left prootic of *Milleropsis* (BP/1/720, this study) which bears a notch for CN VII (Ch. 234, State 1).

The facial nerve (CN VII) is fully enclosed as a foramen (State 0) in most taxa sampled here (e.g., *Leptopleuron*, NMS 1984.20.4), however, it remains a notch (State II) in millerettids (e.g., *Milleropsis*, BP/1/720). This character should only be scored in ontogenetically mature taxa, as it appears to vary through ontogeny (Hamley et al. 2021).

ACCTRAN: Moradisaurinae + *Labidosaurus* (0 to 1), Lepidosauromorpha(0 to 1), Millerettidae (0 to 1).

**Ch. 235. Prootic, contact with its counterpart on the median line of the floor of the endocranial cavity "dorsum sellae: absent (0), present, contributing to dorsum sellae (1).**

Modified from Ezcurra (2016, Ch. 253).

The prootic contacts its counterpart (State 1), forming the dorsum sellae in sphenacomorph synapsids (e.g., *Dimetrodon,* Romer and Price 1940) and pareiasaurs (e.g., *Bunostegos*, MNN MOR47) within this sampling.

UNAMBIGUOUS: Bradysauridae (0 to 1), Sphenacomorpha (0 to 1).

**Ch. 236. Prootic, articulation with epipterygoid: absent (0), present, anterior surface of prootic bears an elongate facet for epipterygoid (1).**

Modified from Pardo et al. (2017, Ch. 230).

The prootic contacts the dorsal process of the epipterygoid (State 1) in leptopleuronines (e.g., *Leptopleuron* NMS 1984.20.4) and archosauriforms (e.g., *Euparkeria,* SAM-PK-7696) in the taxa sampled here.

UNAMBIGUOUS: Leptopleuroninae (0 to 1). DELTRAN Allokotosauria + *Mesosuchus* + Archosauriformes (0 to 1). ACCTRAN: Archosauriformes (0 to 1), Lepidosauromorpha (0 to 1), Temnospondyli (0 to 1).

**Ch. 237. Exoccipital, lateral wing: tall and narrow (0), broad, extending ventral to paroccipital process (1).**

Modified from Brocklehurst et al. (2022, Ch. 119).

**Ch. 238. Exoccipitals enclose the complete ventral margin of the foramen magnum: absent, margin provided by the basioccipital (0), present (1).**

Modified from Brocklehurst et al. (2022, Ch. 120).

**Ch. 239. Exoccipitals, number of metotic foramina: single foramen (0), paired foramina for XII located ventrolateral to wing (1).**

Modified from Gower & Sennikov (1996), Ezcurra, Lecuona & Martinelli (2010, Ch. 23), Dilkes & Arcucci, (2012, Ch. 30).

The exoccipitals of early tetrapods (e.g., *Greererpeton* CMNH 11068, *Edops* MCZ 1378) possess a single opening for the hypoglossal nerve (State 0). In contrast, the exoccipitals of amniotes (e.g., *Ophiacodon*, Romer and Price 1940) and captorhinids (*C. laticeps*, Heaton 1979) possess paired foramina for CN XII (State 1).

UNAMBIGUOUS: *Haptodus garnettensis* (1 to 0). DELTRAN: *Tanystropheus hydroides* (1 to 0) Amniota + Protorothyrididae + Captorhinidae (0 to 1), *Limnoscelis paludis* (0 to 1). ACCTRAN: Diadectomorpha + Protorothyrididae + Amniota (0 to 1).

**Ch. 240. Exoccipitals, fusion to opisthotic: unfused (0), fused (1).**

Simões et al. (2022, Ch. 155).

The exoccipitals of sphenacodontine synapsids (e.g., *Dimetrodon,* Romer and Price 1940) and archosauromorphs (e.g., *Azendohsaurus*, UA-7-20-99-653) are fused to the opisthotics (State 1) in the taxonomic sampling here. The fusion of the otic complex (prootic, opisthotic, and exoccipitals) is independent from one another in several taxonomic groups within this matrix.

UNAMBIGUOUS: Brachystelechidae (0 to 1). DELTRAN: Dinocephalia + *Biseridens* (0 to 1), *Acleistorhinus pteroticus* (0 to 1), *Bunostegos akokanensis* (0 to 1), Pareiasauria (0 to 1), *Tanystropheus hydroides* (0 to 1), *Trilophosaurus buettneri* (0 to 1), *Azendohsaurus madagaskarensis* (0 to 1), *Euparkeria capensis* (0 to 1). ACCTRAN: *Mesosuchus browni* (1 to 0), *Proterosuchus* spp. (1 to 0), Allokotosauria + *Mesosuchus*+ Archosauriformes (0 to 1), Sphenacodontoidea + *Haptodus* + *Cutleria* + *Tetraceratops* (0 to 1).

**Ch. 241. Exoccipitals, fusion to prootic: unfused (0), fused (1).**

Simões et al. (2022, Ch. 155).

The exoccipitals of sphenacodontine synapsids (e.g., *Dimetrodon*, Romer and Price 1940) are fused to the prootic (State 1), whereas the exoccipitals of archosauromorphs (Ezcurra 2016) and most other early amniotes remain unfused to the prootic (State 0). The fusion of the otic complex (prootic, opisthotic, and exoccipitals) are independent of one another in several taxonomic groups within this matrix.

UNAMBIGUOUS: Brachystelechidae (0 to 1). DELTRAN: Dinocephalia + *Biseridens* (0 to 1). ACCTRAN: *Secodontosaurus obtusidens* (1 to 0), Sphenacodontia (0 to 1).

**Ch. 242. Exoccipital, accessory articulation facets for proatlas: absent (0), present (1).**

Modified from Pardo et al. (2017, Ch. 254).

The posterior surface of the exoccipitals bears facets for the proatlas in most reptiliomorphs sampled here (State 0). In contrast, the exoccipital of ankyramorph neoreptiles (e.g., *Delorhynchus,* OMNH 73515) do not bear a facet indicating an osseous contact with the proatlas (State 1).

UNAMBIGUOUS: Ankyramorpha + *Erpetonyx* (1 to 0). DELTRAN: Tanystropheidae + Crocopoda (0 to 1), Temnospondyli (1 to 0), (0 to 1). ACCTRAN: Crocopoda (0 to 1), Amniota + Protorothyrididae + Diadectomorpha + Recumbirostra + Captorhinidae (0 to 1), Embolomeri (0 to 1).

**Ch. 243. Exoccipital, contribution to occipital condyle: absent (0), present (1).**

Modified from Pardo et (2017, Ch. 255).

The exoccipital of recumbirostrans are incorporated into the occipital condyle (e.g., *Euryodus dalyae,* FM UR 2296).

UNAMBIGUOUS: Recumbirostra (0 to 1).

**Ch. 244. Exoccipital, tubercle ventral to proatlantal facet absent (0), present, a tubercle is present ventral to the articulation for the proatlas (1).**

Based on observations of Liu et al. (2022).

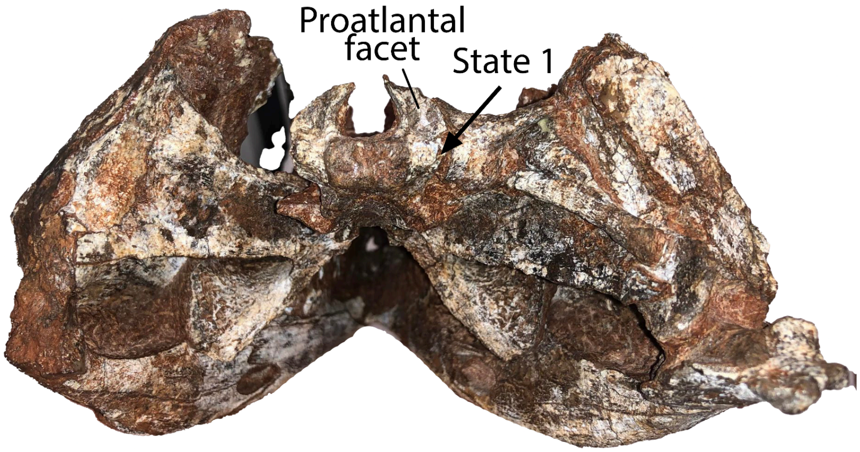


**Fig. S42.** Skull of *Moradisaurus grandis* (MNN MOR149) in occipital view demonstrating the presence of an exoccipital tubercle ventral to the proatlantal facet (Ch. 244, State 1).

The Lopingian moradisaurine captorhinids *Moradisaurus* and *Gansurhinus* bear a tubercle ventral to the proatlantal facet (Sidor et al. 2021 fig. 1A, Lui et. al. 2022).

UNAMBIGUOUS: *Moradisaurus* + *Gansurhinus* (0 to 1).

**Ch. 245. Basioccipital, fusion to exoccipital: unfused (0), fused (1).**

Modified from Simões (2018, Ch. 218).

The exoccipitals fuse to the (State 1) to the basioccipital in many early tetrapods, including some temnospondyls (e.g., *Dendrerpeton,* Robinson et al. 2005), recumbirostrans (e.g., *Carrolla*, TMM 40031-54) and several groups of synapsids (e.g., *Mycterosaurus*, FMNH-UC 692) and reptiles such as pareiasaurs (e.g., *Bunostegos,* MNN MOR47). The fusion of the otoccipital complex (prootic, opisthotic, and exoccipitals) are independent of one another in several taxonomic groups within this matrix.

UNAMBIGUOUS: *Dendrerpeton helogenes* (0 to 1), Brachystelechidae (0 to 1) *Captorhinus laticeps* (1 to 0), *Moradisaurus grandi*s (1 to 0), *Embrithosaurus schwarz*i (1 to 0) Pareiasauromorpha + *Australothyris* (0 to 1). ACCTRAN: Millerettidae (0 to 1), Mesosauridae (0 to 1), Protorothyrididae (0 to 1), Captorhinidae (0 to 1), Ophiacodontidae (1 to 0), Eupelycosauria + Varanopidae (0 to 1).

**Ch. 246. Basioccipital, occipital condyle orientation: posteriorly directed (0), posteroventrally directed (1).**

Modified from Brocklehurst et al. (2022, Ch. 121).

**Ch. 247. Basioccipital, paired ventrolateral tubera: absent or poorly developed, not extending well ventral of occipital condyle (0), well developed, extending ventral to level of occipital condyle (1).**

Modified from Ezcurra (2016, Ch. 226).

The basioccipital of eupelycosaurs (e.g., *Dimetrodon,* Romer and Price 1940), procolophonoids (e.g., *Owenetta*, BP/1/4195), and saurians (e.g., *Euparkeria,* SAM-PK-7696) possess paired ventrolateral tubera that extend ventrally further than the occipital condyle in occipital view (State 1).

DELTRAN: Therapsida (1 to 0), Eupelycosauria + Varanopidae (0 to 1), *Araeoscelis* spp. (0 to 1), Procolophonoidea (0 to 1), *Claudiosaurus* + Sauria (0 to 1), *Thadeosaurus colcanapi* (0 to 1) *Youngina capensis* (0 to 1). ACCTRAN: Dinocephalia + *Biseridens* (1 to 0), Eupelycosauria (0 to 1) Bolosauridae (0 to 1), Procolophonia (0 to 1), Weigeltisauridae (1 to 0), *Hovasaurus boulei* (1 to 0).

**Ch. 248. Basioccipital-parasphenoid/parabasisphenoid, contact with each other in mature individuals: loose, overlapping suture (0), tightly sutured, sometimes by an interdigitated suture, or both bones fused to each other (1).**

Modified from Ezcurra (2016, Ch. 255).

In most Paleozoic tetrapods, contact between the parasphenoid and basioccipital consists of an overlapping suture (State 0) between the two elements (e.g., *Captorhinus laticeps*, OMNH 44816) Heaton 1979). In synapsids (e.g., *Cotylorhynchus romeri*, Romer and Price 1940), pareiasaurs (e.g., *Bunostegos* MNN MOR47) and archosauromorphs, the suture between these two elements is strongly interdigitated or these elements are fused together (State 1).

DELTRAN: Brachystelechidae (0 to 1), *Nannaroter mckinzei* (0 to 1), *Cotylorhynchus romeri* (1 to 0), Pareiasauromorpha + *Australothyris* (0 to 1), *Bellairsia gracilis* (0 to 1), *Euparkeria capensis* (0 to 1), Archosauriformes (0 to 1), Bolosauridae (0 to 1). ACCTRAN: Chthonosauria (0 to 1) *Nannaroter mckinzei* (0 to 1), *Cotylorhynchus* + *Angelosaurus* (1 to 0), Owenettidae (0 to 1), *Bellairsia* + Squamata (0 to 1), *Proterosuchus* spp. (1 to 0), Allokotosauria + *Mesosuchus* + Archosauriformes (0 to 1) Synapsida (0 to 1)

**Ch. 249. Basioccipital, relation to fenestra ovalis: Basioccipital possesses a ventromedial flange contributing to the FO (0), basioccipital lacks a ventromedial flange and a contribution to the FO is absent or reduced (1).**

Modified from Pardo et al. (2017, Ch. 202).

The basioccipital contributes to the ventromedial margin of the fenestra ovalis (State 0) in most tetrapods within this sampling. However, the basioccipital does not contribute to the fenestra ovalis (State 1) in some pareiasaurs (e.g., *Deltavjatia,* Tsuji 2013) and many neodiapsids excluding *Mesosuchus* (SAM-PK-6536).

UNAMBIGUOUS: *Acheloma cumminsi* (0 to 1), Bradysauridae (0 to 1). DELTRAN: *Greererpeton burkemorani* (0 to 1), *Mesosuchus browni* (1 to 0), *Seymouria* spp. (0 to 1). ACCTRAN: *Mesosuchus browni* (1 to 0), *Seymouria* spp. (0 to 1), *Crassigyrinus* + *Greererpeton* + *Baphetes* (0 to 1), Parapleurota (0 to 1).

**Ch. 250. Basioccipital, articular surface of the occipital condyle: concave (0), convex and hemispherical (1).**

Modified from Ezcurra (2016, Ch. 229)

The occipital condyle of the basioccipital is generally concave (State 0) in Paleozoic and Mesozoic tetrapods, possessing a notochordal pit in occipital view (e.g., *Planocephalosaurus* Fraser 1982). In therapsids (e.g., *Biarmosuchus,* Kemp 1982) and archosauromorphs (e.g., *Azendohsaurus* UA-7-20-99-653) within this sampling, the occipital condyle is strongly convex (State 1).

UNAMBIGUOUS: Archosauromorpha (0 to 1). DELTRAN: Therapsida (0 to 1). ACCTRAN: Dinocephalia + *Biseridens* (0 to 1).

**Ch. 251. Basioccipital, dorsolateral surface, conical recess: absent (0), present (1).**

Modified from Pardo et al. (2017, Ch. 252).

The conical recess is a fossa on the dorsolateral surface of the basioccipital which receives an expansion of vestibule (State 1) in recumbirostran ‘microsaurs’ (e.g., *Brachydectes*, USNM 32149).

UNAMBIGUOUS: Recumbirostra (0 to 1).

**Ch. 252. Basioccipital, ventral surface anterior to occipital condyle: smooth (0), prominent anteroposterior ridges (1).**

Modified from Brocklehurst et al. (2022, Ch. 313).

**Ch. 253. Basal articulation, position: approximately level with posterior rim of the transverse flange of pterygoid (0) anterior to rim of the transverse flange (1), posterior to the rim transverse flange (2).**

Modified from Brocklehurst et al. (2022, Ch. 122).

**Ch. 254. Basal articulation, orientation of basipterygoid process: anteroventrolaterally directed (0), laterally directed (1), distinctly ventrally or posteroventrally directed (2).**

Modified from Brocklehurst et al. (2022, Ch. 123).

**Ch. 255. Basal articulation, morphology of articular surface of basipterygoid process: single, rounded articular surface (0), flat anterior facet (1), possesses an anterodorsal and anteroventral facet (2).**

Modified from Brocklehurst et al. (2022, Ch. 315).

**Ch. 256. Basal articulation, fusion between pterygoid/epipterygoid and parabasisphenoid: absent, pterygoid/epipterygoid unfused and articulates with the parabasisphenoid in a ball-in-socket or overlapping joint (0), present, pterygoid/epipterygoid sutured/fused with the Parabasisphenoid ventrally (1).**

Reworked from Brocklehurst et al. (2022, Ch. 309).

Rationale: The original verbiage of this character concerns the presence or absence of a basicranial articulation, following the terminology of therapsid work. However, most work on reptiles concerns the mobility of the basicranial articulation or the fusion of the elements involved in it, and we rephrase to here.

**Ch. 257. Parabasisphenoid, dentition posterior to level of transverse flange: absent (0), present (1).**

Modified from Brocklehurst et al. (2022, Ch. 124).

**Ch. 258. Parabasisphenoid, if dentition posterior to transverse flange of the pterygoid present: abundant, denticle and/or tooth fields extending between the cristae ventrolaterales (0), dentition reduced to rows on lateral margins of cristae ventrolaterales (1), row of 3-6 teeth on ventral plate medial to cristae ventrolaterales (2), isolated denticles restricted to anterior margin of ventral plate (3) . Inapplicable (-) in taxa lacking dentition posterior to level of transverse flange (Ch. 257).**

This study.

When there is dentition on the parabasisphenoid posterior to the level of the transverse flange of the pterygoid, it is usually present as an abundant denticle field that extends between the crista ventrolaterales (State 0) in early tetrapods and amniotes (e.g., *Varanops*, OMNH 73157). In contrast, the parabasisphenoid of the acleistorhinid *Acleistorhinus* (FM UR 1038), mesenosaurines (e.g., *Mesenosaurus*, PIN 3706/11) and millerettids (e.g., *Milleropsis*, BP/1/720) possess a discrete row of dentition just medial to the crista ventrolaterales (State 1).

DELTRAN: *Orovenator* + *Ascendonanus* + *Cabarzia* (1 to 3), Millerettidae (1 to 2), Reptilia (0 to 1).
ACCTRAN: Parapleurota (1 to 2), *Orovenator* + *Ascendonanus* + *Cabarzia* (1 to 3), Neoreptilia (0 to 1).

**Ch. 259. Parabasisphenoid, dentition on ventral surface of cultriform process: absent (0), present (1).**

Modified from Brocklehurst et al. (2022, Ch. 125).

**Ch. 260. Parabasisphenoid, entry foramen for the internal carotid arteries: on the ventral surface of the parabasisphenoid, medial to the basipterygoid process (0), within the lateral wall of braincase, dorsal to the basipterygoid process (1).**

Modified from Brocklehurst et al. (2022, Ch. 126).

**Ch. 261. Parabasisphenoid, lateral parasphenoidal crests present as prominent lateral extensions framing the ventromedial floor of the vidian sulcus, and obscuring the entry foramen of the internal carotid artery from ventral view (0), absent, such that there is no ventral floor to the passage of the internal carotid artery, and the entry foramina of the internal carotid are visible in ventral view (1).**

Modified from Brocklehurst et al. (2022, Ch. 127).

**Ch. 262. Parabasisphenoid, body shape: transversely broad, width greater than length from basipterygoid processes to posterior end (0), transversely narrow, length greater than width (1).**

Modified from Brocklehurst et al. (2022, Ch. 128).

**Ch. 263. Parasphenoid body, posteroventral emargination [basisphenoid shelf]: absent (0), present (1).**

Modified from Brocklehurst et al. (2022, Ch. 316).

**Ch. 264. Parabasisphenoid, outline of lateral margin, anterior ‘shoulder’ level with the basal articulation in ventral view: absent (0), present (1).**

Modified from Brocklehurst et al. (2022, Ch. 129).

**Ch. 265. Parabasisphenoid, ventral plate, medial excavation: depression or concavity between the cristae ventrolaterales (0), deep median groove present (1), absent, ventral surface flat (2).**

Modified from Brocklehurst et al. (2022, Ch. 130).

**Ch. 266. Parabasisphenoid, ventral plate, median longitudinal septum or ridge: absent (0), present (1).**

Modified from Brocklehurst et al. (2022, Ch. 131).

**Ch. 267. Parabasisphenoid, cultriform process, length: short or absent, less than 3/4 the anteroposterior length of main body of parasphenoid (0), between one 3/4 and 2 times length of main body of parasphenoid (1), more than 2x the length of the main body of Parabasisphenoid (2). ORDERED.**

Modified from Laurin and Reisz (1995, Ch. 53)

The cultriform process of ankyramorph neoreptiles is anteroposteriorly reduced (e.g., *Saurodektes,* BP/1/4195), shorter than the main body of the Parabasisphenoid (State 0), whereas the cultriform process of ophiacodontids is elongate, more than 2x the length of the main body of the parasphenoid (State 2, Romer and Price 1940). In most other taxa within this matrix, the cultriform process is intermediate in length, between 75% and 200% the length of the main body of the parasphenoid (State 1).

DELTRAN: Ophiacodontidae (1 to 2), Therapsida (1 to 0), *Feeserpeton oklahomense* (0 to 1), Ankyramorpha + *Erpetonyx* (1 to 0). ACCTRAN: Dinocephalia + *Biseridens* (1 to 0), Ophiacodontidae (1 to 2), *Feeserpeton* + *Delorhynchus*, (0 to 1), *Megachirella* + *Marmoretta* (1 to 0), *Trilophosaurus buettneri* (1 to 0), *Orovenator* + *Ascendonanus* + *Cabarzia* (1 to 2)

**Ch. 268. Parabasisphenoid, cultriform process, orientation: cultriform process extends anteriorly (0), cultriform process extends slightly dorsally at roughly 15 degrees to the basal plane (1), cultriform process extends anterodorsally at more than 45 degrees to the basal plane (2). ORDERED.**

Modified from Modesto et al. (2018). Ch. 45

**Ch. 269. Parabasisphenod, cultriform process, cristae dorsolaterales: absent, the dorsal surface of the cultriform process is smooth or mildly concave (0), present, the cultriform process bears two anteroposteriorly running crest which forms a v-shaped trough on the dorsal surface of the cultriform process, in many cases supporting the sphenethmoid and presumably the unossified interorbital cartilage (1).**

This study.

The cultriform process of most tetrapods (e.g., *Archeria*, Clack and Holmes 1988) within this matrix is V-shaped in cross section due to the development of two anteroposteriorly running crests, the crista dorsolaterales (State 1), forming a trough that likely supports the cartilaginous interorbital septum. In contrast, the cultriform process of recumbirostrans (e.g., *Euryodus*, FM UR 2296) is flat or mildly concave dorsally, possibly related to miniaturization in these forms.

UNAMBIGUOUS: Recumbirostra (1 to 0).

**Ch. 270. Parabasisphenoid, cultriform process, sutural contact with vomer: absent, cultriform process ends freely (0), present, cultriform process extends far anteriorly and sutures with posterior end of vomer (1). (-) in taxa possessing an anteroposteriorly reduced cultriform process (Ch. 267, State 0).**

Modified from Pardo et al. (2017, Ch. 202).

In most early tetrapods and amniotes, the cultriform process of the Parabasisphenoid ends freely anteriorly, either rising dorsally above the palatal surface or ending in the interpterygoid vacuities (e.g., *Captorhinus*, Heaton 1979). The cultriform process of temnospondyls (e.g., *Edops*, MCZ 1378) within this sampling sutures with the posterior margin of the vomer (State 1).

UNAMBIGUOUS: Temnospondyli (0 to 1).

**Ch. 271. Parabasisphenoid, relation to the fenestra ovalis: Parabasisphenoid possesses a posteroventral flange contributing to the FO (0), contribution to FO absent, parabasisphenoid lacks a posteroventral flange (1).**

Modified from Pardo et al. (2017, Ch. 220).

Primitively in amniotes, the parabasisphenoid, basioccipital, prootic, and opisthotic contribute to the fenestra ovalis. The parabasisphenoid is excluded from the ventral margin of the fenestra ovalis (State 1) by the prootic and opisthotic in some saurian reptiles (e.g., *Clevosaurus*, NHMUK PV R36832).

DELTRAN: *Clevosaurus hudsoni* (0 to 1), *Tanystropheus hydroides* (0 to 1), *Trilophosaurus buettneri* (0 to 1). ACCTRAN: *Trilophosaurus buettneri* (0 to 1), Crocopoda (1 to 0), Weigeltisauridae + *Claudiosaurus* + Sauria (0 to 1).

**Ch. 272. Parasphenoid: contribution to the anterior surfaces of the basal tubera: flat or convex (0), parasphenoid forms posterolaterally-facing 'semilunar depression' that warp around the anterior surfaces of the basal tubera (1). (-) in taxa lacking a basioccipital tuber (Ch. 247).**

Modified from Gower and Sennikov (1996), Nesbitt (2011, Ch. 98), Dilkes and Arcucci (2012 Ch. 34), Ezcurra (2016 Ch. 238).

The parasphenoid of crocopod archosauromorphs (e.g., *Mesosuchus*, SAM-PK-6536 ) possess a posterolaterally facing, crescent-shaped fossa that wraps around the anterior surfaces of the basal tubera (State 1).

DELTRAN: *Clevosaurus hudsoni* (0 to 1), *Trilophosaurus buettneri* (1 to 0), Tanystropheidae + Crocopoda (0 to 1). ACCTRAN: *Clevosaurus hudsoni* (0 to 1), *Trilophosaurus buettneri* (1 to 0) Crocopoda (0 to 1).

**Ch. 273. Parabasisphenoid, posterodorsal portion at contact with prootic: completely ossified, braincase ossified along suture between prootic and parabasisphenoid (0), incompletely ossified throughout ontogeny, such as that there is a gap between the prootic and parasphenoid posterior to clinoid processes (1).**

Modified from Ezcurra (2016 Ch. 236).

In most tetrapods, there is an unossified gap between the clinoid processes of the Parabasisphenoid ventral to the prootic (e.g., *Macronemus*, PIMUZ T 2477). In some eupelycosaurs (e.g., *Dimetrodon),* pareisauromorphs (e.g., *Bunostegos* MNN MOR47), and crocopod archosauromorphs (e.g., *Euparkeria,* SAM-PK-7696) this "gap" is completely closed, filled by extensions of the prootic and parabasisphenoid.

UNAMBIGUOUS: Chthonosauria (1 to 0) DELTRAN: Pareiasauria + *Emeroleter* (1 to 0), *Tanystropheus hydroides* (1 to 0), Crocopoda (1 to 0), Bolosauridae (1 to 0), Embolomeri (0 to 1). ACCTRAN: Pareiasauromorpha (1 to 0), *Tanystropheus hydroides* (1 to 0), Allokotosauria + *Mesosuchus* + Archosauriformes (1 to 0), Synapsida (1 to 0), *Archeria crassidiscus* (1 to 0).

**Ch. 274. Parabasisphenoid, retractor pit, midline ridge: absent (0), present, ridge extends anteriorly from dorsum sellae partially dividing the fossae into two (1).**

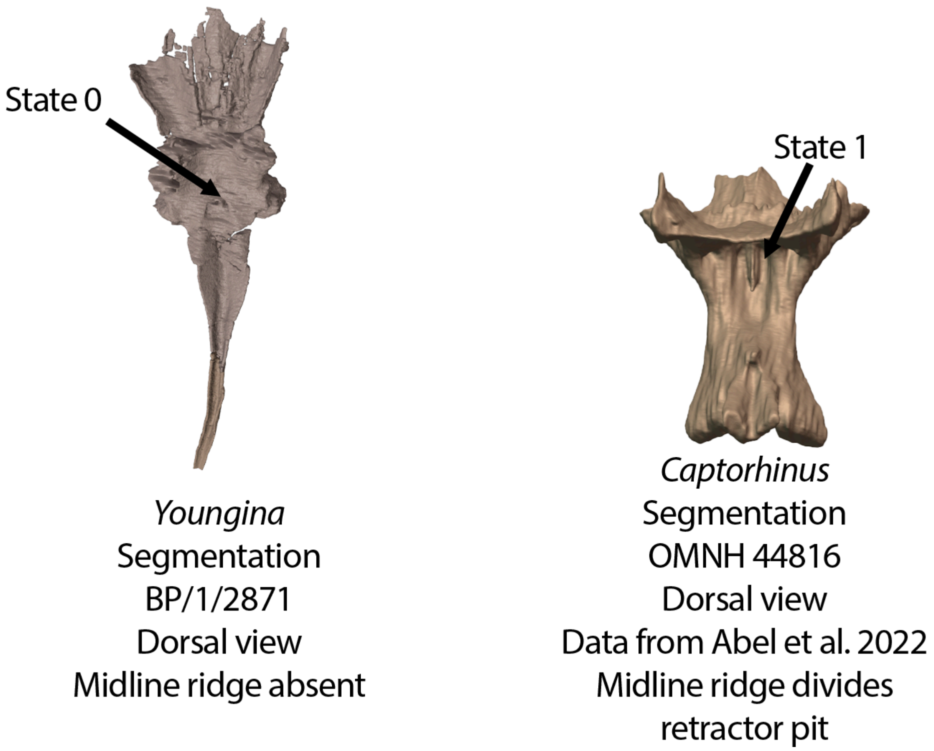


**Fig. S43.** Parabasisphenoids of *Youngina* (BP/1/2871) and *Captorhinus* (OMNH 44816) in dorsal view. The pituitary fossa of *Youngina* lacks a midline ridge (Ch. 274, State 0) whereas a midline ridge is present in *Captorhinus* dividing the pituitary fossa into left and right halves (Ch. 274, State 1).

This study.

The presence of a midline ridge (State 1) dividing the retractor pit is highly plastic in reptiliomorphs, but is nearly always present in several clades including captorhinids (e.g., *Captorhinus* OMNH 44816) and procolophonids (e.g., *Eomurruna*, QMF 59501).

DELTRAN: Moradisaurinae + *Labidosaurus* + *Gansurhinus* + *Captorhinikos* + *Labidosaurikos* (0 to 1), *Varanops brevirostris* (0 to 1), *Mycterosaurus longiceps* (0 to 1), *Varanosaurus acutirostris* (0 to 1), *Belebey vegrandis* (0 to 1), Procolophonidae (0 to 1), *Mesosuchus browni* (0 to 1), *Euparkeria capensis* (0 to 1). ACCTRAN: *Aerosaurus wellesi* (1 to 0), *Varanosaurus acutirostris* (0 to 1), Varanopinae + *Apsisaurus* (0 to 1), Bolosauridae (0 to 1) Procolophonoidea + *Nyctiphruretus* (0 to 1), *Trilophosaurus buettneri* (1 to 0), Allokotosauria + *Mesosuchus* + Archosauriformes (0 to 1), Captorhinidae (0 to 1).

**Ch. 275. Parabasisphenoid, location of hypophyseal (pituitary) fossa: anterior margin of hypophyseal fossa well posterior to or level with basipterygoid process (0), located anteriorly, anterior margin of hypophyseal fossa extends anterior to basipterygoid processes (1).**

This study, based on observations noted in Szostakiwskyj et al. (2015).

The hypophyseal fossa of most early tetrapods is located posterior to the basipterygoid processes (State 0, e.g., *Limnoscelis,* YPM 811). In recumbirostran ‘microsaurs’, the anterior margin of the hypophyseal fossa extends anterior to the basipterygoid processes (e.g., *Llistrofus* OMNH 79031).

UNAMBIGUOUS: *Edops craigi* (0 to 1), *Seymouria* spp (0 to 1), *Procolophon trigoniceps* (0 to 1),Temnospondyli + Reptiliomorpha + *Crassigyrinus* + *Greererpeton* + *Baphetes* (1 to 0). DELTRAN: Rhynchonkidae (0 to 1), *Hapsidopareion* (0 to 1), *Deltavjatia rossica* (0 to 1), . ACCTRAN: Chthonosauria (1 to 0), *Huskerpeton englehorni* (1 to 0), Recumbirostra (0 to 1), Pareiasauromorpha (0 to 1).

**Ch. 276. Parabasisphenoid, ossification flanking dorsum sellae: none, dorsum sellae is absent or the sole ossification in this region (0), paired ossifications parallel with sagittal plane "pleurosphenoids" (1), oblique or perpendicular to sagittal plane "laterosphenoid" or "pila antotica" of some studies (2).**

Modified from Pardo et al. (2017, Ch. 214).

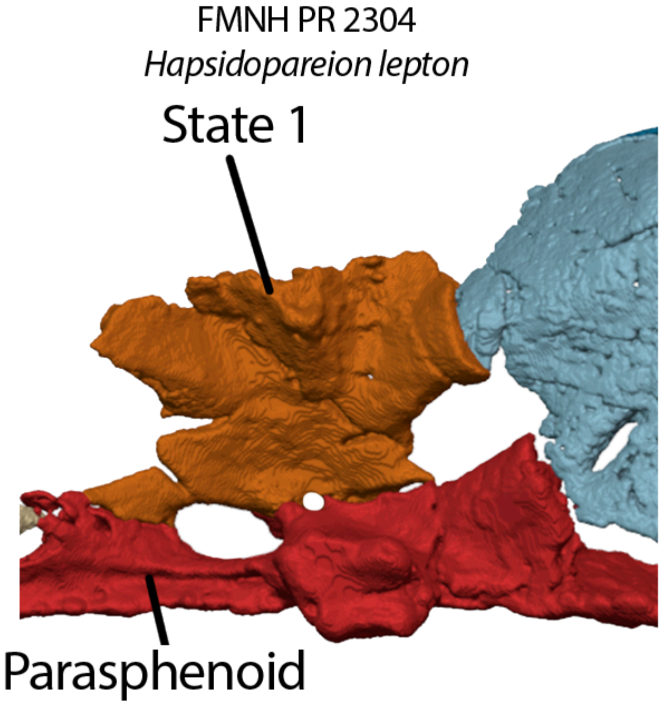


**Fig. S44.** Braincase of *Hapsidopareion* (FMNH PR 2304) in lateral view, demonstrating the presence of a paired ossification of the pila antotica often referred to as a 'pleurosphenoid' (Ch. 276, State 1).

The pila antotica, an orbital cartilage in the otic region, remains unossified in most taxa within this matrix (State 0). In recumbirostrans (e.g., *Euryodus* FM UR 2296), paired ossifications (often called 'pleurosphenoids' in the literature) flank the region of the dorsum sellae (State 1). In many archosauromorphs, the pila antotica is ossified as a single midline element, the laterosphenoid (State 2, e.g., *Euparkeria*, SAM-PK-5867).

UNAMBIGUOUS: Recumbirostra (0 to 1). DELTRAN: *Tanystropheus hydroides* (0 to 2), *Trilophosaurus buettneri* (2 to 0), Crocopoda (0 to 2). ACCTRAN: *Prolacerta broomi* (2 to 0), Archosauromorpha (0 to 2),

**Ch. 277. Parabasisphenoid, crista sellaris, development of dorsum sellae: dorsum sellae low, dorsoventrally equal to or lower than 3x the dorsoventral height of the basipterygoid process (0), tall, dorsum sellae dorsoventrally higher than 3x the dorsoventral height of the basipterygoid processes (1). (-) in taxa lacking a dorsum sellae developed from the parabasisphenoid.**

This study.

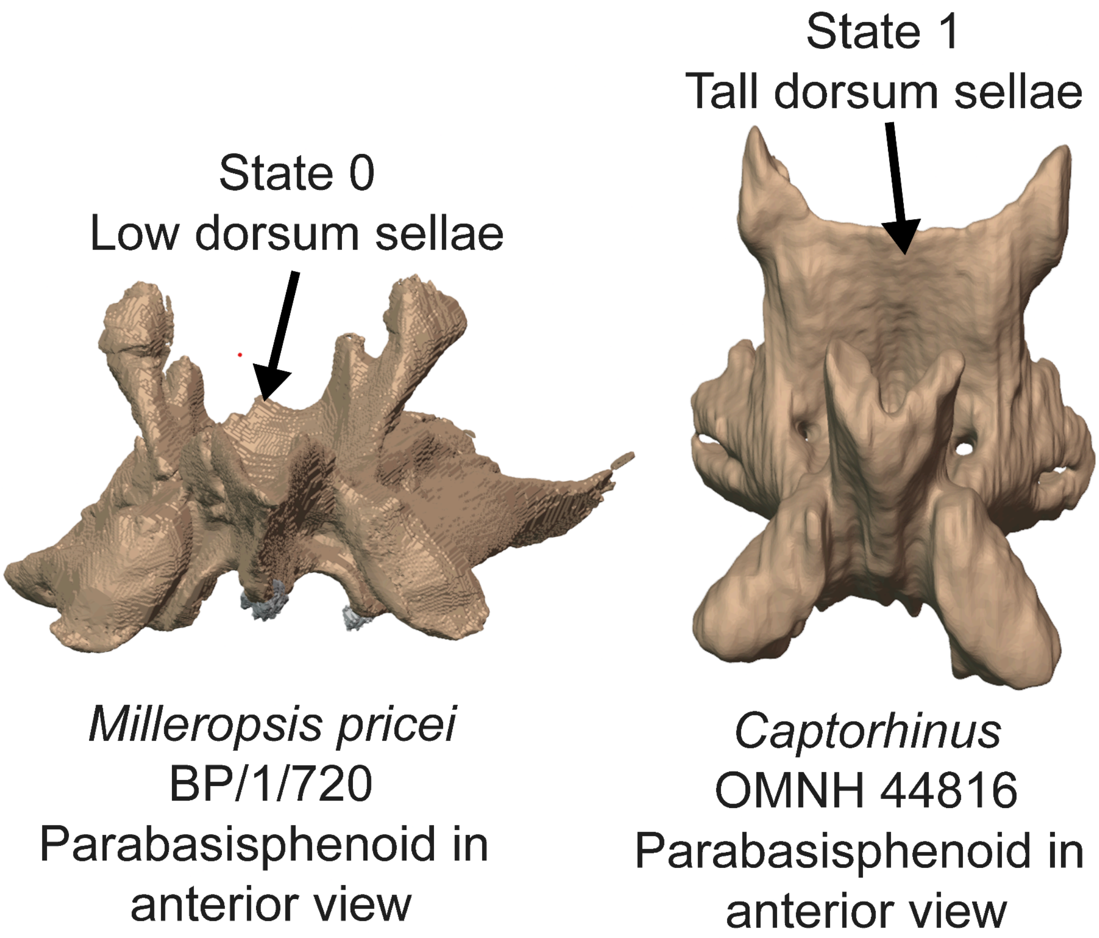


**Fig. S45.** Parabasisphenoids of *Milleropsis* (BP/1/720, this study) and *Captorhinus* (OMNH 44816) in anterior view demonstrating a low dorsum sellae (Ch. 277, State 0) and a tall dorsum sellae (Ch. 277, State 1), respectively.

The crista sellaris or dorsum sellae is relatively low (State 0) in most early tetrapods and amniotes (e.g., *Milleropsis* BP/1/720) within this matrix. In captorhinids (e.g., *Captorhinus*, OMNH 44816) and pantylid ‘microsaurs’ (e.g., *Pantylus*, MCZ 2040), the dorsum sellae is markedly expanded (State 1), more than 3x the height of the basipterygoid processes, nearly reaching the skull roof dorsally.

UNAMBIGUOUS: *Pantylus cordatus* (0 to 1). DELTRAN: *Labidosaurus hamatus* (1 to 0), *Thuringothyris* + Captorhinidae (0 to 1). ACCTRAN: *Labidosaurus hamatus* (1 to 0).

**Ch. 278. Parabasisphenoid, abducens nerve foramina (CN VI), location: within wall of dorsum sellae (0), pierces clinoid process or prootic articulation (1)**

Modified from Nesbitt (2011, Ch. 122); Ezcurra, Scheyer & Butler (2014, Ch. 156), and Pritchard et al., (2015, Ch. 71).

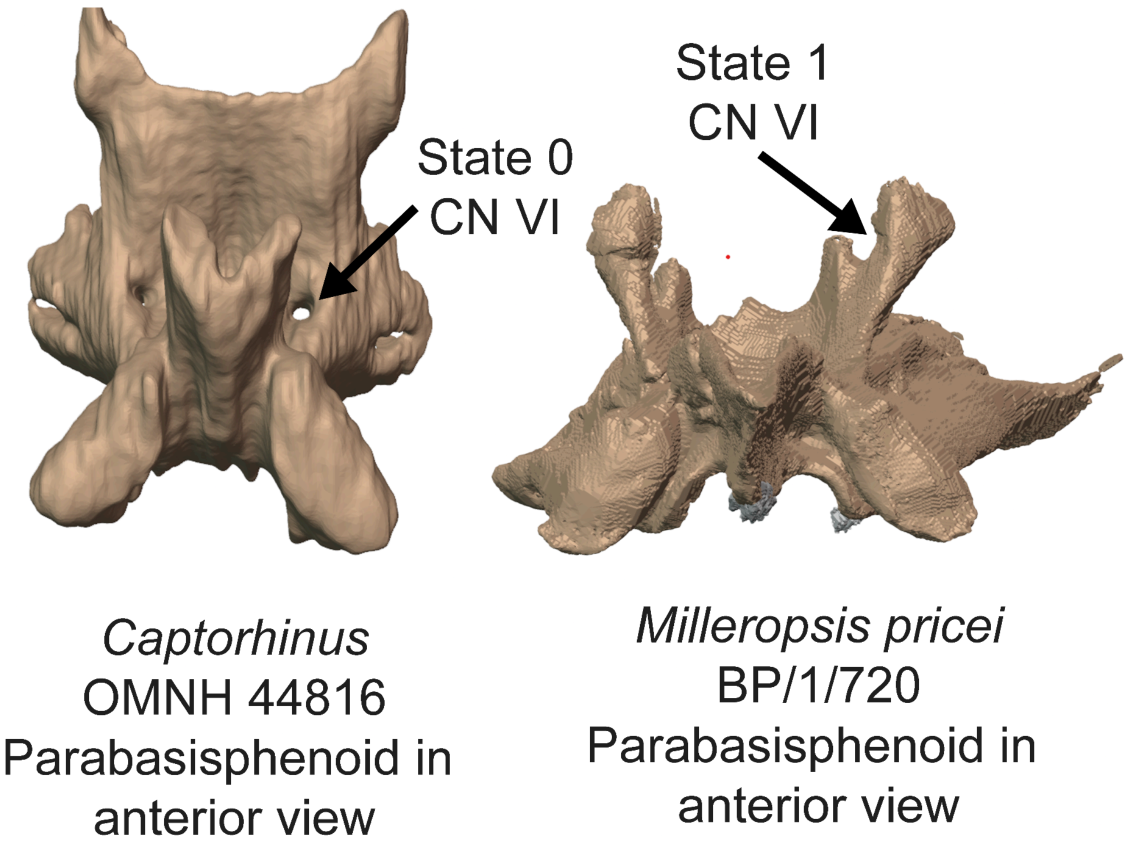


**Fig. S46.** Parabasisphenoids of *Captorhinus* (OMNH 44816) and *Milleropsis* (BP/1/720, this study) in anterior view demonstrating the position of CN V through the dorsum sellae (Ch. 278, State 0) and through the clinoid processes (Ch. 278, State 1), respectively.

In most early or stem amniotes (e.g., *Captorhinus* OMNH 44816) or synapsids such as *Dimetrodon* (Romer and Price 1940 ), or *Heleosaurus* (SAM-PK-K8305, this study), the opening of the abducens nerve (CN VI) pierces the wall of the dorsum sellae (State 0). In most parapleurotan reptiles within this sampling, CN VI is present as a foramen piercing the clinoid processes immediately adjacent to the articulation for the prootic (State 1, *Milleropsis* BP/1/720, *Youngina* BP/1/2871).

DELTRAN: *Planocephalosaurus robinsonae* (1 to 0), Parapleurota( 0 to 1). ACCTRAN: Lepidosauromorpha (1 to 0), Mesosauridae + *Ascendonanus* + *Orovenator* + *Cabarzia* + Parapleurota (0 to 1).

**Ch. 279. Parasphenoid, clinoid processes: present as dorsal pedicels that support the ventral ramus of the prootic (0), absent, prootic is not supported ventrally by parabasisphenoid (1).**

Modified from Pardo et al. (2017, Ch. 248).

The clinoid process of the parabasisphenoids, which form a pedicel for the ventral ramus of the prootic, are absent (State 1) in most early tetrapods (e.g., *Lethiscus*, Pardo et al. 2017) within this sampling but are present (State 0) in most recumbirostrans and amniotes.

DELTRAN: Recumbirostra (1 to 0), *Seymouria* spp. (0 to 1). ACCTRAN: Gymnarthridae + *Nannaroter* + Rhynchonkidae + *Huskerpeton* (1to 0), *Seymouria* spp. (0 to 1)

**Ch. 280. Posterior median braincase ossification or "subiculum infundibulum" of ethmoid trabeculae absent (0), present (1).**

This study.

An ossification of the subiculum infundibulum, a median ossification located anteriorly to the basipterygoid region, is present in gymnarthrid (e.g., *Euryodus* FM UR 2296) and ostodelepid (e.g., *Rhynchonkos,* FM-UR 1039) recumbirostrans.

DELTRAN: Rhynchonkidae + *Huskerpeton* (0 to 1), Gymnarthridae + *Nannaroter* + Rhynchonkidae + *Huskerpeton* (0 to 1).

**Ch. 281. Ossification between optic foramen (CN II) and pila antotica: pila metoptica and associated cartilaginous taeniae unossified (0), ossification complete between optic foramen and pila antotica (1).**

Modified from Pardo et al. (2017, Ch. 188).

The presence of ossification between the pila antotica and CN II (State 1), is present in many early tetrapods including embolomeres (e.g., *Archeria*, Clack and Holmes 1987), temnospondyls (e.g., *Edops* MCZ 1378) and some recumbirostrans (e.g., *Dvellecanus* UCMP 202940).

DELTRAN: Gymnarthridae + *Nannaroter* (0 to 1), *Eothyris parkeyi* (0 to 1), *Bunostegos akokanensis* (0 to 1), *Embrithosaurus schwarzi* (0 to 1), *Azendohsaurus madagaskarensis* (0 to 1), *Archeria crassidiscus* (0 to 1). ACCTRAN: *Dvellecanus carrolli* (0 to 1), Gymnarthridae + *Nannaroter* (0 to 1), *Thuringothyris* + Captorhinidae (0 to 1), *Deltavjatia rossica* (1 to 0), Pareiasauromorpha (0 to 1), *Trilophosaurus* + *Mesosuchus* (1 to 0), *Archeria crassidiscus* (0 to 1).

**Ch. 282. Ossification within columella ethmoidalis: absent or unossified (0), present and ossified (1).**

Modified from Pardo et al. (2017, Ch. 189).

An ossification within the columella ethmoidalis, or nasal septum region, is present (State 1) in chthonosaurian recumbirostrans (e.g., *Brachydectes* KUVP 49541).

UNAMBIGUOUS: *Captorhinus laticeps* (0 to 1), Chthonosauria (0 to 1), DELTRAN: *Archeria crassidiscus* (0 to 1).

**Ch. 283. Ossification of antorbital region dorsal to cultriform process ("sphenethmoid" or "orbitosphenoid"): unossified (0), ossified (1).**

Modified from deBraga and Rieppel (1997), Mülller (2004, Ch. 138), Huttenlocker et al. (2013, Ch. 113).

An ossification dorsal to the cultriform process, generally referred to as a sphenethmoid, orbitosphenoid, or 'ossified interorbital cartilage' in the literature, is present in mature individuals of most Paleozoic tetrapods (e.g., *Elliotsmithia*, TM 1483) (State 1). There is no evidence of this ossification (State 0) in any early parapleurotan reptile (e.g., *Milleropsis,* BP/1/720).

UNAMBIGUOUS: *Dendrerpeton helogenes* (1 to 0), Temnospondyli + Reptiliomorpha + *Crassigyrinus* + *Greererpeton* + *Baphetes* (0 to 1), Parapleurota (1 to 0). DELTRAN. *Elliotsmithia* (0 to 1).

**Ch 284. Orbitosphenoid or sphenethmoid, if present, medial surface expanded for olfactory bulb: absent (0), present (1) .**

**Modified from Pardo et al. (2017, Ch. 194).**

The medial surface of the orbitosphenoid in non-chthonosaurian recumbirostrans (e.g., *Rhynchonkos* FM-UR 1039) is expanded (and medially concave) for the olfactory bulbs (State 1).

DELTRAN: Rhynchonkidae + *Huskerpeton* (0 to 1), *Pantylus cordatus* (0 to 1). ACCTRAN: *Llistrofus* + Chthonosauria (1 to 0), Recumbirostra (0 to 1).

**Ch 285. Orbitosphenoid or sphenethmoid, if present, processus vomeronasalis: absent (0), present, medial process separates pathway of CN 0 and olfactory tract (1).**

This study.

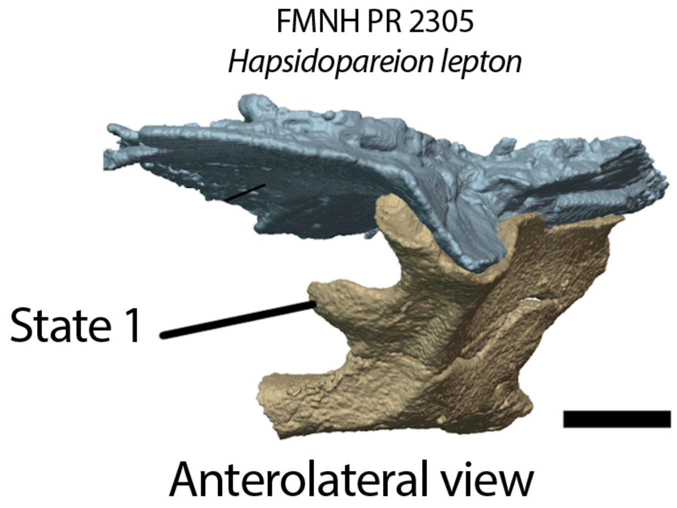


**Fig. S47.** Frontal and orbitosphenoid of *Hapsidopareion* (FMNH PR 2305) in anterolateral view, showing the processus vomeronasalis of the orbitosphenoid (Ch. 285, State 1).

The processus vomeronasalis, an anteromedial process extending on the medial surface of the orbitosphenoid, separating the pathway of the cranial nerve 0 and olfactory tract, is present (State 1) in chthonosaurian recumbirostrans and their stem-taxa (e.g., *Hapsidopareion*, FMNH 2303).

DELTRAN: Chthonosauria (0 to 1), *Hapsidopareion* (0 to 1). ACCTRAN: Gymnarthridae (0 to 1), *Llistrofus* + Chthonosauria (0 to 1).

**Ch. 286. Orbitosphenoid or sphenethmoid, If present, forms interorbital septum: absent (0), present (1). (-) in taxa lacking an ossification of the orbitosphenoid or sphenethmoid (Ch. 271).**

Modified from Pardo et al. (2017, Ch. 200).

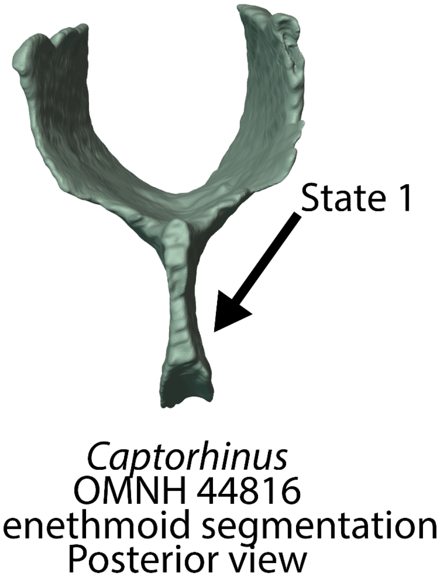


**Fig. S48.** The sphenethmoid of *Captorhinus* (OMNH 44816) in posterior view, bearing an interorbital septum (Ch. 286, State 0).

An interorbital septum (e.g., a y-shaped sphenethmoid as opposed to paired orbitosphenoids) is absent (State 0) in early tetrapods including temnospondyls (e.g., *Dendrerpeton*, NSM 987 GF 99.1) and recumbirostrans (e.g., *Hapsidopareion,* FMNH 2305), but is present in early or near-amniotes (e.g., *Ophiacodon,* Romer and Price 1940) and captorhinids (e.g., *Captorhinus* OMNH 44816). An interorbital septum is also absent in *Saurodektes* (scans of BP/1/1395) and *Procolophon* (Carroll and Lindsay 1985). We note that the interorbital septum is convergently lost in many theriodont synapsids, a group not sampled in this matrix (Benoit et al. 2017).

UNAMBIGUOUS: Ankyramorpha (1 to 0). Procolophonidea (0 to 1). DELTRAN: Recumbirostra + Diadectomorpha + Protorothyrididae + Amniota (0 to 1). ACCTRAN: Recumbirostra (0 to 1).

**Ch. 287. Pleurosphenoid, if present, contact with sphenethmoid or orbitosphenoid: absent, a gap remains unossified between these two elements (0), present, pleurosphenoid-sphenethmoid**

**contacts forms a complete wall in anterior braincase (1). (-) in taxa lacking an ossified pleurosphenoid (Ch. 283).**

Observations noted in Pardo et al. (2017) and Mann et al. (2022).

The pleurosphenoid (see Ch. 260) possesses an anterior development which contacts the orbitosphenoid in gymnarthrid (e.g., *Euryodus* FM UR 2296*)*, ostodolepidid (e.g., *Nannaroter*, OMNH 73107), and rhynchonkid (e.g., *Rynchonkos*, FM-UR 1039) recumbirostrans, but not chthonosaurians (e.g., *Brachydectes*, KUVP 49541). We note here that the presence of this contact does not necessarily mean that ossification is complete between the optic foramen (CN II) and pila antotica, observable in *Rhynchonkos* (FM-UR 1039) where there is still an unossified gap between these two elements dorsally.

DELTRAN: Rhynchonkidae (0 to 1). ACCTRAN: *Huskerpeton englehorni* (1 to 0), Gymnarthridae + *Nannaroter* + Rhynchonkidae + *Huskerpeton* (0 to 1).

**Ch. 288. Epipterygoid, contribution to basal articulation: present, epipterygoid participates in the basipterygoid recess of Parabasisphenoid (0), absent, does not contribute to basipterygoid recess of parabasisphenoid (1).**

Modified from Pardo et al. (2017, Ch. 212).

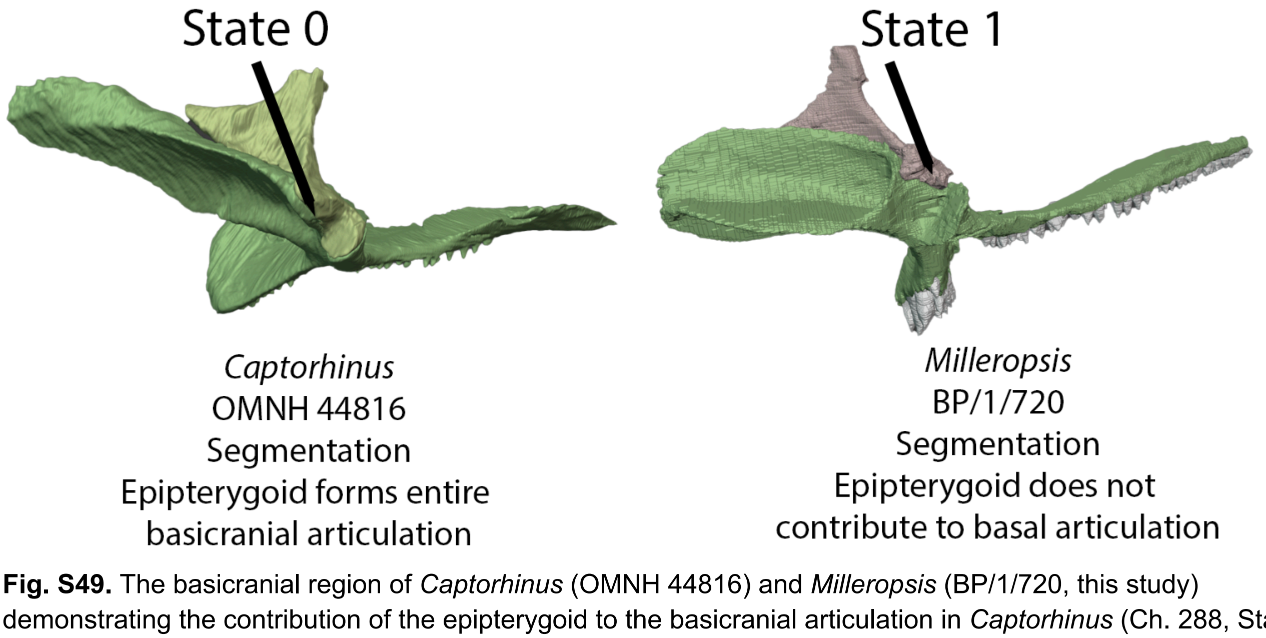


**Fig. S49.** The basicranial region of *Captorhinus* (OMNH 44816) and *Milleropsis* (BP/1/720, this study) demonstrating the contribution of the epipterygoid to the basicranial articulation in *Captorhinus* (Ch. 288, State 0) and the absence of this contribution in *Milleropsis* (Ch. 288, State 1).

The epipterygoid contributes to the basipterygoid recess, conus recessus, or basal articulation in most paleozoic tetrapods including temnospondyls (e.g., *Edops* MCZ 1378), captorhinids (e.g., *Captorhinus laticeps* OMNH 44816), synapsids (e.g., *Ophiacodon* or *Varanops*, Romer and Price 1940), and araeoscelidians (e.g., *Petrolacosaurus*, Reisz 1981). In contrast, the pterygoid forms the entire basipterygoid recess in parapleurotan reptiles including millerettids (e.g., *Milleropsis*, BP/1/720), and neodiapsids (e.g., *Youngina* BP/1/2871).

UNAMBIGUOUS: Parapleurota+ (*Orovenator* + *Ascendonanus* + *Cabarzia*) + Mesosauridae.

(0 to 1). DELTRAN: Ankyramorpha (1 to 0). ACCTRAN: *Deltavjatia* + *Bunostegos* (0 to 1).

**Ch. 289. Epipterygoid, if the contribution to basal articulation present, extent of contribution: epipterygoid forms the entire conus recessus (0), pterygoid also contributes to conus recessus (1). (-) in taxa lacking an epipterygoid contact with basicranial recess (Ch. 288).**

Modified from Pardo et al. (2017, Ch. 212).

When the epipterygoid contributes to the basicranial recess in non-parapleurotan amniotes, the pterygoid also contributes (State 1) to this recess in most amniotes within this matrix such as diadectomorphs (e.g., *Limnoscelis* YPM 811), synapsids (e.g., *Haptodus*, ROM 43604) including reptiles such as acleistorhinids (e.g., *Delorhynchus,* OMNH 73515) and araeoscelidians (e.g., *Petrolacosaurus,* Reisz 1981). In early tetrapods (e.g., *Archeria* Holmes 1989), temnospondyls (e.g., *Mastodonsaurus*, Schoch 2002) and captorhinids such as *Captorhinus* (OMNH 44816) and *Romeria* (Clark and Carroll 1972), the epipterygoid is the sole contributor to the basal articulation or conus recessus (State 0).

DELTRAN: *Hapsidopareion* (1 to 0), Captorhinidae (1 to 0), Recumbirostra + Captorhinidae + Protorothyrididae + Diadectomorpha (0 to 1). ACCTRAN: *Hapsidopareion* (1 to 0), Captorhinidae (1 to 0), Reptiliomorpha (0 to 1).

**Ch. 290. Epipterygoid, medial process or lip roofing basal articulation: epipterygoid possesses a 'medial process' that 'roofs' the basal articulation but does not contribute to it (0), epipterygoid lacks a medial process (1). (-) in taxa in which the epipterygoid contributes to the basicranial recess (Ch. 288).**

This study.

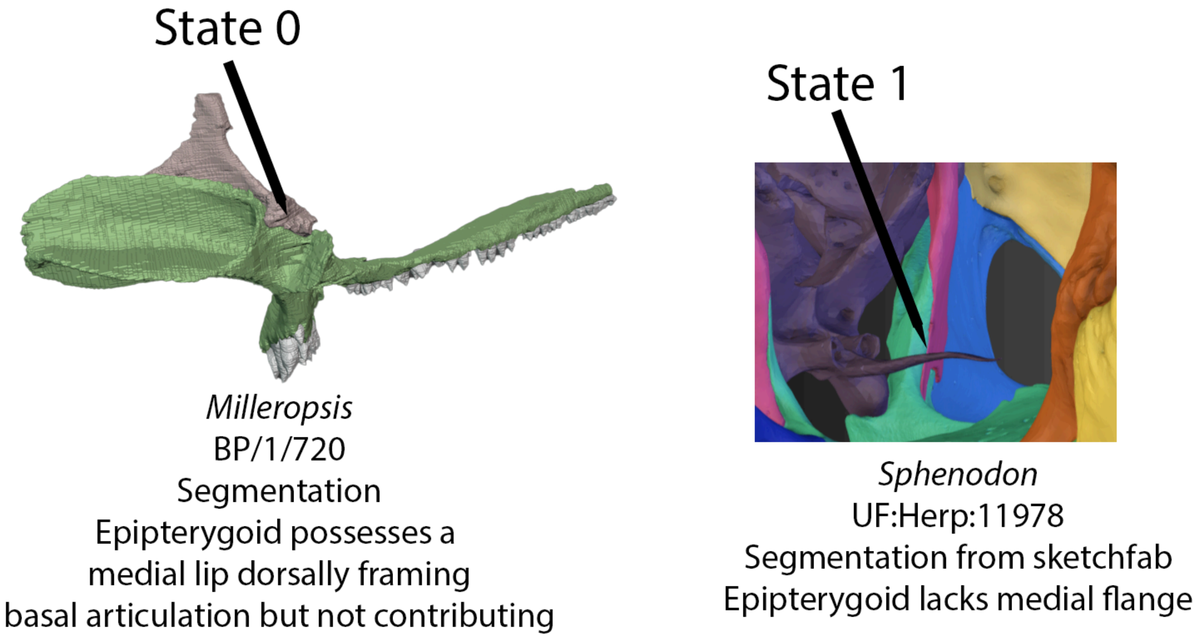


**Fig. S50.** Left, basicranial region of *Milleropsis* in posteromedial view demonstrating the presence of a medial lip of the epipterygoid (Ch. 290, State 0) and right, basicranial region of *Sphenodon* in anterior view with an epipterygoid lacking a medial lip roofing the basal articulation (Ch. 290, State 1).

In reptiles that lack a contribution of the epipterygoid to the basicranial recess, the epipterygoid often possesses a medial process or lip that 'roofs' the basal articulation (Gow 1975). In lepidosauromorphs (e.g., *Clevosaurus*, NHMUK PV R36832), the base of the epipterygoid is located far posterior to the basal articulation, and lacks a medial development roofing this region dorsally (State 1).

UNAMBIGUOUS: Lepidosauromorpha (0 to 1).

**Ch. 291. Epipterygoid, footplate, contact with quadrate: absent, footplate rests solely on pterygoid (0), present, footplate rests on pterygoid and quadrate (1).**

This study.

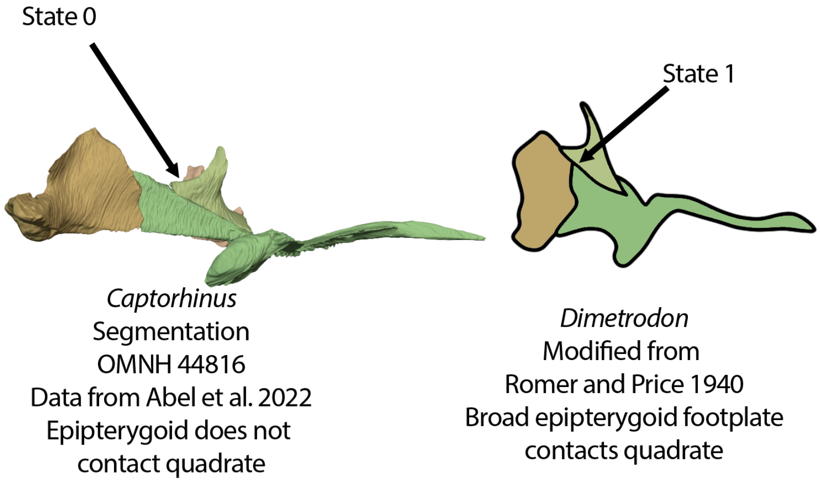


**Fig. S51.** Left, palatoquadrate of *Captorhinus* (OMNH 44816) which bears an epipterygoid that does not contact the quadrate (Ch. 291, State 0). Right, diagram of the palatoquadrate of *Dimetrodon* (modified from Romer and Price 1940) which bears a broad epipterygoid footplate that contacts the quadrate posteriorly (Ch. 291, State 1).

The base or footplate of the epipterygoid contacts the pterygoid ramus of the quadrate in many Paleozoic tetrapods (e.g., *Greererpeton,* CMNH 11234) and pelycosaurs crownward than Caseidae (e.g., Ophiacodon [Romer and Price 1940] or *Mycterosaurus* [FMNH-UC 692]). In contrast, the footplate of the epipterygoid is restricted to the quadrate ramus of the pterygoid in captorhinids (e.g., *Captorhinus* OMNH 44816, recumbirostrans (e.g., *Rhynchonkos*, FM-UR 1039), and early caseasaurs (e.g., *Eothyris*, MCZ 1161).

DELTRAN: *Acheloma cumminsi* (1 to 0), *Cotylorhynchus romeri* (0 to 1), Synapsida (0 to 1) *Clevosaurus hudsoni* (0 to 1), *Tseajaia* + *Orobates* (1 to 0), *Seymouria* spp. (1 to 0). ACCTRAN: *Cotylorhynchus* + *Angelosaurus* (0 to 1) Eupelycosauria + Varanopidae (0 to 1), Rhynchocephalia (0 to 1), Diadectomorpha (0 to 1) Temnospondyli + Reptiliomorpha + *Gephyrostegus* (1 to 0).

**Ch. 292. Epipterygoid, dorsal columella, distal end: unexpanded and rod-like (0), expanded and plate-like (1).**

Modified from Rieppel (2011,Ch. 39) and Gauthier (2012, Ch. 283).

The dorsal columella of most paleozoic tetrapods is rod-like, approaching but not making osseous contact with the skull roof (State 0). In some early tetrapodomorphs (e.g., *Whatcheeria* FM PR 1634 or *Greererpeton*, CMNH 11068) and leptopleuronine procolophonids (e.g., *Leptopleuron,* Spencer 2000), the distal end of the dorsal columella is expanded distally and rectilinear in lateral view (State 1).

UNAMBIGUOUS: *Tetraceratops insignis* (0 to 1), Leptopleuroninae + *Procolophon* (0 to 1). DELTRAN: *Tanystropheus hydroides* (0 to 1), Temnospondyli + Reptiliomorpha (1 to 0). ACCTRAN: Temnospondyli + Reptiliomorpha + *Gephyrostegus* (1 to 0).

**Ch. 293. Epipterygoid, columella, shaft: rodlike and circular in cross section (0), anteroposteriorly expanded and sheet-like in cross-section (1).**

**This study.**

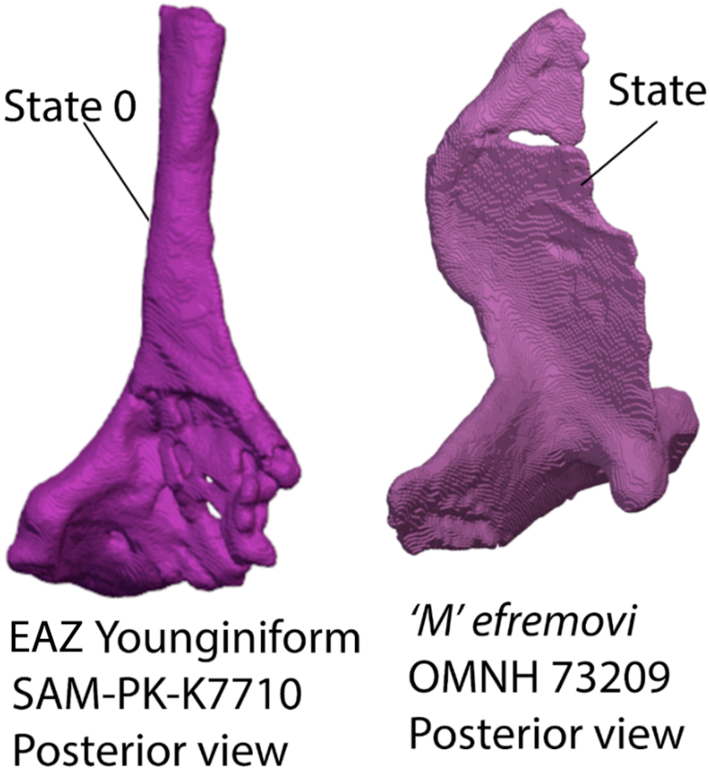


**Fig. S52.** Left, epipterygoid of SAM-PK-7710 in posterior view, which bears a subcircular dorsal columella (Ch. 293, State 0). Right, epipterygoid of *Mesenosaurus efremovi* (OMNH 73209) demonstrating the derived condition (Ch. 293, State 1) bearing a sheetlike dorsal columella of the epipterygoid.

The epipterygoids of most early tetrapods bear a subcircular dorsal columella (e.g., *Edops,* Romer and Witter 1942). In contrast, the dorsal columella of the epipterygoid of many synapsids including the varanopid *Heleosaurus* (this study), the edaphosaurid *Gordodon* (Lucas et al. 2018), and *Dimetrodon milleri* (Brink and Reisz 2021) is sheet-like in cross-section.

**Ch. 294. Epipterygoid, posterior ramus contacting squamosal: absent (0), present (1).**

This study.

The footplate of early tetrapods (e.g., *Greererpeton* or *Edops*) possesses a posterior expansion or ramus which contacts the squamosal posterodorsally which is separate from the dorsal columella

(State 1). This process is absent in all known early reptiliomorphs within this matrix barring embolomeres (State 0).

UNAMBIGUOUS: Reptiliomorpha (0 to 1).

**Ch. 295. Braincase, basicranial fissure between sphenoid and otoccipital elements: absent (0), present (1)**

Modified from Pardo et al. (2017, Ch. 210).

The presence of a basicranial fissure between the basisphenoid and otoccipital region is plesiomorphic in limbed vertebrates (Ahlberg, 1991). Within this taxonomic sampling, the basicranial fissure is present (State 1) in early tetrapodomorphs such as *Whatcheeria* (FM PR 1701). The basicranial fissure is absent in all reptiliomorphs and temnospondyls within this sampling (State 0).

UNAMBIGUOUS: Temnospondyli + Reptiliomorpha + *Gephyrostegus* (0 to 1).

**Ch. 296. Vomer, teeth or shagreen: present (0), absent (1).**

Modified from Brocklehurst et al. (2022, Ch. 132).

**Ch. 297. Vomer, fangs: present (0), absent (1).**

Modified from Brocklehurst et al. (2022, Ch. 133).

**Ch. 298. Vomer, asymmetric bifurcated anterior processes: present, vomer terminates anteriorly in short lateral and elongate medial projections (0), absent (1).**

Modified from Brocklehurst et al. (2022, Ch. 134).

**Ch. 299. Vomer, internarial shape: widest posteriorly (0), widest near middle (1).**

Modified from Brocklehurst et al. (2022, Ch. 317).

**Ch. 300. Vomer, medial contact with pterygoid: absent (0), present (1).**

Modified from Brocklehurst et al. (2022, Ch. 135).

**Ch. 301. Vomer, if medial contact with pterygoid is present it is: extensive ~50% medial border of vomer (0), short (~33% medial border of vomer) (1).**

Modified from Brocklehurst et al. (2022, Ch. 136).

**Ch. 302. Vomer, lateral (alar) flange: absent (0), present, rounded lateral process partially divides choana (1).**

Modified from Tsuji (2006, Ch. 50), MacDougall et al. (2019, Ch. 64, Cisneros et al. (2020, Ch. 62).

The alar flange of the vomer is a lateral process that partially divides the choana giving it a roughly 'kidney-bean' shaped in dorsal or lateral view. The alar flange is present (State 1) in pareiasauromorphs (e.g., *Macroleter,* Tsuji 2006) and the procolophonian *Nyctiphruretus* (PIN 4659/1). DELTRAN: Pareiasauromorpha + *Australothyris* (0 to 1), *Nyctiphruretus acudens* (0 to 1). ACCTRAN: Procolophonoidea (1 to 0), Ankyramorpha (0 to 1).

**Ch. 303. Vomer, medial tooth row: discrete row present (0), teeth absent in this region or shagreen present (1). (-) in taxa bearing a denticle field or shagreen on ventral surface of vomer.**

Modified from Dilkes (1998), Pritchard et al. (2015, Ch. 45), Ezcurra (2016 Ch. 187).

The medial tooth row of the vomer ('V3' of Hunt et al. 2023) is a tooth row located on the medial surface of the vomer along the midline (e.g., *Youngina capensis*, BP/1/2871). This is present in most Paleozoic tetrapods bearing discrete rows of vomerine teeth, but not all (e.g., *Tanystropheus hydroides,* PIMUZ T 2790).

DELTRAN: Hapsidopareion (0 to 1), *Moradisaurus grandis* (0 to 1), *Planocephalosaurus robinsonae* (0 to 1), *Tanystropheus hydroides* (0 to 1), *Hylonomus lyelli* (0 to 1), Diadectomorpha + Protorothyrididae + Amniota + Recumbirostra (1 to 0). ACCTRAN: Captorhinidae (0 to 1), Amniota + Protorothyrididae + Diadectomorpha + Recumbirostra + Captorhinidae (1 to 0).

**Ch. 304. Vomer, lateral (choanal) tooth rows: absent (0), discrete row (s) present (1). (-) in taxa bearing a denticle field or shagreen on ventral surface of vomer.**

Modified from Dilkes (1998), Pritchard et al. (2015, Ch. 45), Ezcurra (2016 Ch. 187).

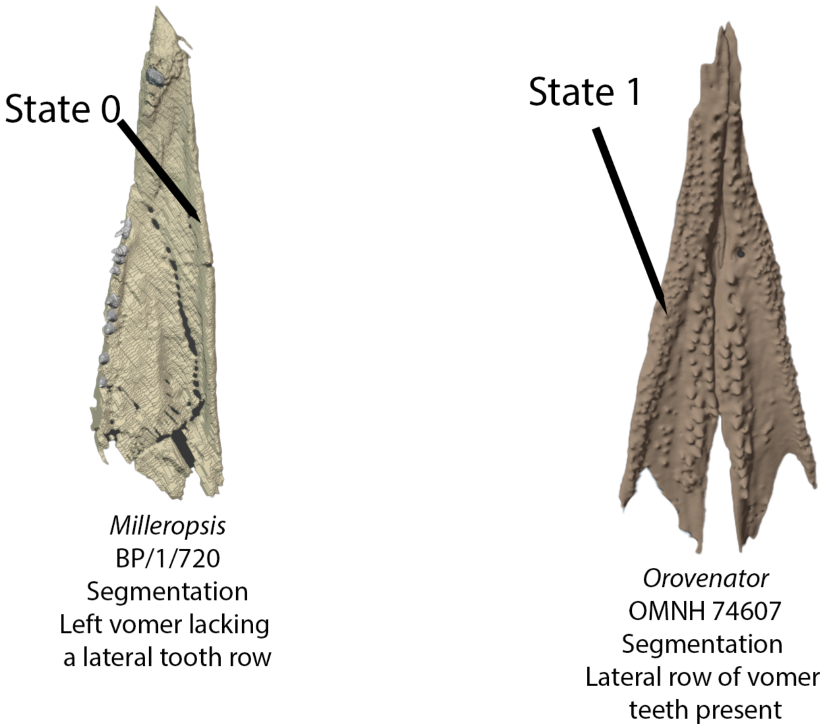


**Fig. S53.** Left, left vomer of *Milleropsis* (BP/1/720, this study) in ventral view demonstrating the absence of a lateral vomerine tooth row (Ch. 304, State 0). Right, vomers of *Orovenator* (OMNH 74607) in ventral view demonstrating the presence of a lateral tooth row (Ch. 304, State 1).

The lateral or choanal tooth rows of the vomer are widespread in early amniotes, particularly reptiles (e.g., *Youngina* BP/1/2871), and are laterally-positioned rows of teeth that often border the choanal margin and can be an anterior continuation of a row of palatine teeth (State 1). This row(s) are absent (State 0) in millerettids (e.g., *Milleropsis* BP/1/720), which only possess a midline row of vomerine teeth.

UNAMBIGUOUS: *Pantylus cordatus* (0 to 1), *Euconcordia cunninghami* (0 to 1), Leptopleuroninae + *Procolophon* + *Eomurruna* (1 to 0), Amniota + Protorothyrididae + *Anthracodromeu*s (0 to 1).
DELTRAN: Millerettidae (1 to 0) Squamata (1 to 0).

**Ch. 305. Vomer, ventral surface: ventral surface flat and unexpanded (0), anterior vomerine buttress bearing enlarged vomerine teeth (1), ventral crests forming midline trough (2).**

This study.

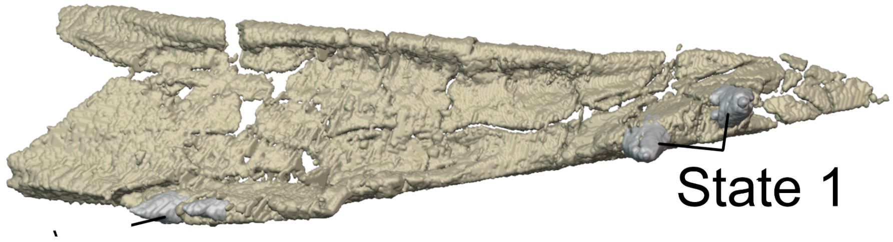

**Fig. S54.** Left vomer of *Milleretta* (BP/1/3822, this study) in ventral view demonstrating the presence of enlarged vomerine teeth supported by a vomerine buttress (Ch. 305, State 1).

The vomer of most amniotes within this sampling is a sheet-like element, with little to no changes in dorsoventral thickness throughout the element (State 0). Millerettid reptiles (e.g., *Milleretta,* BP/1/3822) possess a thickened, vomerine buttress supporting the anteriormost vomerine teeth, which are enlarged relative to the rest of the vomerine tooth row (State 1). This condition differs from the 'vomerine fangs' of early tetrapods in that they are congruent with the medial vomerine tooth row and much smaller than the marginal dentition. The ventral surface of the vomer of many sphenacodontians (e.g., *Dimetrodon*, Romer and Price 1940) and therapsids (e.g., *Titanophoneus*, Efremov 1938) possess a medially-placed midline crest that forms a midline trough with the contralateral vomer (State 2).

UNAMBIGUOUS: *Acheloma* + *Cacops* (0 to 1). DELTRAN: Therapsida (0 to 2). ACCTRAN: Sphenacodontoidea + *Pantelosaurus* (0 to 2). ACCTRAN: Millerettidae (0 to 1).

**Ch. 306. Vomer, ascending flange "dorsolateral process" framing choana: absent (0), present, tall ascending flange of vomer forms medial border of choana (1).**

This study.

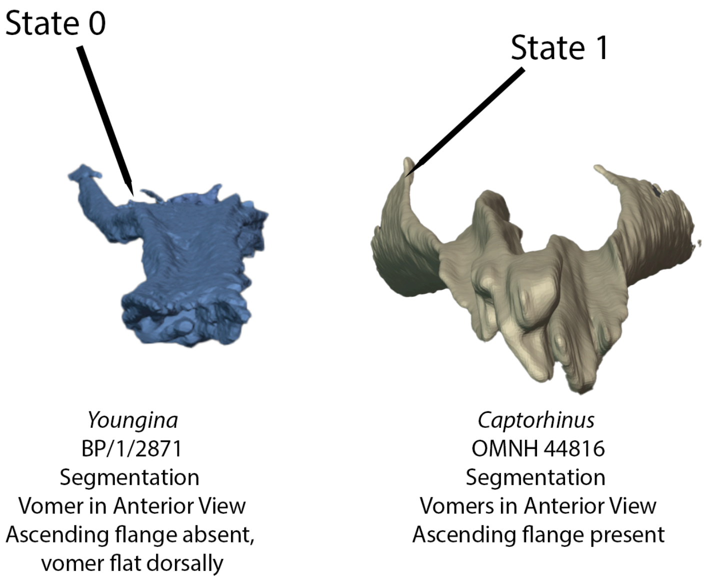


**Fig. S55.** Left, right vomer of *Youngina* (BP/1/2871) in anterior view demonstrating the absence of an ascending flange of the vomer (Ch. 306, State 0) and right, vomers of *Captorhinus* (OMNH 44816) in anterior view bearing ascending lateral flanges of the vomer (Ch. 306, State 1).

The vomer of many early tetrapods (e.g., *Acheloma*, OMNH 79318), recumbirostrans (e.g., *Hapsidopareion* FMNH 2303), and captorhinids (e.g., *Captorhinus*, OMNH 44816) possess an ascending flange which frames the medial border of the choana (State 1), whereas the vomer of most crown amniotes lacks it (e.g., *Milleropsis*, BP/1/720 or *Youngina*, BP/1/2871). This ascending flange possibly delimited the lateral margin of the paraseptal cartilage.

DELTRAN: *Raranimus dashankouensis* (0 to 1), Protorothyrididae (1 to 0), Recumbirostra + Diadectomorpha + Protorothyrididae + Amniota (0 to 1). ACCTRAN: Reptiliomorpha (0 to 1).

**Ch. 307. Palatine, teeth or shagreen: present (0), absent (1).**

Modified from Brocklehurst et al. (2022, Ch. 137).

**Ch. 308. Palatine, fangs: present (0), absent (1).**

Modified from Brocklehurst et al. (2022, Ch. 138).

**Ch. 309. Palatine, incision of anterior surface to accommodate internal naris ( choana): shallow to moderate anterior incision (0), deep anterior incision (1), palatine is not incised, or only a small notch is present (2).**

Modified from Brocklehurst et al. (2022, Ch. 139).

**Ch. 310. Palatine, posterior process, contact with ectopterygoid: present, contacts ectopterygoid (0), reduced or absent, no contact with ectopterygoid (1). (-) in taxa with an absent ectopterygoid (Ch. 336).**

Modified from Gauthier (1988, Ch. 39), Hugo-Reynoso (1998 Ch. 156).

The posterior process of the palatine contacts the ectopterygoid (State 0) in most Paleozoic tetrapods within this matrix (e.g., *Limnoscelis* YPM 811). However, the palatine does not contact the ectopterygoid (State 1) in archosauromorphs, which possess an extremely enlarged suborbital fenestra (e.g., *Macrocnemus,* PIMUZ T 2477).

UNAMBIGUOUS: Archosauromorpha (0 to 1), *Araeoscelis* (0 to 1).

**Ch. 311. Palatine, ventral surface, pocket or fossae: absent, ventral surface of palatine flat and sheetlike (0), present, palatine possesses a deep pocket or emargination medial to maxillary ramus (1).**

This study.

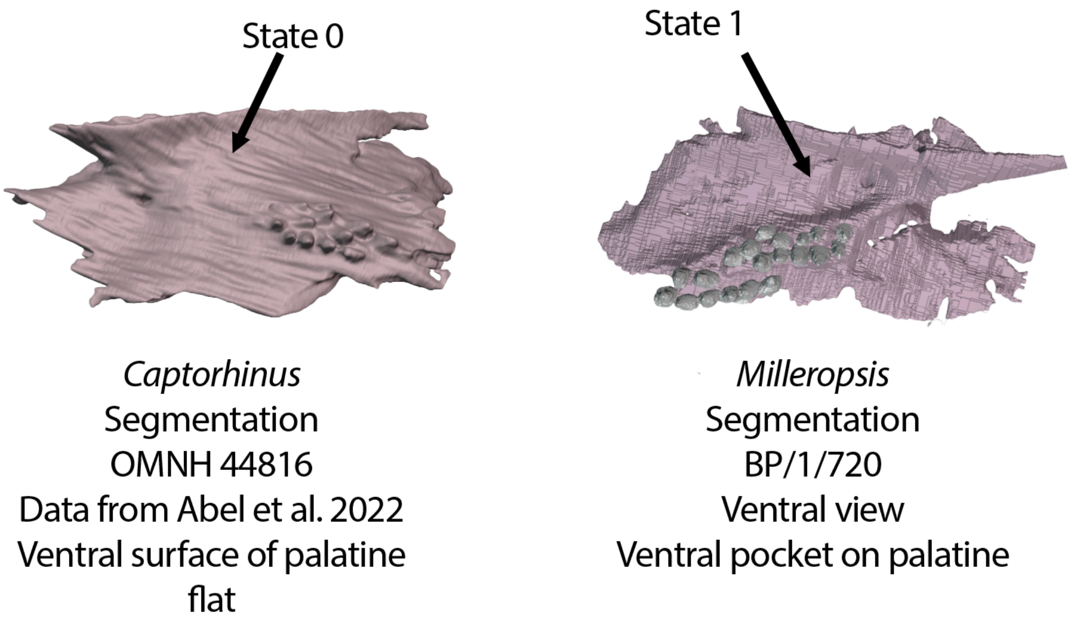


**Fig. S56.** Left palatines of *Captorhinus* (OMNH 44816) and *Milleropsis* (BP/1/720) demonstrating the absence (Ch. 311, State 0) and presence of a ventral pocket on the palatine (Ch. 311, State 1) respectively.

The ventral surface of the palatine is flat (State 0) in most early amniotes (e.g., *Captorhinus* OMNH 44816). In contrast, the palatine of millerettid reptiles (e.g., *Milleretta,* BP/1/3822) and many neodiapsids (e.g., *Youngina* BP/1/2871 or *Clevosaurus*, NHMUK PV R36832) possesses (State 1) a distinct concavity or pocket medial to the maxillary ramus.

DELTRAN: Squamata (0 to 1), Tanystropheidae + Crocopoda (0 to 1), Tangasauridae (1 to 0), Parapleurota + *Orovenator* + *Ascendonanus* + *Cabarzia* (0 to 1) *Orobates pabsti* (0 to 1). ACCTRAN: Rhynchocephalia (0 to 1), Crocopoda (0 to 1), Weigeltisauridae + Tangasauridae + *Claudiosaurus* + *Acerosodontosaurus* + Sauria (1 to 0), *Orobates pabsti* (0 to 1).

**Ch. 312. Palatine, dorsal surface, ascending process: absent or weakly developed, dorsal surface of palatine is flat or weakly convex (0), strongly developed, dorsal surface of palatine possesses an anterodorsally-oriented ascending process or ridge (1).**

Modified from Brocklehurst et al. (2022, Ch. 59).

**Ch. 313. Pterygoid, transverse flange: large, approaches cheek, a noticeable lateral projection (0), small, does not approach cheek (1).**

Modified from Brocklehurst et al. (2022, Ch. 140).

**Ch. 314. Pterygoid, teeth on transverse flange (=T1): present (0), absent (1).**

Modified from Brocklehurst et al. (2022, Ch. 141).

**Ch. 315. Pterygoid, arrangement of teeth on transverse flange: a field or shagreen of teeth covering the ventral surface of the flange (0), additional teeth anterior to a single row of large teeth on ventral rim of flange (1), single row of teeth on ventral rim of flange (2). ORDERED.**

Modified from Brocklehurst et al. (2022, Ch. 142).

**Ch. 316. Pterygoid, orientation of the transverse flange: directed predominately laterally or posterolaterally (0), directed anterolaterally (1), directed anteriorly (2).ORDERED.**

Modified from Brocklehurst et al. (2022, Ch. 143).

**Ch. 317. Pterygoid, teeth in a distinct, anterolaterally oriented field (=T2): pterygoid heavily denticulated, no discrete field present (0), anterolaterally oriented discrete field/row of teeth present (1), teeth and denticles reduced or absent in this region, no discrete row present (2). ORDERED.**

Reworked from Brocklehurst et al. (2022, Ch. 318).

Rationale: Added the T2 specifier used by Welman (1998) for consistency with other early reptile work.

**Ch. 318. Pterygoid, palatal ramus, tooth row "T3": present, tooth row "t3" extends along palatal ramus (0), discrete field or tooth tow "t3" absent (1).**

This study.

Rationale: Added the T3 specifier used by Welman (1998) for consistency with other early reptile work.

**Ch. 319. Pterygoid, a row of fang-like teeth on the medial edge of the anterior ramus (=palatal process) (T4 of Welman, 1998): absent (0), present (1).**

Modified from Ezcurra (2016) Ch. 199.

An additional row of pterygoid dentition along the medial surface of the palatal ramus is present in several archosauromorphs within this sampling including *Prolacerta* and *Teyujagua* (Pinheiro et al. 2020).

UNAMBIGUOUS: *Prolacerta* (0 to 1), *Teyujagua* (0 to 1).

**Ch. 320. Pterygoid, anterior extent of palatal process: terminates anteriorly level with or posterior to the anterior margin of the palatine (0), extends anterior to the palatine (1).**

Modified from Brocklehurst et al. (2022, Ch. 144).

**Ch. 321. Pterygoids, median contact between each other: no median contact between the pterygoids (0), median contact present only anteriorly (1), median contact extensive, pterygoids meet medially along > 50% of their anteroposterior length (2). ORDERED.**

Modified from Brocklehurst et al. (2022, Ch. 145).

**Ch. 322. Pterygoid, quadrate ramus, medial shelf or "arcuate/tympanic" flange: absent (0), present, morphology: little more than a medially developed keep or thickening, projects weaker than a 60 degree angle relative to dorsal process (1), a broad, transversely developed platform extending medially more than 70 degrees relative to dorsal process of quadrate ramus (2). ORDERED.**

Modified from Brocklehurst et al. (2022, Ch. 147).

**Ch. 323. Pterygoid, teeth on the quadrate flange: absent (0), present (1).**

Modified from Brocklehurst et al. (2022, Ch. 148).

**Ch. 324. Pterygoid, ascending dorsal lamina of the anterior ( palatal) ramus: absent or low (0), tall (1).**

Modified from Brocklehurst et al. (2022, Ch. 149).

**Ch. 325. Pterygoid, interpterygoid vacuities: absent, palate closed (0), present (1).**

Modified from Ruta et al. (2003, Ch. 153), Ruta and Coates (2007, Ch. 160), Huttenlocker et al. (2015, Ch. 115), and Marjanovic and Laurin (2019, Ch. 105).

UNAMBIGUOUS: *Greererpeton* + Temnospondyli + Reptiliomorpha (0 to 1).

**Ch. 326. Pterygoid, anterior extent of interpterygoid vacuity: reaches beyond posterior margin of palatine (0) reaches level of palatine or less (1).**

Modified from Brocklehurst et al. (2022, Ch. 150).

**Ch. 327. Pterygoid, mediolateral width of interpterygoid vacuity: narrow, mediolateral width of interpterygoid vacuity smaller than or equal to maximum width between basipterygoid processes (0) enlarged, mediolateral width of interpterygoid vacuity wider than maximum mediolateral width of basipterygoid processes (1). (-) in taxa lacking an interpterygoid vacuity (Ch. 325).**

Modified from Clack et al. (2012, Ch. 89).

The interpterygoid vacuity in most early tetrapods is mediolaterally thin, generally similar to the width of the Parabasisphenoid at the level of the basipterygoid processes (State 0, e.g., *Youngina* BP/1/2871). In the temnospondyls within this sampling (barring *Edops* MCZ 1378), the interpterygoid vacuity are enlarged, much wider than the width of the parasphenoid and general concave laterally (e.g., *Acheloma*, AMNH FARB 42). An enlarged interpterygoid vacuity has also been reconstructed in the weigeltisaurid *Coelurosauravus* (Buffa et al. 2021).

UNAMBIGUOUS: *Dendrerpeton* + *Balanerpeton* + *Acheloma* + *Cacops* (0 to 1). DELTRAN: *Coelurosauravus elivensis* (0 to 1). ACCTRAN: Weigeltisauridae (0 to 1).

**Ch. 328. Pterygoid, ventral projection of transverse flange: extends ventrally no further than maxillary dentition (0), extends well below non-caniniform maxillary dentition (1).**

Modified from Jenkins et al. (2024, Ch. 209).

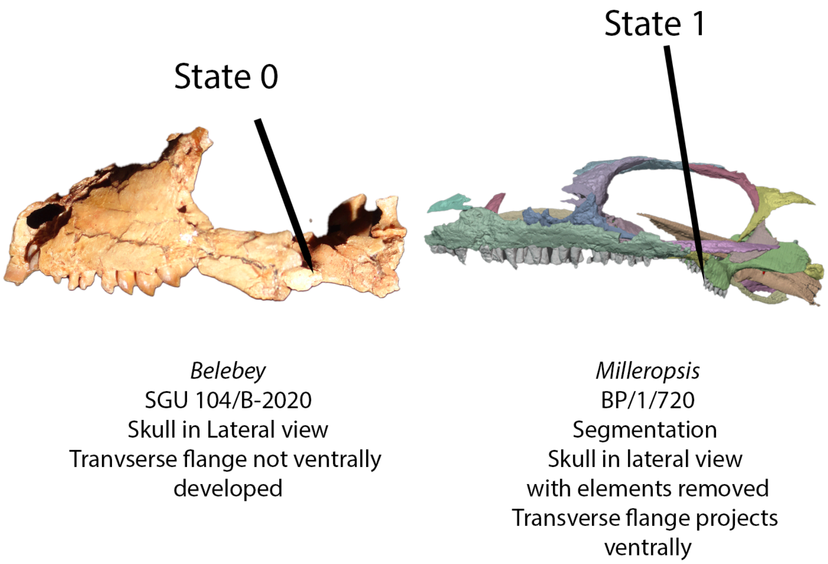


**Fig. S57.** Skulls of *Belebey* (SGU 104/B-2020) and *Milleropsis* (BP/1/720) in lateral view, demonstrating the poor ventral deflection of the transverse flange of the pterygoid (Ch. 328, State 0) and a transverse flange that descends ventrally below the maxillary dentition (Ch. 328, State 1), respectively.

The transverse flange (or similar, transverse development of the pterygoid in taxa which lack a transverse flange) of the pterygoid is flat ventrally, not extending below the maxillary tooth row in most non-amniote taxa, such as early tetrapodomorphs (e.g., *Greererpeton* CMNH 11068), recumbirostrans (e.g., *Carrolla craddocki*, TMM 43001-54), and bolosaurids (e.g., *Belebey*, SGU 104/B-2020) (State 0). In contrast, the transverse flange of the pterygoid in most early amniotes (barring exceptions such as *Mesenosaurus* PIN 3706/11 or the neodiapsids *Claudiosaurus* or *Thadeosaurus* Carroll, 1981) is ventrally developed, extending further ventrally than the maxillary dentition (State 1), and is readily visible in lateral view (e.g., *Youngina*, BP/1/2871).

UNAMBIGUOUS: Bolosauridae (1 to 0). DELTRAN: Captorhinidae (0 to 1), Procolophonia (0 to 1), *Deltavjatia* + *Bunostegos* (1 to 0), Mesenosaurinae (1 to 0), Allokotosauria + *Mesosuchus* (0 to 1), *Prolacerta broomi* (0 to 1), *Claudiosaurus* + Sauria (1 to 0), Mesosauridae + *Ascendonanus* +

*Orovenator* + *Cabarzia* + Parapleurota (0 to 1), Bolosauridae (1 to 0), Diadectomorpha + Protorothyrididae + Amniota (0 to 1). ACCTRAN: *Claudiosaurus* + Sauria (1 to 0), Mesosauridae (1 to 0), Owenettidae (1 to 0).

**Ch. 329. Pterygoid, quadrate ramus, relationship with epipterygoid: epipterygoid sits in a dorsolateral groove or facet (0), quadrate ramus is transversely expanded dorsally, bearing a shelf or platform for the epipterygoid (1), epipterygoid ventrally fused with pterygoid (2).**

This study.

The relationship between quadrate ramus of the pterygoid and the epipterygoid varies strongly in paleozoic amniotes. In most early taxa, such as caseasaurians (e.g., *Casea*, Olson 1968) or araeoscelidians (e.g., *Araeoscelis*, AMNH 4685), the epipterygoid sits on a dorsolaterally-facing facet on the quadrate ramus (State 0). In early procolophonoids (e.g., *Saurodektes*, BP/1/4195) and the Permian synapsid *Edaphosaurus* (Modesto, 1995), the quadrate ramus bears a transverse expansion for the epipterygoid (State 1). State 2, in which the epipterygoid is ventrally fused with the quadrate ramus of the pterygoid is seen in leptopleuronine procolophonids (e.g., *Leptopleuron*, Spencer 2000).

UNAMBIGUOUS: *Greererpeton burkemorani* (0 to 2), Leptopleuroninae (1 to 2), Temnospondyli (0 to 2). DELTRAN: *Edaphosaurus boanerges* (0 to 1), Procolophonoidea (0 to 1). ACCTRAN: Procolophonia (0 to 1).

**Ch. 330. Pterygoid, contact with premaxilla: absent, pterygoid-premaxilla contact prevented by vomer (0), present (1).**

This study.

The pterygoid of the early tetrapodomorph *Whatcheeria* contacts the palatal ramus of the premaxilla.

**Ch. 331. Pterygoid, quadrate ramus, contact with contralateral pterygoid: absent, quadrate ramus does not contact other quadrate ramus posteriorly (0), quadrate rami of pterygoids possess a medial flange that are in contact anterior to the parasphenoid (1).**

Modified from Li et al. (2008, Ch. 72).

The quadrate rami of the pterygoids are separated by the cultriform process of the Parabasisphenoid or interpterygoid vacuity in nearly all tetrapods within this sampling (State 0). In therapsids (e.g., *Biarmosuchus* or *Archaeosyodon*, Chudinov 1960), the quadrate rami of both pterygoids make midline contact (State 1).

UNAMBIGUOUS: Therapsida (0 to 1).

**Ch. 332. Pterygoid, quadrate ramus, medial surface: straight or convex throughout its length (0), deep excavation present on posterolateral surface (1).**

Modified from Laurin and Reisz (1995, Ch. 43) Cisneros et al. (2020, Ch. 71).

The quadrate ramus of the pterygoid is straight or convex laterally (State 0) in most taxa within this sampling (e.g., *Captorhinus,* OMNH 44816). In ankyramorphan reptiles (e.g., *Acleistorhinus*, FM UR 1038), the medial surface of the quadrate rami bears a deep excavation on the posterolateral surface (State 1).

UNAMBIGUOUS: *Greererpeton burkemorani* (0 to 1), *Lanthanosuchus* (0 to 1), *Procolophon trigoniceps* (0 to 1), *Deltavjatia* + *Bunostegos* (0 to 1), *Tanystropheus hydroides* (0 to 1).

**Ch. 333. Pterygoid, quadrate ramus, orientation: primarily in the sagittal plane, directed posteriorly or weakly posterolateral (0), strongly posterolateral such that the posterior margin of the quadrate ramus is oriented primarily transversely (1).**

Modified from Laurin and Reisz (1995) and Cisneros et al. (2020 Ch. 43).

The quadrate ramus of the pterygoid in ankyramorphan reptiles (e.g., *Macroleter,* Tsuji 2006) is oriented posterolaterally, such that the posterior margin of the quadrate ramus is oriented primarily transversely (State 1).

UNAMBIGUOUS: *Lanthanosuchus* (0 to 1), Pareiasauria (1 to 0), *Australothyris* (1 to 0), DELTRAN: Ankyramorpha + *Erpetonyx* (0 to 1).

**Ch. 334. Pterygoid, quadrate ramus, descending flange:** absent (0), present, quadrate ramus possesses a descending flange that extends ventrally below the cheek (1).

The presence of a descending flange of the quadrate ramus is a autapomorphy of embolomeres within this study (State 1).

UNAMBIGUOUS: Embolomeri (0 → 1).

**Ch. 335. Ectopterygoid: present (0), absent (1).**

Reworked from Brocklehurst et al. (2022, Ch. 125).

**Ch. 336. Ectopterygoid, dentition: present (0), absent (1).**

Modified from Brocklehurst et al. (2022, Ch. 151).

**Ch. 337. Ectopterygoid, contact with jugal: absent (0), present (1).**

Modified from Brocklehurst et al. (2022, Ch. 152).

**Ch. 338. Ectopterygoid, anteroposterior length: as long as half the length of the palatines or longer (0), shorter than half the length of the palatines (1). (-) in taxa with an absent ectopterygoid (Ch. 335).**

Modified from Clack et al. (2012 ch. 3), Pardo et al.(2017, Ch. 260).

The ectopterygoid of early tetrapods is a large element of the palate and nearly the size of the palatine (State 0). In near-amniotes, including some (but not all) recumbirostrans (e.g., *Hapsidopareion*, FMNH 2303) and crown amniotes (e.g., *Ophiacodon*, Romer and Price 1940), the ectopterygoid is anteroposteriorly reduced, less than half the length of the palatines or small (State 1).

DELTRAN: Rhynchonkidae (1 to 0), Recumbirostra + Diadectomorpha + Protorothyrididae + Amniota (0 to 1). ACCTRAN: Gymnarthridae + *Nannaroter* + Rhynchonkidae + *Huskerpeton* (1 to 0).

**Ch. 339. Ectopterygoid, fangs: present (0), absent (1). (-) in taxa with an absent ectopterygoid (Ch. 335)**

Modified from Clack et al. (2012,Ch. 30) and Pardo et al. (2017, Ch. 283).

Ectopterygoid fangs are present (State 0) in most early tetrapodomorphs including embolomeres (e.g., *Proterogyrinus*, Holmes 1984) and temnospondyls (e.g., *Acheloma* OMNH 79318) but are absent (State 1) in reptiliomorphs (e.g., *Seymouria* BMS E3500) within this sampling.

UNAMBIGUOUS: Reptiliomorpha (0 to 1).

**Ch. 340. Ectopterygoid, contribution to subtemporal fenestra: absent (0), present (1). (-) in taxa with an absent ectopterygoid (Ch. 335).**

Clack et al. (2012, Ch. 04) and Pardo et al. (2017, Ch. 261).

The ectopterygoid is excluded (State 0) from the subtemporal fenestra by an anterolateral development of the pterygoids in recumbirostrans (e.g., *Euryodus*, FM UR 2296) and eupelycosaurs (e.g., *Edaphosaurus*, MCZ 1680) within this sampling.

DELTRAN: *Greererpeton burkemorani* (0 to 1), *Baphetes* spp. (0 to 1), Recumbirostra (1 to 0), Therapsida (0 to 1), Synapsida (1 to 0). Dinocephalia + *Biseriden*s (0 to 1), Eupelycosauria + Varanopidae (1 to 0), Recumbirostra (1 to 0), Embolomeri (1 to 0).

**Ch. 341. Ectopterygoid, contribution to transverse flange: Distal to transverse flange, does not contribute to lateral portion of flange (0), makes contact with lateral portion of transverse flange (1). (-) in taxa with an absent ectopterygoid (Ch. 335).**

Modified from deBraga and Reisz (1996, Ch. 33) and Cisneros et al. (2020, Ch. 76).

The ectopterygoid is located distally to the transverse flange in most amniotes within this sampling (e.g., *Milleropsis*, BP/1/720 or *Dimetrodon*, Romer and Price 1940) (State 0). In procolophonoids (e.g., *Saurodektes kitchingorum* BP/1/4195), the ectopterygoid makes contact with the lateral portion of the transverse flange, visible in ventral or lateral view (State 1).

UNAMBIGUOUS: *Lanthanosuchus* (0 to 1), *Emeroleter levis* (0 to 1), Procolophonoidea (0 to 1). DELTRAN: *Acleistorhinus pteroticus* (0 to 1), Squamata (0 to 1), *Trilophosaurus buettneri* (1 to 0), Crocopoda (0 to 1), *Coelurosauravus elivensis* (0 to 1). ACCTRAN: Rhynchocephalia (0 to 1), *Trilophosaurus buettneri* (1 to 0), Allokotosauria + *Mesosuchus*+ Archosauriformes (0 to 1), Weigeltisauridae (0 to 1).

**Ch. 342. Ectopterygoid, contact with lacrimal: absent (0), present (1). (-) in taxa lacking an ectopterygoid (Ch. 336).**

**Modified from Säilä (2010, Ch. 57).**

The ectopterygoid contacts the lacrimal (State 1) in procolophonid ankyramorphs within this sampling (e.g., *Eomurruna* QMF 59501).

UNAMBIGUOUS: Procolophonoidea + *Nyctiphruretus* (0 to 1). DELTRAN: Gymnarthridae (0 to 1), *Rhynchonkos stovalli* (0 to 1). ACCTRAN: Gymnarthridae + *Nannaroter* + Rhynchonkidae + *Huskerpeton* (0 to 1).

**Ch. 343. Ectopterygoid, posterolateral process: absent, lateral margin of ectopterygoid lacks a posterior extension (0), present, process extends posterolaterally from main body of ectopterygoid (1). (-) in taxa with an absent ectopterygoid (Ch. 336).**

This study.

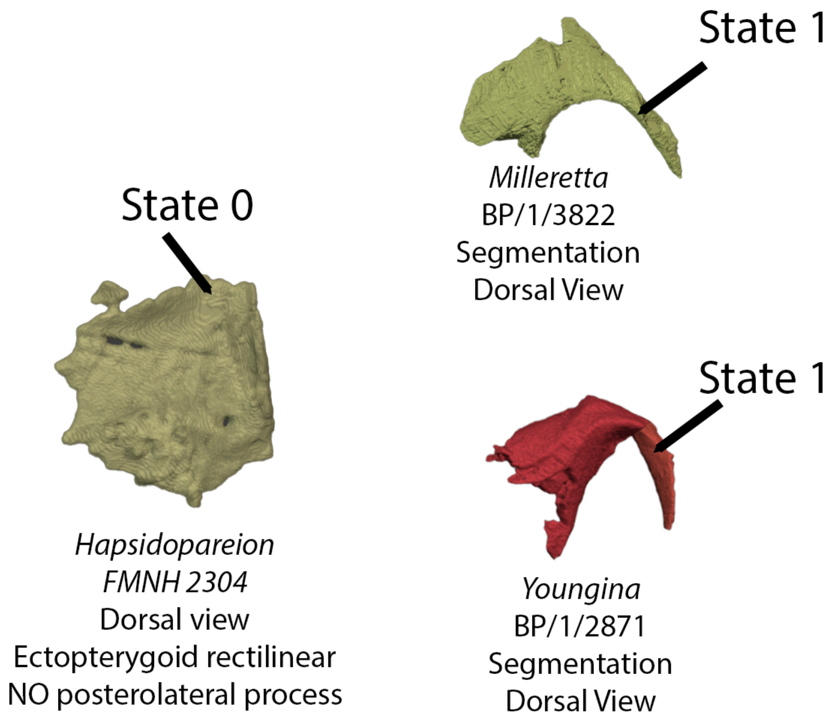


**Fig. S58.** Left, left ectopterygoid of *Hapsidopareion* in dorsal view lacking a posterolateral process (Ch. 343, State 0). Top right (*Milleretta*, BP/1/3822) and bottom right (*Youngina*, BP/1/2871) bearing ectopterygoids with distinct posterolateral processes (Ch. 343, State 1).

The posterolateral process of the ectopterygoid extends posterolaterally from the main body of the ectopterygoid, generally forming the anterior margin of the subtemporal fossa and making contact with the bones of the cheek laterally. This feature is present (State 1) in all known early parapleurotans (e.g., *Youngina* BP/1/2871*)* and *Orovenator* (Ford and Benson 2018), but is absent (State 0) in all other known early amniotes (e.g., *Delorhynchus,* Rowe et al. 2023) in which the ectopterygoid is rectilinear and various early saurians, in which the ectopterygoid becomes strut-like.

UNAMBIGUOUS: Rhynchocephalia (0 to 1), Parapleurota + *Orovenator* + *Ascendonanus* (0 to 1). DELTRAN: Lepidosauria (1 to 0), *Trilophosaurus buettneri* (1 to 0) *Azendohsaurus madagaskarensis* (1 to 0), *Trilophosaurus* + *Mesosuchus* (1 to 0). ACCTRAN: Lepidosauria + *Marmoretta* + *Megachirella* (1 to 0), *Mesosuchus browni* (0 to 1), Allokotosauria + *Mesosuchus* (1 to 0), Tanystropheidae (1 to 0).

**Ch. 344. Suborbital opening: absent (0), present (1).**

Modified from Brocklehurst et al. (2022, Ch. 153).

**Ch. 345. Suborbital opening, morphology if present: present as anteroposteriorly elongate fenestra (0), present as small foramen (1).**

Modified from Brocklehurst et al. (2022, Ch. 154).

**Ch. 346. Suborbital opening, if present, contribution from the maxilla: present (0), absent (1)**

This study.

In araeoscelidians, millerettids, and neodiapsid reptiles, the maxilla contributes to the suborbital foramen or fenestra. In contrast, the suborbital foramen is restricted between the palatine, pterygoid, and ectopterygoid in procolophonians (Carroll and Lindsay 1985).

**Ch. 347. Quadrate, jaw articulation position ( anteroposterior aspect): posterior to occiput (0), approximately level with occiput (1), anterior to occiput (1). ORDERED.**

Modified from Brocklehurst et al. (2022, Ch. 155).

**Ch. 348. Quadrate, jaw articulation position (dorsoventral aspect): at level of alveolar margin of maxillary tooth row (0), quadrate projects far ventral to level of alveolar margin of maxillary tooth row (1). ORDERED.**

Modified from Brocklehurst et al. (2022, Ch. 156).

**Ch. 349. Quadrate, condyles: separate, condyles are asymmetrical ventrally (0), confluent, forming a saddle- shaped articular facet (1).**

Modified from Brocklehurst et al. (2022, Ch. 157).

**Ch. 350. Quadrate, slope of occipital margin: inclined posterodorsally (0), approximately vertical (1),inclined anterodorsally by 10-55 degrees (2), strongly inclined anterodorsally (3), ORDERED.**

Modified from Brocklehurst et al. (2022, Ch. 158).

**Ch. 351. Quadrate, posterior surface, emargination: absent, quadrate is straight posteriorly (0), fossa present but restricted to ventral margin of quadrate shaft (1), present, quadrate shallowly emarginated or pneumatized (2). ORDERED.**

Modified from Brocklehurst et al. (2022, Ch. 159).

We've added State 1 to reflect the reduced quadrate emargination present in non-saurian neodiapsids (e.g., *Thadeosaurus*, Buffa et al. 2025) as well as millerettids (e.g., *Milleretta,* BP/1/3822), in which the emargination is restricted to the lateral surface of the quadrate shaft.

**Ch. 353. Quadrate, posterior surface, if emargination present: shallow emargination (0), distinct excavation or conch framed by two tympanic crests (1). Inapplicable in taxa lacking an emargination on the quadrate (Ch. 346).**

This study.

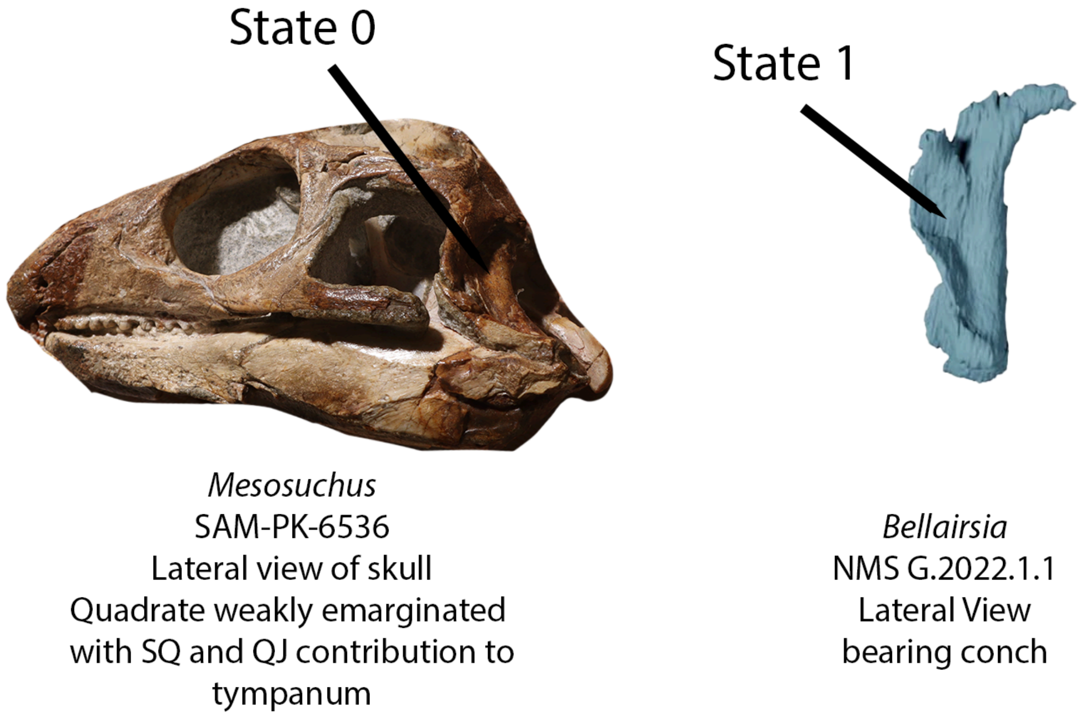


**Fig. S59.** Left, skull of *Mesosuchus* (SAM-PK-6536) in left lateral view, bearing a shallow quadrate emargination (Ch. 353, State 0). Right, quadrate of *Bellairsia* in lateral view bearing a distinct quadrate conch (Ch. 353, State 1).

The quadrate emargination of the EAZ younginiform and early archosauromorphs, presumably housing the tympanic membrane, is relatively shallow and often bears a single, lateral tympanic crest (e.g., *Prolacerta* BP/1/5880, but not in the EAZ younginiform, in which the lateral tympanic crest is located on quadratojugal). The quadrate emargination of lepidosauromorphs (e.g., *Megachirella* PZO628 or *Bellairsia*, modified from Tałanda et al. 2022) is deeply emarginated, taking the form of a conch in lateral view and is framed by a medial and lateral tympanic crest (State 1).

DELTRAN: Lepidosauromorpha (0 to 1). ACCTRAN: Lepidosauria + *Marmoretta* + *Megachirella* (0 to 1).

**Ch. 353. Quadrate, medial surface: a posteriorly-opening groove or concavity forming a "stapedial (=columellar) recess" (0), bears a raised convexity or ridge "stapedial shelf, stapedial boss" (1), flat or featureless (2), distinct, circular "stapedial pit" (3). ORDERED.**

This study, based on observations by Romer and Price (1940) and Carroll (1981).

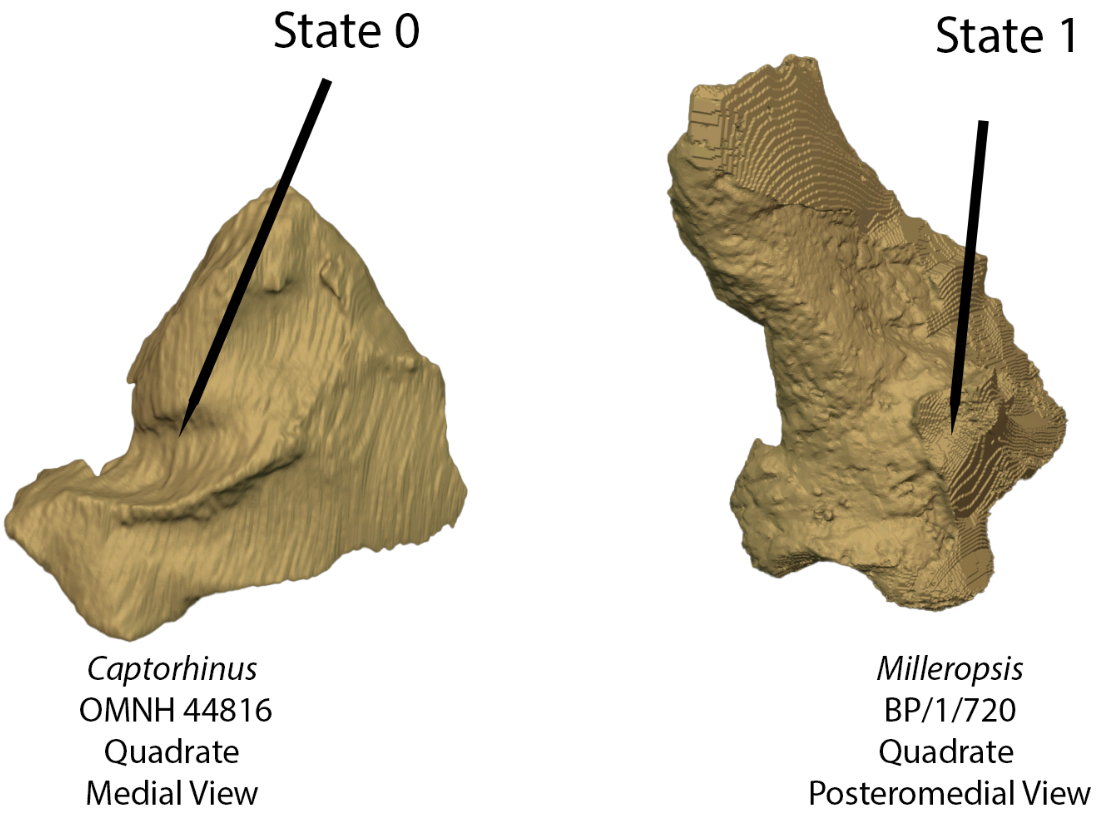


**Fig. S60.** Left, left quadrate of *Captorhinus* (OMNH 44816) in medial view, demonstrating the presence of a stapedial recess (Ch. 353, State 0) and right, left quadrate of *Milleropsis* (BP/1/720) in posteromedial view demonstrating the presence of a raised stapedial boss (Ch. 353, State 1).

The medial surface of the quadrate possesses a groove or fossa, the stapedial (=columellar) recess (State 0), in most early and near-amniotes within this sampling, including captorhinids (e.g., *Captorhinus,* OMNH 44816) protorothyridids (e.g., *Protorothyris* Clark and Carroll 1973), varanopids (e.g., *Heleosaurus*, this study), ophiacodontids (Romer and Price 1940) and araeoscelidians (e.g., *Araeoscelis*, AMNH 4685). In contrast, early neoreptiles, including ankyramorphs (e.g., *Delorhynchus* OMNH 73515), millerettids (e.g., *Milleropsis*, BP/1/720), and neodiapsids (e.g., *Youngina*, Gow 1975) possess a raised ridge, often referred to as a stapedial shelf or boss which cart (State 1). The medial surface of the quadrate of many saurian reptiles (e.g., *Prolacerta* BP/1/5880) is featureless, lacking an indication of osseous contact with the stapes (State 2). Crocopod archosauromorphs (e.g., *Mesosuchus* SAM-PK-6536) possess a small ‘stapedial pit’ which only cartilaginously receives the distalmost end of the stapes (State 3).

UNAMBIGUOUS: Reptiliomorpha (1 to 0) DELTRAN: *Mesosuchus browni* (1 to 3), *Azendohsaurus madagaskarensis* (1 to 3), Sauria (2 to 1). ACCTRAN: Neoreptilia (0 to 2), Chthonosauria (0 to 2).

**Ch. 354. Quadrate, pterygoid ramus, length: long, extends anteriorly at least 40% the length of the quadrate ramus of the pterygoid (0), short, extends anteriorly for less than or equal to 40% the length of the quadrate ramus of the pterygoid (1).**

Modified from Cisneros et al. 2020 (Ch. 29).

The pterygoid ramus of the quadrate is elongate (State 0) in ankyramorphan neoreptiles (e.g., *Saurodektes*, BP/1/4195), sometimes longer than the quadrate ramus of the pterygoid. In most other taxa within this sampling, the pterygoid ramus is reduced, extending less than 40% the length of the quadrate ramus of the pterygoid (State 1).

UNAMBIGUOUS: *Dendrerpeton* + *Balanerpeton* +*Acheloma* + *Cacops* (0 to 1), Moradisaurinae + *Labidosaurus* + *Gansurhinus* + *Captorhinikos* + *Labidosaurikos* (1 to 0). DELTRAN: *Mesosuchus browni* (1 to 0), Allokotosauria + *Mesosuchus*+ Archosauriformes (1 to 0), *Prolacerta broomi* (1 to 0). ACCTRAN: Crocopoda (1 to 0).

**Ch. 355. Quadrate, pterygoid ramus, elevation: ventral margin of pterygoid ramus rises gradually and is ventral to or in line with quadrate condylar surface (0), pterygoid ramus is distinctly elevated relative to quadrate condylar surface (1).**

Modified from Pritchard (2016,Ch. 259).

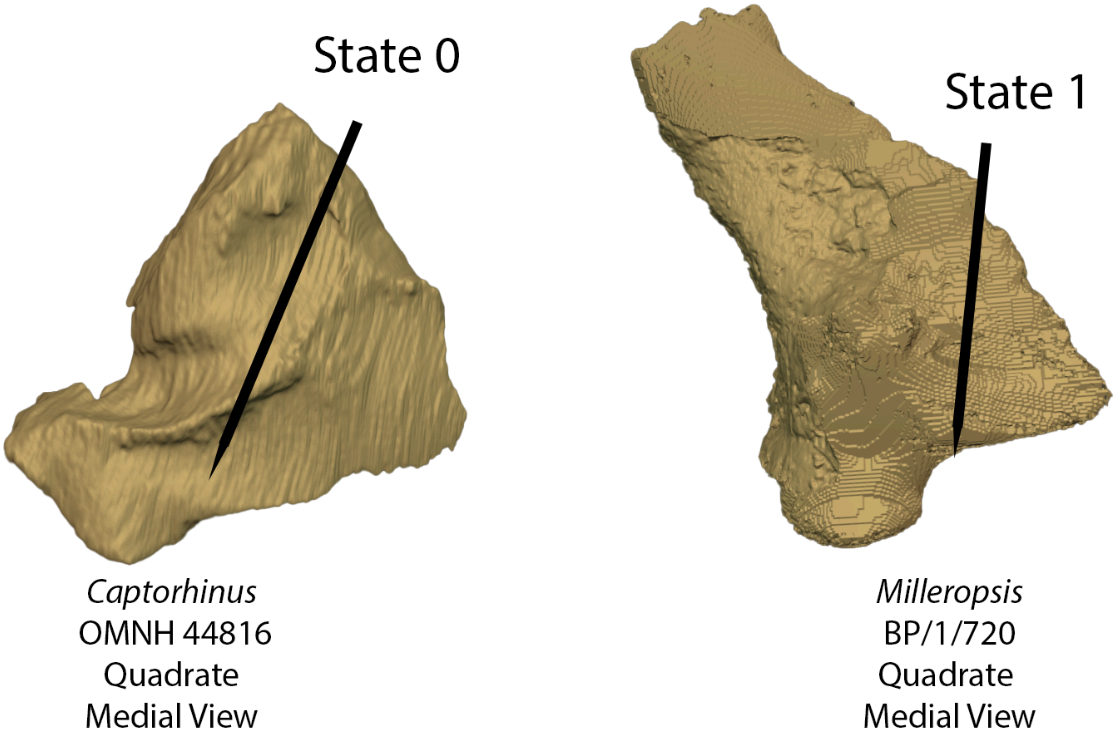


**Fig. S61.** Left, left quadrate of *Captorhinus* (OMNH 44816) in medial view, demonstrating a low pterygoid ramus (Ch. 355, State 0) and right, left quadrate of *Milleropsis* (BP/1/720) in posteromedial view bearing a distinctly raised pterygoid ramus relative to the quadrate condyles (Ch. 354, State 1).

The pterygoid ramus of the quadrate is generally in-line with the quadrate condylar surface in early amniotes (State 0), however, the quadrate ramus of caseid synapsids (Olson 1968) and neoreptiles (including neodiapsids, Pritchard and Nesbitt 2017) are distinctly elevated dorsally relative to the quadrate condylar surface (State 1).

UNAMBIGUOUS: Caseidae (0 to 1). DELTRAN: *Edaphosaurus boanerges* (0 to 1) *Delorhynchus cifellii* (0 to 1), *Colobomycter pholeter* (0 to 1), Ankyramorpha (0 to 1) Parapleurota + *Orovenator* + *Ascendonanus* + *Cabarzia* (0 to 1). ACCTRAN: Neoreptilia (0 to 1), Edaphosauridae + *Ianthodon* (0 to 1).

**Ch. 356. Quadrate, dorsal shaft, dorsal development: dorsal shaft is unexpanded and continuous throughout its length (0), dorsally expanded and hooked (1).**

Modified from Pritchard and Sues (2019, Ch. 75).

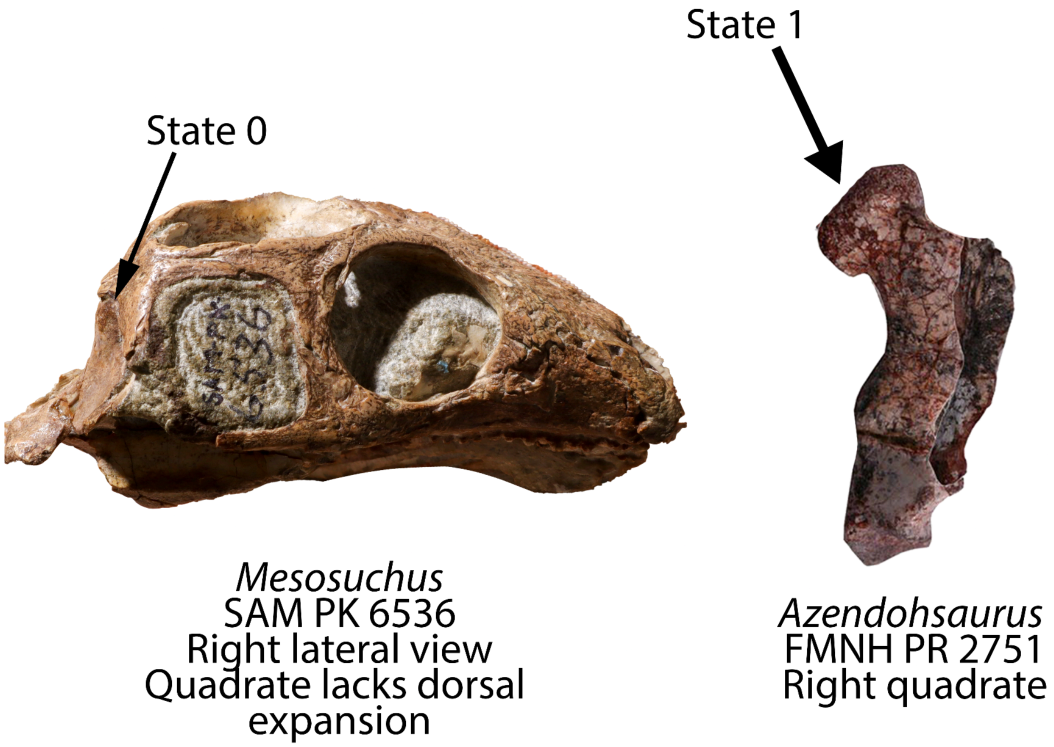


**Fig. S62.** Left, skull of *Mesosuchus* (SAM-PK-6536) in right lateral view bearing a quadrate without a distinct head (Ch. 356, State 0). Right, right quadrate of *Azendohsaurus* (FMNH PR 2751) bearing a distinct condylar head (Ch. 356, State 1).

The dorsal shaft of the quadrate of most early reptiles is unexpanded and straight in lateral view (e.g., *Youngina* BP/1/2871). In contrast, the dorsal shaft of the quadrate in crocopod archosauromorphs bears a distinct 'head' or a mediolateral and anteroposterior expansion (e.g., *Azendohsaurus* UA-7-20-99-653).

DELTRAN: *Tanystropheus hydroides* (0 to 1), *Trilophosaurus buettner*i (0 to 1), *Azendohsaurus madagaskarensis* (0 to 1). ACCTRAN: *Tanystropheus hydroides* (0 to 1), *Mesosuchus browni* (1 to 0), Allokotosauria + *Mesosuchus* (0 to 1).

**Ch. 357. Quadrate, dorsal articular 'cephalic' condyle for squamosal: absent (0), present, quadrate extends dorsally to fit into fossa on the ventral surface of squamosal exposure on skull roof (1).**

Modified from Pritchard and Sues (2019) Ch. 62.

UNAMBIGUOUS: EAZ younginiform (0 to 1), Sauria (0 to 1).

**Ch. 358. Quadrate, foramen for intermandibular vein 'paraquadrate foramen' 'quadratojugal foramen': absent (0), present (1).**

Modified from Gauthier et al. (1988, Ch 21)

The quadrate foramen, sometimes called the 'paraquadrate foramen' or 'quadratojugal foramen' marks the pathway of the intermandibular vein and is present between the quadrate medially and the squamosal and/or quadratojugal laterally. Previous considered by some studies to be a feature of reptiles (e.g., Simões et al. 2022), this foramen is present (State 1) in most early tetrapods (e.g., *Greererpeton* CMNH 11079), captorhinids (e.g., *Captorhinis laticeps*, Heaton 1979, although it is partially obscured in posterior view), synapsids (e.g., *Euromycter*, Sigoneau-Russell and Russell 1974), and reptiles (e.g., *Milleropsis,* BP/1/720), although it is unreported in recumbirostran 'microsaurs' (e.g., *Hapsidopareion,* FMNH 2303), possibly a synapomorphy of that clade (State 0).

DELTRAN: Recumbirostra (1 to 0), *Labidosaurikos meachami* (1 to 0), Therapsida (1 to 0), *Saurodektes kitchingorum* (1 to 0), *Tanystropheus hydroides* (1 to 0), Amniota + Protorothyrididae + Diadectomorpha + Recumbirostra + Captorhinidae (0 to 1), Temnospondyli (0 to 1). ACCTRAN: Dinocephalia + *Biseridens* (1 to 0), Leptopleuroninae (1 to 0), *Tanystropheus hydroides* (1 to 0), Diadectomorpha + Protorothyrididae + Amniota (0 to 1), Amniota + Protorothyrididae + Diadectomorpha + Recumbirostra (1 to 0), Temnospondyli + Reptiliomorpha (0 to 1).

**Ch. 359 Quadrate, if foramen for intermandibular vein present, lateral margin formed by: squamosal (0), quadratojugal only (1). (-) in taxa lacking a quadrate foramen (Ch. 359).**

This study.

When present, the lateral or dorsolateral boundary of the quadrate foramen is formed by the squamosal (State 0) in most Paleozoic tetrapods (e.g., *Ophiacodon,* Romer and Price 1940). In millerettids (e.g., *Milleropsis* BP/1/720) and neodiapsids (e.g., *Prolacerta*, Gow 1975), the lateral margin of the quadrate foramen is formed solely by the quadratojugal (State 1).

DELTRAN: *Ennatosaurus tecton* (0 to 1), *Dimetrodon limbatus* (0 to 1). ACCTRAN: *Ennatosaurus tecton* (0 to 1), Sphenacodontia (0 to 1), Parapleurota (0 to 1).

**Ch. 360. Stapes, quadrate process: absent (0), blade-like with prominent quadrate process, substantially longer than dorsal process (1), quadrate process small or rod-like (2). ORDERED.**

Modified from Brocklehurst et al. (2022, Ch. 312).

**Ch. 361. Stapes, stapedial foramen: present (0), absent (1).**

Modified from Brocklehurst et al. (2022, Ch. 161).

**Ch. 362. Stapes, footplate, anterior suture with braincase: absent, the anterior surface of the stapes footplate is not sutured or fused with surrounding elements (0), present, the anterior surface of the stapes is sutured or fused with the prootic or Parabasisphenoid (1). (-) in taxa lacking an ossified footplate.**

This study.

The stapes of most tetrapods ends freely or cartilaginously within the fenestra ovalis (State 0). In derived captorhinids (e.g., *Labidosaurikos,* OMNH 04331), the footplate of the stapes is sutured or fused to the prootic and/or the parabasisphenoid (State 1).

UNAMBIGUOUS: Temnospondyli (0 to 1), Captorhinidae (0 to 1).

**Ch. 363. Stapes, stapedial footplate, size relative to braincase: massive, stapedial footplate is larger than or equal to the ventral ramus of the opisthotic (0), reduced, stapedial footplate smaller than the ventral ramus of the opisthotic (1). (-) in taxa lacking an ossified footplate.**

This study.

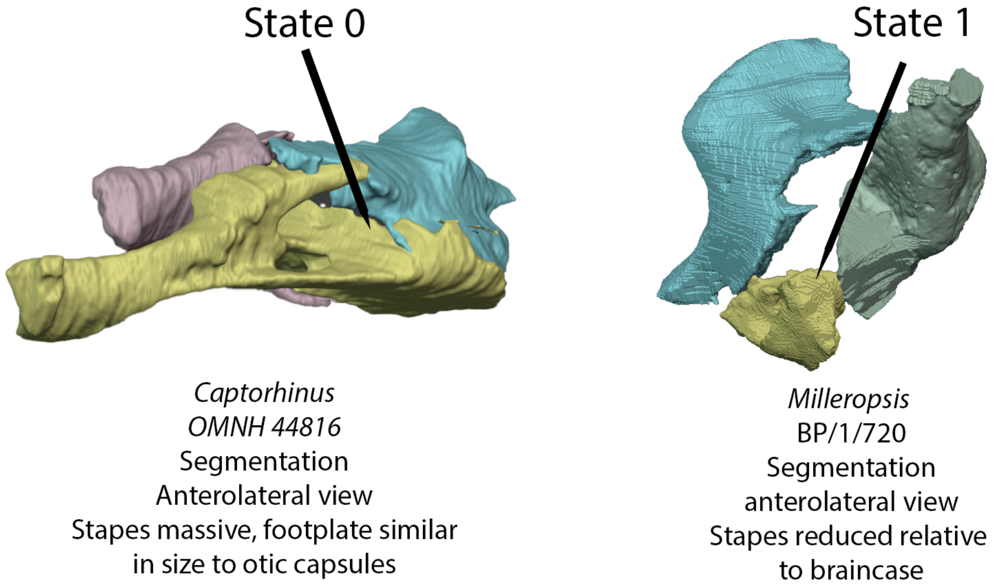


**Fig. S63.** Left, stapes and otic capsule of *Captorhinus* (OMNH 44817) in anterolateral view, which bears an expanded stapedial footplate that is similar in size to the otic capsules (State 0). Right, stapes and otic capsule of *Milleropsis* (BP/1/720, this study) which bears a reduced stapedial footplate relative to the otic capsules (State 1).

The stapedes of early and near-amniotes possess an enlarged footplate that is larger than the ventral ramus of the opisthotic (e.g., *Captorhinus*, Heaton 1979 or *Edaphosaurus*, Modesto 1995). In many reptiles (e.g., *Milleretta,* BP/1/3822 or *Youngina* BP/1/2871) and independently in therapsids (e.g., *Biarmosuchus*, Tchudinov 1960), the stapedial footplate becomes reduced in size relative to the braincase (State 1).

UNAMBIGUOUS: Sphenacodontia (0 to 1), *Australothyris* (1 to 0). DELTRAN: *Deltavjatia* + *Bunostegos* (0 to 1). ACCTRAN: *Deltavjatia* + *Bunostegos* (1 to 0), Reptilia (0 to 1).

**Ch. 364. Stapes, footplate, ventral process: absent, footplate single-headed (0), present, such as that stapedial footplate bears two heads or articular surfaces (1). (-) in taxa lacking an ossified footplate.**

Modified from Schoch (2018, Ch. 151).

The stapedial footplate is single-headed in most tetrapods within this sampling (e.g., *Edaphosaurus* Modesto 1995 or *Prolacerta* BP/1/5880). In temnospondyls, the stapedial footplate bears an accessory ventral process, such that the footplate bears two articular surfaces or heads (e.g., *Dendrerpeton*, Robinson et al. 2005).

UNAMBIGUOUS: *Cacops morrisi* (1 to 0), *Labidosaurus hamatus* (0 to 1), Temnospondyli (0 to 1). DELTRAN: *Deltavjatia rossica* (0 to 1). ACCTRAN: Pareiasauromorpha (0 to 1)

**Ch. 365. Stapes, columella, distal end: columella thins distally or is slightly expanded (0), columella expands distally, becoming greater in height than footplate (1).**

This study.

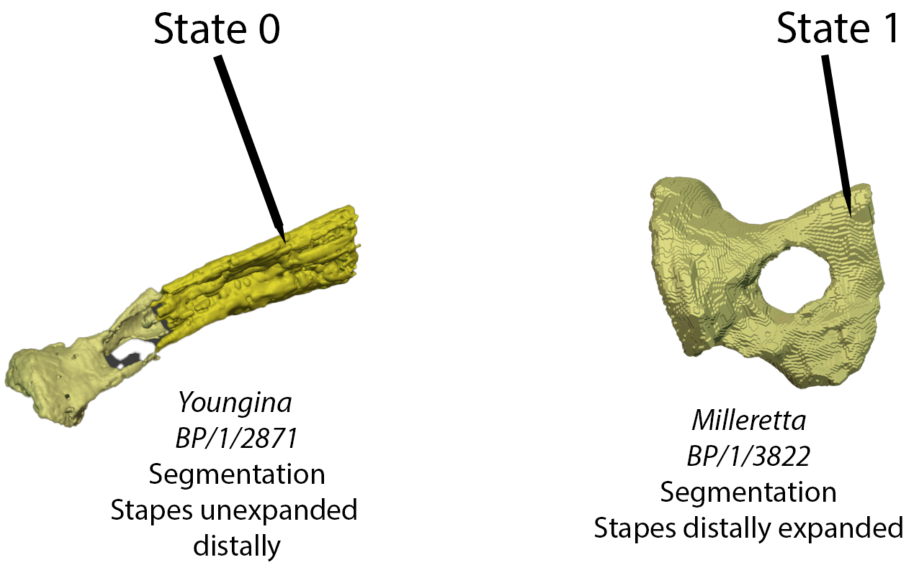


**Fig. S64.** Left, right stapes of *Youngina* (BP/1/2871) in posterior view bearing an unexpanded distal end (Ch. 365, State 0) and right, right stapes of *Milleretta* (BP/1/2871) in posterior view bearing marked distal expansion (Ch. 365, State 1).

The distal end of the stapedial columella is dorsally expanded in millerettids (e.g., *Milleretta,* BP/1/3822) and the enigmatic reptile *Eunotosaurus* (CM 777), such that the distal end of the columella is greater in dorsoventral height than the footplate (State 1).

ACCTRAN: Millerettidae (0 to 1).

**Ch. 366. Stapes, columella, proportions: robust, mediolateral length of columella is less than 4x dorsoventral height at midshaft (0), slender, length more than 4x the dorsoventral height at midshaft (1).**

This study.

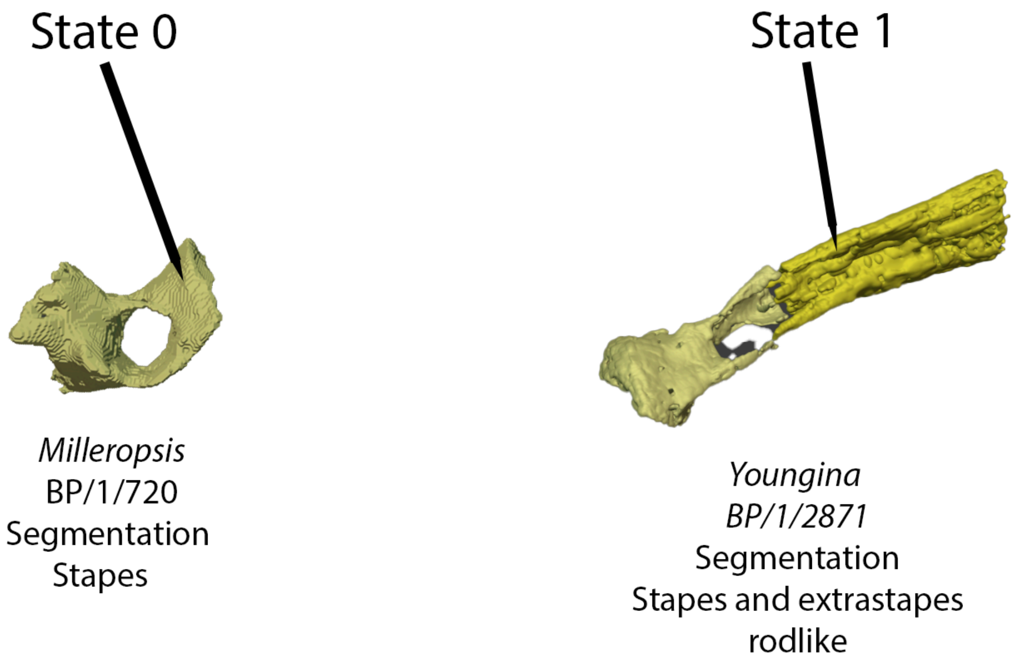


**Fig. S65.** Left, left stapes of *Milleropsis pricei* (mirrored, BP/1/720, this study) demonstrating the plesiomorphic condition of a relatively robust stapes (Ch. 366, State 0) and right, the right stapes of *Youngina* (BP/1/3871, this study) demonstrating the derived condition of a relatively gracile stapes (Ch. 366, State 1).

In most early amniotes, the stapes is a robust element of the occiput, bearing a columella that is less than 4x long as it is the height of its midlength (e.g., *Ophiacodon*, Romer and Price 1940). In most neodiapsids other than *Hovasaurus* (MNHN.1908-32-99), the stapedial columella is slender, being longer than 4x as tall (State 1), although we note that in *Youngina* this is achieved by the accessory ossification of cartilaginous extrastapes (e.g., BP/1/2871).

DELTRAN: *Acheloma* + *Cacops* (0 to 1), *Edops craigi* (0 to 1), Tangasauridae (0 to 1). ACCTRAN: *Dendrerpeton* + *Balanerpeton* (1 to 0), Weigeltisauridae + *Claudiosaurus* + Sauria (0 to 1), Temnospondyli (0 to 1).

**Ch. 367. Stapes, columella, orientation: dorsal, towards skull roof (0), ventrolateral towards quadrate (1).**

Modified from Anderson et al. (2008, Ch. 109), Huttenlocker et al. (2013, Ch. 109).

The stapedial columella is oriented dorsally towards the skull roof (State 0) in many early tetrapods (e.g., *Baphetes*, Clack 2003 or *Cacops,* Reisz et al. 2009), whereas the stapes is oriented towards the suspensorium (quadrate) in reptiliomorphs (e.g., *Araeoscelis*, AMNH FARB 4686) within this sampling.

DELTRAN: Recumbirostra + Captorhinidae + Diadectomorpha + Protorothyrididae + Amniota (0 to 1). ACCTRAN: Reptiliomorpha excluding *Seymouria* (0 to 1).

**Ch. 368. Stapes, dorsal process: present and clearly separate from quadrate process/columella (0), present and confluent with quadrate shaft, grading smoothly into lateral edge of stapes (1), absent or greatly reduced (2). ORDERED.**

Modified from Brocklehurst et al. (2022, Ch. 162).

**Ch. 369. Stapes, dorsal process, articulation with the paroccipital process of the opisthotic: absent (0), present (1).**

Modified from Brocklehurst et al. (2022, Ch. 163).

**Ch. 370. Stapes, dorsal process, distal end: dorsal process unexpanded, thinning distally (0), expands dorsally and wider than its proximal base (1). (-) in taxa lacking the dorsal process of the stapes (Ch. 354).**

This study, based on observations noted by Romer and Price 1940.

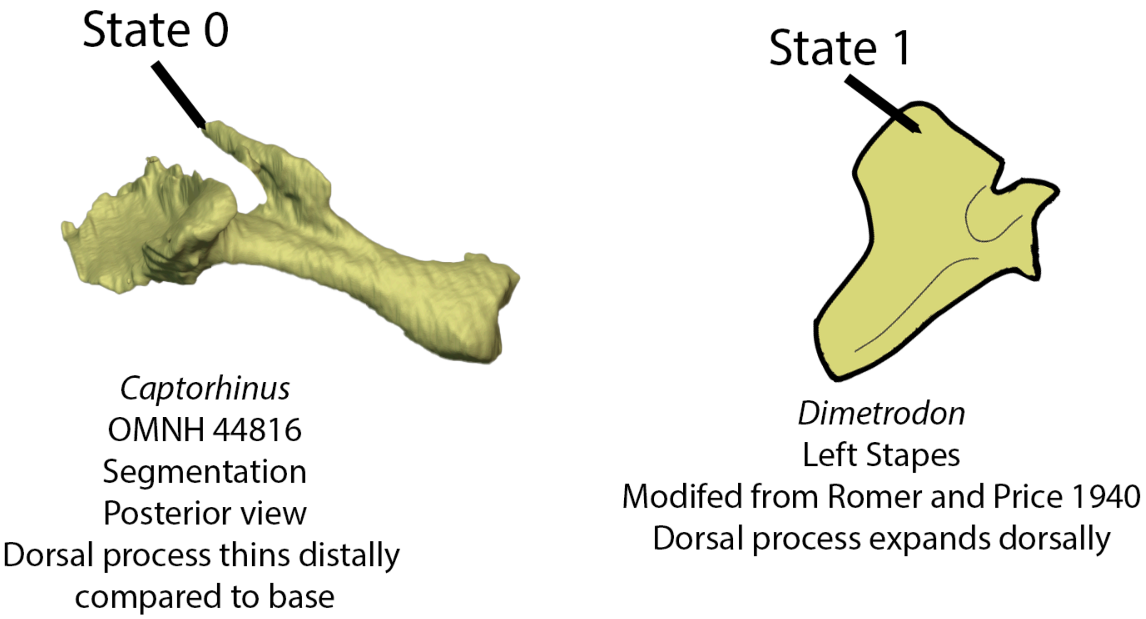


**Fig. S66.** Left, right stapes of *Captorhinus* (OMNH 44816) bearing a distally-thinning dorsal process (Ch. 370, State 0). Right, left stapes of *Dimetrodon* (modified from Romer and Price 1940) bearing a distally expanded dorsal process (Ch. 370, State 1).

The dorsal process of the stapes expands dorsally, being wider than its proximal (ventral) base (State 1) in sphenacomorph synapsids within this sampling (e.g., *Haptodus*, ROM 43604).

UNAMBIGUOUS: Sphenacomorpha (0 to 1).

**Ch. 371. Hyoid apparatus, ossification of basihyal 'basibranchial 1', 'copula' 'corpus hyoideum in mature individuals : unossified in late ontogeny (0), median hyoid element 'basihyal' or 'copula' ossified (1).**

This study.

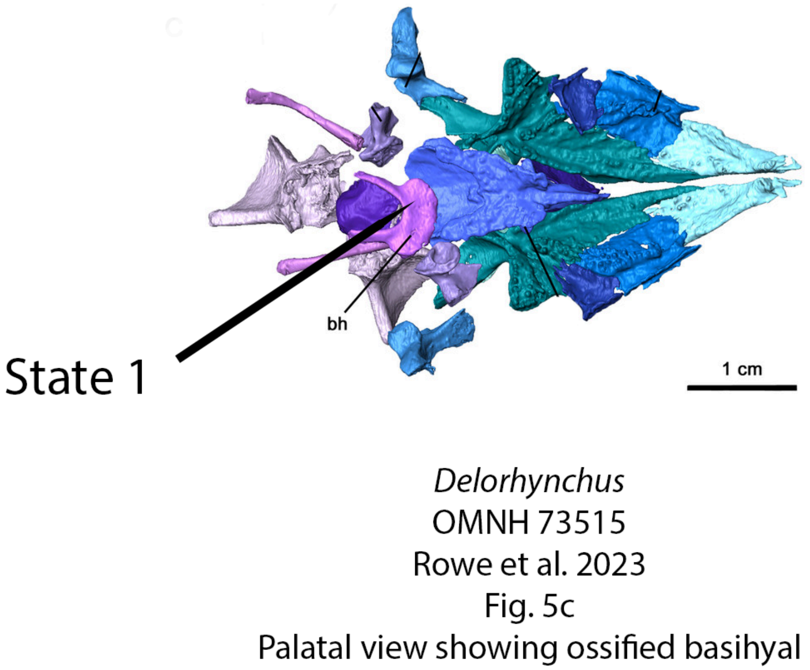


**Fig. S67.** Palate of *Delorhynchus* (OMNH 73515, Rowe et al. 2023) in ventral view bearing an ossified basihyal (Ch. 371, State 1).

The basihyal, also known as as the first basibranchial or hyoid copula, is a midline element of the hyoid that is nearly always ossified (State 1) in ankyramorphan reptiles (e.g., *Procolophon* NHMUK PV R. 10254) but is rarely ossified in other amniotes (State 0).

UNAMBIGUOUS: *Greererpeton burkemorani* (0 to 1), *Brachydectes newberry*i (0 to 1). DELTRAN: Varanops + *Varanodon* (1 to 0), Ankyramorpha + *Erpetonyx* (0 to 1), *Protorothyris archeri* (0 to 1). ACCTRAN: Eupelycosauria + Varanopidae (1 to 0), Mesosauridae + *Ascendonanus* + *Orovenator* + *Cabarzia* +Parapleurota (1 to 0), Araeoscelidia (1 to 0), Amniota + Protorothyrididae + *Anthracodromeus* (0 to 1)

**Ch. 372. Hyoid apparatus, length of first ceratobranchial: short, not longer than quadrate ramus of pterygoid (0), elongate, longer than quadrate ramus and extends posterior to articular region (1).**

Modified from Reisz & Dilkes (2003. Ch. 40), Reisz, Laurin & Marjanović (2010, Ch. 67); Ezcurra, Scheyer & Butler, (2014, Ch. 66).

The first ceratobranchial or ceratohyal, when ossified, is approximately the anteroposterior length of the quadrate ramus of the pterygoid in ventral view (e.g., *Milleropsis,* BP/1/720). Previously interpreted a feature supporting Varanopidae including the Mesenosaurinae (e.g., Reisz et al. 2010), only the ceratobranchial of varanopines is anteroposteriorly elongate (State 1). In *Heleosaurus* (SAM-PK-K8305) and *Elliotsmithia* (TM -1483) the ceratobranchials are dislodged posteriorly, but still more elongate than quadrate ramus of the pterygoid, which not uncommon in paleozoic reptiles where the hyoid elements are not in an osseous contact with the rest of the skull. The first ceratobranchial is also elongated in archosauromorphs within this sampling (e.g., *Mesosuchus* SAM-PK-6536).

UNAMBIGUOUS: Archosauromorpha (0 to 1). DELTRAN: *Varanodon agilis* (0 to 1), *Mesenosaurus romeri* (0 to 1), *Huehuecuetzpalli mixtecus* (0 to 1), Weigeltisauridae + Tangasauridae + *Claudiosaurus* + Sauria (0 to 1). ACCTRAN: Varanopinae + *Apsisaurus* (0 to 1), *Mesenosaurus romeri* (0 to 1), *Bellairsia* + Squamata (0 to 1), Tangasauridae (0 to 1).

**Ch. 373. Mandible, proportions ( not including laminar portion of the angular): very long and dorsoventrally slender, maximum height <0.18 of total length (0), intermediate proportions, dorsoventral height 0.20-0.26 of total length (1), short and robust, dorsoventral height >0.30 of total length (2).**

Modified from Brocklehurst et al. (2022, Ch. 164).

**Ch. 374. Mandible, anterodorsal curvature of anterior mandible in lateral view: absent, alveolar margin of anterior mandible approximately horizontal (0), present, dorsal margin of anterior mandible shows distinct concavity (1).**

Modified from Brocklehurst et al. (2022, Ch. 165).

**Ch. 375. Mandible, posterior margin: rectilinear in lateral view (0), acuminate (1).**

Modified from Brocklehurst et al. (2022, Ch. 166).

**Ch. 376. Mandible, position of coronoid eminence: posteriorly, within posterior 1/3 of total length (0), anteriorly, approximately 2/5 of total length from posterior end (1).**

Modified from Brocklehurst et al. (2022, Ch. 167).

**Ch. 377. Mandible, shape of coronoid eminence: slightly convex or low, barely rises above dorsal surface of surangular or dentary (0), strongly convex (1), subhorizontal/flat (2), dorsoventrally tall (3). ORDERED.**

Modified from Brocklehurst et al. (2022, Ch. 168).

**Ch. 378. Mandible, contribution of dentary to coronoid eminence: absent (0), present (1).**

Modified from Brocklehurst et al. (2022, Ch. 169).

**Ch. 379. Mandible, retroarticular process: absent (0), present (1).**

Modified from Brocklehurst et al. (2022, Ch. 170).

**Ch. 380. Mandible, retroarticular process if present: small (0), large (1).**

Modified from Brocklehurst et al. (2022, Ch. 171).

**Ch. 381. Mandible, retroarticular process, composition: composite, formed by articular, angular and surangular (0), formed only by articular (1).**

Modified from Brocklehurst et al. (2022, Ch. 172).

**Ch. 382. Mandible, retroarticular process, orientation: posteriorly directed (0), upturned posterodorsally (1), curves posteroventrally (2).**

Modified from Brocklehurst et al. (2022, Ch. 173).

**Ch. 383. Mandible, lateral mandibular fenestra between dentary and angular: absent (0), present (1).**

Modified from Brocklehurst et al. (2022, Ch. 174).

**Ch. 384. Mandible, posterior inframeckelian foramen ( foramen intermandibulare caudalis): present (0), absent (1).**

Modified from Brocklehurst et al. (2022, Ch. 175).

**Ch. 385. Mandible, contribution of the splenial or post-splenial to the posterior inframeckelian foramen: absent (0), present (1).**

Modified from Brocklehurst et al. (2022, Ch. 176).

**Ch. 386. Mandible, contribution of the prearticular to the posterior inframeckelian foramen: absent (0), present (1).**

Modified from Brocklehurst et al. (2022, Ch. 177).

**Ch. 387. Mandible, posterior inframeckelian foramen ( foramen intermandibulare caudalis), size: large, greater than or equal to 16% of jaw length (0), small, less than 15% of jaw length (1).**

Reworked from Brocklehurst et al. (2022, Ch. 178).

Rationale: The original wording of this character used the specifiers 25% and 16%. We rephrase here to ensure that intermediate taxa can be scored.

**Ch. 388. Mandible, symphysis: dorsoventrally low, mandible tapers anteriorly (0), dorsoventrally thick, symphysis almost as deep as mandible at midlength of the tooth row (1).**

Modified from Brocklehurst et al. (2022, Ch. 179).

**Ch. 389. Mandible, anterior inframeckelian foramen: absent (0) present (1).**

Modified from Huttenlocker et al. (2013, Ch. 130), Pardo et al. (2017, Ch. 94).

The mandible of early tetrapods often possess more than one inframeckelian foramen or a series of meckelian fenestrae (e.g., *Seymouria* MCZ 1083). All amniotes or near-amniotes within this sampling lack an anterior inframeckelian foramen (e.g., *Orovenator,* Ford and Benson 2018).

DELTRAN: *Seymouria* spp. (0 to 1), Temnospondyli + Reptiliomorpha (1 to 0). ACCTRAN: *Seymouria* spp. (0 to 1), Temnospondyli + Reptiliomorpha (1 to 0).

**Ch. 390. Mandible, lateral surface, sensory canal: absent, ventrolateral surface of mandible smooth or convex (0), present, sensory canal runs along ventrolateral surface of mandible (1).**

Modified from Clack et al. (2012, Ch. 53).

The ventrolateral surface of the mandible in early tetrapodomorphs (e.g., *Crassigyrinus*, Cowdenbeath skull of Panchen 1985) possess a sensory canal for the lateral line system (State 1). It is lost in all temnospondyls and reptiliomorphs within this sampling (State 0).

ACCTRAN: Temnospondyli + Reptiliomorpha (1 to 0). *Archeria crassidiscus* (0 to 1).

**Ch. 391. Mandible, if sensory canal present, oral sulcus of surangular: absent (0), distinct 'oral sulcus' branches of main lateral line onto the dorsolateral surface of the surangular (1).**

From Pardo et al. 2017 (Ch. 278).

An oral sulcus is present on the lateral surface of the surangular (State 1) in the earliest-diverging tetrapodomorphs within this sampling (e.g., *Whatcheeria* FMNH PR 1701). This feature is absent (State 0) in embolomeres (e.g., *Archeria*, Holmes 1989).

DELTRAN: *Archeria crassidiscus* (1 to 0). ACCTRAN: Temnospondyli + Reptiliomorpha (1 to 0).

**Ch. 392. Mandibular ramus in ventral view: straight (0), sigmoid (1).**

Modified from Brocklehurst et al. (2022, Ch. 296).

**Ch. 393. Mandibular ramus: narrow, 8% or less of jaw length (0), broad, at least 14% of jaw length (1).**

Modified from Brocklehurst et al. (2022, Ch. 297).

**Ch. 394. Adsymphysial or ‘parasymphyseal tooth plate’: present (0), absent (1).**

Modified from Clack et al. (2012, Ch. 82) and Pardo et al. (2017, Ch. 335).

The adsymphyseal or ‘parasymphyseal tooth plate’ (although it does not necessarily possess dentition) is an accessory ossification dorsal to the mandibular symphysis that bears teeth/fangs. An adsymphyseal ‘parasymphyseal’ tooth plate is present (State 0) in early tetrapodomorphs (e.g., *Whatcheeria* FM PR 1634 or *Archeria* Holmes 1989) but is absent (State 1) in all known crown-group tetrapods within this sampling (e.g., *Edops* MCZ 1378).

DELTRAN: Embolomeri (0 to 1). ACCTRAN: Temnospondyli + Reptiliomorpha (0 to 1).

**Ch. 395. Adsymphysial, fang pair: fang pairs present (0), absent (1). (-) in taxa lacking an ossified adsymphysial (Ch. 395).**

Modified from Pardo et al. 2017 (Ch. 336), Clack et al. 2012 (Ch. 83).

UNAMBIGUOUS: Temnospondyli + Reptiliomorpha (0 → 1).

**Ch. 396. Dentary size: comprises equal to or greater than 70% of the anteroposterior length of the mandible (0), equal to or less than 65% (1).**

Modified from Brocklehurst et al. (2022, Ch. 180).

**Ch. 397. Dentary, anterior end, split by meckelian canal: absent (0), present (1).**

Modified from Simões et al. (2022, Ch. 167).

UNAMBIGUOUS: Reptilia (0 → 1), Archosauromorpha (1 → 0), Squamata (1 → 0).

**Ch. 398. Dentary, anterior end, symphysial articulatory facet, position: on dorsal margin of Meckelian canal only (0), on dorsal and ventral margins (0), on ventral margin only (1).**

Modified from Simões et al. (2022, Ch. 168).

**Ch. 399. Dentary orientation of anteriormost marginal tooth: mainly vertically (0), leans anteriorly (1), leans strongly anteriorly, at least 45 degrees from perpendicular of the alveolar margin (2). ORDERED.**

Modified from Brocklehurst et al. (2022, Ch. 181).

**Ch. 400. Dentary, anteroposterior length of suture with surangular in lateral view: equal to or shorter than the dentary/angular suture (0), longer that the dentary/angular suture (1).**

Modified from Brocklehurst et al. (2022, Ch. 182).

**Ch. 401. Dentary, posterior extent: extends posteriorly no further than the level of coronoid eminence or slightly beyond (0), extends posteriorly more than halfway between coronoid eminence and articular condyle (1).**

Modified from Ford et al. (2020, Ch. 361).

The dentary of rhynchocephalian lepidosaurs (e.g., *Planocephalosaurus*, Fraser 1982) is posteriorly elongated, extending more than halfway the distance between the coronoid eminence and articular condyles (State 1). In contrast, the dentary of other lepidosaurs and tetrapods within this sampling does not extend much further posteriorly to the coronoid eminence (e.g., *Bellairsia* NMS G.2022.1.1), if at all (State 0).

UNAMBIGUOUS: *Acheloma* + *Cacops* (0 to 1), *Huskerpeton englehorni (*0 to 1), *Opisthodontosaurus carrolli (*0 to 1). DELTRAN: *Euryodus dalyae* (0 to 1), *Hapsidopareion* (0 to 1), Squamata (0 to 1). ACCTRAN: Rhynchocephalia (0 to 1).

**Ch. 402. Dentary, parasymphyseal fangs: present (0), absent (1).**

Modified from Clack et al. (2012, Ch. 279), Pardo et al. (2017, Ch. 332).

Parasymphyseal fangs of the dentary are additional teeth mesial to the dentary tooth row. They are present (State 0) in early tetrapodomorphs (e.g., *Whatcheeria* FM PR 1701) and temnospondyls (e.g., *Acheloma* OMNH 79318), but are absent (State 1) in all stem-amniotes within this matrix (e.g., *Seymouria* BMS E3500).

UNAMBIGUOUS: Reptiliomorpha (0 to 1).

**Ch. 403. Dentary, teeth, asymmetry of dorsoventral height of labial and lingual walls of alveolar groove: lingual wall subequal in height to labial wall (0), lingual wall absent or low, less that half the height of the labial wall (1). (-) in taxa possessing teeth fused to the crista dorsalis (Ch. 28).**

Modified from Ford et al. (2021, Ch. 380).

UNAMBIGUOUS: Parapleurota (0 to 1). DELTRAN: *Dendrerpeton helogenes* (1 to 0), *Cacops morrisi* (1 to 0), *Ennatosaurus tecton* (0 to 1), Eupelycosauria (0 to 1), Leptopleuroninae + *Procolophon* + *Eomurruna* (1 to 0), Tanystropheidae (0 to 1), *Weigeltisaurus jaekeli* (0 to 1), *Feeserpeton* + *Delorhynchus*(1 to 0), Bolosauridae (1 to 0), *Seymouria* spp. (1 to 0), ACCTRAN: *Acheloma cumminsi* (0 to 1), *Ennatosaurus tecton* (0 to 1), Sphenacomorpha (0 to 1), Procolophonoidea + *Nyctiphruretus* (1 to 0), Lepidosauromorpha (0 to 1), Weigeltisauridae (0 to 1), Neoreptilia (0 to 1), Amniota (1 to 0), Temnospondyli + Reptiliomorpha (1 to 0).

**Ch. 404. Dentary, teeth, dorsoventral height of lingual wall of alveolar groove: low, less than half the height of the labial wall (0), absent (1). (-) in taxa possessing teeth fused to the crista dorsalis (Ch. 28)**

Modified from Ford et al. (2021, Ch. 381).

DELTRAN: *Acheloma cumminsi* (1 to 0), *Nannaroter mckinzei* (1 to 0), Edaphosauridae + *Ianthodon* (0 to 1), Pareiasauria + *Emeroleter* (0 to 1), *Mesosaurus tenuidens* (0 to 1), *Marmoretta oxoniensis* (0 to 1), Protorothyrididae (1 to 0). ACCTRAN: Owenettidae (0 to 1) Lepidosauria + *Marmoretta* + *Megachirella* (0 to 1), Mesosauridae (0 to 1), Amniota (1 to 0), Diadectomorpha (1 to 0), Temnospondyli (1 to 0).

**Ch. 405. Dentary, teeth, interdental plates: contact both labial and lingual walls of alveolar groove, fully separating adjacent teeth (0), do not contact both labial and lingual walls of alveolar groove or are absent (1). (-) in taxa possessing teeth fused to the crista dorsalis (Ch. 28).**

Modified from Ford et al. (2021, Ch. 379).

UNAMBIGUOUS: Caseidae (1 to 0), Archosauromorpha (1 to 0). DELTRAN Sphenacodontoidea (1 to 0), Bolosauridae (1 to 0).

**Ch. 406. Dentary, anterior end, ventral keel (= mentonian process): absent (0), present (1).**

Modified from Apesteguia & Novas (2003, Ch. 33), Simões et al. (2020, Ch. 169).

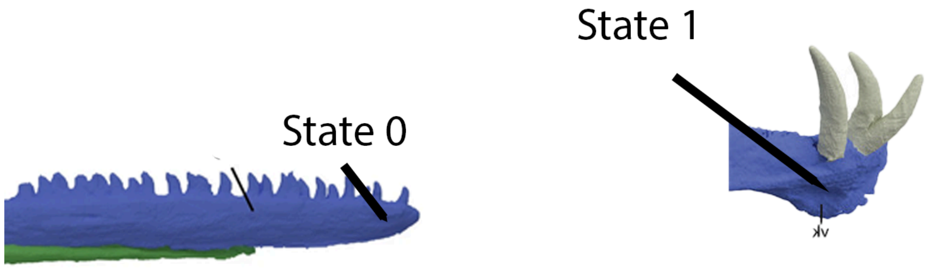

**Fig. S68.** Left, anterior end of dentary of *Macrocnemus* (PIMUZ T 2477, Miedema et al. 2020) with an unexpanded anterior end (Ch. 406, State 0) and right, anterior dentary of *Tanystropheus* (Spiekman et al. 2020) bearing a mentonian-like process of the dentary (Ch. 406, State 1).

The anterior end of the dentary of rhynchocephalian lepidosauromorphs (e.g., *Clevosaurus,* NHMUK PV R36832) possess a ventral keel near the level of the symphysis, giving the dentary an appearance of a 'chin' in lateral or medial view (State 1).

DELTRAN: Squamata (1 to 0), *Tanystropheus hydroides* (0 to 1). ACCTRAN: Rhynchocephalia (0 to 1)

**Ch. 407. Dentary, caniniform teeth: caniniform teeth absent (0), caniniform region present anteriorly (1), Single caniniform tooth present anteriorly (2).**

Modified from Brocklehurst et al. (2022, Ch. 319).

**Ch. 408. Dentary, mesialmost tooth: not enlarged (0), enlarged (1).**

Modified from Brocklehurst et al. (2022, Ch. 320).

**Ch. 409. Dentary, lingual shelf overhanging jaw medially: absent (0), present (1).**

Modified from Modesto et al. 2018 (Ch. 76).

The medial margin of the dentary is expanded, forming a medial or lingual shelf that overhangs the dentary medially. In moradisaurine captorhinids, this feature supports additional rows of dentary dentition (Modesto et al. 2018).

UNAMBIGUOUS: Moradisaurinae (0 to 1).

**Ch. 410. Splenial, contribution to mandibular symphysis: present, symphysis formed from dentary and splenial (0), absent, symphysis formed solely by dentary (1).**

Modified from Brocklehurst et al. (2022, Ch. 183).

**Ch. 411. Splenial, anterior dorsoventral height : narrow, splenial tapers anteriorly (0), deep, maintains dorsoventral height anteriorly (1).**

Modified from Brocklehurst et al. (2022, Ch. 184).

**Ch. 412. Splenial, exposure on lateral surface of mandible: present (0), absent (1).**

Modified from Brocklehurst et al. (2022, Ch. 185).

**Ch. 413. Splenial, contact with posterior coronoid: absent (0), present (1).**

Modified from Brocklehurst et al. (2022, Ch. 186).

**Ch. 414. Postsplenial: present (0), absent (1).**

Modified from Brocklehurst et al. (2022, Ch. 187).

**Ch. 415. Postsplenial, if present, contact with prearticular: absent (0), present (1). (-) in taxa lacking a postsplenial (Ch. 414).**

Modified from Clack et al. (2012, Ch. 67), Pardo et al. (2017, Ch.318).

The postsplenial, when present, makes contact with the prearticular (State 1) in all tetrapodomorphs more derived than *Crassigyrinus* (Cowdenbeath skull of Panchen 1985) and *Greererpeton* (CMNH 11068) within this sampling.

UNAMBIGUOUS: Temnospondyli + Reptiliomorpha (0 to 1).

**Ch. 416. Coronoids, number in mandible: three (0), two (1), one (2), coronoids absent (3). ORDERED.**

Modified from Brocklehurst et al. (2022, Ch. 188).

**Ch. 417. Coronoid, dentition: present (0), absent (1).**

Modified from Brocklehurst et al. (2022, Ch. 189).

**Ch. 418. Coronoid, if dentition present, fangs: absent (0), present, large fangs/teeth present taller than rest of tooth row or field (1). Inapplicable in taxa lacking coronoid dentition (Ch. 416).**

Modified Clack et al. (2012, Ch. 73), Pardo et al. (2017, Ch. 326).

The coronoid of early tetrapodomorphs (e.g., *Archeria* Holmes 1989) possess enlarged fangs (State 1) in addition to a shagreen of denticles. The coronoid of all temnospondyls and reptiliomorphs within this sampling (e.g., *Seymouria* MCZ 1083) lack fangs (State 0).

UNAMBIGUOUS: Temnospondyli + Reptiliomorpha (1 to 0), *Pantylus* (0 to 1).

**Ch. 419. Posterior coronoid, anterior process underlying tooth row: absent, anterior extension of posterior coronoid in medial view ends posterior to or level with posteriormost tooth (0), short, posterior coronoid extends anteriorly no more than 1/3 of the dentary tooth row (1), elongate, posterior coronoid extends anteriorly more than 1/3 the length of the dentary tooth row (2).ORDERED.**

This study.

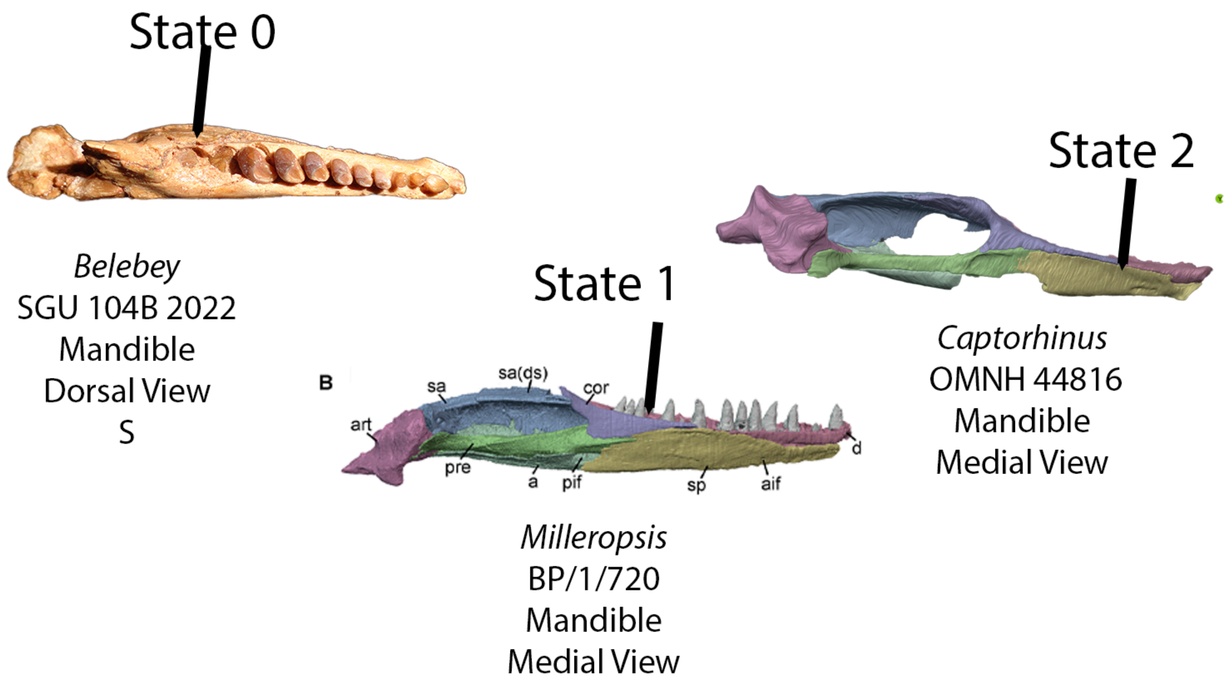


**Fig. S69.** Left, right mandible of *Belebey* (SGU 104B 2022) in dorsal view, demonstrating (Ch. 419, State 0), or an anterior process of the coronoid that does not reach the posterior margin of the tooth row. Middle, *Milleropsis* (BP/1/720, this study) bearing a short anterior process that extends less than a third the length of the dentary (Ch. 419, State 1) and right, left mandible of *Captorhinus* (OMNH 44816) bearing an anteroposteriorly elongated anterior process of the coronoid (Ch. 419, State 2)
.

The anterior process of the coronoid is intermediate in length, not extending more anteriorly than the posterior third of the dentary tooth row (State 1. e.g., *Milleropsis* BP/1/720). In contrast, the posterior coronoid of procolophonoids (e.g., *Eomurruna* QMF 6702,) and bolosaurids (e.g., *Belebey*, SGU 104B/2022) tends to be extremely short or absent, not extending anterior to the posteriormost tooth of the dentary (State 0). The ‘posterior coronoid’ of captorhinids (e.g., *Captorhinus* OMNH 44816), is extremely elongated, sometimes nearly approaching the mandibular symphysis (State 2), and is topologically similar to the extent of the middle coronoid, which is reportedly absent in captorhinids.

UNAMBIGUOUS: *Pantylus cordatus* (1 to 2). DELTRAN: Captorhinidae (1 to 2), Araeoscelidia (1 to 0), Procolophonidae (1 to 0), *Bellairsia gracilis* (1 to 0), *Macrocnemus bassanii* (1 to 0).

**Ch. 420. Posterior coronoid, posteroventromedial process forming the ventral rim of the adductor fossae: absent (0), present (1).**

Modified from Simões et al. (2022, Ch. 203).

UNAMBIGUOUS: Rhynchonkidae (0 to 1). ACCTRAN: Lepidosauromorpha (0 to 1), Mesosauridae + *Ascendonanus* + *Orovenator* + Parapleurota (1 to 0), Amniota + Protorothyrididae + Captorhinidae + Diadectomorpha (0 to 1), Temnospondyli (0 to 1), Embolomeri (0 to 1).

**Ch. 421. Angular, cross-section shape of posteroventral border of angular: ridge or keel absent from ventral surface of angular (0), weakly developed ridge or keel (1), prominent, sheet-like keel with strongly convex posterior edge forming reflected lamina (2). ORDERED.**

Modified from Brocklehurst et al. (2022, Ch. 190).

**Ch. 422. Angular, lateral exposure: exposed along 1/3 of the lateral face of the mandible (0), exposed only as a small sliver along the lateral face (1).**

Modified from Laurin (1991, Ch. J4), Müller ( 2004, Ch 167); Reisz et al. (2010, Ch. 62), Griffiths et al. (2022, Ch. 351).

UNAMBIGUOUS: *Nannaroter mckinzei* (0 to 1), Lepidosauromorpha (0 to 1).

**Ch. 423. Angular, reflected lamina, posterior notch: absent (0), present (1). (-) in taxa lacking a reflected lamina of the angular (Ch. 421).**

Modified from Li et al. (2008, Ch. 55).

UNAMBIGUOUS: Sphenacodontoidea + *Pantelosaurus* (0 to 1).

**Ch. 424. Angular, reflected lamina, posterior emargination: short (0), long with free posterodorsal margin (1).**

Modified from Brocklehurst et al. (2022, Ch. 3212).

**Ch. 425. Angular, ventral margin, ventrally projecting boss or ridge: absent (0), present (1).**

Modified from Laurin (1991, J4), Müller (2004, Ch 167); Reisz et al. (2010, Ch. 62).

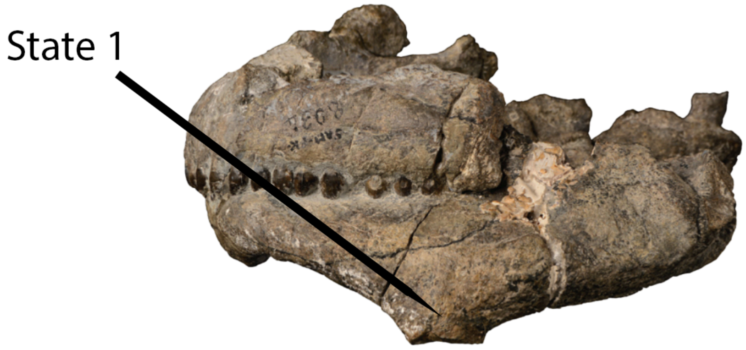


*Embrithosaurus*
Van Den Brandt et al. 2020
Fig. 8
Angular with typical ventral boss
of pareiasaurs

**Fig. S70.** Skull of *Embrithosaurus* in left lateral view demonstrating the ventral angular boss (Ch. 425, State 1).

The ventral margin of the angular of pareiasaurs (e.g., *Embrithosaurus* SAM-PK-8034) possess a ventrally project ridge or boss (State 1), whereas the angular of non-pareiasaurian pareiasauromorphs and other tetrapods remains flat ventrally (State 0).

UNAMBIGUOUS: Pareiasauria + *Emeroleter* (0 to 1).

**Ch. 426. Surangular, anterior extension: extends anteriorly beyond coronoid eminence (0), terminates posterior to or level with the anterior border of the coronoid eminence (1).**

Modified from Brocklehurst et al. (2022, Ch. 191).

**Ch. 427. Surangular, anterior surangular foramen: absent (0), present (1).**

Modified from Brocklehurst et al. (2022, Ch. 192).

**Ch. 428. Surangular, posterior surangular foramen: absent (0), present (1).**

Modified from Brocklehurst et al. (2022, Ch. 193).

**Ch. 429. Surangular, medial expansion of dorsal surface: thin, sheet-like surangular, unexpanded (0), medially expanded along the dorsal margin, forming a broad dorsally facing platform (1).**

Modified from Brocklehurst et al. (2022, Ch. 194).

**Ch. 430. Surangular, lateral adductor crest : absent (0), present (1).**

**Ch. 431. Surangular, anterodorsal process preventing posterior contact of dentary and coronoid: absent (0), present (1).**

Modified from Brocklehurst et al. (2022, Ch. 195).

**Ch. 432. Surangular, morphology of anterodorsal process, if present: dorsoventrally broad posteriorly, tapering anteriorly (0), narrow splint-like process (1).**

Modified from Brocklehurst et al. (2022, Ch. 196).

**Ch. 433. Prearticular, medial surface: nearly straight (0), twisted posteriorly (1).**

Modified from Brocklehurst et al. (2022, Ch. 197).

**Ch. 434. Prearticular, posterior contact with surangular: absent, prearticular contacts articular posteriorly (0), present, prearticular sutures with surangular posteriorly (1).**

Modified from Clack et al. (2012, Ch. 66) and Pardo et al. (2017, Ch. 319).

The prearticular does not contact the surangular posteriorly (State 0) in most reptiliomorphs within this sampling other than *Pantylus* (MCZ 2040), but contact is present in embolomeres (e.g., *Archeria,* Holmes 1989), *Gephyrostegus* (Klembaria et al. 2014), and *Seymouria* (BMS E3500) within this sampling.

UNAMBIGUOUS: Temnospondyli (1 to 0), Taxa more derived than *Whatcheeria* (0 to 1). DELTRAN: *Pantylus cordatus* (0 to 1). Recumbirostra + Captorhinidae+ Diadectomorpha + Protorothyrididae + Amniota (1 to 0).

**Ch. 435. Prearticular, dentition: absent (0), present (1).**

Modified from Clack et al. (2012, Ch. 86) and Pardo et al. (2017, Ch. 339).

The prearticular of non-tetrapod tetrapodomorphs (e.g., *Greererpeton* CMNH 11079) possess a shagreen of denticles on the medial surface (State 0).

UNAMBIGUOUS: Temnospondyli + Reptiliomorpha (1 to 0).

**Ch. 436. Prearticular, medially projecting flange along posterior margin of adductor fossa: absent (0), present (1).**

Modified from Clack et al. (2012, Ch. 69) and Pardo et al. (2017, Ch. 322).

DELTRAN: *Edaphosaurus boanerges* (1 to 0), Bolosauridae (0 to 1), Temnospondyli + Reptiliomorpha (1 to 0). ACCTRAN: Edaphosauridae + *Ianthodon* (1 to 0), Synapsida (0 to 1), Temnospondyli + Reptiliomorpha (1 to 0).

**Ch. 437. Articular, jaw articulation position: At level of posterior alveolar margin of dentary tooth row (0), articular projects ventral to level of posterior alveolar margin of dentary tooth row (1), articular projects far ventrally, below the ventral margin of the anterior mandible (2). ORDERED.**

This study.

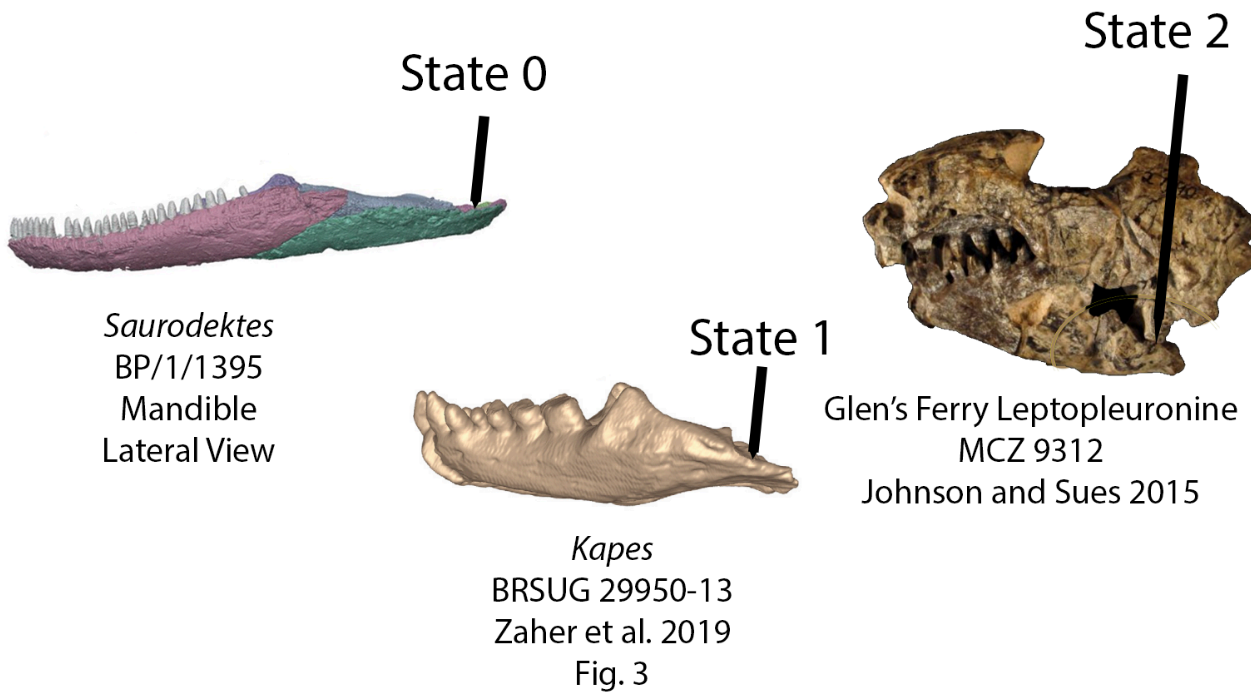


**Fig. S71.** Left, left mandible of *Saurodektes* (BP/1/1395) in lateral view, demonstrating a jaw articulation that is in line with the alveolar margin of the anterior tooth row (Ch. 437, State 0). Middle, left mandible of *Kapes* in lateral view, bearing a jaw articulation ventral to the tooth row (Ch. 437, State 1). Right, left mandible of the Glen's Ferry leptopleuronine (MCZ 8312) demonstrating (Ch. 437, State 2), or a jaw articulation that is well-ventral to the dentary tooth row.

The jaw articulation is level (State 0) with the alveolar margin of the anterior tooth row in most tetrapods within this sampling (e.g., *Saurodektes kitchingorum*, BP/1/4195). The articular projects ventral to the tooth row, but is still dorsal to the ventral margin of the mandible (State 1), in many classically herbivorous taxa such as *Edaphosaurus* (Modesto 1995). The jaw articulation far ventral to the dentary tooth row, projecting ventral to the anterior jaw margin, in extreme instances such as leptopleuronine procolophonids (e.g., *Hypsognathus* YPM 55831).

UNAMBIGUOUS: Caseidae (0 to 1), Leptopleuroninae (1 to 2). DELTRAN: Dinocephalia + *Biseridens* (0 to 1), Edaphosauridae + *Ianthodon* (0 to 1), *Ianthodon schultzei* (0 to 1), *Trilophosaurus buettneri* (0 to 1). ACCTRAN: *Secodontosaurus obtusidens* (1 to 0), Sphenacodontoidea (0 to 1).

**Ch. 438. Articular, orientation of glenoid: dorsal (0), dorsomedial (1).**

This study.

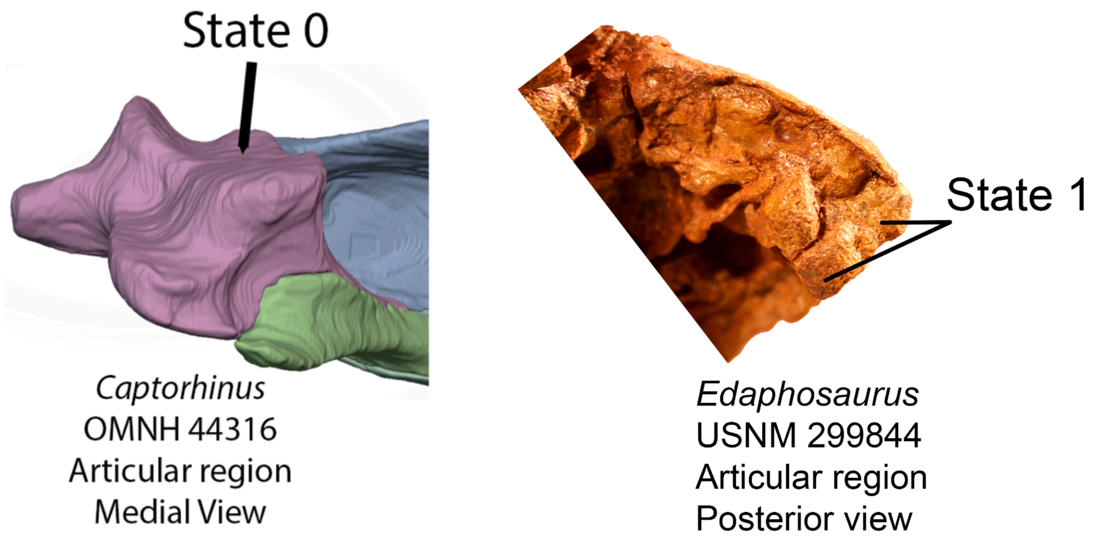


**Fig. S71.** Left, articular region of the left mandible in medial view of *Captorhinus* (OMNH 44316) bearing articular condyles that face dorsally (Ch. 438, State 0). Right, articular of *Edaphosaurus* in posterior view bearing dorsomedially facing articular condyles (Ch. 438, State 1).

The glenoid fossa in most tetrapods is oriented dorsally to articulate with the quadrate condyles (e.g., *Sauropareion*, SAM-PK-11192) or *Ophiacodon,* Romer and Price 1940). The glenoid fossa in sphenacomorph synapsids is oriented strongly dorsomedially and twisted, causing the medial articular cotyle to be placed ventral relative to the lateral cotyle (e.g., *Dimetrodon*, Romer and Price 1940).

DELTRAN: Caseidae: (0 to 1), Eupelycosauria (0 to 1). ACCTRAN: *Euromycter rutenus* (1 to 0), Caseidae (0 to 1), Sphenacomorpha (0 to 1).

**Ch. 439. Articular, lateral shelf: absent (0), present (1).**

Modified from Laurin and Reisz (1995, Ch. 78).

The lateral surface of the articular in most taxa within this sampling (e.g., *Sphenacodon*, UCMP 83459) is smooth (State 0). In contrast, a shelf or crest is present on the lateral surface of the articular (State 1) in pareiasaurs (e.g., *Nochelesaurus*, (SAM-PK-k6238), leptopleuronine procolophonids (e.g., *Hypsognathus*, YPM 55831), and several archosauromorphs (e.g., *Azendohsaurus* UA-7-20-99-653).

UNAMBIGUOUS: Leptopleuroninae + *Procolophon* (0 to 1). DELTRAN: Pareiasauria + *Emeroleter* (0 to 1), *Azendohsaurus madagaskarensis* (0 to 1), *Euparkeria capensis* (0 to 1). ACCTRAN: Pareiasauromorpha (0 to 1), *Azendohsaurus madagaskarensis* (0 to 1), *Euparkeria capensis* (0 to 1).

**Ch. 440. Vertebrae, notochordal canal: present throughout ontogeny (0), absent in adults (1).**

Modified from Brocklehurst et al. (2022, Ch. 199).

**Ch. 441. Vertebrae, pleurocentra, fusion: paired (0), fused with counterpart (1).**

Modified from Huttenlocker et al. (2013, Ch. 155).

UNAMBIGUOUS: *Crassigyrinus* + *Greererpeton* + *Baphetes* (1 to 0). DELTRAN: *Dendrerpeton* + *Balanerpeton* + *Acheloma* + *Cacops* (1 to 0). ACCTRAN: Temnospondyli (1 to 0).

**Ch. 442. Vertebrae, pleurocentra and intercentra: rhachitomous, intercentra dominant weight-bearing element (0), embolomerous, intercentra and pleurocentra equal in size (1), pleurocentra dominant weight-bearing element in vertebral column (1).**

Modified from Huttenlocker et al. (2013, Ch. 155).

UNAMBIGUOUS: Temnospondyli + Reptiliomorpha (0 to 1). DELTRAN: *Dendrerpeton* + *Balanerpeton* + *Acheloma* + *Cacops* (1 to 0), Recumbirostra + Captorhinidae + Protorothyrididae + Diadectomorpha + Amniota (1 to 2) . ACCTRAN: Reptiliomorpha (0 to 2).

**Ch. 443. Dorsal vertebrae, count: 21 or more (0), 20-11 (1), 10 or fewer (2). ORDERED.**

Reworked from Brocklehurst et al. (2022, Ch. 200).

Rationale: Changed state information to refer to dorsal vertebrae as opposed to presacral vertebrae more broadly. Dorsal vertebrae and cervical vertebrae counts vary independently.

**Ch. 444. Presacral/sacral vertebrae, intercentra: present along entire series (0), present only in parts of series, cartilaginous intercentra may be present in places (1) absent (2). ORDERED.**

Modified from Brocklehurst et al. (2022, Ch. 323).

**Ch. 445. Cervical vertebrae, atlantal ribs: present (0), absent (1).**

Modified from Brocklehurst et al. (2022, Ch. 201).

**Ch. 446. Cervical vertebrae, atlas-axis complex, atlantal and axial intercentra: contact ventrally or in very close proximity (0), widely separated by ventral extension of the atlantal centrum (1).**

Modified from Brocklehurst et al. (2022, Ch. 179).

**Ch. 447. Cervical vertebrae, atlantal centrum: separate from axial intercentrum (0), fused to axial intercentrum (1).**

Derived from Pritchard and Nesbitt (2017, Ch. 243).

UNAMBIGUOUS: Parapleurota (1 to 0), DELTRAN: Sphenacomorpha (1 to 0), Lepidosauria + *Marmoretta* + *Megachirella* (0 to 1), Parapleurota (1 to 0). ACCTRAN: Sphenacomorpha (1 to 0), Lepidosauria (0 to 1), Allokotosauria + *Mesosuchus* (0 to 1).

**Ch. 448. Cervical vertebrae, axial intercentrum: not fused to axial centrum (0), fused to axial centrum (1).**

Derived from Pritchard and Nesbitt (2017, Ch. 269).

DELTRAN: Squamata (0 to 1). ACCTRAN: Lepidosauria (0 to 1).

**Ch. 449. Cervical vertebrae, axial neural spine, anteroposterior length of apex: longer than or equal to the centrum (0), shorter than centrum (1), short and spine inclined anterodorsally (2).**

Modified from Brocklehurst et al. (2022, Ch. 203).

**Ch. 450. Cervical vertebrae, axial neural spine, dorsoventral height: low, subequal to the centrum height (0), tall, approximately 1.5 times the height of the centrum (1), very tall, many times the centrum height (2). ORDERED.**

Modified from Brocklehurst et al. (2022, Ch. 204).

**Ch. 451. Cervical vertebrae, third cervical ( = first postaxial cervical), centrum length: subequal to axis centrum length (0), longer than axis centrum length (1).**

**Modified from Ezcurra (2016, Ch. 655).**

UNAMBIGUOUS: Archosauromorpha (0 to 1). ACCTRAN: *Mesosuchus* + *Teyujagua* + Archosauriformes (1 to 0).

**Ch. 452. Cervical vertebrae, ribs, slender and tapering at low angle to vertebrae: absent (0), present (1).**

Modified from Brocklehurst et al. (2022, Ch. 205).

**Ch. 453. Cervical vertebrae, count: 4 or fewer (0), 5 to 7 (1), 8 (2), 9 (3), 10 or more (4). ORDERED.**

Reworked from Brocklehurst et al. (2022, Ch. 200).

Rationale: See Ch. 444 above.

**Ch. 454. Cervical vertebrae, centra length: shorter than or subequal to the dorsal centra (0), markedly longer than dorsal centra (1).**

Reworked from Brocklehurst et al. (2022, Ch. 206).

**Ch. 455. Cervical vertebrae, ventral surface of non-axial cervical centra: rounded (0), bearing a low keel or ridge (1), bearing a prominent, transversely narrow, sheet-like keel (2).**

Reworked from Brocklehurst et al. (2022, Ch. 207).

**Ch. 456. Cervical vertebrae, neural arch excavation ( lateral to the base of the neural spine): absent (0), present (1).**

Reworked from Brocklehurst et al. (2022, Ch. 208).

**Ch. 457. Cervical vertebrae, outline of neural spines in lateral view: sub- rectangular (0), sub-triangular (1),**

Reworked from Brocklehurst et al. (2022, Ch. 209).

**Ch. 458. Cervical ribs, accessory process on anterolateral surface: absent (0), present (1).**

Reworked from Brocklehurst et al. (2022, Ch. 210).

**Ch. 459. Cervical ribs, proximal heads: some or all holocephalous (single- headed) (0), all dichocephalous (1).**

Reworked from Brocklehurst et al. (2022, Ch. 211).

**Ch. 460. Cervicodorsal vertebrae, cervicals and anterior dorsal ribs bear dorsoventral expansion distally: absent (0), present (1).**

Modified from Simões et al. (2022, Ch. 264).

UNAMBIGUOUS: *Greererpeton burkemorani* (1 to 0). DELTRAN: Amniota + Protorothyrididae (1 to 0), *Limnoscelis paludis* (1 to 0). ACCTRAN: Amniota + Protorothyrididae (1 to 0), Diadectomorpha (1 to 0).

**Ch. 413. Cervicodorsal vertebrae, mammillary processes on posterior cervical and anterior dorsal neural spines: absent (0), present (1).**

Modified from Brocklehurst et al. (2022, Ch. 212).

**Ch. 462. Dorsal vertebrae, anteroposterior length of centra relative to height: short, subequal to height (0), 1.5-2.5 times as long as high (1), very long, length is more than 2.5 x centrum height (2). ORDERED.**

Modified from Brocklehurst et al. (2022, Ch. 213).

**Ch. 463. Dorsal vertebrae, ventral surface of dorsal centra: rounded (0), bearing a low ridge or keel (1), bearing a prominent, transversely narrow, sheet-like keel (2), bearing a ventrally raised platform or keel with longitudinal trough (3). ORDERED.**

Modified from Brocklehurst et al. (2022, Ch. 214).

**Ch. 464. Dorsal vertebrae, transverse processes in trunk: short, projecting equal with or only slightly beyond the lateral surface of the neural arch (0), moderately long, projecting clearly beyond the lateral surface of the neural arch and readily visible in dorsal or ventral view (1).**

Modified from Brocklehurst et al. (2022, Ch. 215).

**Ch. 465. Dorsal vertebrae, outline of anterior and middle dorsal neural spines in lateral view: sub-triangular (0), sub-rectangular (1), dorsal neural spines are long and thin (2).**

Modified from Brocklehurst et al. (2022, Ch. 216).

**Ch. 466. Dorsal vertebrae, posterior dorsal neural spines, orientation: approximately vertical (0), moderately anterodorsally inclined (1), strongly posterodorsally inclined (2).**

Modified from Brocklehurst et al. (2022, Ch. 217).

**Ch. 467. Dorsal vertebrae, anterodorsally inclined posterior dorsal neural spines: posteriormost one or two (0), several posterior dorsal neural spines (1).**

Modified from Brocklehurst et al. (2022, Ch. 218).

**Ch. 468. Dorsal vertebrae, neural spines, height: short, approximately 1.5 times centrum height or lower (0), intermediate, 2-3 times centrum height (1), very tall (2). ORDERED.**

Modified from Brocklehurst et al. (2022, Ch. 219).

**Ch. 469. Dorsal vertebrae, neural spines, lateral tubercles: absent (0), present (1).**

Modified from Brocklehurst et al. (2022, Ch. 220).

**Ch. 470. Dorsal vertebrae, neural spines morphology: consistent along column (0), alternating (1).**

Modified from Brocklehurst et al. (2022, Ch. 221).

**Ch. 471. Dorsal vertebrae, anterior centrodiapophyseal lamina: absent (0), present (1).**

Modified from Brocklehurst et al. (2022, Ch. 222).

**Ch. 472. Dorsal vertebrae, dorsolateral surface: flat or weakly concave (0), swollen and convex (1), excavated by a depression lateral to the base of the neural spine (2). ORDERED.**

Modified from Brocklehurst et al. (2022, Ch. 223).

**Ch. 473. Dorsal vertebrae, postzygapophyses: widely spaced (0), contact on midline (1).**

Modified from Brocklehurst et al. (2022, Ch. 224).

**Ch. 446. Dorsal vertebrae, trunk ribs: mostly dichocephalous (double-headed) (0), mostly holocephalous (single-headed) (1).**

Modified from Brocklehurst et al. (2022, Ch. 225).

**Ch. 475. Dorsal vertebrae, trunk ribs, curvature: curved proximally, only weakly curved distally (0), strongly arched proximally, curved throughout length enclosing expanded, “barrel- shaped” trunk (1).**

Modified from Brocklehurst et al. (2022, Ch. 226).

**Ch. 476. Dorsal vertebrae, posteromost dorsals, ribs: ribs present on all dorsal vertebrae (0), two or more posteriormost dorsal ribs absent, creating a "lumbar" region (1).**

This study.

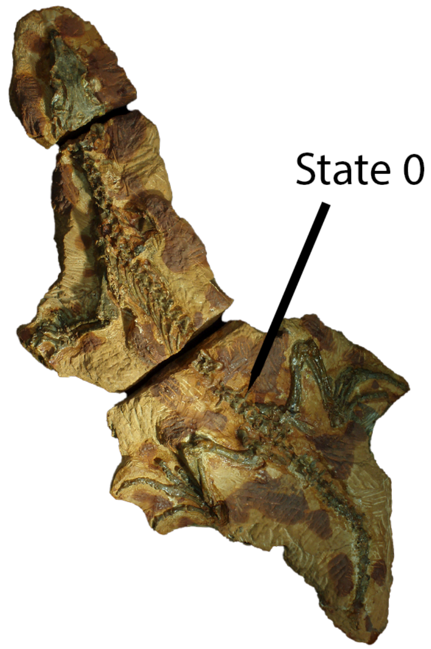


*Mesenosaurus*
PIN 3706/4
Dorsal View

**Fig. S72.** Left, skeleton of *Mesenosaurus* (PIN 3706/4) in dorsal view, bearing ribs on all dorsal vertebrae (Ch. 476, State 0).

In most Paleozoic tetrapods, dorsal ribs are present throughout the dorsal vertebral column (State 0, e.g., *Stereosternum*, BHM 999). In contrast, dorsal ribs are lacking on the posteriormost dorsal centra in procolophonoids (*Saurodektes*, BP/1/4195) creating a 'lumbar' region (State 1).

UNAMBIGUOUS: Pareiasauria (1 to 0). DELTRAN: Ankyramorpha (0 to 1).

**Ch. 477. Dorsal vertebrae, neural arches, articulation with centra: loosely articulated with centrum in mature individuals (0) sutured with centrum in mature individuals (1) fused to centrum in mature individuals (2). ORDERED.**

Modified from Huttenlocker et al. (2013,Ch.149).

DELTRAN: Protorothyrididae (1 to 2), *Thuringothyris* + Captorhinidae (1 to 2), Recumbirostra + Captorhinindae+ Diadectomorpha + Protorothyrididae + Amniota (0 to 1). ACCTRAN: Amniota (1 to 2).

**Ch. 478. Dorsal transverse process, location: approximately at midlength of neural arch, anterior centrodiapophyseal lamina oriented anteroventrally (0), located anteriorly, anterior centrodiapophyseal lamina vertical (1).**

**Ch. 479. Dorsal prezygapophyses: planar, do not contact on midline (0), transversely concave, contact on midline (1), planar and inclined strongly medially, contact on midline (2).**

Modified from Brocklehurst et al. (2022, Ch. 325).

**Ch. 480. Dorsal postzygapophyses, hyposphene between postzygapophyses: absent (0), present and prominent (1).**

Modified from Brocklehurst et al. (2022, Ch. 224).

**Ch. 481. Dorsal neural arches, zygosphene-zygantrum as bilateral accessory articulations: absent (0), present (1).**

Derived from Rieppel et al. (1999), Muller (2004, Ch. 44), Pritchard et al. (2015, Ch. 130), Ezcurra (2016, Ch. 360).

DELTRAN: Lepidosauria + *Marmoretta* + *Megachirella* (0 to 1). ACCTRAN: Lepidosauria (0 to 1), Mesosauridae (0 to 1).

**Ch. 482. Dorsal and sacral neural spines, dorsal end: unexpanded (0), transversely expanded forming spine table (1).**

Modified from Brocklehurst et al. (2022, Ch. 225).

**Ch. 483. Dorsal ribs, tuberculum (contacts diapophysis) morphology: well-developed and flange-like (0), reduced to low tuberosity (1), low tuberculum with expanded, concave, cup-like articular facet (2).**

Modified from Brocklehurst et al. (2022, Ch. 226).

**Ch. 484. Patagial ossifications (elongate bony spars positioned lateral/superficial to dorsal ribs: absent (0), present (1).**

Modified from Pritchard et al. (2021 Ch. 204).

Patagial ossifications are a unique autapomorphy of weigeltisaurids within this taxonomic sampling (Pritchard et al. 2021, Buffa et al. 2022).

UNAMBIGUOUS: Weigeltisauridae (0 to 1).

**Ch. 485. Osteoderms, midline rows dorsal to vertebral column: absent, no osteoderms present (0), absent, unorganized osteoderms present dorsal to vertebral column (1), present, midline row (2), present, two rows present with their median margins approaching the vertebral midline ( = paramedian rows) (3).**

Modified from Brocklehurst et al. (2022, Ch. 227).

**Ch. 486. Sacral vertebrae, morphology of first sacral rib: greatly enlarged in comparison with the second sacral rib(0), subequal to or only slightly larger than the second sacral rib (1).**

Modified from Brocklehurst et al. (2022, Ch. 228).

**Ch. 487. Sacral vertebrae, second sacral rib: not bifurcate (0), bifurcate (1).**

Modified from Brocklehurst et al. (2022, Ch. 229).

**Ch. 488. Sacral vertebrae, second sacral rib, contacts first sacral rib anterolaterally: absent (0), present, sacral ribs are in contact for much of their distal extent (1).**

This study.

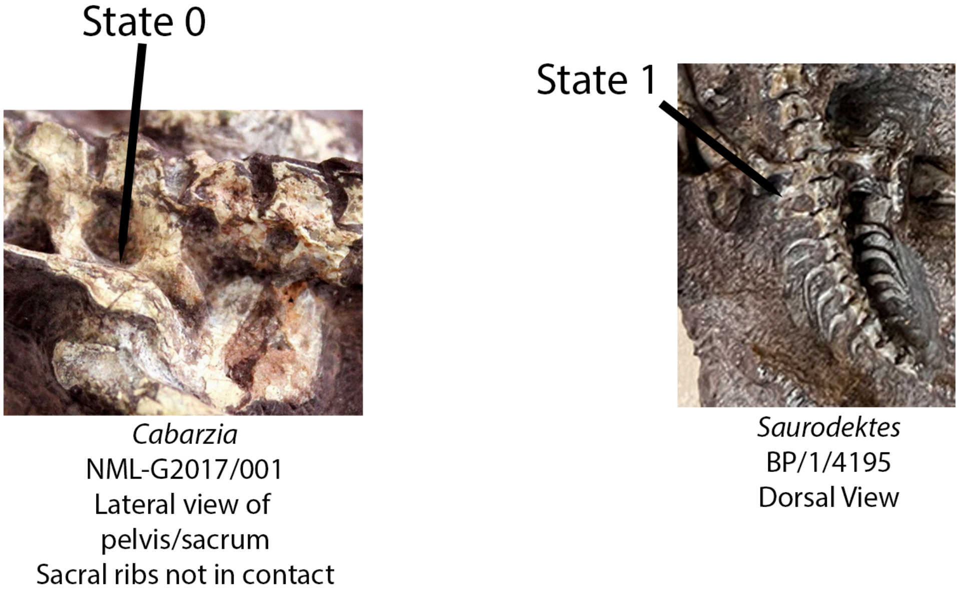


**Fig. S73.** Left, sacral ribs of *Cabarzia* (NML-G2017/001) in dorsolateral view, which do not contact distally (Ch. 488, State 0). Right, sacral region of *Saurodektes* (BP/1/4195) in dorsal view, with sacral ribs in broad contact distally (Ch. 488, State 1).

The second sacral rib generally contacts the first sacral rib anterolaterally in early amniotes (e.g., *Petrolacosaurus*, Reisz 1981). The second sacral rib does not contact the first sacral rib in the early reptile *Ascendonanus* (Spindler et al. 2018) or mesenosaurines (Botha-Brink and Modesto 2009) (Spindler et al. 2019).

DELTRAN: *Deltavjatia rossica* (1 to 0), *Owenetta kitchingorum* (1 to 0), *Nyctiphruretus acudens* (1 to 0), *Euparkeria capensis* (1 to 0), *Coelurosauravus elivensis* (1 to 0), Sauropsida (1 to 0). ACCTRAN: Procolophonoidea + *Nyctiphruretus (*0 to 1), Ankyramorpha (1 to 0), Archosauriformes (1 to 0), Weigeltisauridae (1 to 0).

**Ch. 489. Sacral vertebrae, count: one (0), two (1), three or greater (2).**

Reworked from Brocklehurst et al. (2022, Ch. 230).

Rationale: Added State 1, as several tetrapodomorphs within this sampling possess a single sacral vertebrae.

**Ch. 490. Sacral and caudal neural spines: smooth (0), rugose with longitudinal ridges on lateral surface and tapering apex (1).**

Modified from Brocklehurst et al. (2022, Ch. 329).

**Ch. 491. Caudal vertebrae, anterior caudal neural spine height: low with height/length <1.0 (0), moderately tall with height/length between >1.0 and <2.0 )(1), tall with height/length >2.0 and <3.0 (2), very tall with height/length >3.0 (3). ORDERED.**

Modified from Brocklehurst et al. (2022, Ch. 231).

**Ch. 492. Caudal vertebrae, chevrons, distal width of haemal spine: wider than proximal width(0), equivalent to proximal width (1), tapering (2). ORDERED.**

Modified from Brocklehurst et al. (2022, Ch. 232).

**Ch. 493. Caudal vertebrae, autotomic septum: absent (0), present (1).**

Modified from Pinna (1986, Ch. 52), Ford et al. (2021, Ch. 234), Simões et al.( 2022, Ch. 236).

DELTRAN: Moradisaurinae + *Labidosaurus* + *Gansurhinus* + *Captorhinikos* + *Labidosaurikos* (0 to 1), Lepidosauria + *Marmoretta* + *Megachirella* (0 to 1). ACCTRAN: Lepidosauromorpha (0 to 1), Captorhinidae (0 to 1).

**Ch. 494. Caudal vertebrae, orientation of ribs or transverse processes, when fused: posterolaterally directed and "hook-like", extending posteriorly past the centrum of the succeeding vertebrae (0), laterally directed and straight (1).**

Modified from Pritchard and Nesbitt (2017, Ch. 133), MacDougall et al. (2019, Ch. 132).

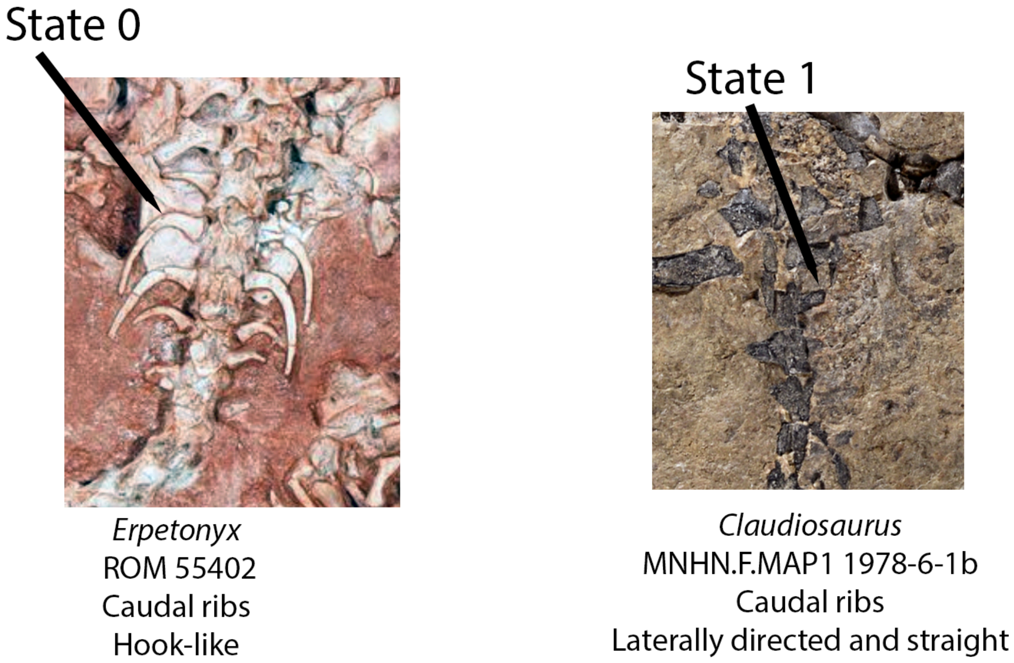


**Fig. S75.** Left, caudal ribs of *Erpetonyx* (ROM 55402) in dorsal view, exhibited the plesiomorphic condition of hook-like caudal ribs (Ch. 494, State 0) and right, caudal ribs of *Claudiosaurus* (MNHN.F.MAP1 1978-06-1b) which bears laterally-directed caudal ribs (Ch. 494, State 0).

The caudal ribs of early amniotes (e.g., *Saurodektes*, BP/1/4195) are hooklike, extending posteriorly past the centra of the succeeding vertebrae (State 0). In contrast, the caudal ribs of millerettids (e.g., *Broomia*, NHMUK 4065) and neodiapsids (Carroll 1981) are laterally directed and straight, completely fused to the caudal central (State 1).

UNAMBIGUOUS: Paurapleurota (0 to 1) DELTRAN: Therapsida (0 to 1), Pareiasauria + *Emeroleter* (0 to 1), *Ascendonanus* + *Cabarzia* (0 to 1). ACCTRAN: Dinocephalia + *Biseridens* (0 to 1), Mesosauridae (0 to 1).

**Ch. 495. Caudal vertebrae, ribs or transverse processes, number: present on five or fewer anterior caudals (0), present on 6-12 (1), present on 13 or more (2).**

Modified from Laurin and Reisz (1995), Cisneros et al. (2020,Ch. 129).

Millerettids (e.g., *Milleretta*, BP/1/3821 or *Broomia* NHMUK 4065) possess five or fewer caudal ribs restricted to the proximal tail (State 0).

DELTRAN: Therapsida (1 to 2), Neodiapsida (0 to 2), Millerettidae (1 to 0), *Saurosternon bainii* (0 to 1).

**Ch. 496. Caudal vertebrae, transverse processes in mature individuals: sutured to vertebra (0), fused to vertebra (1).**

**This study.**

In most early tetrapods, the caudal ribs or transverse processes are weakly sutured to the neural arch of centrum. In parapleurotan reptiles such as *Broomia perplexa* (Thomassen and Carroll 1981), the suture between the transverse processes and caudal vertebrae is absent in mature individuals.

UNAMBIGUOUS: Diadectomorpha (0 to 1), Mesenosaurinae (0 to 1), Mesosauridae (0 to 1), Parapleurota (0 to 1).

**Ch. 497. Caudal vertebrae, supraneural radials: present, supraneural radials dorsal to caudal neural spines (0), absent (1)**

Derived from Pardo (2024).

Derived from observations in Pardo (2024), which note that portions of the caudal tail fin remain in early tetrapodomorphs such as *Whatcheeria* and *Archeria* (Pardo 2024) .
DELTRAN: Temnospondyli + Reptiliomorpha (0 to 1).

**Ch. 498. Gastralia: present (0), absent (1).**

Modified from Clack et al. (2012, Ch. 111), Pardo et al. (2017, Ch. 362).

UNAMBIGUOUS: *Brachydectes newberryi* (0 to 1). Taxa more derived than *Whatcheeria* (1 to 0). DELTRAN: Pareiasauria + *Emeroleter* (0 to 1), Lepidosauria (0 to 1). ACCTRAN: *Emeroleter levis* (1 to 0) *Bellairsia* + Squamata (0 to 1).

**Ch. 499. Gastralia, if present, morphology: present as rounded or rectilinear 'gastral scales' (0), present as paired rows of discrete, rod-like bones forming a 'gastral basket' (1).**

Modified from Clack et al. (2012, Ch. 111), Pardo et al. (2017, Ch. 362).

The ventral or ‘gastral’ scales of early tetrapodomorphs (e.g., *Proterogyrinus*, MCZ 4537) are present as scales (State 0). In contrast, the gastralia of known reptiliomorphs (when present) are present as rod-like bones that form a ‘gastral basket’ ventrally (e.g., *Batropetes*, SMNS 5588).

DELTRAN: Recumbirostra (0 to 1), Amniota + Protorothyrididae + Captorhinidae (0 to 1). ACCTRAN: Diadectomorpha + Protorothyrididae + Amniota + Recumbirostra (0 to 1), *Pantylus* (1 to 0).

**Ch. 500. Gastralia, lateral gastrale, number of segments per row: five or more (0), three to four (1), two (2), one (3). ORDERED.**

This study.

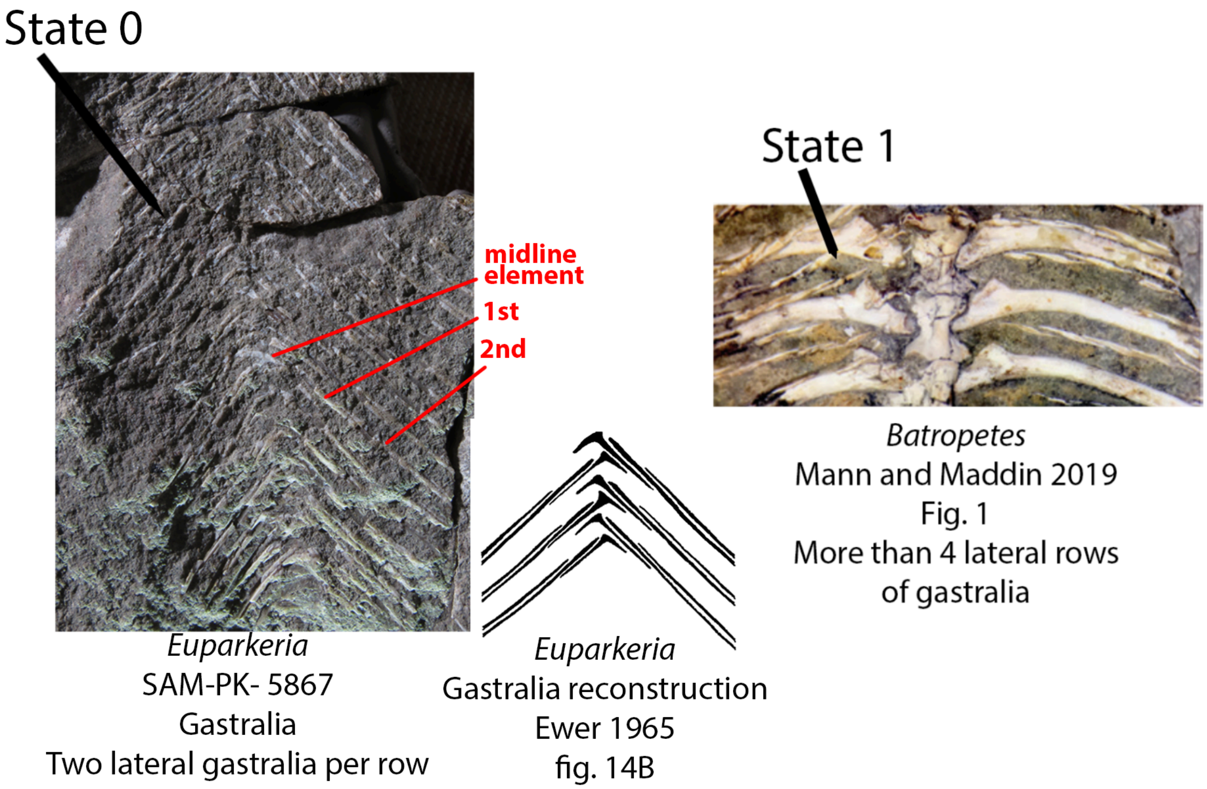


**Fig. S76.** Left, gastral basket of *Euparkeria* (SAM-PK-5867) which bears two lateral gastralia per row (Ch. 500, State 0) and right, gastral basket of *Batropetes* (Mann and Maddin 2019) which bears upwards of four gastralia per row (Ch. 500, State 1).

The gastralia of early reptiliomorphs (e.g., *Proterogyrinus*, MCZ 4537) possess upwards of three lateral gastralial elements per row (State 1). In contrast, millerettids (e.g., *Milleropsis,* BP/1/4203) and neodiapsids possess two lateral gastralia elements or fewer (State 0). The millerettid *Milleretta* (e.g., BP/1/3822) and the probable millerettid *Eunotosaurus* (e.g. NMQR 3299) possess a single lateral gastrale per row (State 3), a feature shared with some testudines among early reptiles, an observation noted by Lyson et al. (2013).

UNAMBIGUOUS: Parapleurota (1 to 2). ACCTRAN: Amniota + Protorothyrididae (0 to 1), Captorhinidae (0 to 1), *Milleretta* + *Millerosaurus* (2 to 3).

**Ch. 501. Gastralia, if present, midline element: absent (0), midline element overlapped laterally by rows of gastralia present (1)**

This study.

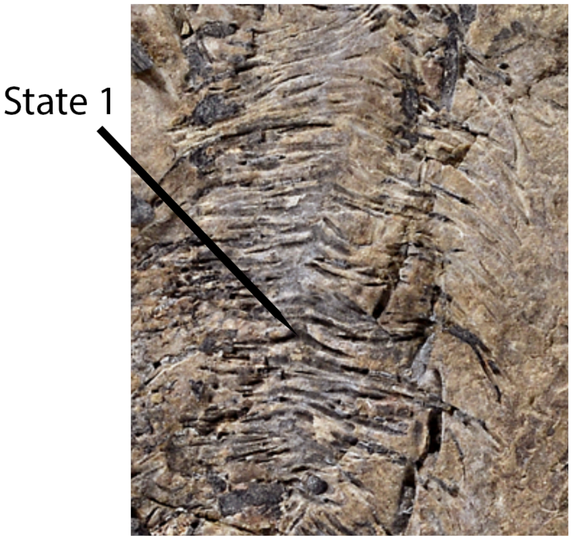


**Fig. S77.** Skeleton of *Claudiosaurus* (MNHN.F.MAP1 1978-6-1b) bearing a midline gastral element (Ch. 501, State 1).

*Millereopsis* (BP/1/4203) and early neodiapsids (e.g., *Claudiosaurus*, Carroll 1981) possess a midline element that is overlapped by both rows of lateral gastralia (State 1).

UNAMBIGUOUS: Parapleurota (0 to 1). DELTRAN: Eupelycosauria + Varanopidae (0 to 1). ACCTRAN: Eupelycosauria (0 to 1).

**Ch. 502. Posternal inscription ribs: absent or cartilaginous (0), present, inscriptional ribs articulate with dorsal ribs distally (1)**

Modified from Estes (1988, Ch. 110).

The ossification of poststernal inscriptional ribs is widespread in stem-squamates and crown Squamata, including *Bellairsia* (Talanda et al. 2022) and *Huehuecuetzpalli* (Reynoso 1998) within the current sampling.

UNAMBIGUOUS: *Bellairsia* + Squamata (0 to 1).

**Ch. 503. Clavicle, orientation of long axis of ventromedial plate relative to shaft: highly obtuse angle (0), almost perpendicular (1).**

Modified from Brocklehurst et al. (2022, Ch. 233).

**Ch. 504. Clavicle, anteroposterior width of ventromedial plate: narrow, less than three times the width of the midpoint of the dorsal shaft (0), narrow, but with a distinct anterior process of the**

**ventromedial plate (1), intermediate, equal to or more than 3 times the width of the midpoint of the dorsal shaft (2), deep, at least 4 times the width of the midpoint of the dorsal shaft (3).**

Modified from Brocklehurst et al. (2022, Ch. 234).

**Ch. 505. Clavicle, contact with contralateral clavicle on midline: present (0), absent (1).**

Modified from Huttenlocker et al. (2013, Ch. 195), Pardo et al. (2017, Ch.157).

UNAMBIGUOUS: Reptiliomorpha (1 to 0). ACCTRAN: Recumbirostra (0 to 1), *Echinerpeton* + *Archaeothyris* (0 to 1), *Broomia* (1 to 0), *Euparkeria capensis* (1 to 0), Tangasauridae (1 to 0), Parapleurota (0 to 1), Araeoscelidia (0 to 1), *Protorothyris archeri* (0 to 1).

**Ch. 506. Interclavicle, extensive dermal sculpturing on ventral surface: absent, interclavicle unornamented (0), present, ventral surface of interclavicle bears dermal sculpturing (1)**

Derived from Huttenlocker et al. (2013, Ch. 189), Pardo et al. (2017, Ch. 151).

DELTRAN: *Brachydectes newberryi* (1 to 0), *Opisthodontosaurus carrolli* (1 to 0), *Protocaptorhinus price*i (1 to 0), Amniota + Protorothyrididae (1 to 0), Diadectomorpha + Protorothyrididae + Amniota (1 to 0). ACCTRAN:Moradisaurinae + *Labidosaurus* + *Captorhinikos* + *Labidosaurikos* (0 to 1).

**Ch. 507. Interclavicle, head, anterior fimbriation: absent (0), present, interclavicle head bears fimbriation on anterior surface (1). (-) in taxa with a midline contact of the clavicles.**

Modified Huttenlocker et al. (2013, Ch. 189), Pardo et al. (2017, Ch.151).

The interclavicular head of early tetrapods often possess fimbriation or striations on the ventral surface (State 1, e.g., *Gephyrostegus* Carroll 1970 or *Archeria* Romer 1957). In contrast, the interclavicular head of stem-amniotes within this sampling lack this fimbriation (e.g., *Pantylus* MCZ 2040).

UNAMBIGUOUS: Reptiliomorpha (1 to 0), *Proterogyrinus scheelei* (1 to 0).

**Ch. 508. Interclavicle, median notch on anterior margin: absent (0), present (1).**

Modified from Brocklehurst et al. (2022, Ch. 235).

**Ch. 509. Interclavicle, posterior stem: little change in width along entire length or attenuates posteriorly (0), expansion present (1).**
Modified from Brocklehurst et al. (2022, Ch. 236).

**Ch. 510. Interclavicle, ventral surface of posterior stem: smooth or marked by a low, anteriorly restricted ridge (0), marked by a strongly developed ridge that extends posteriorly (1).**

Modified from Cisneros (2008, Ch. 47).

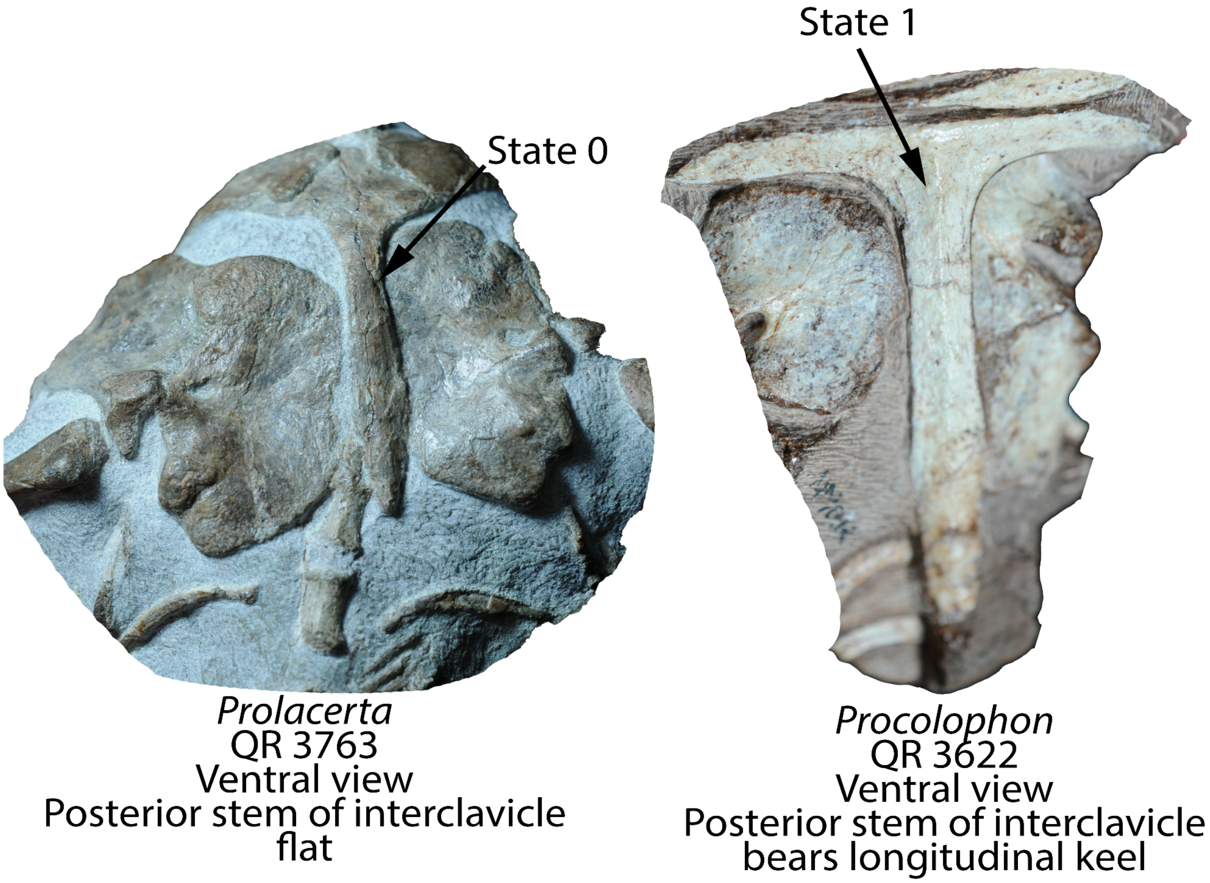


**Fig. S78.** Left, interclavicle of *Prolacerta* (NMQR 3763) in ventral view bearing a flat ventral surface on the posterior stem (Ch. 510, State 0). Right, interclavicle of *Procolophon* (NMQR 3622) bearing a longitudinal keel on the ventral surface of the posterior stem (Ch. 510, State 1).

Procolophonids within this sampling (e.g., *Eomurruna*, Hamley et al. 2021) possess a midline ridge or keel on the ventral surface of the posterior ramus of the interclavicle.

**Ch. 511. Interclavicle , shape of interclavicular head in ventral view: paddle- shaped (0), diamond or spade shaped with anteroposterior length sub-equal to or greater than the transverse width (1), lateral processes or wings present, with transverse width approximately two times greater than anteroposterior length of interclavicular head (2).**

Modified from Brocklehurst et al. (2022, Ch. 237).

**Ch. 512. Interclavicle, if lateral processes present, shape: straight or weakly concave, do not curve posteriorly (0), curve strongly posteromedially or anchor-shaped (1). (-) in taxa that possess an interclavicle lacking lateral processes (Ch. 511)**

Modified from deBraga and Reisz (1996).

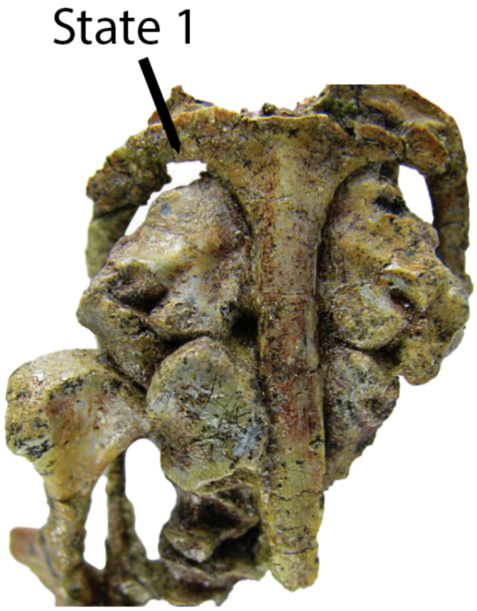


**Fig. S79.** Interclavicle of *Eomurruna* (Hamley et al. 2021) in ventral view, bearing distinctly curving lateral processes that give the interclavicle an anchor-like appearance (Ch. 512, State 1).

In taxa that possess lateral processes of the interclavicle, these lateral processes in ankyramorphan neoreptiles (e.g., *Delorhynchus*, OMNH 73515) curve strongly posteromedially, given an 'anchor-like' appearance in ventral or dorsal view (State 1).

UNAMBIGUOUS: Ankyramorpha + *Erpetonyx* (0 to 1).

**Ch. 513. Interclavicle, if lateral processes or wings are present, an anterior process extends between the clavicles: absent, or reduced to a small nub/mild convexity (0), present (1).**

Modified from Brocklehurst et al. (2022, Ch. 238).

**Ch. 514. Interclavicle, angle of head: low angle, interclavicle weakly curved in lateral view (0), head sharply upturned (1).**

**Ch. 515. Interclavicle, shape of posterior margin of head: distinctly offset from shaft by posterolateral emargination (0), grades gradually into shaft (1).**

Modified from Brocklehurst et al. (2022, Ch. 38).

**Ch. 516. Scapula, supraglenoid foramen: present (0), absent (1).**

Modified from Brocklehurst et al. (2022, Ch. 239).

**Ch. 517. Scapula, supraglenoid foramen, location: posterior to the apex of the supraglenoid buttress (0), located on or anterior to the apex of the buttress (1).**

Modified from Brocklehurst et al. (2022, Ch. 240).

**Ch. 518. Scapula, dorsoventral location of supraglenoid foramen: immediately dorsal to glenoid (0), ⅓-1/4 blade height dorsal to glenoid (1).**

Modified from Brocklehurst et al. (2022, Ch. 334).

**Ch. 519. Scapula, anterior margin: straight or slightly concave, at least dorsally (0), convex along entire length (1).**

Modified from Brocklehurst et al. (2022, Ch. 241).

**Ch. 520. Scapula, deep notch in anterior margin around midheight: absent (0), present (0).**

Modified from Brocklehurst et al. (2022, Ch. 334).

**Ch. 521. Scapula, scapulocoracoid notch on anterior margin on scapula / coracoid contact: absent (0), present (1).**

Modified from Brocklehurst et al. (2022, Ch. 242).

**Ch. 522. Scapula, acromion process: absent (0), present on the anterior surface of the scapula (1).**

**Ch. 523. Scapula, anterodorsal margin of scapular blade: bears a thin groove or articular surface for the cleithrum (0), anterior margin convex, lacking a groove, slot, or articular surface (1).**

This study.

The scapula of non-saurian amniotes bears a thin groove or articular surface for the cleithrum (State 0), whereas the scapula does not support a cleithrum in all known saurians other than possibly the Permian archosauromorph *Protorosaurus* (Ezcurra et al. 2016). We prefer this scoring over a simple presence/absence of the cleithrum character because this element is often disarticulated.

DELTRAN: *Protorosaurus speneri* (1 to 0), Weigeltisauridae (0 to 1), *Thadeosaurus colcanapi* (0 to 1), ACCTRAN: *Claudiosaurus* + Sauria (0 to 1).

**Ch. 524. Scapulocoracoid, glenoid shape: anteroposteriorly elongate and helical, lying along the external face and postaxial margin of the scapula (0), short, lies along the postaxial margin of the scapula (1).**

Modified from Brocklehurst et al. (2022, Ch. 330).

**Ch. 525. Scapula, anteroposterior breadth of distal end: broad (0), narrow (1).**

Modified from Brocklehurst et al. (2022, Ch. 331).

**Ch. 526. Scapula, anteroposterior breadth of proximal end (base): broad (0), pinched/narrow (1).**

Modified from Brocklehurst et al. (2022, Ch. 232).

**Ch. 527. Scapula, posterolateral surface of blade immediately dorsal to glenoid: deep, triangular concavity bounded anteriorly by prominent supraglenoid buttress (0), weakly concave (1), distinct supraglenoid buttress absent (2).**

Modified from Brocklehurst et al. (2022, Ch. 333).

**Ch. 528. Cleithrum, if present, dorsal extent of head: caps dorsal scapula (0), does not cap dorsal scapula (1). (-) in taxa lacking an ossified cleithrum.**

Derived from Huttenlocker et al. (2013, Ch. 191).

UNAMBIGUOUS: *Crassigyrinus* + *Greererpeton* (0 to 1), *Pantylus cordatus* (1 to 0). DELTRAN: *Dendrerpeton* + *Balanerpeton* (0 to 1), Amniota + Protorothyrididae + Diadectomorpha + Recumbirostra + Captorhinidae (0 to 1). ACCTRAN: *Acheloma* + *Cacops* (1 to 0), Temnospondyli + Reptiliomorpha + *Gephyrostegus* (0 to 1).

**Ch. 529. Cleithrum, if present, morphology of dorsal expansion: dorsal width greater than shaft (0), simple rod without or slight dorsal expansion (1). (-) in taxa lacking an ossified cleithrum.**

Derived from Huttenlocker et al. (2013, Ch. 192).

DELTRAN: Diadectomorpha + Protorothyrididae + Amniota (0 to 1).

**Ch. 530. Cleithrum, if present, postbranchial lamina: present (0), absent (1). (-) in taxa lacking an ossified cleithrum.**

Modified from Clack et al. (2012, Ch. 95).

The cleithrum of early tetrapodomorphs (e.g., *Whatcheeria* FMNH PR 1766) possesses a postbranchial lamina (State 0) that forms the posterior wall of the opercular chamber.

UNAMBIGUOUS: *Crassigyrinus scoticus* (0 to 1), Temnospondyli + Reptiliomorpha (0 to 1).

**Ch. 531. Coracoids, number of ossification centers: one (0), two (2).**

Modified from Brocklehurst et al. (2022, Ch. 244).

**Ch. 532. Coracoid, triceps process: absent or small (0), large (1).**

Modified from Brocklehurst et al. (2022, Ch. 245).

**Ch. 533. Coracoid, foramen on posterodorsal surface between glenoid and triceps process: absent (0), present (1).**

Modified from Brocklehurst et al. (2022, Ch. 336).

**Ch. 534. Coracoid (= procoracoid), anterior coracoid emargination: absent (0), present, emargination present and separated from the scapulocoracoid emargination by an anterior scapular ray (1).**

Modified from Pinna (1986, Ch. 56), Simões et al. (2022, Ch. 285).

DELTRAN: Lepidosauria (0 to 1). ACCTRAN: *Bellairsia* + Squamata (0 to 1).

**Ch. 535. Sternum: not mineralised (0), mineralized (1).**

Modified from Brocklehurst et al. (2022, Ch. 246).

**Ch. 536. Humerus, capitellum (radial condyle) and trochlea ( ulnar condyle): strongly developed as distinct ball-shaped and saddle-shaped structures (0), poorly developed but distinct from the ectepicondyle and entepicondyle (1), indistinct from the ectepicondyle and entepicondyle (2). ORDERED.**

Modified from Brocklehurst et al. (2022, Ch. 247).

**Ch. 537. Humerus, secondary ossification of epiphyses: absent (0), present, epiphyses remain as separate ossification centers until late ontogeny (1) .**

Derived from Benton (1985, Ch. X1)

UNAMBIGUOUS: Lepidosauromorpha (0 to 1).

**Ch. 538. Humerus, ridge connecting deltopectoral crest to head: double, paired ridge enclosing proximolateral fossa (deltopectoral crest anteroposteriorly expanded and tuberous (0) single, fossa is absent (1).**

Modified from Brocklehurst et al. (2022, Ch. 248).

**Ch. 539. Humerus, anterior surface of deltopectoral crest: weakly concave (0), strongly concave, bounded dorsally by a prominent, proximodistally elongate ridge (1).**

Modified from Brocklehurst et al. (2022, Ch. 340).

**Ch. 540. Humerus, position of latissimus dorsi attachment: proximal, adjacent to internal epicondyle (0), distal (1).**

Modified from Brocklehurst et al. (2022, Ch. 341).

**Ch. 541. Humerus, morphology of latissimus dorsi attachment: step-like transverse ridge or mound (0), prominent, posteriorly-directed tubercle (1).**

Modified from Brocklehurst et al. (2022, Ch. 342).

**Ch. 542. Humerus, diaphysis ( shaft): absent, distal and proximal heads merge and lack discrete shaft (0), distinct shaft present (1).**

Modified from Brocklehurst et al. (2022, Ch. 249).

**Ch. 543. Humerus, ventral surface of proximal end: extends proximally forming a low, anteroposteriorly oriented crest posteroventral to head (0), extends far proximally, forming a prominent crest (1).**

Modified from Brocklehurst et al. (2022, Ch. 339).

**Ch. 544. Humerus, torsion between proximal and distal ends: strong, 80 degrees or more (0), intermediate, 45-79 degrees (1), weak, less than 45 degrees (1). ORDERED.**

Modified from Muller (2004, Ch. 145, Ezcurra et al. (2016, Ch. 415).

DELTRAN: Embolomeri (2 to 1), Parapleurota (0 to 1), Crocopoda (1 to 2), Lepidosauromorpha  (1 to 0). ACCTRAN: Araeoscelidia (0 to 1), Archosauromorpha (0 to 2), Ankyramorpha  (0 to 1).

**Ch. 545. Humerus, ratio of width of distal head to humerus length : equal to or greater than 0.40 (0), less than 0.40 (1).**

Modified from Brocklehurst et al. (2022, Ch. 343).

**Ch. 546. Humerus, entepicondylar foramen: present (0), absent (1).**

**Ch. 547. Humerus, posterior surface of shaft around exit of entepicondylar foramen, if present : convex (0), exit foramen very large and rimmed by a longitudinal depression, foramen only enclosed by a narrow strip of bone (1).**

**Ch. 548. Humerus, entepicondyle foramen, if present, position: proximally, positioned proximal to or level with supinator/ectepicondylar groove, just distal to the humeral shaft (0) distally, located distal to ectepicondyle expansion or groove, just proximal to the trochlea (1).**

This study, based on observations in Spindler et al. (2019).

The entepicondylar foramen, when present, is located just distal to the humeral shaft in most taxa within this sampling (e.g., *Eomurruna* QMF 18335). In millerettids (e.g., *Milleretta*, BP/1/3822), the entepicondylar foramen is located more distal on the humerus, just proximal to the cochlea (State 1).

UNAMBIGUOUS: Mesosauridae + *Ascendonanus* + *Orovenator* + Parapleurota (0 to 1). DELTRAN: Squamata (0 to 1), Parapleurota (1 to 0). ACCTRAN: Rhynchocephalia (0 to 1).

**Ch. 549. Humerus, entepicondyle (ulnar epicondyle), transverse width: enlarged, makes up more than 2/3 of the distal transverse width (0), moderate makes up just less than half of transverse width of distal expansion (1), reduced or confluent with the ectepicondyle (2). ORDERED.**

Modified from Brocklehurst et al. (2022, Ch. 252).

**Ch. 550. Humerus, entepicondyle, distal extent: terminates proximal to the distal margin of the ulnar condyle (0), extends distally relative to ulnar condyle (1).**

This study.

The entepicondylar of weigeltisaurid neodiapsids (Buffa et al. 2021) terminates distally relative to the ulnar condyle (State 1).

UNAMBIGUOUS: Weigeltisauridae (0 to 1), *Eomurruna* + *Procolophon* + *Leptopleuron* (0 to 1).

**Ch. 551. Humerus, ventral surface (faces anteroventrally) of entepicondyle: flat or weakly convex (0), low, anteroproximally directed ridge on posterior margin (1).**

Modified from Brocklehurst et al. (2022, Ch. 344).

**Ch. 552. Humerus, ectepicondylar foramen: present (0), long supinator process, but epicondylar foramen not enclosed (1), absent, ectepicondylar groove not enclosed and supinator process proximodistally short (2), supinator process very low or absent (3). ORDERED.**

Modified from Brocklehurst et al. (2022, Ch. 253).

**Ch. 553. Humerus, supinator process: transversely broad and extends laterally or anterolaterally (0), transversely narrow and almost confluent with humeral shaft (1).**

Modified from Brocklehurst et al. (2022, Ch. 254).

**Ch. 554. Radius-humerus, length ratio: less than 0.68 (0), 0.68 to 0.82 (1), greater than 0.82 (2). ORDERED.**

Modified from Brocklehurst et al. (2022, Ch. 255).

**Ch. 555. Radius, length: shorter than ulna (0), longer than or approximately of the same length as the ulna (1).**

Modified from Brocklehurst et al. (2022, Ch. 256).

**Ch. 556. Radius, shape: straight (0), distinctly sigmoidal in lateral view (1).**

Modified from Brocklehurst et al. (2022, Ch. 257).

**Ch. 557. Ulna, olecranon process: absent or very low (0), prominent but lower than its transverse depth at base (1), strongly developed, being higher than its transverse depth at base (2). ORDERED.**

Modified from Brocklehurst et al. (2022, Ch. 2258).

**Ch. 558. Carpus, width to length ratio of radiale: subequal or less than 1.0 (0), greater than 1.0 (1).**

Modified from Brocklehurst et al. (2022, Ch. 259).

**Ch. 559. Carpus, ulnare proximodistal proportions: longer than wide, width less than 0.7 times length (0), short, width equal to or greater than 0.7 times length (1).**

Modified from Brocklehurst et al. (2022, Ch. 260).

**Ch. 560. Carpus, transverse width across carpus at widest point ( excluding pisiform): subequal to or greater than the length of the 4th metacarpal (0), less than 4th metacarpal (1),**

Modified from Brocklehurst et al. (2022, Ch. 261).

**Ch. 561. Manus, intermedium size: larger than medial centrale (0), smaller than medial centrale (1).**

Modified from Brocklehurst et al. (2022, Ch. 246).

**Ch. 562. Manus, intermedium, proximal articulation: contacts ulnare only (0), contacts ulnare and radius (1).**

**Ch. 563. Carpus, pisiform, if present, size: subequal to or smaller than the 4th distal carpal (0), conspicuously larger than the 4th distal carpal (1).**

Modified from Brocklehurst et al. (2022, Ch. 262).

**Ch. 564. Carpus, radiale: present as a distinct ossification (0), absent or unossified (1).**

**This study.**

The radiale is absent as a distinct ossification in mesosaurids and *Euparkeria* within this study.

UNAMBIGUOUS: Mesosauridae (0 to 1), *Euparkeria* (0 to 1). .

**Ch. 565. Carpus, radiale contacts distal carpal one: absent, contact is prevented by medial extension of the medial centrale (0), present, medial centrale is reduced allowing distal contact with DC1 (1). (-) in taxa lacking a radiale (Ch. 564).**

The radiale contacts distal carpal one due to reduction in the transverse width of the medial centrale in varanopine synapsids (e.g., *Varanops*, Romer and Price 1940), weigeltisaurid neodiapsids (e.g., *Coelurosauravus*, Buffa et al. 2022), and several archosauromorphs (e.g., *Prolacerta* UWBM 95529, *Azendohsaurus*, Nesbitt et al. 2015) within this sampling.

DELTRAN: Moradisaurinae + *Labidosaurus* + *Gansurhinus* + *Captorhinikos* + *Labidosaurikos* (1 to 0), *Azendohsaurus madagaskarensis* (1 to 0). ACCTRAN: Captorhinidae (1 to 0).

**Ch. 566. Carpus, medial centrale, contact with DC4: absent (0), present, medial centrale expanded, contacting DC4 (1).**

This study.

**Ch. 567. Metacarpus, length of McIV compared to radius: less than 45% of radius length (0), greater than 45% (1).**

Modified from Brocklehurst et al. (2022, Ch. 241).

**Ch. 568. Metacarpus, MC IV, length compared to that of MC III: longer (0), equal or shorter (10.**

Modified from Ezcurra (2016, Ch. 450).

**Ch. 569. Metacarpus, McV to McIV length ratio: greater than 0.65 (0), less than 0.65 (1).**

Modified from Brocklehurst et al. (2022, Ch. 264).

**Ch. 570. Manus, lateral centrale overlaps proximal surface of third distal carpal: no (0), yes (1).**

Modified from Brocklehurst et al. (2022, Ch. 345).

**Ch. 571. Manus, metapodial shape: long and slender, two-three times longer than maximal width (0), short and fat with small diaphysis (1).**

Modified from Brocklehurst et al. (2022, Ch. 347).

**Ch. 572. Manus, ungual ( terminal) phalanges, height to width ratio: low, ratio <1.1 and blood vessel grooves may be visible on the dorsolateral surface of the phalanx (0), high, ratio >1.5, strongly recurved and blood vessel grooves are located on the lateral surface of the phalanx (1).**

Modified from Brocklehurst et al. (2022, Ch. 265).

**Ch. 573. Manus, ungual ( terminal) phalanges, shape: hoof-like or bluntly spade- shaped distally with no ventral curvature (0), taper distally to a point, and curved ventrally (1).**

Modified from Brocklehurst et al. (2022, Ch. 266).

**Ch. 574. Manus, ungual phalanges, flexor tubercle: single bulbous eminence (0), paired, medial and lateral eminences (1), absent (2).**

Modified from Brocklehurst et al. (2022, Ch. 352).

**Ch. 575. Manus, digits, number or count: six or more (0),five (1), four or fewer (2). ORDERED.**

Huttenlocker et al. (2013, Ch. 218).

Most tetrapods possess 5 digits (State 1) within their manus (e.g., *Whatcheeria* FMNH PR 1816). Many microsaurs (e.g., *Pantylus* UT 4000-1) and temnospondyls (e.g., *Dendrerpeton*. NSM 987 GF 99.1) within this sampling possess four or fewer digits (State 0).

DELTRAN: *Dendrerpeton* + *Balanerpeton* + *Acheloma* + *Cacops* (1 to 0), Recumbirostra (1 to 0).
ACCTRAN: Recumbirostra (1 to 0), Temnospondyli (1 to 0).

**Ch. 576. Manus, phalanges in digit II: three (0), two (1).**

Modified from Maddin et al. (2008, Ch. 89,90,92).

The reduction in phalangeal count from three (State 0) phalanges to two (State 1) is considered here to be an autapomorphy of derived caseids (e.g., *Ennatosaurus*, Olson 1958) within this sampling.

DELTRAN: *Cotylorhynchus* + *Angelosaurus* + *Ennatosaurus* + *Euromycter* + *Oromycter* + *Casea* + *Datheosaurus* (0 to 1).

**Ch. 577. Manus, phalanges in digit lil: four (0), three (1), two (2). ORDERED.**

**Ch. 578. Manus, phalanges in digit IV: five (0), four or fewer (1).**

Modified from Brocklehurst et al. (2022, Ch. 349).

**Ch. 579. Manus, phalanges in digit V: three (0), two (1), four (2).**

Modified from Maddin et al. (2008, Ch. 89,90,92).

The manual phalangeal count is reduced to two (State 1) from the primitive three (State 0) in several procolophonians (e.g., *Deltavjatia,* Tsuji 2013) and caseids (e.g., *Cotylorhynchus,* Romer and Price 1940). The presence of four phalanges in digit V is considered here to be a unique autapomorphy of weigeltisaurid neodiapsids (e.g., *Coelurosauravus*, Buffa et al. 2022).

UNAMBIGUOUS: Pareiasauria + *Emeroleter* (0 to 1). DELTRAN: Caseidae (0 to 1), *Procolophon trigoniceps* (0 to 1), Weigeltisauridae + *Claudiosaurus* + Sauria (0 to 2). ACCTRAN: Procolophonidae (0 to 1), Weigeltisauridae (0 to 2).

**Ch. 580. Manus, phalanges, distal articular surface orientation: distal (0), ventrodistal (1).**

Modified from Brocklehurst et al. (2022, Ch. 351).

**Ch. 581. Pelvic girdle, acetabulum: sub-oval and shallow (0), subcircular and deep (1).**

Modified from Brocklehurst et al. (2022, Ch. 267).

**Ch. 582. Pubis, anteroposterior length relative to acetabulum: > 1.5 times (0), 1.0- 1.5 times (1).**

Modified from Brocklehurst et al. (2022, Ch. 268).

**Ch. 583. Pubis, ventrolateral surface of pubic apron: flat or convex, pubes extends ventromedially (0), strongly concave, pubes extends approximately medially (1).**

Modified from Brocklehurst et al. (2022, Ch. 269).

**Ch. 584. Pubis, pubic tubercle anteroventral to acetabulum: absent (0), present (1).**

Reworked from Brocklehurst et al. (2022, Ch. 270).

Rationale: The original framing of this character concerned both the presence and the morphology of the pubic tubercle. We split this character into its components: the presence of the pubic tubercle and the direction of its projection.

**Ch. 585. Pubis, morphology of pubic tubercle, if present: projects laterally (0), projects dorsally (1), pubic tubercle is weakly developed and is bounded by a laterally directed crest dorsally (2).**

Reworked from Brocklehurst et al. (2022, Ch. 270).

**Ch. 586. Pubis, midline symphysial contact: enlarged, dorsoventrally broad (0), subequal to height of ischial midline symphysis, restricted to peripheral margin of medial surface (1) .**

Modified from Brocklehurst et al. (2022, Ch. 253).

**Ch. 587. Pubis, morphology of obturator opening: fully-canalized foramen within the pubis (0), expressed as a foramen along the ischium-pubis suture (1).**

The position of the obturator foramen is highly variable amongst amniotes in this matrix. In most taxa herein, the obturator foramen fully pierces the pubis (State 0). In therapsids in this matrix, the obturator foramen is present along the ischia-pubic suture (State 1, e.g., *Titanophoneus*, Efremov 1938).

ACCTRAN: Therapsida (0 to 1).

**Ch. 588. Puboischiadic plate, ventromedial keel: absent (0), present, a raised ridge or keel present on the midline of the pubes and ischia (1).**

Modified from Li et al. (2018, Ch. 279).

The puboischiadic plate of pareiasaurs (e.g., *Embrithosaurus* SAM-PK-8034) possesses a strongly developed ventromedial keel along the ventral midline (State 1). This feature has been used classically to support a turtle-pareiasaur relationship (Li et al. 2018).

UNAMBIGUOUS: Pareiasauria + *Emeroleter* (0 to 1).

**Ch. 589. Puboischiadic plate, thyroid fenestra between pubis and ischium: absent (0), present (1)**

Modified from Pritchard and Sues (2019, Ch. 268).

The thyroid fenestra is present as an opening between the suture of the pubis and ischium and first appears in early crown saurians (e.g., *Bellairsia* NMS G.2022.1.1), although it is lost in most archosauromorphs crownward of tanystropheids (e.g., *Azendohsaurus*, Nesbitt et al. 2015).

DELTRAN: Lepidosauria + *Marmoretta* + *Megachirella* (0 to 1), *Trilophosaurus* + *Mesosuchus* (0 to 1). ACCTRAN: Tanystropheidae (0 to 1), Lepidosauromorpha (0 to 1).

**Ch. 590. Ilium, medial surface: weakly concave or flat (0) anteroventrally oriented ridge contacts pubic articulation (1).**

Modified from Brocklehurst et al. (2022, Ch. 271).

**Ch. 591. Ilium, mediolateral expansion on dorsal surface: present, and forms a trough or well-defined fossa (0), present, and forms a shelf with no discernable fossa (1), absent, ilium is plate- like with no mediolateral expansion along the dorsal surface (2). ORDERED.**

Modified from Brocklehurst et al. (2022, Ch. 272).

**Ch. 592. Ilium, main axis of the postacetabular process in lateral or medial view:mainly vertically orientated (0), posterodorsally oriented (1), mainly posteriorly oriented (2). ORDERED.**

Modified from Pritchard et al. (2015, Ch. 164, Ezcurra (2016, Ch. 464).

The postacetabular process of the ilium is posterodorsally directed (State 0) in most paleozoic tetrapods within this sampling (e.g., *Captorhinus aguti,* OMNH 44816). In weigeltisaurid neodiapsids (e.g., *Coelurosauravus,* Buffa et al. 2022), procolophonians (e.g., *Bunostegos*, MNN MOR33), and recumbirostrans (e.g., *Brachydectes* KUVP 49541), the ilium is oriented primarily vertically. The postacetabular process of archosauromorphs (e.g., *Mesosuchus*, SAM 6046) and sphenacomorph synapsids (Romer and Price 1940) is oriented primarily posteriorly.

DELTRAN: *Milleretta rubidge*i (0 to 2), Crocopoda (0 to 2), Mesenosaurinae (0 to 2), Recumbirostra (0 to 1). ACCTRAN: Ankyramorpha (0 to 1), Sphenacomorpha (0 to 2).

**Ch. 593. Ilium, postacetabular process, length: short, ends anterior to or level to posterior margin of iliac body (0), long, extends posterior to iliac body (1).**

This study.

The postacetabular process of the enigmatic *Hylonomus* (e.g., Carroll 1964), *Anthracodromeus* (Mann et. al 2021,recumbirostran ‘microsaurs’ (e.g., *Llistrofus*, Carroll and Gaskill 1978) and some procolophonians (e.g., *Eomurruna* QMF 18335) is poorly developed (State 0), not extending posterior to the iliac body.

UNAMBIGUOUS: Reptiliomorpha (1 to 0). DELTRAN: Diadectomorpha + Protorothyrididae + Amniota (0 to 1), Captorhinidae (0 to 1), Caseasauria (1 to 0), Ankyramorpha (1 to 0).

**Ch. 594. Ilium, preacetabular process or anterodorsal blade: absent or incipient, not extending past anteriormost extent of acetabulum (0) present, extending anterior to the acetabulum but not the ventral margin of the lium(1), extends anterior to the ventral margin of the ilium (2). ORDERED.**

Modified from Muller (2004, Ch. 67, Nesbitt (2011, Ch. 268), Ezcurra (2016, Ch. 460).

The preacetabular, or anterior, process of the ilium projects anterior to the acetabulum (State 1) and is a noted feature of Archosauromorpha (e.g., *Euparkeria,* SAM-PK-K5807) although it is absent in *Trilophosaurus* (TMM 31025-140) and some specimens of *Protorosaurus* (Gottmann-Quesada and Sander 2009).

UNAMBIGUOUS: Archosauromorpha (0 to 1). DELTRAN: Sphenacomorpha (0 to 1). Caseasauria (0 to 1), Pareiasauria + *Emeroleter* (0 to 1).

**Ch. 595. Ilium, anteroventral process: absent, anteroventral margin of ilium unexpanded (0), present, anteroventrally-oriented process extends from anterior margin of pubic peduncle (1).**

Modified from Benton (1985 Ch. J12) and Pritchard and Sues (2019 Ch. 269).

The anteroventral process of the ilium is an anteroventrally directed process that extends from the anterior margin of the pubic peduncle and is only present in lepidosauromorphs among the taxa sampled here (e.g., *Planocephalosaurus,* Fraser and Walkden 1984).

ACCTRAN: Lepidosauromorpha (0 to 1).

**Ch. 596. Ilium, posterodorsal surface of acetabulum: irregular, marked by posterodorsal invasion by finished bone (0), smooth, no posterodorsal invasion of bone (1).**

Modified from Benton (1985), Pritchard et al. (2015 Ch. 168), Ezcurra (2016, Ch. 469).

The acetabulum of early amniotes bears a posterodorsal invasion of bone making it appear irregular in lateral view (State 0, *Petrolacosaurus,* Reisz 1981). In contrast, the acetabulum of millerettids and neodiapsids is smooth and circular with no invasion by bone (State 1, *Youngina capensis*, Gow 1975).

DELTRAN: Parapleurota (0 to 1), Pareiasauria + *Emeroleter* (0 to 1). ACCTRAN: Total-group reptiles (0 to 1).

**Ch. 597. Ilium, dorsal and postacetabular processes: separate such that the dorsal margin of the ilium is bifurcate (0), confluent, dorsal margin of ilium smooth (1).**

Modified from Huttenlocker et al. (2013, Ch. 211), Pardo et al. (2017, Ch. 171).

The early tetrapod ilium is bifurcated, characterized by a dorsal ‘forking’ of the ilium caused by the possession of a posterior and dorsal process (State 0, e.g., *Proterogyrinus,* Holmes 1984). In contrast, the ilia of most temnospondyls and reptiliomorphs (e.g., *Dendrerpeton*, Holmes et al. 1998) possess a single posterior process and a smooth dorsal margin of the ilium (State 1).

UNAMBIGUOUS: Temnospondyli + Reptiliomorpha + *Gephyrostegus* (0 to 1), *Greererpeton burkemorani* (0 to 1).

**Ch. 598. Ischium, shape: slender, tapering posteriorly (0), expanded posterodorsally (1).**

Modified from Brocklehurst et al. (2022, Ch. 274).

**Ch. 599. Hindlimb-trunk, length ratio: hindlimb much shorter than trunk (0), hindlimb not less than 90% of the anteroposterior length of the trunk.**

Modified from Brocklehurst et al. (2022, Ch. 275).

**Ch. 600. Femur, distal articular surface: uneven, posterior (fibular) condyle projecting distinctly beyond the anterior (tibial) condyle (0), both condyles extend distally to approximately at same level (1).**

Modified from Brocklehurst et al. (2022, Ch. 276).

**Ch. 601. Femur, length to humerus ratio: greater than or equal to 1.2 (0), less than 1.2 (1).**

Modified from Brocklehurst et al. (2022, Ch. 277).

**Ch. 602. Femur, proportions: short and broad, maximum length less than three times distal width (0), long and slender, maximum length more than three times distal width (1).**

Modified from Brocklehurst et al. (2022, Ch. 278).

**Ch. 603. Femur, 4th trochanter (insertion of caudofemoralis muscle): present as a raised ridge on the ventral surface of the femoral shaft (0), absent (1).**

Modified from Brocklehurst et al. (2022, Ch. 279).

**Ch. 604. Femur, distal adductor crest on ventral surface: prominent (0), low and feebly developed (1), absent or reduced to a rugosity on the surface of the bone (2).**

Modified from Brocklehurst et al. (2022, Ch. 280).

**Ch. 605. Femur, anterior condyle: dorsoventrally thick, at least 0.6 of the dorsoventral depth of the posterior condyle (0), dorsoventrally compressed, approx. 0.5 or less of the dorsoventral depth of the posterior condyle (1).**

Modified from Brocklehurst et al. (2022, Ch. 281).

**Ch. 606. Femur, dorsal surface, intercondylar fossa: present, proximodistally extending depression separating anterior (medial) and posterior (lateral) condyles (0), absent or reduced, condyles not separated by fossa (1).**

This study.

The intercondylar fossa of the femur, located between the anterior and posterior condyles, is present in most early tetrapods in this matrix (e.g., *Limnoscelis* YPM 811). In contrast, the intercondylar fossa is reduced or absent in some neodiapsids (e.g., *Acerosodontosaurus*, MNHN.1908-32-57) and saurians (e.g., *Clevosaurus,* Fraser 1988).

DELTRAN: *Mesosaurus tenuidens* (0 to 1), *Claudiosaurus* + Sauria (0 to 1), *Acerosodontosaurus longipes* (0 to 1). ACCTRAN: *Claudiosaurus* + Sauria (0 to 1), Mesosauridae (0 to 1).

**Ch. 607. Femur, intertrochanteric fossa, depth: prominent and deep (0), shallow or absent (1).**

Modified from Brocklehurst et al. (2022, Ch. 358).

**Ch. 608. Femur, intertrochanteric fossa if present, proximodistal length: large, extends distally more than 20% of femur length (0), small, does not extend distally further than proximal end (1).**

This study.

In early tetrapods, the intertrochanteric fossa of the femur is large and extends distally more than 20% of the length of the femur (State 0). In many reptiles (e.g., *Araeoscelis,* Vaughn 1955), the intertrochanteric fossa is reduced, not extending further distally than the proximal end of the femur (State 1).

DELTRAN: Therapsida (0 to 1), Araeoscelidia (0 to 1), Reptilia (0 to 1), *Paleothyris acadiana* (0 to 1). ACCTRAN: *Apsisaurus witteri* (0 to 1), *Echinerpeton* + *Archaeothyris* (0 to 1), Dinocephalia + *Biseridens* (0 to 1), *Petrolacosaurus kansensis* (1 to 0), Synapsida (1 to 0).

**Ch. 609. Femur, morphology of head: terminal and anteroposteriorly elongate (0) subspherical (1).**

Modified from Brocklehurst et al. (2022, Ch. 355).

**Ch. 610. Femur, greater trochanter: absent (0), present (1).**

Modified from Brocklehurst et al. (2022, Ch. 356).

**Ch. 611. Femur, mound-like eminence on dorsal surface of proximal end: extensive, prominent and longitudinally elongate (0), small or absent (1).**

Modified from Brocklehurst et al. (2022, Ch. 357).

**Ch. 612. Femur, posterior longitudinal ridge located proximally on ventral surface: absent, internal fossa not enclosed posteriorly (0), present, enclosing posterior margin of internal fossa (1).**

Modified from Brocklehurst et al. (2022, Ch. 359).

**Ch. 613. Femur, posterior condyle, dorsal surface: convex (0), transversely concave, bearing longitudinal trough (1).**

Modified from Brocklehurst et al. (2022, Ch. 360).

**Ch. 614. Tibia, proximal end, anteriorly projected process beyond the articular portion for the femur ( = cnemial crest): low (0), prominent and distinct, straight (1), prominent and distinct, curved anterolaterally (2). ORDERED.**

Modified from Brocklehurst et al. (2022, Ch. 282).

**Ch. 615. Tibia, lower leg-foot length, ratio: shorter (0), articulated tibia + tibiale/astragalus longer than articulated 4th metatarsal + digit (1).**

Modified from Brocklehurst et al. (2022, Ch. 283).

**Ch. 616. Tibia, contact with lateral centrale (centrale 2) even if fused with astragalus: absent (0), present (1).**

Modified from Ford et al. (2021, Ch. 374).

The tibia contacts the pedal centrale in the archosauromorphs sampled here (e.g., *Prolacerta* AMNH FARB 9502). In some archosauromorphs, the single pedal centrale fuses or partially fuses with the astragalus during ontogeny (e.g., *Azendohsaurus*, Nesbitt et al. 2015), however, the centrale still contacts the tibia in these taxa.

Unambiguous: *Procolophon trigoniceps* (0 to 1), Archosauromorpha excluding *Protorosaurus* (0 to 1).

**Ch. 617. Fibula, transverse width at mid-length: subequal to or wider than transverse width of the tibia (0), distinctly narrower than transverse width of the tibia (1).**

Ezcurra (2016, Ch. 528).

The fibula in most early tetrapods is similar in transverse width to the tibia (State 0, e.g., *Heleosaurus* SAM-PK-K8305) whereas it is distinctly narrower than the tibia (State 1) in procolophonians (e.g., *Embrithosaurus,* SAM-PK-8034) and saurians (e.g., *Trilophosaurus*, TMM 31025-140).

UNAMBIGUOUS: Pareiasauria + *Emeroleter* (0 to 1), Procolophonoidea + *Nyctiphruretus* (0 to 1). DELTRAN: *Claudiosaurus* + Sauria (0 to 1). ACCTRAN: Leptopleuroninae + *Procolophon* (1 to 0), *Eichstaettisaurus schroederi* (1 to 0).

**Ch. 618. Tarsus, tibio-astragalar joint: flat (0), tibial ridge fits into astragalar groove (1).**

Modified from Brocklehurst et al. (2022, Ch. 284).

**Ch. 619. Tarsus, transverse width across tarsus at widest point: subequal to or greater than length of 4th metatarsal (0), less than 4th metatarsal (1).**

Modified from Brocklehurst et al. (2022, Ch. 285).

**Ch. 620. Tarsus, astragalus: absent (0), present (1).**

Modified from Brocklehurst et al. (2022, Ch. 286).

**Ch. 621. Tarsus, astragalus, if present, number of ossification centers: sutures or ossification centers delineate the tibiale, intermedium, and proximal centrale at some point during ontogeny (0) no evidence of multiple ossification centers remain (1). (-) in taxa that lack an astragalus (Ch. 620).**

Modified from Laurin and Reisz (1995 Ch. 115), MacDougall et al. (2019, Ch. 159), Cisneros et al. (2020, Ch. 157).

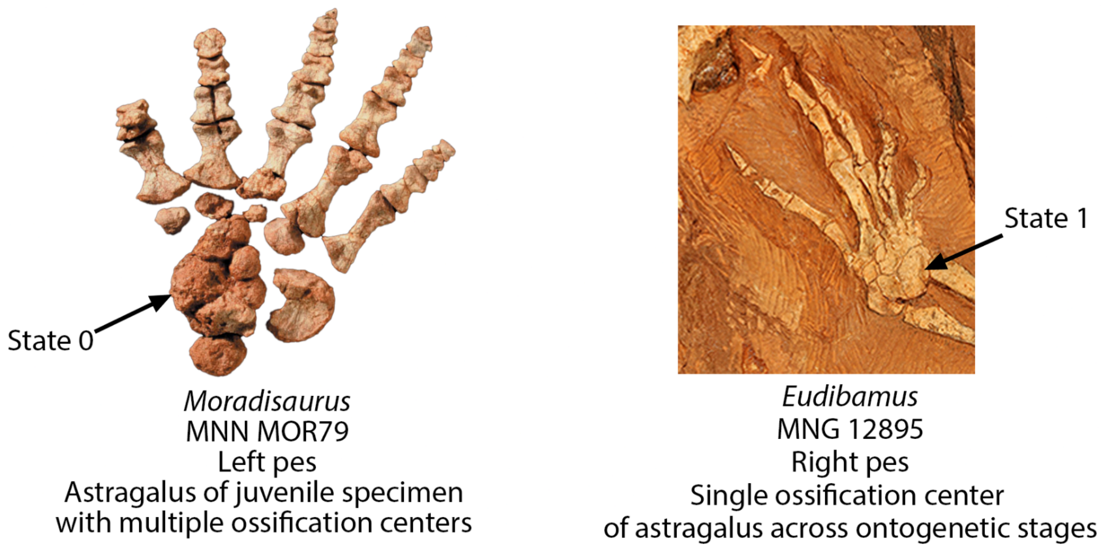


**Fig. S80.** Left, left pes of *Moradisaurus* (MNN MOR79) bearing an astragalus with multiple centers of ossification (Ch. 621, State 0). Right, right pes of *Eudibamus* (MNH 12895) bearing an astragalus with a single apparent ossification center (Ch. 621, State 1).

The astragalus, a feature once thought to be diagnostic of Amniota (e.g., Romer and Price 1940, Carroll 1968), is present in many ‘microsaurs’ and other stem-aminiotes as reviewed by Meyer and Anderson (2013). In early-diverging taxa possessing an astragalus, the astragalus often bears multiple ossification centers marking the early tetrapod tibiale, intermedium, and centrale (e.g., *Moradisaurus*, MNN MOR79, *Orobates*, Berman et al. 2004) and even ‘eureptiles’ such as *Hylonomus* (Meyer and Anderson 2013), although these centers are often only evident in juvenile taxa. However, there is no evidence of multiple ossification centers (State 1) of the astragalus in taxa approaching the reptile (e.g., Neoreptilia) or synapsid crown (e.g., therapsids, Romer and Price 1940), suggesting that the loss of multipartite development of the astragalus was independently acquired in several lineages of amniotes. We note here that future work, particularly on juvenile synapsids or captorhinids, that uses methodology such as histology or scanning techniques will clarify the evolution of this feature.

UNAMBIGUOUS: Amniota + Protorothyrididae + *Anthracodromeus* (0 to 1).

**Ch. 622. Tarsus, astragalus, proximal neck region: short (0), long, equal to or > 0.4 of the proximodistal length of astragalus (1).**

Modified from Brocklehurst et al. (2022, Ch. 287).

**Ch. 623. Astragalus, margin between tibial and fibular facet: margin grades smoothly into anterior hollow (0), prominent ridge separates margin from anterior hollow (1).**

**Ch. 624. Tarsus, calcaneum ( fibulare), proportions: length approximately equal to width (0), length conspicuously greater than width (1).**

Modified from Brocklehurst et al. (2022, Ch. 288).

**Ch. 625. Tarsus, calcaneum, lateral margin, expansion or "tuber": absent, lateral margin of calcaneum unexpanded (0), lateral margin of calcaneum bears a calcaneal tuber (1)**

Modified from Laurin (1991, Ch F9) and Ezcurra (2016, Ch. 545).

The calcaneum of crocopod archosauromorphs (e.g., *Mesosuchus*, SAM-PK-7416) bear a distinct lateral expansion or ‘calcaneal tuber’, probably for insertion of the gastrocnemius and peroneus longus musculature as in extant clades. (State 1).

DELTRAN: Tanystropheidae + Crocopoda (0 to 1). ACCTRAN: Crocopoda (0 to 1).

**Ch. 626. Calcaneum-astragalus, articulation. (-) in taxa with a fused astragalocalcaneum : concavoconvex with concavity on the calcaneum (0), roughly flat (1), concavoconvex with concavity on the astragalus (2).**

Modified from Ezcurra (2016, Ch. 532). Ordered.

UNAMBIGUOUS: CROCOPODA (0 TO 2).

**Ch. 627. Tarsus, notch or trough for perforating foramen: on astragalus only (0), on the astragalus and calcaneum (fibulare) (1), perforating foramen absent (2).**

Modified from Brocklehurst et al. (2022, Ch. 289).

**Ch. 628. Tarsus, perforating foramen, if present, proximodistal location: located distally, on distal third of astragalus or calcaneum (0), located at proximodistal midlength of astragalus and/or calcaneum (1). (-) in taxa that lack an astragalus (Ch. 620).**

This study.

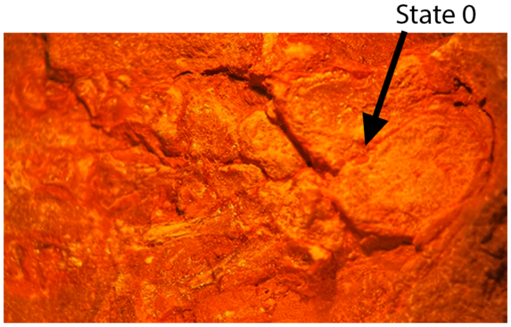


*Mycterosaurus*
AMNH FARB 7002
Left pes
Perforating foramen located distally

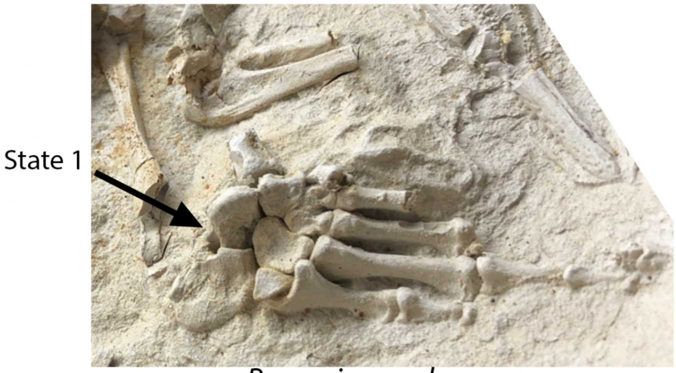


*Broomia perplexa*
NHMUK 4065
Right pes
Perforating foramen located proximally

**Fig. S81.** Left, left pes of *Mycterosaurus* (AMNH FARB 7002) bearing a distally-located perforating foramen of the tarsus (Ch. 628, State 0) and right, right pes of *Broomia* (NHMUK 4065) bearing a centrally positioned perforating foramen (Ch. 628, State 0).

The perforating foramen of the pes is located distally (State 0) on the astragalus and calcaneum articulation in many near- or early amniotes (e.g., *Captorhinus* OMNH 44816 or *Paleothyris* MCZ 3481) and is considered here to be the primitive condition in Amniota. In contrast, the perforating foramen is located near the midlength of this articulation in millerettid (e.g., *Broomia,* NHMUK 4065) and neodiapsid reptiles (e.g., *Hovasaurus*, MNHN.1908-32-68).

UNAMBIGUOUS: Parapleurota (0 to 1). ACCTRAN: Mesosauridae (0 to 1).

**Ch. 629. Tarsus, proximal tarsals, fusion between astragalus and calcaneum: absent (0), present (1). (-) in taxa that lack an astragalus (Ch. 620).**

Modified from Lee (1995, Ch. G5).

The astragalus and calcaneum are co-ossified and fused into a single unit (State 1) in many procolophonians (e.g., *Saurodektes* BP/1/4195, *Emeroleter* KPM uncatalogued) and lepidosauromorphs (e.g., *Clevosaurus*, Fraser 1988) sampled here.

DELTRAN: *Procolophon trigoniceps* (1 to 0), *Eomurruna yurrgensis* (1 to 0), Ankyramorpha (0 to 1), Lepidosauria + *Marmoretta* + *Megachirella* (0 to 1). ACCTRAN: Procolophonidae (1 to 0), Ankyramorpha (0 to 1), Lepidosauromorpha (0 to 1).

**Ch. 630. Tarsus, first distal tarsal: present (0), absent as a discrete ossification (1).**

Modified from Brocklehurst et al. (2022, Ch. 290).

**Ch. 631. Tarsus, fourth distal tarsal, proximal surface : smooth or weakly convex contact for proximal tarsals (0), prominent process for contact with proximal tarsals (1), distinct, proximally raised region on the posterior portion (= heel of Sereno & Arcucci, 1994) (2).**

Modified from Sereno & Arcucci (1994).

**Ch. 632. Tarsus, fourth distal tarsal, articulates with MTV: absent, fourth distal tarsal restricted to contact with digit four (0), present, fourth distal tarsal articulates with MT IV and V (1).**

This study.

**Ch. 633. Tarsus, fifth distal tarsal: present (0), absent (1).**

Modified from Brocklehurst et al. (2022, Ch. 290).

**Ch. 634. Tarsus, number of centralia when astragalus absent: two discrete centralia centers of ossification present throughout ontogeny (0), only one centralia ossification present throughout ontogeny (1), discrete ossification of centralia absent (2). ORDERED.**

Modified from Brocklehurst et al. (2022, Ch. 290).

**Ch. 635. Tarsus, lateral centralia, if present, contact with distal tarsal 1: absent (0), present (1).**

This study.

**Ch. 636. Metatarsus, ratio of lengths of metatarsals I and IV: equal or greater than 0.42 (0), less than 0.42 (1).**

Modified from Brocklehurst et al. (2022, Ch. 292).

**Ch. 637. Metatarsus, metapodials overlapping proximally: absent (0), present (1).**

Modified from Brocklehurst et al. (2022, Ch. 293).

**Ch. 638. Metatarsus, fourth metatarsal: short (0), long, at least 40% of digit IV (1).**

Modified from Brocklehurst et al. (2022, Ch. 294).

**Ch. 639. Metatarsus, metatarsal V, length compared to that of metatarsal III: equal to or longer than .50 (0), metatarsal V shorter than .49 length of MT III (1**).

Modified from Ezcurra et al. (2020) Ch. 823

**Ch. 640. Metatarsus, Metatarsal V, lateral plantar tubercle: absent (0), present (1).**

Modified from Ezcurra, Scheyer & Butler (2014,Ch. 218), Ezcurra (2016, Ch. 580)

The fifth metatarsals of saurian reptiles often possess one or two plantar tubercles for the insertion of peroneus longus, peroneus brevis, and gastrocnemius musculature (Lee, 1997). Of these tubercles, the lateral plantar tubercle is present (State 1) in most early saurian reptiles (e.g., *Prolacerta*, AMNH FARB 9502). In contrast, the lateral plantar tubercle is absent (State 0) in most early amniotes and lost in Archosauriformes sampled here (e.g., *Proterosuchus,* Ezcurra 2016).

Lepidosauria + *Marmoretta* + *Megachirella* (0 to 1), Archosauriformes (0 to 1), Trilophosaurus + Mesosuchus (0 to 1), *Coelurosauravus elivensis* (0 to 1). ACCTRAN: Archosauriformes (1 to 0), *Protorosaurus speneri* (1 to 0), Claudiosaurus germaini (1 to 0), Weigeltisauridae + *Claudiosaurus* + Sauria (0 to 1).

**Ch. 641. Metatarsus, Metatarsal V, medial plantar tubercle: absent (0), present (1).**

Modified from Ezcurra, Scheyer & Butler (2014, Ch. 218), Ezcurra (2016, Ch. 580)

The medial plantar tubercle is here considered to be an autapomorphy of Sauria, being present (State 0) in lepidosauromorphs (e.g., *Bellairsia* NMS G.2022.1.1) and early archosauromorphs (e.g., *Macrocnemus,* PIMUZ T 2477). However, the medial plantar tubercle is lost or absent (State 1) in many members of Archosauromorpha (e.g., *Euparkeria*, SAM-PK-K5807).

DELTRAN: Lepidosauria + *Marmoretta* + *Megachirella* (0 to 1), *Macrocnemus bassanii* (0 to 1). ACCTRAN: SAURIA (0 to 1), *Protorosaurus* (1 to 0).

**Ch. 642. Metatarsal V distal shaft, angle relative to proximal tarsal articulation: straight, with proximal tarsal articulation forming a straight line with primary shaft (0) 'hooked' with proximal tarsal articulation forming a right angle with the primary shaft (1).**

Modified from Pritchard and Nesbitt (2017, Ch. 197).

The distal shaft of MTV in most taxa herein is straight, with the proximal articulation in line with the rest of the shaft (State 0). In most early saurian reptiles (but not all, e.g., *Protorosaurus* Gottmann-Quesada and Sander 2009), the fifth metatarsal is 'hooked' in appearance (e.g., *Trilophosaurus*, TMM 31025-140) due to the proximal tarsal articulation forming a right angle with the shaft.

DELTRAN: *Claudiosaurus* + Sauria (0 to 1). ACCTRAN: Sauria (0 to 1).

**Ch. 643. Metatarsal V, distal process, dorsiflexion: absent, distal process straight (0), present, distal process angles dorsally near proximodistal midpoint (1).**

**Ch. 644. Metatarsus, metatarsal V, expansion of proximal end: absent, proximal end of MTV is smooth and unexpanded (0), 1 present, proximal end of MTV is expanded proximally relative to the articulation with the tarsal 4 or 5 (1).**

This study.

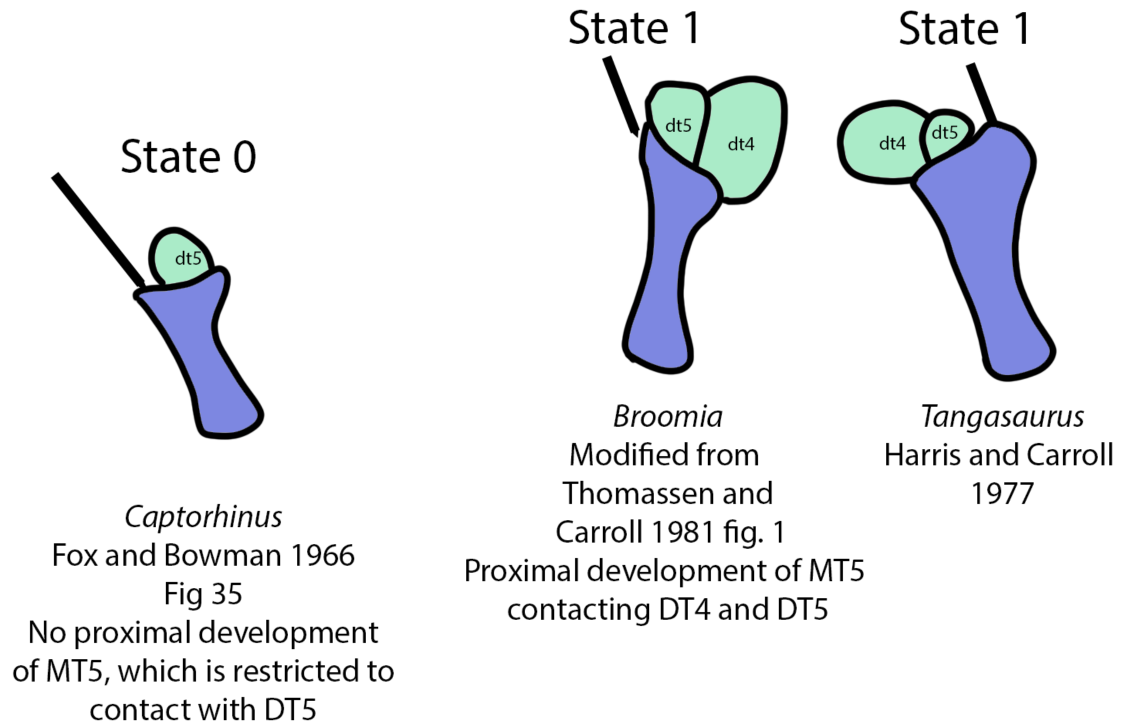


**Fig. S82.** Left, 5th metatarsal of *Captorhinus* (Modified from Fox and Bowman 1966) which bears no proximal expansion (Ch. 644, State 0). Right, 5th metatarsals of *Broomia* (Modified from Thomassen and Carroll 1981) and *Tangasaurus* (Modified from Harrison and Carroll 1977) which bear a proximal expansion contacting the 5th distal tarsal (Ch. 644, State 1).

The fifth metatarsal of millerettid (e.g., *Broomia*, NHMUK 4065) and neodiapsid (e.g., *Kenyasaurus* KNM-MA1) reptiles bear a proximal expansion relative to the articulation with the 5th distal tarsal (or 4th distal tarsal in taxa lacking an ossification of the 5th) (State 1). In extant taxa, this process marks the insertion of the brevis branch of the peroneus muscle.

UNAMBIGUOUS: Parapleurota (0 to 1) DELTRAN: Weigeltisauridae + *Claudiosaurus* + Sauria (1 to 0)

**Ch. 645. Metatarsus, metatarsal V, proximal surface, if proximal expansion present, postaxial margin: marked by a prominent, pointed process (outer process sensu Robinson, 1975) (0), with a posterolateral flange framing the 4th or 5th distal tarsal laterally (1). Inapplicable (-) in taxa lacking a proximal expansion of MTV (Ch. 651).**

Modified from Pritchard and Nesbitt (2017, Ch. 196).

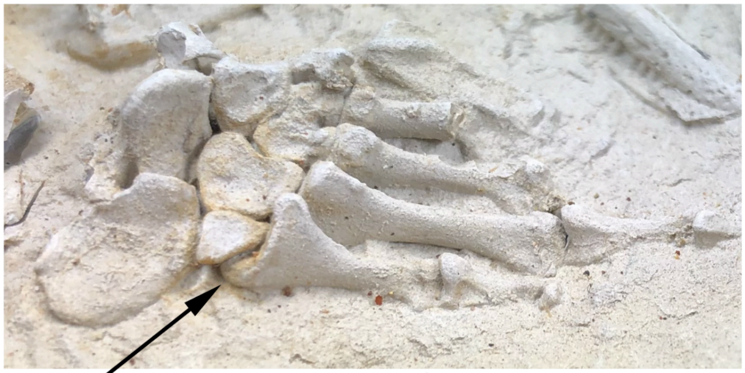


**Fig. S83.** Right pes of *Broomia* (NHMUK 4065) with a posterior flange of MTV (Ch. 645, State 1).

In most neodiapsid reptiles, the proximal expansion of the fifth metatarsal does not contribute to the distal tarsal articulation (State 0), and is instead commonly referred to as the 'outer process' *sensu* Robinson (1975). In contrast, the proximal development of MTV in millerettid reptiles (e.g., *Broomia,* NHMUK 4065) and the early neodiapsid *Saurosternon* (Carroll 1975), contributes to the articulation with the 5th or 4th distal tarsal (State 1).

UNAMBIGUOUS: Neodiapsida (1 to 0).

**Ch. 646. Pes, digits I and II, length proximal phalanges relative to metatarsals: proximal phalanges equal to or longer than 70% the length of their respective metatarsal (0), metatarsals longer than proximal phalanges (1).**

This study.

**Ch. 647. Pes, pedal unguals of digits II-IV: much shorter than all non-ungual phalanges (0), subequal to at least one non-ungual phalanx of the same digit (1), longer than all non-ungual phalanges of the same digit (2). ORDERED.**

Modified from Pritchard and Nesbitt (2017, Ch. 222).

The pedal unguals of early tetrapodomorphs (e.g., *Greererpeton,* Smitshon 1989) are noticeably smaller than non-ungual phalanges, a feature also present in recumbirostrans (e.g., *Pantyus,* Carroll 1968) and *Hylonomus* (Carroll 1974), which is in contrast with most early amniotes in which the unguals are approximately equal to the length of the non-ungual phalanges. The pedal unguals of many procolophonians (e.g., *Deltavjatia,* Tsuji 2013) and early crocopod archosauromorphs (e.g., *Azendohsaurus*, Nesbitt et al. 2015) are more elongated than the non-ungual phalanges in the same digit (State 2).

Unambiguous: Captorhinidae + Diadectomorpha + Protorothyridae + Amniota (0 to 1). DELTRAN: *Dimetrodon milleri* (1 to 2), *Pantelosaurus saxonicus* (1 to 2), *Eomurruna yurrgensis* (2 to 1), *Nyctiphruretus acudens* (2 to1), Archosauriformes (1 to 2), Mesosauridae (1 to 0), *Orobates pabsti* (1 to 2)

### 9. Observed specimens

90% of OTUs were directly observed by the authors of this study. The following lists specimens that we have directly observed for the purpose of this study, not simply specimens we have observed for other purposes. For example, we have observed several specimens of *Baphetes* but not for phylogenetic purposes, so we do not count that taxon as being directly observed for these purposes. Method concerns whether we observed the specimens in-person, via scans, or both.

| Taxon | Observed Specimens | Method |
|---|---|---|
| *Acerosodontosaurus piveteaui* | MNHN.F.MAP359a (holotype) | In-person |
| *Acheloma cumminsi* | AMNH FARB 4205 (holotype), FMNH UC 1584 (referred specimen) | In-person, CT |
| *Acleistorhinus pteroticus* | FMNH UR 1038 (holotype) | In-person, CT |
| *Aerosaurus wellesi* | UCMP 40096 (holotype), UCMP 35762 and UCMP 40097 (referred specimens) | In-person, CT |
| *Angelosaurus romeri* | FMNH UR 827 (holotype), FMNH UR 828, FMNH UR 844 through FMNH UR 854 (referred specimens) | In-person |
| *Anthracodromeus longipes* | AMNH FARB 6940 (holotype), CM 81532 and 25282 (referred specimens) | In-person |
| *Apsisaurus witteri* | MCZ 1474 (holotype) | In-person |
| *Araeoscelis* spp. | *A. grandis* FMNH UR 341 (holotype), FMNH UR 659 through FMNH UR 662 (syntypes). *A. casei* AMNH 4685 (holotype), MCZ 4380, MCZ 8828 (referred specimens) | In-person, CT of *A. casei* |
| *Archaeothyris florensis* | MCZ 4079 (holotype), MCZ 4080:4086 (paratypes) | In-person |
| *Archeria crassidiscus* | AMNH 4550 (Type), AMNH 4123, 4551:4555, | In-person |

| | | |
|---|---|---|
| | 23588:23620, MCZ 2052 (referred specimens) | |
| *Ascendonanus nestleri* | Taxon scored based on description by Spindler et al. (2018) | |
| *Australothyris smithi* | SAM-PK-K8392 | In-person, Synchrotron µCT |
| *Azendohsaurus madagaskarensis* | FMNH PR 2751 (paratype) and other FMNH specimens of this taxon, UA-7-20-99-653 (holotype) | In-person |
| *Balanerpeton woodi* | Taxon scored based on (in part) description in Milner and Sequeira (1994). | |
| *Baphetes* spp. | NHMUK R.4056/R.4183 (holotype) and description of referred specimens by Milner et al. 2009 | In-person |
| *Batropetes niederkirchensis* | Based on description of Glienke (2015) | |
| *Belebey vegrandis* | Photos of SGU 104/B-2020 and SGU 104/B-2021 provided by C. Kammerer | In-person and photographs provided by C. Kammerer |
| *Bellairsia gracilis* | NHMUK R12678 (holotype), NMS G.2022.1.1, NHMUK PV R12678; NMS G.2019.34.1; NMS G1992.47.10 | In-person, Synchrotron µCT |
| *Biarmosuchus tener* | PIN 1785/2 (holotype); 1758/8; 1758/34; 1758/86; (+ exhibited specimens) | In-person |
| *Biseridens qilianicus* | Taxon scored based on description by Li and Cheng (1997) and Liu et al. (2010) | |
| *Brachydectes newberryi* | AMNH FARB 6941 (holotype), USNM V 4509, USNM V 4477, AMNH FARB 6961, FARB 4757, various other AMNH referred specimens. | In-person, CT |
| *Bolosaurus major* | AMNH FARB 4461 | In-person, CT |

| | | |
|---|---|---|
| *Broomia perplexa* | NHMUK PV R. 4065 (holotype), BP/1/6222 (referred specimen) | In-person |
| *Brouffia orientalis* | Taxon scored based on description by Klembara et al. (2024) | |
| *Bunostegos akokanensis* | MNN MOR72 (holotype), MNN MOR25, MNN MOR34, MNN MOR 37, MMN MOR 47, MNN MOR86, MNN MOR101, MNN MOR104 (referred specimens) | In-person |
| *Cabarzia trostheidei* | Taxon scored based on description by Spindler et al. 2019 | |
| *Cacops morrisi* | OMNH 53077 (holotype), OMNH 73206,OMNH 53073 (referred specimens) | In-person |
| *Callibrachion gaudryi* | MNHN.F.AUT490 (holotype_ | In-person |
| *Candelaria barbouri* | DGM 314R (holotype) | Photos provided by J. Cisneros |
| *Captorhinikos valensis* | FMNH UR 101 (holotype), FMNH UR 101-108 (referred specimens) | In-person |
| *Captorhinus aguti* | AMNH 4333 (holotype), KUVP 9978, USNM PAL 606433, USNM PAL 619436, USNM PAL 619437, USNM PAL 619440_3 | In-person, CT |
| *Captorhinus laticeps* | FMNH UC 642 (holotype), FMNH UC 1698, FMNH UC 701 (referred specimens) | In-person, CT |
| *Cardiocephalus sternbergi* | AMNH FARB 3892 (holotype) | In-person |
| *Carrolla craddocki* | TMM 40031-54 (holotype) | In-person, scan |
| *Casea broilii* | FMNH UC 656 (holotype), FMNH UC 698, FMNH UC 1011, and FMNH mount | In-person |
| *Chroniosaurus dongusensis* | THUg 3308, PIN 3585/124, | In-person |

| | | |
|---|---|---|
| | PIN 105B/1097 | |
| *Claudiosaurus germaini* | MNHN 978-6-1, MNHN 1978-6-1, 1978-6-2 | In-person |
| *Clevosaurus hudsoni* | NHMUK PV R36832 (referred specimen) | In-person, CT |
| *Coelostegus prothales* | Taxon scored based on the description of Klembara et al. (2023) | |
| *Coelurosauravus elivensis* | MNHN.F.MAP325a (lectotype), MNHN 1908-5-2 | In-person |
| *Colobomycter pholeter* | FMNH 272 (holotype), BMRP2008.3.1 (referred specimen) | In-person |
| *Cotylorhynchus bransoni* | FMNH UR 835 (holotype), FMNH UR 837, FMNH UR 835, FMNH UR 841, FMNH UR 910 | In-person |
| *Cotylorhynchus hancocki* | FMNH UR 154 (holotype), FMNH UR 581, FMNH UR 622,FMNH UR 703 | In-person |
| *Cotylorhynchus romeri* | USNM V 21317, supplemented by description of Olson 1968 and Reisz et al.'s (2022) description of OMNH 04329 | In-person |
| *Crassigyrinus scoticus* | NHMUK VP R310, NHMUK PV OR30532, NHMUK VP R10000 | CT |
| *Cutleria wilmarthi* | USNM 22099 (holotype) | In-person |
| *Datheosaurus macrourus* | MB.R. 1015.1 (holotype) | In-person |
| *Delorhynchus cifellii* | OMNH 73515 (holotype) | In-person, nCT |
| *Deltavjatia rossica* | PIN 2212/2 (holotype) | In-person |
| *Dendrerpeton helogenes* | NSM 987 GF 99.1 | CT |
| *Dimetrodon limbatus* | AMNH FARB 2275; AMNH | In-person |

| | | |
|---|---|---|
| | FARB 4081, AMNH FARB 25888, FARB 25886, FARB 25884, FARB 25860, FMNH UC 114; FMNH UC 540; FMNH UC 804; FMNH UC 813; FMNH UC 1001 | |
| *Dimetrodon milleri* | MCZ 1365 (holotype), AMNH FARB 4126, AMNH FARB 4290 | In-person |
| *Dvellecanus carrolli* | UCMP 202940 (holotype) | In-person, CT |
| *Echinerpeton intermedium* | RM 206866 (referred specimen) | In-person |
| *Edaphosaurus boanerges* | MCZ 1761, MCZ 1680, AMNH FARB 7131, FARB 7129, FARB 7128, | In-person |
| *Edaphosaurus novomexicanus* | FMNH UC674 (holotype) | In-person |
| *Edops craigi* | AMNH FARB 7614, USNM V 23309 (referred specimens), MCZ 1378 | In-person |
| *Eichstaettisaurus schroederi* | Taxon scored based on description by Simões et al. 2018 | |
| *Elliotsmithia longiceps* | TM-1483 (holotype) | In-person |
| *Embrithosaurus schwarzi* | SAM-PK-8034 (holotype), | In-person |
| *Emeroleter levis* | PIN 2212/92 (holotype), PIN 2212/14 (referred specimens) | In-person,CT scanned Bazzana-Adams et al. (2024). |
| *Ennatosaurus tecton* | PIN 1580/17 (holotype), PIN 1580/101, PIN 1580/106, PIN 1580/107 | In-person |
| *Eocasea martini* | KUVP 9616b (holotype) | In-person |
| *Eomurruna yurrgensis* | Taxon scored based on description by Hamley et al. (2020) | |
| *Eothyris parkeyi* | MCZ 1161 (holotype) | In-person |
| *Erpetonyx arsenaultorum* | ROM 55402 (holotype) | In-person |
| *Euconcordia cunninghami* | KUVP 87102A + B | In-person |

| | | |
|---|---|---|
| | (holotype), KUVP 96164A + B (referred specimen) | |
| *Eudibamus cursoris* | MNG 8852 (holotype), MNG 12895 (referred specimen) | Photographs provided by A. Henrici |
| *Euparkeria capensis* | SAM-PK-5867 (holotype), SAM-PK-6047A&B, SAM-PK-6050, SAM-PK-7699-7710, SAM-PK-7696, SAM-PK-13665, SAM-PK-13666. | In-person |
| *Euromycter rutenus* | MNHN.F.MCL-2 (holotype) | In-person |
| *Euryodus dalyae* | FM UR 2296 (holotype) | In-person, nCT |
| *Feeserpeton oklahomense* | OMNH 73541 (holotype) | In-person, CT |
| *Gansurhinus qingtoushanensis* | Taxon scored by the description of Reisz et al. (2011) | |
| *Gephyrostegus bohemicus* | AMNH FARB 11546 (latex peel of holotype) | In-person |
| *Glaucosaurus megalops* | FMNH UC 691 (holotype) | In-person |
| *Gordodon kraineri* | Specimen scored based on description by Lucas et al. (2018) | |
| *Greererpeton burkemorani* | CMNH 10931 (holotype), USNM V 22576, USNM PAL 419694 | In-person |
| *Hapsidopareion lepton* | FMNH 2303 (holotype), FMNH 2304 and 2305. | In-person,CT |
| *Haptodus garnettensis* | ROM 14156 (holotype), ROM 29872, ROM 30099, ROM 43601; ROM 43602: 43608 | In-person |
| *Heleosaurus scholtzi* | SAM-PK-K8305 (referred specimen) | In-person, Synchrotron µCT |
| *Hovasaurus boulei* | MNHN 1925-5-34 (paratype), MNHN.F.MAP373 (referred specimen with braincase), OUMNH | In-person |

| | | |
|---|---|---|
| | PAL-GX.00071/4/5/6/85) | |
| *Huehuecuetzpalli mixtecus* | Taxon scored based on description by Reynoso (1998) | |
| *Huskerpeton englehorni* | USNM 32144 (holotype) | CT |
| *Hylonomus lyelli* | NHMUK R.4168 (holotype), NHMUK R.4167, NHMUK R.445 | In-person, Synchrotron µCT |
| *Hypsognathus fenneri* | YPM 55831 | In-person |
| *Ianthodon schultzei* | KUVP 133735 (holotype) | In-person |
| *Ianthasaurus hardestiorum* | KUVP 69035 (holotype), ROM 29941 and ROM 29942 (referred specimens) | In-person |
| *Labidosaurikos meachami* | FMNH UR 1031 (referred specimen), FMNH UR 380 (cast of type) | In-person |
| *Labidosaurus hamatus* | AMNH FARB 4341; FMNH UC 634, FMNH UR 161, FMNH UR 273, FMNH UR 634, FMNH UR 643, FMNH UR 696, FMNH UR 727, FMNH UR1198, FMNH UC 1543, FMNH P 12758 | In-person |
| *Lanthanolania ivakhnenkoi* | Taxon scored on description by Modesto and Reisz (2003) | |
| *Lanthanosuchus watsoni* | Cast of holotype | In-person |
| *Leptopleuron lacertinum* | RSM 1891.92.528 (holotype), NHMUK R4779, NHMUK R3919, NHMUK R3917 | In-person |
| *Limnoscelis paludis* | YPM 811 (holotype), FMNH UC 676. | In-person |
| *Lupeosaurus kayi* | AMNH FARB 4006, FARB 4024, MCZ 1455 (holotype). | In-person |
| *Macrocnemus bassanii* | PIMUZ T 2477 | Synchrotron µCT |
| *Macroleter poezicus* | PIN 4353/3, PIN uncatalogued | In-person |
| *Marmoretta oxoniensis* | NMS G1992.47.1a–b, | In-person, Synchrotron µCT |

| | | |
|---|---|---|
| | CAMSM X9991 (referred specimens), NHMUK R.12025, NHMUK R.12026, NHMUK R.12027, NHMUK R.12028, NHMUK 12400-12406, NMS G 1992.47.1, NMS G 1992.47.4, NMS G 1992.47.5. | |
| *Megachirella wachtleri* | PZO628 (holotype) | In-person, CT, Synchrotron µCT |
| *Mesenosaurus romeri* | PIN 3706/4, PIN 3717/1 | In-person |
| *Mesosaurus tenuidens* | BP/1/4332, BP/1/4371, BP/1/4372, BP/1/4715,4719: 4725, AMNH 23809, AMNH 23810, AMNH 23813 | In-person |
| *Mesosuchus browni* | SAM-PK-K 5882 (holotype), SAM-PK-K6536, SAM-PK-K6046, SAM-PK-K7416 | In-person |
| *Milleretta rubidgei* | RC 14 (holotype), R.C. 70, BP/1/3818,BP/1/3821, BP/1/3822, BP/1/2876, NMQR 3317 (Referred specimens) | In-person, CT, Synchrotron µCT |
| *Milleropsis pricei* | BP/1/720 (holotype), BP/1/4203 (referred specimen) | In-person, CT, Synchrotron µCT |
| *Millerosaurus nuffieldi* | RC 78 (holotype) | In-person |
| *Moradisaurus grandis* | MNN MOR149, MNN MOR 150, MNN MOR78, MNN MOR79 | In-person |
| *Mycterosaurus longiceps* | FMNH UC 692 (holotype), FMNH UC 169, AMNH FARB 7002 (referred specimens) | In-person |
| *Nannaroter mckinziei* | OMNH 73107 (holotype) | CT |
| *Nochelesaurus alexanderi* | SAM-PK-6239 (holotype), | In-person |

| | | |
|---|---|---|
| | SAM-PK-6238, SAM-PK-6239A, | |
| *Nyctiphruretus acudens* | BP/1/5622 (referred specimen) | In-person |
| *Oedaleops campi* | UCMP 35758 (holotype) | In-person, CT |
| *Ophiacodon* spp. | AMNH FARB 4143 (neotype of *O. uniformis*) | In-person and based on descriptions by Romer and Price (1940) |
| *Opisthodontosaurus carrolli* | Taxon scored based on description of Reisz et al. 2015 | |
| *Orobates pabsti* | MNG 10181 (holotype), MNG 8980 (paratype) | CT, additional photographs provided by A. Henrici |
| *Oromycter dolesorum* | FMNH PR 2281 (holotype), FMNH PR 2283:2287 (referred specimens) | In-person |
| *Orovenator mayorum* | OMNH 74606 (holotype), OMNH 74607 (referred specimen) | In-person, CT |
| *Paleothyris acadiana* | MCZ 4381 (holotype), MCZ 4382, MCZ 4384 (referred specimens) | In-person |
| *Paliguana whitei* | AM 3585 (holotype) | In-person, Synchrotron µCT |
| *Pantelosaurus saxonicus* | RS 14752 (holotype) | In-person |
| *Pantylus cordatus* | MCZ 2040 (holotype), UT-4001 (referred specimens) | In-person |
| *Petrolacosaurus kansensis* | KUVP 1424 (holotype), KUVP 8351, KUVP 9951 KUVP 33607, KUVP 33602 | In-person |
| *Planocephalosaurus robinsonae* | NHMUK R9953: R9976 (referred specimens) | In-person |
| *Procolophon trigoniceps* | BP/1/5127, BPI/1/4248, NHMUK R1726 (holotype), | In-person, CT |
| *Prolacerta broomi* | UCMP:V:37151, BPI/1/471, BPI/1/2675, BPI/1/2676, | In-person, Synchrotron µCT |

| | | |
|---|---|---|
| | BPI/1/4504a+b, BP/1/5066, BPI/1/5375, BPI/1/5880 (referred specimens) | |
| *Proterogyrinus scheelei* | USNM 22573, MCZ 4537 (referred specimen) | In-person |
| *Proterosuchus* spp. | SAM-PK-K591 (holotype of *P. fergusi*) SAM-PK-11208, BPI/1/3993, BPI/1/4016 (specimens referred to *Proterosuchus* spp. ) | In-person |
| *Protocaptorhinus pricei* | MCZ 1478 (holotype) | In-person |
| *Protorosaurus speneri* | USNM PAL 442453 (referred specimen) and descriptions of Gottman-Queseda and Sander (2009) | In-person |
| *Protorothyris archeri* | MCZ 1532 (holotype), MCZ 2147: 2150 (referred specimens) | In-person |
| *Raranimus dashankouensis* | IVPP V15424 (holotype) | CT |
| *Reiszorhinus olsoni* | FMNH UC 183 (holotype) | In-person |
| *Rhynchonkos stovalli* | FM-UR 1039 (holotype) | In-person, CT |
| *Romeria* spp. | MCZ 1963 (*R. prima* holotype) and MCZ 1480 (MCZ 1480, *R. texana* holotype) | In-person |
| *Rothianiscus* spp. | FMNH UR 713 (referred specimen) | In-person |
| *Saurodektes kitchingorum* | BP/1/3189, BP/1/1395 | In-person, CT, Synchrotron μCT |
| *Sauropareion anoplus* | SAM-PK-11192 (holotype), BP/1/5779, NMQR 3544, NMQR 3556, NMQR 3602 (referred specimens) | In-person, Synchrotron μCT |
| *Saurosternon bainii* | NHMUK 1234 holotype) | In-person |
| *Secodontosaurus obtusidens* | AMNH 4007 (holotype) | In-person |

| | | |
|---|---|---|
| *Seymouria* spp. | FMNH UC 1757, OMNH 79342, USNM V 21902, USNM V 15553 (referred specimens) | In-person, CT |
| *Sphenacodon* spp. | AMNH FARB 4779, AMNH FARB 4798, FMNH UC 10, FMNH UC 35, FMNH UC 736, and FMNH UC 1218 (referred specimens) | In-person |
| *Stereosternum tumidum** | AMNH FARB 2440, AMNH FARB 5748, AMNH FARB 6993, AMNH FARB 11002, AMNH FARB 23816 | In-person. We have kept the mesosaurid genera in this matrix as separate OTUs in this matrix, but acknowledge their hrecent synonymy by Piñeiro et al. 2021). |
| *Sumidadectes chozaensis* | FMNH UR 97 (holotype) | In-person |
| *Tanystropheus hydroides* | PIMUZ T 2790 | CT |
| *Tetraceratops insignis* | AMNH FARB 4526 (holotype) | In-person |
| *Thadeosaurus colcanapi* | MNHN 1908-i1-13a (holotype) | In-person. |
| *Thuringothyris mahlendorffae* | MNG 7729 (holotype), MNG 10183 (Referred specimen) | Based on descriptions by Boy and Martens (1991), Müller et al. (2006). |
| *Titanophoneus potens* | PIN 157/-1 (holotype) | In-person |
| *Trilophosaurus buettneri* | TMM 31025-244, MM 31025-4, TMM 31025-5, TMM 31025-68, TMM 31025-74, TMM 31025-80, TMM 31025-116, TMM 31025-125, TMM 31025-140, TMM 31025-142, TMM 31025-143, TMM 31025-144, TMM 31025-207, | In-person |
| *Tseajaia campi* | UCMP 59012 (holotype) | In-person |
| *Varanodon agilis* | FMNH UR 986 (holotype) | In-person |
| *Varanops brevirostris* | FMNH UC 644 (holotype) and FMNH UC 243, FMNH | In-person |

| | | |
|---|---|---|
| | UC 348, FMNH UC 349 (referred specimens) | |
| *Varanosaurus acutirostris* | AMNH FARB 4174 and FMNH PR 1670 (referred specimens) | In-person |
| *Vaughnictis smithae* | MCZ 2985 (holotype) | In-person, CT |
| *Watongia meieri* | UCMP 143278 (holotype) | In-person |
| *Weigeltisaurus jaekeli* | SSWG 113/7 (holotype) | Laminography scan |
| *Whatcheeria deltae* | FMNH PR 1700 (holotype), FMNH PR 1634, FMNH PR 1814 (referred specimens). | In-person, CT |
| *Youngina capensis* | AMNH 5561 (holotype), BPI/1/70, BPI/1/2459, BPI/1/2871, BPI/1/3859 . | In-person, CT, Synchrotron µCT |

**10. Supporting Video Captions.**

**Supporting Video 1.** Digital rendering of *Milleropsis* BP/1/720, with the three skulls oriented horizontally.

**Supporting Video 2.** Digital rendering of *Milleropsis* BP/1/720, with the three skulls oriented vertically.

**Supporting Video 3.** Digital rendering of the articulated skeleton of *Milleropsis* BP/1/4203.
